# Improving Predictability, Reliability and Generalisability of Brain-Wide Associations for Cognitive Abilities via Multimodal Stacking

**DOI:** 10.1101/2024.05.03.589404

**Authors:** Alina Tetereva, Annchen R. Knodt, Tracy R. Melzer, William van der Vliet, Bryn Gibson, Ahmad R. Hariri, Ethan T. Whitman, Jean Li, Farzane Lal Khakpoor, Jeremiah Deng, David Ireland, Sandhya Ramrakha, Narun Pat

## Abstract

Brain-wide association studies (BWASs) have attempted to relate cognitive abilities with brain phenotypes, but have been challenged by issues such as predictability, test-retest reliability, and cross-cohort generalisability. To tackle these challenges, we proposed a machine-learning “stacking” approach that draws information from whole-brain magnetic resonance imaging (MRI) across different modalities, from task-fMRI contrasts and functional connectivity during tasks and rest to structural measures, into one prediction model. We benchmarked the benefits of stacking, using the Human Connectome Projects: Young Adults (n=873, 22-35 years old) and Human Connectome Projects-Aging (n=504, 35-100 years old) and the Dunedin Multidisciplinary Health and Development Study (Dunedin Study, n=754, 45 years old). For predictability, stacked models led to out-of-sample *r*∼.5-.6 when predicting cognitive abilities at the time of scanning, primarily driven by task-fMRI contrasts. Notably, using the Dunedin Study, we were able to predict participants’ cognitive abilities at ages 7, 9, and 11 using their multimodal MRI at age 45, with an out-of-sample *r* of 0.52. For test-retest reliability, stacked models reached an excellent level of reliability (ICC>.75), even when we stacked only task-fMRI contrasts together. For generalisability, a stacked model with non-task MRI built from one dataset significantly predicted cognitive abilities in other datasets. Altogether, stacking is a viable approach to undertake the three challenges of BWAS for cognitive abilities.

**Significance statement:** Scientists have had limited success in predicting cognitive abilities from brain MRI. We proposed a machine learning method, called stacking, to draw information across different types of brain MRI. Using three large databases (n=2,131, 22–100 years old), we found stacking to make the prediction of cognitive abilities 1) closer to actual cognitive scores when applied to a new individual, not part of the modelling process, 2) reliable over times and 3) applicable to the data collected from different age groups and MRI scanners. Indeed, stacking, especially with fMRI task contrasts, allowed us to use MRI of people aged 45 to predict their childhood cognitive abilities reasonably well. Accordingly, stacking may help MRI realise its potential to predict cognitive abilities.

## Introduction

Individual differences in cognitive abilities are stable across the lifespan(1) and have relatively high heritability(2). They are key indicators of educational achievements(3), career successes(4), well-being(5), socioeconomic stability(6) and health outcomes(7). Recent studies have also demonstrated a widespread relationship between impairments in cognitive abilities and various psychopathological disorders(8, 9). Accordingly, relating individual differences in cognitive abilities to neuroimaging data has been a primary goal for cognitive neuroscientists, from both basic and applied science perspectives(10). This approach allows neuroscientists to scientifically quantify the presence of information related to cognitive abilities from each neuroimaging type or modality. It also paves the way for identifying neural indicators of cognitive abilities, which could be useful for understanding the etiology of neuro- and psychopathology(11). Indeed, a leading transdiagnostic framework for psychiatry, the Research Domain Criteria (RDoC), treats cognitive abilities as one of the main functional domains for psychopathology across diagnoses. Having a robust neural indicator of cognitive abilities, in addition to behavioral and genetic indicators, is central to the RDoC framework(12).

The availability of large-scale neuroimaging databases(13) and the accessibility of predictive modelling methodologies(11, 14) have provided encouraging avenues to pursue a neural indicator of cognitive abilities. Accordingly, several researchers have built prediction models to predict cognitive abilities from brain magnetic resonance imaging (MRI) signals and evaluated the models’ performance on separate, unseen data in the so-called Brain-Wide Association Studies (BWAS)(11, 15). BWAS can be conducted using either univariate (also known as mass-univariate) or multivariate (also known as machine learning) methods to draw MRI information. While univariate methods draw data from one region/voxel at a time, multivariate methods draw MRI information across regions/voxels (11, 16, 17). These multivariate methods, from particular MRI modalities, appear to boost predictability for cognitive abilities (11, 16–18). For examples, Marek and colleagues(16) conducted BWAS on several large datasets can concluded, “More robust BWAS effects were detected for functional MRI (versus structural), cognitive tests (versus mental health questionnaires) and multivariate methods (versus univariate).”

Akin to Genome-Wide Association Studies (GWAS) in genetics(19) that can integrate information across SNPs across the genome to create a predicted, propensity score for a phenotype of interest (e.g., cognitive abilities), known as polygenic scores, BWAS can also be used to create a similar predicted score for each individual based on his/her neuroimaging data. For instance, Marek and colleagues(16) used trained multivariate methods to predict cognitive abilities from brain MRI data using part of the data (known as training set) and applied the trained model to the unseen participants (known as test set). Participants in the test set then had a predicted score of their cognitive abilities, based on their brain MRI data. Yet, BWAS for cognitive abilities has faced several challenges, including but not limited to predictability, test-retest reliability and generalisability, as detailed below(16, 20, 21). These challenges have led to headlines, such as “Cognitive Neuroscience at the Crossroads”(22) and “Scanning the Brain to Predict Behavior, a Daunting ‘Task’ for MRI”(23). To address these issues, we(24, 25) have recently proposed a potential solution, “stacking”(26), which allows us to combine different modalities of MRI into one prediction model. In this study, we aim to formally benchmark the benefits of stacking in improving predictability, test-retest reliability and generalisability, using three large-scale neuroimaging databases(27–29).

First, predictability, or out-of-sample prediction, pertains to the ability of prediction models to predict a target variable, e.g., cognitive abilities, based on features, e.g., functional MRI (fMRI) data, of unseen participants, not part of the model-building processes(30). More specifically, we refer to an application of a validation within one dataset. Here researchers usually take a relatively large dataset, split it into training and test sets, then build a model from the training set and apply the model to the test set. In addition to doing one split, researchers could also apply a cross-validation strategy by splitting a dataset into different non-overlapping training-test folds and looping through folds to calculate the average performance across the test sets(31, 32). Several earlier studies(33–35) did not apply any validation when predicting cognitive abilities from MRI, possibly causing inflated predictability(16). With proper data splitting, a recent meta-analysis(15) estimated the predictability of multivariate methods on brain MRI of different modalities with a validation for cognitive abilities to be a Pearson’s correlation *r* of 0.42 on average.

While this level of predictability is encouraging, there is still room for improvement. Given that different MRI modalities may convey different information about the brain, drawing information across different MRI modalities could allow us to improve predictability further. Stacking enables researchers to draw information across MRI modalities, which seems to improve predictability over relying on any single MRI modality (24–26, 36, 37). In this framework (see Figure 1), researchers first build ‘non-stacked’ prediction models separately for each MRI set of features (e.g., cortical thickness or cortical area), and computed predicted values from each of these non-stacked’ models. They then treat these predicted values as features for ‘stacked’ prediction models, allowing them to draw information across different MRI sets of features. Still, most studies use one single type of MRI to build prediction models. The popular choices include resting-state fMRI functional connectivity (Rest FC, or correlations in blood-oxygen-level-dependent (BOLD) time series across areas during rest)(38–40), task fMRI functional connectivity (Task FC, or correlations in BOLD time series across brain regions during each task)(41–48) and structural MRI (including measures such as thickness, area and volume in cortical/subcortical areas)(49). While it is less common to use task-fMRI contrasts (Task Contrasts, or fMRI BOLD activity relevant to events in each task) to predict cognitive abilities, studies have started to show the superior predictability of contrasts from certain tasks, compared to other MRI modalities(24, 25, 39, 50). Nonetheless, previous attempts at stacking often ignored task contrasts (36, 37), and as a result, while improving over non-stacked models, they have not led to satisfactory predictive performance. We(24, 25) have started to show a boost in predictability when applying stacking to combine task contrasts with other MRI modalities. Here, to ensure the robustness of this approach, we examined the benefits of stacking task contrasts, along with other MRI modalities, on multiple large-scale datasets.

**Figure 1.**
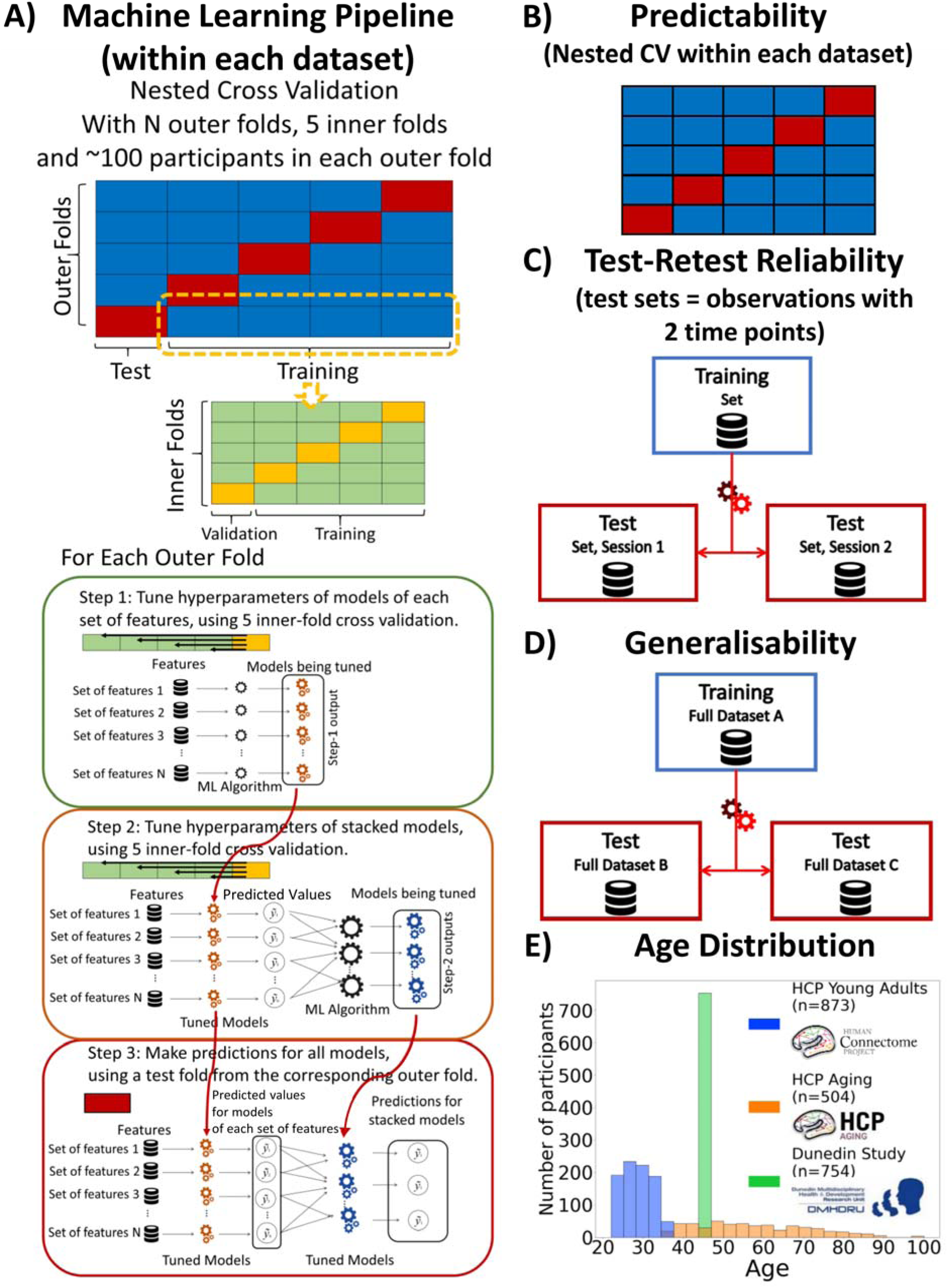
Overview of Study Methodology. We used three datasets: Human Connectome Project Young Adults (HCP Young Adults), Human Connectome Project Aging (HCP Aging) and Dunedin Multidisciplinary Health and Development Study (Dunedin Study). A) Machine Learning Pipeline. Here, we depict the process we used for building prediction models for testing predictability within each dataset. Briefly, we used nested cross-validation (CV) by splitting the data into outer folds with around 100 participants in each. In each outer-fold CV loop, we then treated one of the outer folds as an outer-fold test set and treated the rest as an outer-fold training set. We then divided each outer-fold training set into five inner folds and applied inner-fold CV to build prediction models in three steps. In the first step (known as a non-stacking layer), one of the inner folds was treated as an inner-fold validation set, and the rest was treated as an inner-fold training set in each inner-fold CV. We used grid search to tune prediction models for each set of features. In the second step (known as a stacking layer), we treated different combinations of the predicted values from separate sets of features as features to predict the cognitive abilities in separate “stacked” models. In the third step, we applied the already tuned models from the first and second steps to the outer-fold test set. B) Predictability. Here, we examined the predictive performance across outer-fold test sets within each dataset. C) Test-Retest Reliability. Here, we used HCP Young Adults and Dunedin Study and treated participants who were scanned twice across MRI sessions as the test set and the rest as the training set. We then examined the intraclass correlation (ICC) of the predicted values in the test set between the first and second MRI sessions. D) Generalisability. Here, we examined the predictive performance of the models built from a different dataset. We treated one of the three datasets as a training set and the other two as two separate test sets. E) Age Distribution. Here, we show the age of participants at the time of scanning in each dataset.

Second, test-retest reliability pertains to the rank stability of measurements across different time points, assuming the absence of significant changes between assessments (e.g., treatment exposure, injury and/or disease progression)(48–50)(51, 52). For instance, if some people score higher than their peers at time one, they should also score higher than their peers at time two. To use prediction models as an indicator for individual differences in cognitive abilities, the predicted values should be reliable across time. A recent study challenged the test-retest reliability of task contrasts(20). Here the researchers examined test-retest reliability of Task Contrasts in certain areas, known to be strongly elicited in each task, across two-time points and found a poor level of test-retest reliability across different tasks and two datasets: the Human Connectome Project Young Adult (HCP Young Adults)(29), and the Dunedin Multidisciplinary Health and Development Study (Dunedin Study)(28). This poor level of test-retest reliability from Task Contrasts is concerning, especially when compared to the higher levels of test-retest reliability found from structural MRI, Rest FC and Task FC(20, 53, 54). In fact, structural MRI provided reliability that was almost at the ceiling(20). Yet these studies simply took task contrasts from certain areas; they did not create prediction models or use multivariate methods and stacking to draw information across the whole brain and across different tasks/MRI modalities. It is possible that Task Contrasts could be more reliable once the models consider information across the whole brain, across different tasks and across different MRI modalities. Following this conjecture, we(25) recently showed that multivariate methods and stacking substantively boosted reliability, reaching a much higher level of reliability in HCP Young Adults(29). To ensure the robustness of our findings, we need to test the benefits of this stacking approach in another, independent dataset: the Dunedin Study(28), for example.

Third, generalisability, or more specifically cross-cohort generalisability, pertains to the ability of prediction models built from one dataset to predict the cognitive abilities of participants of another dataset(21). Different datasets, for instance, use different MRI scanners, recruit participants from different cultures and age groups, or implement different cognitive-ability measurements. Thus, while predictability within one dataset provides the performance of prediction models within specific, harmonised contexts of one dataset, generalisability across datasets allows us to gauge the performance of prediction models in broader contexts. This means that generalisability situates closer to how deployable the prediction models are in indicating cognitive abilities in the real world(21). Yet, only a few studies have investigated the generalisability of MRI prediction models for cognitive abilities, and most have focused on functional connectivity during rest and/or tasks(55–57). The generalisability of stacked models is currently unknown.

Our overarching goal is to benchmark the impact of stacking on predictability, test-retest reliability and generalisability of MRI prediction models for cognitive abilities. To achieve this, we used three large-scale neuroimaging databases, including HCP Young Adults(29), Human Connectome Project Aging (HCP-Aging)(27) and Dunedin Study(28). The databases vary in various aspects, such as participants’ age (see Figure 1) and cultures, physical scanners, scanning parameters and cognitive-ability assessments. Note that, unlike our previous implementation of stacking(25), we also included Task FC in addition to Task Contrasts to capture wider information during task scanning. Specifically, here we built stacked models from eight different combinations of functional and structural MRI sets of features: “Task Contrast” including Task Contrasts from all of the tasks, “Task FC” including Task FC from all of the tasks, “Non Task” including Rest FC and structural MRI, “Task Contrast & FC” including task contrasts and task FC from all of the tasks, “All” including all sets of features, “All excluding Task Contrast” including every set of features except Task Contrasts, “All, excluding Task FC” including every set of features except Task FC, “Resting and Task FC” including FC during rest and tasks.

For predictability (see Figure 1), we applied nested cross-validation (CV) within each dataset to evaluate the predictability of stacked models from multimodal MRI. To build the stacked models, we applied 16 combinations of multivariate predictive-modelling algorithms (including Elastic Net(58), Support Vector Regression(59), Random Forest(60) and XGBoost(61)). Moreover, the nature of the Dunedin Study’s longitudinal measurements(28) allowed a unique opportunity for us to predict cognitive abilities from MRI data at the time of scanning (at the age of 45 years old), but also at much earlier times (at the age of 7, 9 and 11 years old), as well as to predict the residual scores that reflect relative changes in cognitive abilities during 36 years, compared to participants’ peers(62). For test-retest reliability (see Figure 1), we examined the rank stability of stacked models from participants who were scanned twice in HCP Young Adults and Dunedin Study. Lastly, for generalisability (see Figure 1), we built stacked models from one dataset and evaluated their performance on the other two. Due to the different tasks used in different datasets, we unfortunately could only examine the generalisability of the “Stacked: Non Task” model, which combined all MRI modalities that did not involve tasks.

## Results

### 1. Predictability

We showed performance indices of stacked and non-stacked models for each predictive-modelling algorithm and dataset in Figure 2 and Figures S1-9 and their bootstrapped 95% CI in Figures S10-18. Overall, when predicting cognitive abilities at the time of scanning, the prediction models from multimodal MRI across different predictive-modelling algorithms varied in their performance, reflected by Pearson’s correlation (*r*) between predicted and observed values, ranging from around 0 to .6, across the three datasets. Notably, combining different sets of MRI features into stacked models constantly led to higher predictive performance. “Stacked: All”, which included all sets of MRI features, gave rise to top-performing models across algorithms and the three datasets. Additionally, using Elastic Net across both non-stacking and stacking layers regularly resulted in prediction models that were either equally good or better than other prediction models based on other algorithms (see the bootstrapped differences in Figure S19-28). For instance, using Elastic Net across both layers for “Stacked: All” led to *r* at mean*(M*)=.60 (95%CI[.56, .64]), *M*=.61 (95%CI[.56, .66]) and *M*=.55 (95%CI[.49, .60]) for HCP Young Adults, HCP Aging and Dunedin Study, respectively.

**Figure 2.**
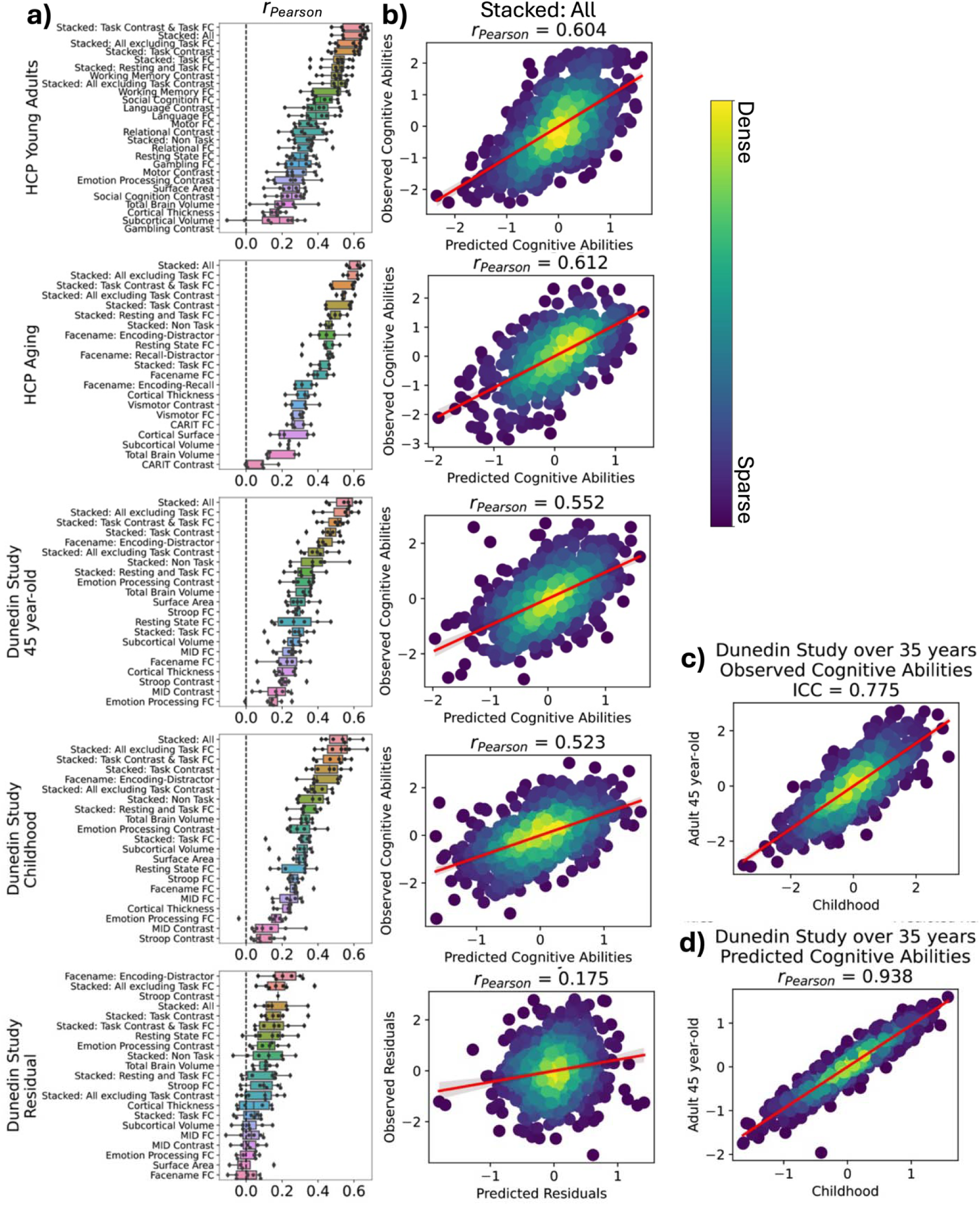
Predictability of stacked and non-stacked models. **a)** Pearson’s correlation, r) of stacked and non-stacked models for each dataset with Elastic Net across the two layers. Higher is better. Each dot represents predictive performance at each outer-fold test set. For other algorithms and other performance indices, the coefficient of determination (R^2^) and mean absolute error (MAE), see Figures S1-S9. For Dunedin Study, childhood scores reflect cognitive abilities, averaged across 7, 9 and 11 years old, and negative residual scores reflect a stronger decline in cognitive abilities, as expected from childhood cognitive abilities, compared to participants’ peers. **b)** Dense scatter plot illustrating observed and predicted cognitive abilities (Z scores) using Stacked-All models with Elastic Net across two layers. Stacked All include all sets of MRI features. **c)** presents observed cognitive abilities at ages 7, 9, and 11 compared to age 45 from the Dunedin Study. The interclass correlation (ICC) reflects the strength of the relationship in the observed cognitive-ability scores between these time points. **d)** displays predicted cognitive abilities at ages 7, 9, and 11 compared to age 45 from the Dunedin Study. Pearson’s correlation reflects the strength of the relationship in the predicted cognitive-ability scores between these time points. The predicted cognitive-ability scores at each of the two time points were trained from the same set of neuroimaging features via the Stacked-All models, albeit with different targets (either the cognitive abilities averaged across ages 7, 9, and 11, or cognitive abilities at age 45). This is because MRI data were only collected at age 45, while cognitive abilities were collected at both time points. Accordingly, it is expected that the ICC of the observed cognitive-ability scores will be higher than the Pearson’s correlation of the predicted cognitive-ability scores. SVR = Support Vector Regression; XGB = XGBoost; FC = Functional Connectivity. ICC=Interclass Correlation.

Among the non-stacked models that predicted cognitive abilities at the time of scanning, we found varied predictive performance associated with different sets of MRI features. On the one hand, Task Contrasts from certain tasks led to top-performing models across the three datasets: the working memory task in HCP Young Adults and the facename task in HCP Aging and Dunedin Study. With Elastic Net, these three task contrasts led to *r* at *M*=.5 (95%CI[.45, .59]) for HCP Young Adults, *M*=.46 (95%CI[.39, .51]) for HCP Aging, *M*=.43 (95%CI[.37, .49]) for Dunedin Study. On the other hand, Task Contrasts from some other tasks led to poor-performing models, such as the gambling task in HCP Young Adults (*r* could not be calculated due to the models resulting in the same predicted values on certain folds), the Conditioned Approach Response Inhibition Task (CARIT) task in HCP Aging, *r* at *M* =.07 (95%CI[-.02, .15]), and the monetary incentive delay (MID) task in Dunedin Study, *r* at *M*=.16 (95%CI[.08, .22]).

For Dunedin Study, the prediction models that predicted cognitive abilities at the time of scanning (i.e., when participants were 45 years old) performed similarly to those that predicted cognitive abilities, collected much earlier than the scanning time (i.e., when they were 7, 9 and 11 years old). For instance, the “Stacked: All” models using Elastic Net across both non-stacking and stacking layers predicted cognitive abilities at 45 years old and at 7, 9 and 11 years old with *r* at *M*=.55 (95%CI [.49, .60]) and *M*=.52 (95%CI [.47, .57]), respectively. And Task Contrasts from the facename task led to top-performing models across two time points: with *r* at *M*=.43 (95%CI[.36, .48]) at 7, 9 and 11 years old, compared to *M*=.43 (95%CI[.37, .49]) at 45 years old.

In contrast, the performance of models predicting the residual scores for cognitive abilities from multimodal MRI was much poorer. Note that the negative residual scores reflect a stronger decline in cognitive abilities, as expected from childhood cognitive abilities, compared to participants’ peers. The highest-performing model predicting the residual scores across algorithms was the model with the encoding vs distractor contrast from the facename task, followed by various stacked models. With Elastic Net, the best model that predicted the residual scores led to *r* at *M*=.21 (95%CI[.14, .27]). And with Elastic Net across both layers for “Stacked: All” led to *r* at *M*=.17 (95%CI[.10, .24]). While these *r* levels were statistically better than chance according to bootstrapping (see Figure S10), it was much lower than those from prediction models that predicted cognitive abilities at the time of scanning or much younger age.

To understand the contribution of each MRI feature, we examined feature importance of each model based on Elastic Net coefficients. Given that Elastic Net features are linear and additive, the linear combination of Elastic Net coefficients reflects how the algorithm makes prediction. For stacked models, Figure S29 shows the feature importance of stacked models for each dataset when predicting cognitive abilities at the time of scanning. The top-performing Task Contrasts contributed stronger in the stacked models across the three datasets. For non-stacked models, S30-S31 shows the feature importance of for each MRI modality, study and target variable. Figure 3 and Table S1-10 show the feature importance of the top-performing, non-stacked models for each study and target variable.

**Figure 3.**
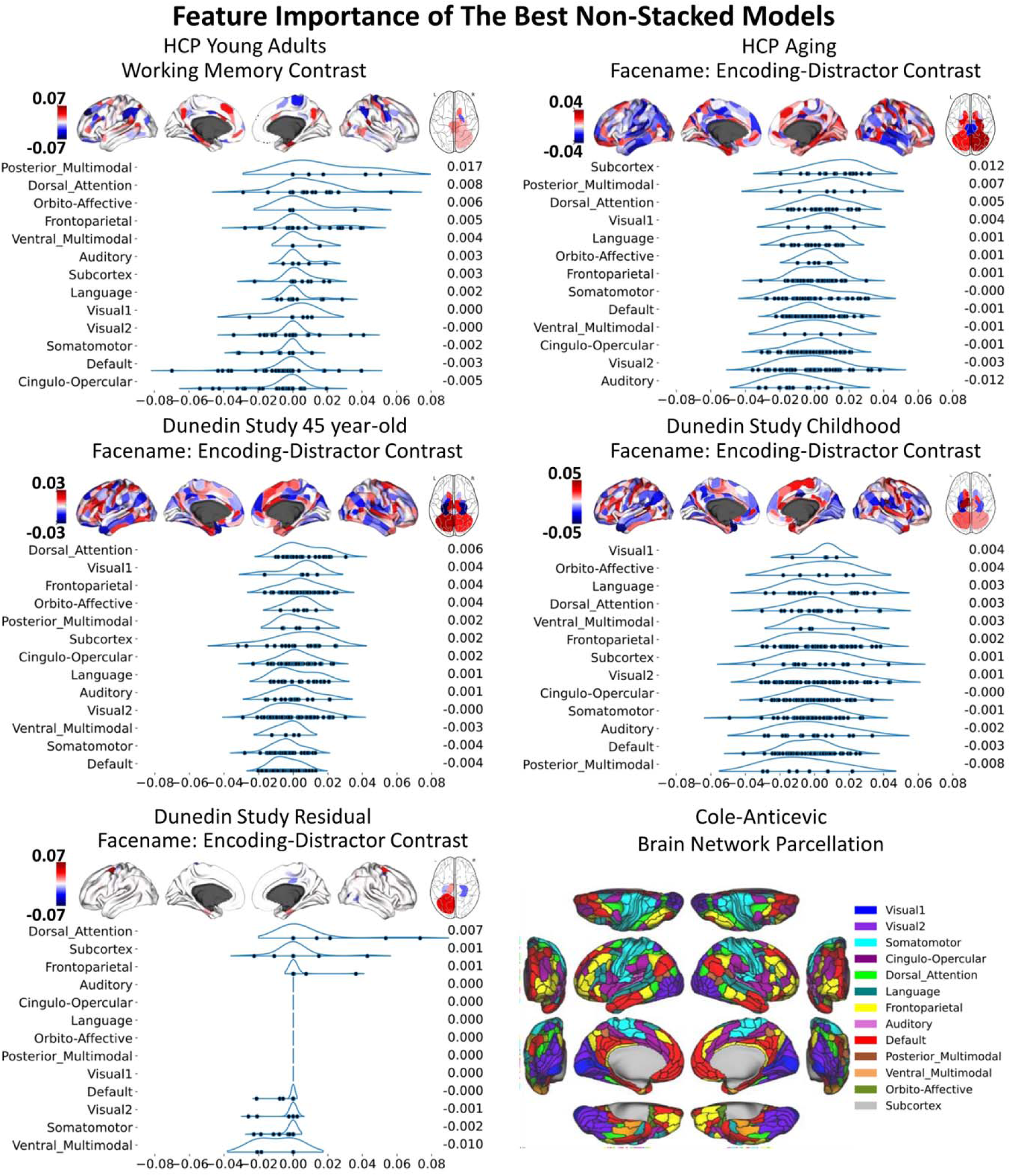
Feature importance of the top-performing non-stacked models with with Elastic Net, as indicated by Elastic Net coefficients. We grouped brain ROIs from the Glasser atlas (67) into 13 networks based on the Cole-Anticevic brain networks (66). In each figure, the networks are ranked by the mean Elastic Net coefficients, with the rankings shown to the right of each figure. The network partition illustration is sourced from the Actflow Toolbox https://colelab.github.io/ActflowToolbox/. We provid actual values of the feature importance in Table S1-10.

Note that we provided tables of the numerical values of the predictability indices on our github page: https://github.com/HAM-lab-Otago-University/Predictability-Reliability-Generalizability/tree/main/4_Supplementary.

### 2. Test-Retest Reliability

Figure 4 shows the test-retest reliability. Here we tested the rank stability of predicted values from prediction models across two sessions, as indicated by interclass correlation (ICC). Given the availability of the test-retest participants, we examined the test-retest reliability of the stacked and non-stacked models from HCP Young Adults and Dunedin Study. We provided predicted values across two scanning sessions for each participant in Figures S33-S34 and ICC for each MRI feature before prediction modelling in Figures S35-S38. Overall, for both datasets, prediction models with structural MRI, including total brain and subcortical volume, surface area and cortical thickness, led to the highest level of test-retest reliability.

**Figure 4.**
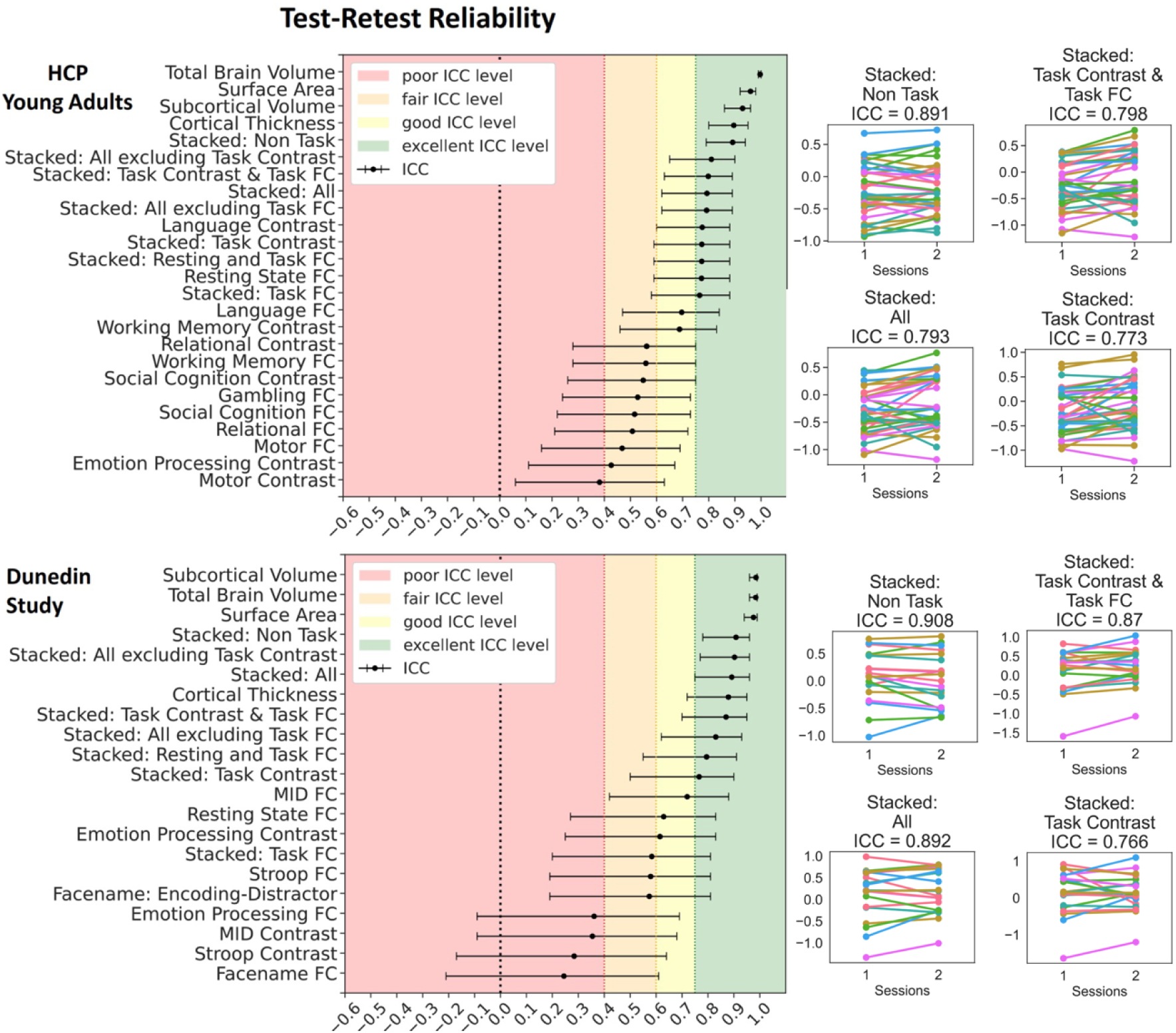
Test-retest reliability of the predicted values of the stacked and non-stacked models, indicated by Interclass Correlation (ICC) for HCP Young Adults and Dunedin Study. **Left panel:** Each dot represents ICC, while each bar represents a 95% Confidence Interval. **Right panel**: Predicted values of some stacked models across two scanning sessions. Each line represents each participant. Lines would be completely parallel with each other in the case of perfect test-retest reliability. For other stacked and non-stacked models, see Figures S33-S34.

Similar to predictability, combining different sets of MRI features into stacked models mostly gave rise to high test-retest reliability. “Stacked: All”, which included all sets of MRI features, resulted in an excellent ICC at .79 and .89 for HCP Young Adults and Dunedin Study, respectively. Moreover, we also found the boosting effect of stacking even when only fMRI during tasks was included in the models. For instance, “Stacked: Task Contrast and Task FC”, which included the contrasts and functional connectivity (FC) from all of the fMRI tasks within each dataset, led to an excellent ICC at .8 and .87 for HCP Young Adults and Dunedin Study, respectively. Similarly, “Stacked: Task Contrast”, which included the contrasts, but not functional connectivity (FC), from all of the fMRI tasks within each dataset, still led to an excellent ICC at .77 and .77 for HCP Young Adults and Dunedin Study, respectively.

Among the non-stacked models that predicted cognitive abilities from fMRI (including Task Contrasts, Task FC and Rest FC), some models showed a good-to-excellent level of ICC. These include a contrast from the language task (ICC=.77) and FC during rest (ICC=.77) in HCP Young Adults and FC during the monetary incentive delay (MID) task (ICC=.72) and rest (ICC=.63) in Dunedin Study. Yet, some models from fMRI provided a poor level of ICC, including a contrast during the motor task (ICC=.38) in HCP Young Adults and FC, a contrast during the MID (ICC=.35) and Stroop (ICC=.28) tasks and FC (ICC=.24) during the emotion processing and facename tasks in Dunedin Study.

### 3. Generalisability

Figure 5 shows the generalisability among the three datasets. Here we tested the performance of the prediction models trained from one dataset in predicting the cognitive abilities of participants from another separate dataset, as indicated by Pearson’s correlation (*r*) between predicted and observed cognitive abilities. Given the different fMRI tasks used in different datasets, we only examined the generalisability of the prediction models using non-task sets of features (including rest FC, cortical thickness, cortical surface area, subcortical volume, total brain volume and their combination, or “Stacked: Non Task”). Note that even the fMRI tasks with the same name, “the facename task” and “face or emotion processing” tasks were implemented differently across different datasets (see Methods).

**Figure 5.**
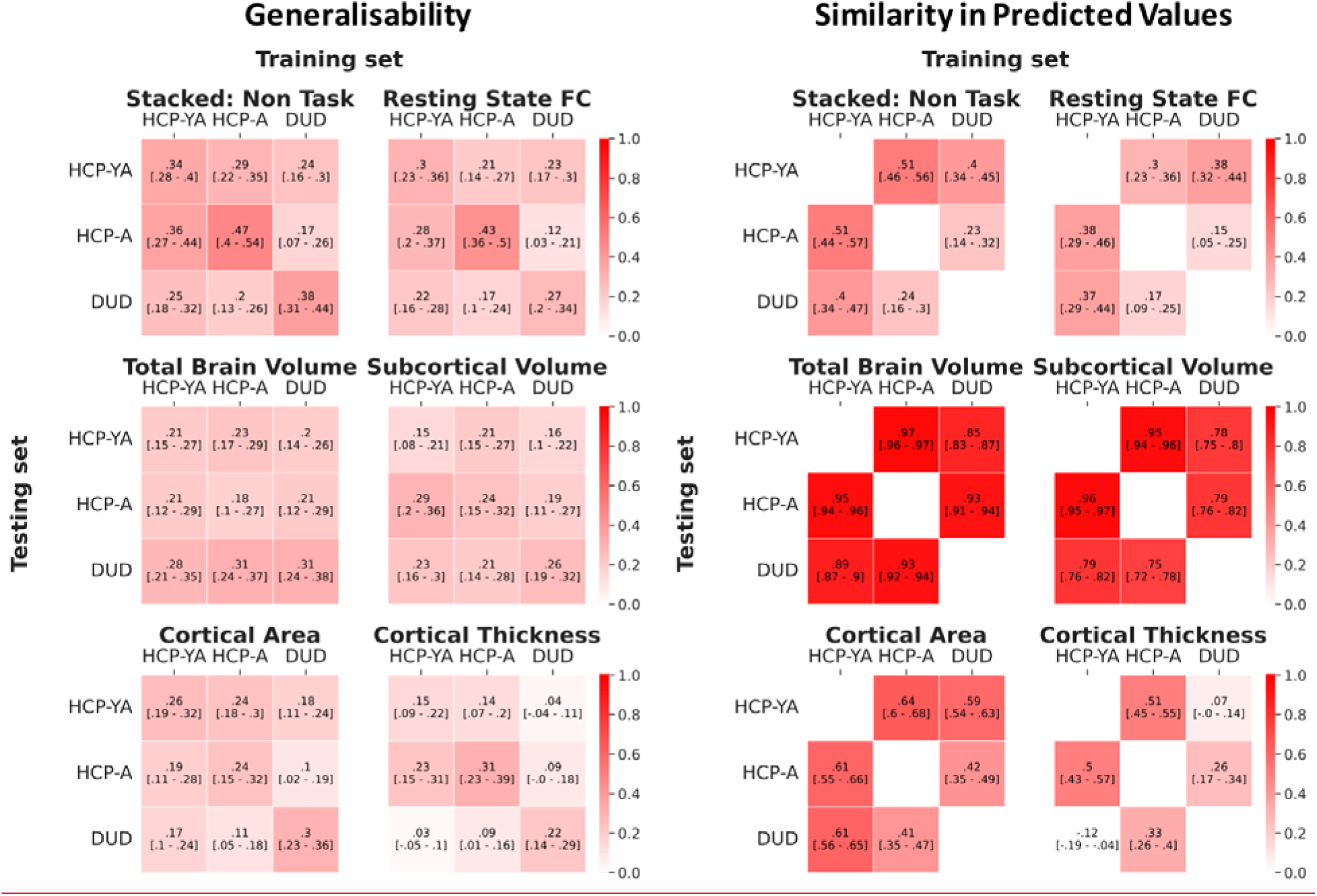
Generalisability and similarity in predicted values among the three datasets, as indicated by Pearson’s correlation, r. Note that due to the different tasks used in different datasets, we only examined the generalisability of prediction models built from non-task sets of features (including rest FC, cortical thickness, cortical surface area, subcortical volume, total brain volume and their combination, or “Stacked: Non Task”). For generalisability, the off-diagonal values reflect the level of generalisability from one dataset to another, while the diagonal values reflect the predictability of the models built from the same dataset via nested cross-validation (CV). For the similarity in predicted values, the off-diagonal values reflect the level of similarity in predicted values between two datasets. Higher values are better. The values in square blankets reflect a bootstrapped 95% Confidence Interval (CI). If 95% CI did not include zero, then generalisability/similarity in predictive values was better than chance. HCP-YA = HCP Young Adults; HCP-A = HCP Aging; DUD = Dunedin Study.

The “Stacked: Non Task” model showed generalisability at *M*=.25 (*SD*=.06). This level of cross-dataset generalisability was significantly better than chance (see the 95%CI in Figure 5) and was similar to, albeit numerically smaller than, the within-dataset predictability of the models built from nested cross-validation (CV) (*M*=.4*, SD*=.05). There were some differences in generalisability among pairs of studies. Generalisability was numerically higher between HCP Young Adults and HCP Aging (*M*=.33*, SD*=.04), as compared to between Dunedin Study and the other two datasets (*M*=.22*, SD*=.03).

Similarly, the non-stacked models with non-task sets of features showed generalisability at *M*=.18 (*SD*=.05). Apart from cortical thickness, the generalisability of every other non-task set of features was significantly better than chance (see the 95% CI in Figure 5). Additionally, this level of cross-dataset generalisability from the non-task sets of features was similar to the within-dataset predictability of the models built from nested CV (*M*=.25*, SD*=.05).

We also examined the similarity in predicted values among the three datasets. Here we tested the Pearson’s correlation (*r*) in predicted values between the prediction models built from the same dataset and those built from another dataset (Figure 5). The “Stacked: Non Task” model showed similarity in predicted values at *M*=.38 (*SD*= .12). This level of similarity was significantly better than chance (see the 95%CI in Figure 5). As for the non-stacked models with non-task sets of features, we found the similarity in predicted values on average at *M*=.57 (*SD*=.11) and all of them were significantly better than chance (see the 95%CI in Figure 5). Yet, some non-task sets of features appeared to be stronger in similarity than others. For instance, the similarity in predicted values for the total brain volume (*M*=.92, *SD*=.04) and subcortical volume (*M*=.84, *SD*=.09) were numerically higher than those for rest FC (*M*= .29, *SD*= .10), cortical area (*M*= .55, *SD*= .09) and cortical thickness (*M*= .26, *SD*= .22).

## Discussion

Here, we examined stacking as a potential solution for improving BWAS for cognitive abilities in three key aspects: predictability, test-retest reliability and generalisability(16, 20, 21). Most BWASs use one single modality of MRI to build prediction models, but here, we drew information across different MRI modalities via stacking(26). Stacked models demonstrated improvement in all three aspects and performed better than any individual modality in isolation. For predictability, stacked models led to high predictability, relative to what has been reported in the literature, across the three datasets when predicting cognitive abilities at the time of scanning. Notably, using the Dunedin Study, we were able to predict participants’ cognitive abilities at ages 7, 9, and 11 using their multimodal MRI at age 45, relatively well (*r*=.52). We found this predictive performance was driven by task contrasts, followed by task connectivity, across the three datasets. For test-retest reliability, stacked models reached an excellent level of reliability across HCP Young Adults and Dunedin Study, even when we only included fMRI during tasks in the models. For generalisability, combining non-task MRI features into a stacked model led to models that were applicable to other datasets, giving a level of performance that is better than chance. Altogether, the results optimistically support stacking as a viable approach to address the three challenges of BWAS for cognitive abilities.

### Stacking Improved Predictability

Combining MRI across different sets of features via stacking consistently and substantially improved predictability within each dataset. Indeed, we found this improved predictability across three large-scale datasets that varied in age, culture, scanner manufacturer, scanning parameters and cognitive-ability assessments(27–29). This is consistent with our previous findings(24, 25). For cognitive abilities at the time of scanning, stacked models with all MRI sets of features led to *r* up to around .6, higher than those of non-stacked models in the current study, as well as those reported in a recent meta-analysis (*r*=.42 with CI_95%_=[0.35,0.50])(15). As reviewed in the meta-analysis (15), most studies applied similar machine learning algorithms to those used here but relied on a single MRI modality, most commonly functional connectivity (FC) during rest or task, followed by structural MRI. The use of task-fMRI contrasts is less common(39). Our non-stacked models with task FC, rest FC, and sMRI showed similar performance to previous findings. Importantly, our stacked models that included Task Contrasts along with other modalities showed a much higher predictive performance (e.g., *r*=.604, R^2^=. 352 based on “Stacked: All” with Elastic Net across both layers done on the HCP Young Adults), compared to the stacked models that did not include Task fMRI in the current study as well as to the models in a previous study that also modelled the data from HCP Young Adults (R^2^ = .078)(37). This confirms a) that different MRI sets of features provide independent but complementary information about individual differences in cognitive abilities and b) that Task Contrasts, which are often ignored, could significantly help improve the predictive performance.

The superior predictability of Task Contrasts from certain tasks, especially when predicting cognitive abilities at the time of scanning, was also consistent with previous work(24, 25, 39, 50). The working-memory task in HCP Young Adults and the facename task in HCP Aging and Dunedin Study created non-stacked models with the highest predictability for each dataset. The more popular MRI modalities, Task FC and Rest FC(15), did not perform as strongly as the Task Contrasts from working-memory and facename tasks. And structural MRI seemed to provide much poorer predictability across datasets, consistent with earlier work(49). It is important to note that not all Task Contrasts produced prediction models with high predictability. In fact, the worst prediction models across the three studies were also Task Contrasts (e.g., the gambling, CARIT and MID tasks in HCP Young Adults, HCP Aging and Dunedin Study, respectively). This suggests the selectivity of the fMRI BOLD activity relevant to events for different tasks--some tasks were related to individual differences in cognitive abilities and some tasks were not. The best tasks here were related to either working memory or episodic memory, which might reflect what was being measured with the cognitive-ability assessments(17, 63), i.e., through NIH Toolbox(64) or Wechsler Adult Intelligence Scale (WAIS)(65). Accordingly, to further improve the predictability of BWAS for cognitive abilities via task contrasts, future research will need to determine which tasks are more relevant to cognitive abilities. If the tasks used have not yet been established to predict cognitive abilities, researchers may still consider stacking to determine if combining Task Contrasts from such tasks with other MRI modalities improves predictability. By using multivariate predictive modeling algorithms, such as Elastic Net, to stack different Task Contrasts, tasks that do not enhance predictability will have less weight in the final stacked model due to regularization. This makes it safe to include any available tasks. Researchers may then decide to drop the less contributing tasks to create a more parsimonious model.

We also examined the predictability of stacked models in light of the Dunedin Study’s longitudinal measurements for cognitive abilities(28). Stacked models with all MRI sets of features were able to predict cognitive abilities, collected 36 years before the scanning time, at a similarly high level of performance to those at the time of scanning (*r*=.55 vs. *r*=.52, respectively). Yet, when predicting the residual scores, the stacked models with all MRI sets of features gave much lower predictability, albeit still significant, at *r*=.17. These residual scores reflect changes in cognitive abilities from childhood to middle age, compared to participants’ peers. This pattern of results may suggest that brain information revealed by multimodal MRI, obtained in the middle age (i.e., 45 years old), is more related to the stable trait of cognitive abilities, but less to the changes over 35 years. Perhaps this is because individual differences in cognitive abilities were stable over the lifespan(1), making it easier for multimodal MRI to capture their inter-variability over intra-individual variability. Using the Dunedin Study, we indeed found a high rank stability of this trait: that childhood cognitive abilities were related to middle-aged cognitive abilities at ICC=.78, and that the multimodal MRI predicted values of cognitive abilities based on either time points led to very similar scores at *r*=.94 (see Figure 2c and 2d). Accordingly, if the aim of BWAS is to capture the stable trait of cognitive abilities, the current approach of stacking multimodal MRI data from one time point seems appropriate. It is important to note that our work only demonstrates the retrospective prediction of cognitive abilities (i.e., using MRI at age 45 to predict cognitive abilities 36 years prior). Future work is needed to examine if our method can be extended to forecast cognitive abilities. Fortunately, the Dunedin Study is still ongoing, and we hope to test if our method can use MRI at age 45 to predict future cognitive decline as well as cognitive-related neurological disorders, such as mild cognitive impairments, dementia, and Alzheimer’s, as the participants age.

To explain how each model made predictions, we treated Elastic Net coefficients as indicators of feature importance. Examining the feature importance in the stacked models revealed that the top-performing modality, specifically the best Task Contrasts, was the strongest contributor. This pattern was consistent across datasets. This suggests that the top-performing Task Contrasts, such as the working-memory task (reflected by high over lower working-memory load conditions) in HCP Young Adults and the facename task (reflected by encoding over distractor/control conditions) in HCP Aging and the Dunedin Study, provided strong and unique contributions to the overall prediction of the stacked models. Examining the feature importance of these top-performing Task Contrasts illustrates the contribution from each brain area.

We grouped brain areas into 13 different networks based on the Cole-Anticevic definition (66). Contributions from different brain networks varied depending on the specific Task Contrasts, datasets, and cognitive target variables. Nonetheless, some patterns emerged. For instance, our prediction models indicated that participants with more positive Task Contrasts from brain areas within the default mode network tended to have worse cognitive abilities. Conversely, participants with more positive Task Contrasts from brain regions within the dorsal attention network tended to have better cognitive abilities at the time of scanning, cognitive abilities 35 years prior, and residual scores. Accordingly, we demonstrated the contribution of certain networks within the context of specific tasks in predicting cognitive abilities.

### Stacking Improved Test-Retest Reliability

Creating prediction models from separate MRI sets of features and combining them via stacking also improved reliability for Dunedin Study(28), especially for Task Contrasts, similar to our previous findings(25) with HCP Young Adults(29). This approach, in effect, addresses the poor reliability of Task Contrasts, found earlier in the same two datasets(20). That is, previous work(20) focused on the reliability of Task Contrasts at specific brain areas from certain tasks and found low test-retest reliability (also demonstrated here in Figure S35-36). Here instead of focusing on specific areas, we used multivariate methods and stacking to draw information across the whole brain and tasks and found a boost in reliability. Indeed, stacked models that combined only Task Contrasts and that combined Task Contrasts and FC together both gave the ICC at excellent levels (i.e., ICC≥.75) across the two datasets.

While the prediction models from structural MRI sets of features, e.g., surface area, total brain volume, subcortical volume, led to the highest level of test-retest reliability, these models provided poorer predictability for cognitive abilities. This high level of test-retest reliability from structural MRI is not surprising since we should not expect drastic changes in brain anatomy in a short period of time, assuming no major brain incidents (e.g., concussion or stroke) (20). In contrast, the stacked models from Task Contrasts and FC also provided an excellent level of test-retest reliability (albeit not as high as structural MRI models), but they gave much higher predictability. Accordingly, future BWAS for cognitive abilities that would like to optimise both reliability and predictability might prefer stacking Task Contrasts and FC, or better yet stacking all the MRI data available, rather than relying on structural MRI.

### Stacking of Non-Task MRI Sets of Features Led to Better-Than-Chance Generalisability

Unlike predictability and reliability, we could only focus on the Non-Task MRI sets of features (including rest FC, cortical thickness, cortical surface area, subcortical volume and total brain volume) for cross-cohort generalisability, given the differences in fMRI tasks used in each dataset. We found that the “Stacked: Non Task” models, built from one dataset, predicted the cognitive abilities of the participants in the other two datasets better than chance. Still, if we treated the within-dataset predictability as the ceiling of cross-dataset generalisability, the cross-dataset generalisability of the “Stacked: Non Task” Models (*r*=.25) was numerically lower than, the ceiling (*r*=.4).

One caveat is that the generalisability of the “Stacked: Non Task” models between HCP Young Adults and HCP Aging (*r*=.33) is numerically higher than those between Dunedin Study and the two HCP datasets (*r*=.22). Consistent with this is the numerically higher similarity in predicted values between HCP Young Adults and HCP Aging (*r*=.51) compared to between Dunedin Study and the two HCP datasets (*r*=.32). This may reflect a higher homogeneity between the two HCP datasets. While the two HCP datasets differed in the age of participants and certain scanning parameters (e.g., TR length), HCP Aging(27) was modelled after the earlier success of HCP Young Adults(29). The two HCP datasets, for instance, used the NIH Toolbox(64) to access cognitive abilities, while Dunedin Study(28) used WAIS(65). Nonetheless, testing generalisability on Dunedin study that was conducted independently from the Human Connectome Projects may provide a more realistic picture of how deployable the “Stacked: Non Task” models to indicate cognitive abilities in the real world.

As for the non-stacked models, cross-cohort generalisability was mostly significant, except for cortical thickness. This is in line with previous studies focusing on cross-dataset generalisability of Rest FC(55–57). The generalisability of structural MRI sets of features was more varied. Some structural MRI sets gave generalisability close to predictability and provided high similarity in predicted values: total brain volume (generalisability=.24, predictability=.23 and similarity=.92) and subcortical brain volume (generalisability=.21, predictability=.22 and similarity=.84). But other structural MRI sets did not: cortical areas (generalisability=.16, predictability =.27 and similarity=.55) and cortical thickness (generalisability=.10, predictability=.23 and similarity r=.26). It is hard to pinpoint whether this is due to the differences in scanning parameters between datasets or, instead, due to the nature of the sets of features. Future research with a larger number of datasets is needed to pinpoint the characteristics of the datasets and/or features that could lead to better generalisability.

### Limitations and Future Directions

The current study has several limitations. First, it would have been desirable to examine the generalisability of stacked models involving Task Contrasts and Task FC in all cohorts. Stacked models involving Task Contrasts and FC scored high in both predictability (especially when compared to “Stacked: Non Task” models) and reliability. The inability to test their generalisability means that we cannot know for sure how deployable these highly predictable models are. Thus, for the time being, we advise researchers who would like to apply the stacked models with task fMRI on new data to follow the procedures of the original datasets as much as possible. This could be task design and scanning parameters among others.

Second, we mainly relied on the fMRI tasks and pre-processing pipelines chosen by the original investigators of each dataset. However, the fMRI tasks they chose might not be optimised for predictability, reliability and generalisability for cognitive abilities. As suggested elsewhere(52), perhaps fMRI tasks need to be designed from the ground up, using tools such as item response theory, to ensure that they capture individual differences well. Fortunately, some of the fMRI tasks (e.g., the working-memory and language tasks) provided relatively high predictability and reliability for cognitive abilities. Based on our results, in a situation where optimisation of the tasks is unknown, stacking Task Contrasts and Task FC across all of the available fMRI tasks should provide the best performance possible, given the choice of the tasks used. Similarly, each dataset’s pre-processing approach might not be optimised for BWAS with cognitive abilities as a target. For instance, for Rest FC in HCP Young Adults, we treated a choice of the two denoising strategies as another hyperparameter to select from the training sets: the investigators’ recommended method, ICA-FIX (67) and an alternative method aCompCor (68). We found that aCompCor(68) performed better in the training sets across different prediction algorithms (see Figure S32), despite not being used in the original pre-processing pipeline(29, 69, 70). While using the recommended pre-processing pipeline for each dataset allowed for easier reproducibility, we still need to test if predictability, reliability and generalisability could be further improved with more refined pipelines, optimised for predicting cognitive abilities.

Third, while predictability, reliability and generalisability are important for multimodal MRI to be applied as a neural indicator for cognitive abilities(21), other aspects still need to be accomplished for cognitive neuroscientists to truly understand the relationship between cognitive abilities and multimodal MRI measures. For instance, to reveal how the prediction models draw information from each MRI set of features, we need prediction models with good explainability(17, 71). Yet, the current prediction models are optimised for predictability, but not explainability. We previously proposed several methods to improve the explainability of prediction models (17). For example, to provide statistical inference for feature importance, future researchers could create a null distribution of feature importance via permutation, allowing them to determine whether the contribution of each feature is significantly better than chance, a technique called eNetXplorer (17, 72). Similarly, to demonstrate the pattern (i.e., linearity vs. nonlinearity) and directionality (i.e., positive vs. negative) of the relationship between each MRI feature and the prediction, future researchers could apply a visualisation technique called Accumulated Local Effects (ALE) (17, 73). Lastly, for interactive algorithms (e.g., XGBoost and Random Forest), future researchers could use Friedman’s H-statistic (17, 74) to quantify the interaction strength between each MRI feature and all other MRI features in making the prediction. However, optimising explainability is beyond the scope of this study.

## Conclusions

Cognitive neuroscientists have long dreamt of the ability to associate individual differences in cognitive abilities with brain variations(10). Yet, BWASs need to be improved in their predictability, test-retest reliability and generalisability before they can produce a robust neural indicator for cognitive abilities(16, 20, 21, 75, 76). Based on our benchmark, combining different modalities of MRI into one prediction model via stacking seems to be a viable approach to realise this dream of cognitive neuroscientists.

## Materials and Methods

### 1. Datasets

In this study, we used three datasets with 2,131 participants (1,139 females) in total (see Figure 1 for their age distribution; see Supplementary Methods for participants’ details). These three datasets have been approved by the Institutional Review Boards at the institutions where the data were collected, and informed consent was obtained (see references (27, 29, 77)).

#### Test-Retest Subsets

HCP Young Adults and Dunedin Study had a subset of participants who completed the entire MRI procedure twice. In HCP Young Adults, 45 participants were scanned *M*=139 (*SD*=67.3) days apart, and the exclusion criteria left 34 participants. In Dunedin Study, 20 participants were scanned *M*=79 (*SD=*10.4) days apart.

### 2. Features: Multimodal MRI

We used the following MRI modalities: task-fMRI contrasts, task-fMRI functional connectivity, resting state fMRI connectivity and structural MRI.

#### 2.1. Task-fMRI contrasts (Task Contrasts)

Task Contrasts reflect fMRI BOLD activity relevant to events in each task. We used Task Contrasts, pre-processed by each of the three studies. See Supplementary methods for the details about task-contrast features. Briefly, for each study, we extracted one set of 379 (i.e., ROIs) per contrast, leaving seven, five and four sets of 379 task-contrast features for HCP Young Adults, HCP Aging and Dunedin Study, respectively.

#### 2.2. Task-fMRI functional connectivity (Task FC)

Task FC reflects functional connectivity during each task. Studies have considered task FC as an important source of individual differences(53, 78, 79). As opposed to creating contrasts from fMRI time series during each task as in Task Contrasts, here we computed functional connectivity, controlling for HRF-convolved events from each task. See Supplementary methods for the details about Task FC features. Briefly, for each study, we extracted one set of 75 task FC features (principal components) per contrast, leaving seven, five and four sets of 75 task-FC features for HCP Young Adults, HCP Aging and Dunedin Study, respectively.

#### 2.3. Resting-state fMRI functional connectivity (Rest FC)

Rest FC reflects functional connectivity during rest. Both HCP Young Adults and HCP Aging included four runs of rest FC, each at 14:33 min and 6:42 min long, respectively. Dunedin Study included only one run of rest FC with 8:16 min long. See Supplementary methods for the details about Rest FC features. Briefly, we obtained one set of 75 rest FC features (principal components) for each of the three datasets.

#### 2.4. Structural MRI

Structural MRI reflects individual differences in brain anatomy. The three studies applied Freesurfer(80) to quantify these individual differences. Here we focused on four sets of features: cortical thickness, cortical surface area, subcortical volume and total brain volume. Specifically, for cortical thickness and cortical surface area, we created 148 vertex-based ROIs using the Destrieux atlas(81), while for subcortical volume, we created 19 voxel-based ROIs using the ASEG atlas(80). For total brain volume, we used summary indices provided by Freesurfer(80). See Supplementary Methods for details.

### 3. Target: Cognitive abilities

Cognitive abilities were measured outside of the MRI. HCP Young Adults and HCP Aging measured cognitive abilities using the NIH Toolbox (64). Here we used a summary score (CogTotalComp_Unadj) that covered behavioural performance from several tasks, including picture sequence memory, Flanker, list sorting, dimensional change card sort, pattern comparison, reading tests and picture vocabulary.

Dunedin Study measured cognitive abilities in several visits. We computed three scores and used them as separate targets. The first score is cognitive abilities, collected as part of the MRI visit at 45 years old via the Wechsler Adult Intelligence Scale (WAIS) IV scale(65). The second score is cognitive abilities, averaged across 7, 9 and 11 years old, collected using the Wechsler Intelligence Scale for Children - Revised (WISC-R)(82). The third score is the residual scores for the cognitive abilities(62), calculated as follows. We, first, used linear regression to predict cognitive abilities at 45 years old from cognitive abilities at 7, 9, and 11 years old. We, then, subtracted the predicted values of this linear regression from the actual cognitive abilities at 45 years old, creating the residual cognitive abilities. Negative scores of these residual cognitive abilities reflect a stronger decline in cognitive abilities, as expected from childhood cognitive abilities, compared to participants’ peers. Note that, due to the differences in the cognitive measures used for age 45 versus ages 7, 9, and 11, we could not simply subtract the scores between the two time points. While using the residual scores did not provide a change in cognitive abilities in absolute terms, they still indicate the relative changes in an individual’s cognitive abilities compared to their peers. See Prediction models below for our approach to prevent data leakage when calculating this residual score.

### 4. Prediction models

Similar to our previous work(25), we employed nested cross-validation (CV) to predict cognitive abilities from multimodal MRI data (see Figure 1). Initially, we divided the data from each study into outer folds. The number of outer folds was determined to ensure at least 100 participants per fold. Consequently, we had eight outer folds for HCP Young Adults, five for HCP Aging, and seven for the Dunedin Study. For HCP Young Adults, which included participants from the same families, we created the eight outer folds based on approximately 50 family groups, ensuring that members of the same family were in the same outer fold.

We then iterated through the outer folds, treating one fold as the test set and the remaining folds as the training set. This approach resulted in around 100 participants in each outer-fold test set for all three studies, with approximately 700, 400, and 600 participants in the outer-fold training sets for HCP Young Adults, HCP Aging, and the Dunedin Study, respectively. Next, we split each outer-fold training set into five inner folds. We iterated through these inner folds to tune the hyperparameters of the prediction models, selecting the final models based on the coefficient of determination (R²), a default option in sklearn. To prevent data leakage between the outer-fold training and test sets when calculating residual scores for cognitive abilities in the Dunedin Study, we created linear regression models to predict cognitive abilities at age 45 from abilities at ages 7, 9, and 11 using the outer-fold training set, and applied these models to the corresponding outer-fold test set.

Apart from training each of the sets of multimodal MRI features to predict cognitive abilities in separate prediction models, known as “non-stacked” models, we also combined different sets together via stacking, creating “stacked” models (see Figure 1). To train the stacked models, once we finished training all of the non-stacked models from every set of features, we computed predicted values from these non-stacked models. Specifically, we used only the data from each outer-fold training set to train the stacked models, and treated the predicted values from the non-stacked models as features to predict cognitive abilities. For example, to create a stacked model to combine Task Contrasts for HCP Young Adults, we first created non-stacked models for each of the seven sets of 379 task-contrast features available in this dataset. Then we computed the predicted values of each of these seven non-stacked models, using them as seven features in a stacked model that was trained to predict cognitive abilities in the outer-fold training sets. We tuned these stacked models using the same inner-fold CV as the non-stacked models. Accordingly, the training of stacked and non-stacked models did not involve outer-fold test sets, preventing data leakage. Note that by creating just one predicted value per non-stacked model, our stacking approach is sometimes called “late fusion,” which differs from simply concatenating features from different sets, known as “early fusion” or “flat model” (25, 83). One benefit of stacking predicted values over concatenating features is the ability to use different machine learning algorithms within and across sets of features.

We created eight stacked models: “Task Contrast” including Task Contrasts from all of the tasks, “Task FC” including Task FC from all of the tasks, “Non Task” including Rest FC and structural MRI, “Task Contrast & FC” including task contrasts and task FC from all of the tasks, “All” including all sets of features, “All excluding Task Contrast” including every set of features except Task Contrasts, “All, excluding Task FC” including every set of features except Task FC, “Resting and Task FC” including FC during rest and tasks. Note that we used two strategies, ICA-FIX(67) and aCompCor(68), to denoise Rest FC for HCP Young Adults. We ultimately picked aCompCor(68) to be included in the stacked models since it led to a better predictive performance in the outer-fold training sets (see Figure S32).

We applied corrections to reduce the influences of potential confounds. First, for all three studies, we controlled for biological sex by residualising biological sex from all MRI features and cognitive abilities. For HCP Young Adults and HCP Aging, we also controlled for age, in addition to biological sex, from all MRI features and cognitive abilities. We did not control for age in Dunedin Study because all of the participants were scanned and measured their cognitive abilities at roughly the same age. Additionally, we residualised motion (average of relative displacement, Movement_RelativeRMS_mean) from task contrasts for HCP Young Adults and Dunedin Study. We did not residualise motion from task contrasts for HCP Aging as well as task FC and rest FC for all studies since either ICA-FIX(67) or aCompCor(68) was already applied to each participant. We also standardised all MRI features. To avoid data leakage, we first applied all residualisation and standardisation on each outer-fold training set. We then applied the parameters of these residualisation and standardisation to the corresponding outer-fold test set.

As in our previous article(25), we implemented four multivariate, predictive-modelling algorithms via Scikit-learn (84): Elastic Net(58), Support Vector Regression (SVR)(59, 85), Random Forest(60) and XGBoost(61). For stacked models, we needed to apply the algorithm to two layers: 1) non-stacked layer (Step 1, Figure 1), or on each set of features and 2) stacked layer (Step 2, Figure 1), or on the predicted values from each set. Accordingly, we implemented 16 (i.e., four-by-four across two layers) combinations of algorithms for the stacked models. See the details about predictive-modelling algorithms in Supplementary Methods.

### 5. Predictability

To evaluate the predictability of prediction models, we computed the predicted values of the models at each outer-fold test set and compared them with the observed cognitive abilities. We calculated three performance indices for predictability: Pearson’s correlation (*r*), the coefficient of determination (*R*^2^) and mean absolute error (MAE). Note for *R*^2^, we applied the sum of squares definition (i.e., *R*^2^ = 1 – (sum of squares residuals/total sum of squares)) and not the square of *r*, following a previous recommendation(31).

To quantify the uncertainty around these performance indices, we calculated bootstrapped 95% confidence intervals (CI)(86). Here, we combined predicted and observed cognitive abilities across outer-fold test sets, sampled these values with replacement 5,000 times and computed the three performance indices each time, giving us a bootstrapped distribution for each index. If the 95% CI of the *r* or *R*^2^ bootstrapped distribution were higher than zero, then the predictability from a particular prediction model was better than chance.

To compare predictability among prediction models, we also used the bootstrapping approach(86). Similar to the above, we sampled, with replacement for 5,000 times, the observed cognitive abilities along with their predicted values from different prediction models across outer-fold test sets. In each sample, we computed each performance index of each prediction model and subtracted this index from that of the prediction model with the highest predictability of each dataset. If the 95% CI of this distribution of the subtractions were higher than zero, then we concluded that the prediction model we tested had significantly poorer performance than the prediction model with the highest predictability. We applied this approach separately for non-stacked and stacked models, allowing us to evaluate the best non-stacked and stacked models for each dataset.

To understand how the prediction models drew information across multimodal MRI features, we plotted Elastic-Net coefficients. We chose Elastic-Net coefficients because 1) Elastic Net led to high predictability, as high as or higher than other algorithms (see Results), and 2) the Elastic Net coefficients are readily interpretable. Elastic Net creates a predicted value from a weighted sum of features, and therefore a stronger magnitude of an Elastic Net coefficient means a higher contribution to the prediction.

Our use of nested CV led to separate Elastic-Net models, one for each outer fold, making it hard to visualise Elastic-Net coefficients across all participants in each dataset. To address this, we retrained Elastic Net using the whole data (i.e., without splitting the data into outer folds) in each dataset and applied five CVs to tune the model. We then plotted the Elastic-Net coefficients on brain images using brainspace(87) and nilearn(88). Note we modelled Task FC and Rest FC after reducing their dimension via PCA. To extract the feature importance at each ROI-pair index, we multiplied the absolute PCA scores with Elastic Net coefficients and then summed the multiplied values across the 75 components, resulting in 71,631 ROI-pair indices.

### 6. Test-Retest Reliability

Given the high predictability of Elastic Net (see Figures S19-28), we evaluated the test-retest reliability of the prediction models based on Elastic Net. To evaluate test-retest reliability, we used HCP Young Adults and Dunedin Study test-retest subjects (i.e., participants who were scanned twice) as the test set and the rest of the participants in each dataset as a training set. Within the training set, we used the same five CVs to tune the Elastic Net models, as described above. We then examined the test-retest reliability of the predicted values between the first and second MRI sessions, as quantified by intraclass correlation (ICC) 3,1(89) via pingouin (https://pingouin-stats.org/):

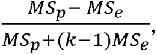

where *MSp* is mean square for participants, *MSe* is mean square for error, and *k* is the number of time points. We used the following criteria to interpret ICC(90). ICC < 0.4 as poor, ICC ≥ 0.4 and < 0.6 as fair, ICC ≥ 0.6 and < 0.75 as good and ICC ≥ 0.75 as excellent reliability.

### 7. Generalisability

Similar to test-retest reliability, we evaluated the generalisability of the prediction models based on Elastic Net given the high predictability of Elastic Net based on bootstrapped comparisons (see Figures S19-28). For features, because the three datasets used mostly different fMRI tasks, we focused on the generalisability of non-task sets of features (including rest FC, cortical thickness, cortical surface area, subcortical volume and total brain volume) and their stacked model, “Stacked: Non Task”. For the target, we standardised cognitive abilities using a Z-score within each dataset, so that the target for each dataset was at the same standardised scale before model fitting. This is because Dunedin Study used WAIS-IV for measuring cognitive abilities, while HCP-YA and HCP-A used NIH toolbox(64). Note, for Dunedin

Study, we only focused on cognitive abilities collected during the MRI visit at age 45 (as opposed to during earlier visits) as the target, given that the other two studies only provided cognitive abilities during the MRI visit.

To evaluate generalisability across datasets, we treated one of the three datasets as a training set and the other two as two separate test sets. We computed the predicted values of the models at each test dataset and compared them with the observed cognitive abilities using Pearson’s correlation (*r*). To examine if the generalisability was statistically significant, we bootstrapped *r* 5000 times. If the 95% bootstrapped CI were higher than zero, the *r* was statistically significantly better than chance. We also compared generalisability across datasets to predictability within each dataset using nested CVs. For predictability within each dataset, we combined predicted values across outer test sets and compared them with the observed cognitive abilities. We considered the predictability within each dataset as the ceiling of how high generalisability across datasets could be.

To further understand the extent to which the prediction models built from one dataset are different from those built from another, we also examined the similarity between predictive values. Here, using Pearson’s correlation (*r*), we compared the correlation in predictive values from the prediction models built from the same dataset and those built from another dataset. Similar to generalisability, to test if the similarity between predictive values was statistically significant, we bootstrapped *r* 5000 times. If the 95% bootstrapped CI were higher than zero, the *r* was statistically significantly better than chance.

## Data sharing plans

All codes are available at https://github.com/HAM-lab-Otago-University/Predictability-Reliability-Generalizability.

Instructions for data access can be found here:

HCP Young Adults https://www.humanconnectome.org/study/hcp-young-adult, HCP Aging https://www.humanconnectome.org/study/hcp-lifespan-aging and Dunedin Study https://dunedinstudy.otago.ac.nz/.

While we did not preregister our data-analysis plan for HCP Young Adults and HCP Aging, we preregistered our plan to test predictability and test-retest reliability for the Dunedin study prior to having access to the dataset at https://dunedinstudy.otago.ac.nz/files/1639954373_Pat%20Multimodal%20brain%20concept%20paper_NP%20TEMsigned.docx.pdf.

## Acknowledgements

We thank the Dunedin Study members, their families and friends for their long-term involvement. Thank Dunedin Unit Director, Professor Reremoana (Moana) Theodore, Unit research staff, Professor Terrie E. Moffitt, Professor Avshalom Caspi, previous Study Director, Emeritus Distinguished Professor, the late Richie Poulton, for his leadership during the Study’s research transition from young adulthood to aging (2000–2023) and Study founder, Dr Phil A. Silva. Professor Terrie E. Moffitt was especially helpful in providing comments on earlier drafts. The Dunedin Multidisciplinary Health and Development Research Unit is supported by the New Zealand Health Research Council and New Zealand Ministry of Business, Innovation and Employment. Funding support was also received from The New Zealand Health Research Council Programme Grant (16–604), US-National Institute on Aging grants AG049789-06, and R01AG032282-11 and UK Medical Research Council grants MR/X021149/1. Additional data were provided by the Human Connectome Project – Young Adults and the Human Connectome Project in Aging. The Human Connectome Project – Young Adults, WU-Minn Consortium (Principal Investigators: David Van Essen and Kamil Ugurbil; 1U54MH091657) was funded by the 16 NIH Institutes and Centers that support the NIH Blueprint for Neuroscience Research; and by the McDonnell center for Systems Neuroscience at Washington University. The Human Connectome Project in Aging was supported by the National Institute on Aging of the National Institutes of Health under Award Number U01AG052564 and by funds provided by the McDonnell Center for Systems Neuroscience at Washington University in St. Louis. The content is solely the responsibility of the authors and does not necessarily represent the official views of the National Institutes of Health. The author(s) wish to acknowledge the use of New Zealand eScience Infrastructure (NeSI) high performance computing facilities, consulting support and/or training services as part of this research. New Zealand’s national facilities are provided by NeSI and funded jointly by NeSI’s collaborator institutions and through the Ministry of Business, Innovation & Employment’s Research Infrastructure programme. URL https://www.nesi.org.nz. A.T., T.M. and N.P. were supported by Health Research Council Funding (grant number 21/618 and 24/838), by the University of Otago and by Neurological Foundation of New Zealand (grant number 2350 PRG).

