## Supplementary Methods, Figures and Tables for "Improving Predictability, Reliability and Generalisability of Brain-Wide Associations for Cognitive Abilities via Multimodal Stacking"

**Participants**

1.1. **Human Connectome Project Young Adults (HCP Young Adults):** HCP Young Adults S1200 Release included multiple MRI modalities and cognitive measurements from 1,206 healthy participants (22-35 years old)(1, 2). We excluded participants with the “A” (anatomical anomalies) or “B” (segmentation and surface) flag, with any known major issues(3) or with any incomplete MRI or cognitive measurements(4). These exclusions left 873 participants (473 females, *M*=28.7 (*SD*=3.7) years old) in our analysis.
1.2. **Human Connectome Project Aging (HCP Aging):** HCP Aging 2.0, released on 24^th^ February 2021, consisted of ‘typical-aging’ participants, from 36 to 100 years old, without identified pathological causes of cognitive decline (e.g., stroke and clinical dementia)(5). This release provided data from 725 participants. After applying the same exclusion criteria as HCP Young Adults, 504 participants (293 females, *M*=57.83 (*SD*=14.25) years old) remained in our analysis.
1.3. **The Dunedin Multidisciplinary Health and Development Study (Dunedin Study):** Dunedin Study is a longitudinal study of the health and behaviours of 1,037 individuals, born between April 1972 and March 1973 in Dunedin, a small city in the South Island of New Zealand(6). Dunedin Study has conducted many assessments on the participants since birth. The study collected MRI data of multiple modalities from 875 participants when they were 45 years old. After excluding incomplete MRI or cognitive measurements, 754 participants (373 females) remained in our analysis.

**Task-fMRI contrasts (Task Contrasts)**

**HCP Young Adults:** HCP Young Adults provided complete details of scanning parameters and pre-processing pipeline elsewhere(1, 7, 8). Briefly, they implemented 720-ms TR, B0 distortion correction, motion correction, gradient unwrap, boundary-based co-registration to T1-weighted image, non-linear registration to MNI152 space, grand-mean intensity normalization, high-pass filtering with a cut-off at sigma = 200s, surface generation and the multimodal alignment protocol (MSMAll)(9). The study included seven fMRI tasks, each with two runs with different phase encodings: left-to-right (LR) and right-to-left (RL). HCP Young Adults applied a general-linear model (GLM) to combine the two runs and computed GLM contrasts in a Connectivity Informatics Technology Initiative (CIFTI) format, containing both cortical surface and subcortical volume. We parcellated these contrasts into 379 regions of interest (ROIs), consisting of 360 cortical-surface ROIs from the Glasser atlas(9) and 19 subcortical ROIs from the Freesurfer’s Automatic subcortical SEGmentation (ASEG) atlas (10). We then extracted the average value for each ROI.

In each of the seven tasks, we chose only one GLM contrast between the main experimental vs. control conditions. The study provided full task descriptions for these conditions in the release documentation(7, 11). We used the following contrasts: face vs. shape for the face, or emotion-processing, task(12), reward vs. punishment for the gambling task(13–16), story vs. math for the language task(17), averaged movement vs. cue for the motor task(18–21), relational vs. match for the relational task(22), theory of mind vs. random for the social cognition task(23–26) and 2-back vs. 0-back for the working memory task(27). Accordingly, we obtained seven sets of 379 (i.e., ROIs) task-contrast features for HCP Young Adults.

**HCP Aging:** HCP Aging used similar scanning parameters and a pre-processing pipeline to HCP Young Adults(5, 8, 28). There were a few differences: task fMRI was collected using TR at 800 ms (as opposed to 720 ms) and using the posterior-to-anterior (PA) phase (as opposed to LR and RL), and the investigators applied linear detrend (as opposed to high-pass filtered cut-off at sigma = 200s ) and ICA-FIX(9).

HCP Aging did not provide task contrasts, but rather pre-processed time series in the CIFTI format for each task. To extract task contracts from the time series, we applied the “fake” NIFTI approach(29) using FMRI Expert Analysis Tool (FEAT) from FMRIB Software Library (FSL)(30). Here we first transformed the CIFTI file into a 32767x3 NIFTI array, then conducted GLM and converted the output contrasts back into the CIFTI format. For the GLM, we modified the fsf-template file, given by the HCP Aging (<https://github.com/Washington-University/HCPpipelines/tree/master/Examples/fsf_templates/HCP-Aging>). Specifically, we regressed the time series on the convolved task events using a double-gamma canonical hemodynamic response function (HRF). We used a default high pass cut-off in FSL at sigma = 200s. Similar to HCP Young Adults, we parcellated the contrasts into 379 ROIs using the Glasser(9) and ASEG(10) atlases for cortical and subcortical regions, respectively.

HCP Aging included three fMRI tasks(5). Similar to HCP Young Adults, we chose GLM contrasts between the main experimental vs. control conditions. We used the following contrasts for each task: encoding vs. distractor, recall vs. distractor and encoding vs. recall for the facename task(31–33), NoGo vs. Go for the Conditioned Approach Response Inhibition Task (CARIT) go-nogo task (34), and stimulus vs. baseline for the visual motor (VisMotor) task(5). Note the facename task taps into episodic memory. There were three blocks: “Encoding” when participants needed to memorise the names of faces, “Distractor” when participants were distracted by an irrelevant task and “Recall” when participants needed to recall the names of the previously shown faces. Because each block reflects different processes of episodic memory, we chose all possible pairs of contrasts for this task. Accordingly, we obtained five sets of 379 task-contrast features for HCP Aging.

**Dunedin Study**: Dunedin Study provided complete details of the scanning parameters and pre-processing pipeline elsewhere(35, 36). Briefly, the Dunedin Study investigators implemented 2000-ms TR, B0 distortions correction(37), despike(38), slice-timing and motion correction(38), boundary-based co-registration to T1-weighted image(39) and non-linear registration to MNI space(40, 41). The study applied a general-linear model (GLM) using the AFNI 3dREMLfit(38)and applied the HCP Minimal Preprocessing Pipeline (MPP) (<https://github.com/Washington-University/HCPpipelines/releases>) to project the contrast images into Cifti format. Similar to the HCP Young Adults and HCP Aging, the study parcellated the contrasts into 379 ROIs using the Glasser(9) and ASEG(10) atlases using the Ciftify toolbox (https://github.com/edickie/ciftify).

Dunedin Study included four fMRI tasks(36). The study provided us with the following contrasts: encoding vs. distractor for the facename task(33), face vs. shape for the face, or emotion processing, task(12), gain vs. neutral anticipation for the monetary incentive delay (MID) task(42) (see <https://www.haririlab.com/methods/dbis_vs.html>) and incongruent vs. congruent for the Stroop task (see <https://www.haririlab.com/methods/dbis_control.html>). Note the facename task in Dunedin Study was different from that in HCP Aging. During “Distractor”, participants in the Dunedin study had to perform an odd/even-number identification task (<https://www.haririlab.com/methods/dbis_hippocampus.html>) whereas participants in the HCP Aging had to perform a Go/NoGo task(5, 33). The face, or emotion processing, task in Dunedin Study was also different from that in HCP Young Adults. The Dunedin Study version included four different facial expressions: fearful, angry, surprised or neutral (see <https://www.haririlab.com/methods/dbis_amygdala.html>) and lasted 6:40 minutes, whereas the HCP Young Adults version had only two facial expressions: angry and fearful and lasted only 2:16 minutes. Altogether, we obtained four sets of 379 task-contrast features for Dunedin Study.

**Task-fMRI functional connectivity (Task FC)**

**HCP Young Adults**: Because HCP Young Adults did not provide denoised fMRI time series for each task, we denoised the time series ourselves. We first converted raw task fMRI data into Brain Imaging Data Structure (BIDS)(43) using the hcp2bids function from MICA (<https://github.com/MICA-MNI/micapipe-supplementary/blob/main/functions/hcp2bids>). To pre-process the BIDS data, we applied fMRIPrep(44, 45) without slice-timing correction. We then used XCP-D(46) to further process the data. Specifically, we used the aCompCor(47) flag and created customised regressors for task events, leaving the following regressors of no interest in our GLM: HRF-convolved task events, 12 motion estimates (i.e., x, y, z rotation and translation and their derivatives), five white-matter and five cerebrospinal fluid aCompCor components, a linear trend, a cosine (1/128Hz) and an intercept. We, then, concatenated LR and RL runs and parcellated the time series into 379 ROIs using the Glasser(9) and ASEG(10) atlases. Subsequently, we computed r-to-z transformed Pearson’s correlations across all possible pairs, resulting in 71,631 non-overlapping FC indices for each task. To reduce the dimensionality of task FC, we applied principal component analysis (PCA) of 75 components (48–50). To avoid data leakage, we conducted PCA on each training set and applied its definition to the corresponding test set. Note we calculated task FC for all of the tasks in HCP Young Adults except the emotion processing task due to its short duration (2:16 mins). Accordingly, we obtained six sets of 75 task FC features for HCP Young Adults.

**HCP Aging**: Unlike HCP Young Adults, the fMRI time series for each task in HCP Aging was denoised by ICA-FIX(9). Accordingly, we did not implement fMRIprep(44, 45) and XCP-D(46) to clean the HCP Aging time series further. Instead, we only regressed out the HRF-convolved task events from the time series and applied a high-pass filter at 0.008 Hz following previous work(35) using nilearn(51). For each of the three tasks, we applied the same parcellation, Pearson’s correlation, r-to-z transformation and PCA as we did for HCP Young Adults. Accordingly, we obtained three sets of 75 task FC features for HCP Aging.

**Dunedin Study:** The study provided denoised time series and details on their denoising strategies elsewhere(35). Briefly, they applied the following regressors of no interest in their GLM: 12 motion estimates with derivatives, five CompCor components from white-matter and from cerebrospinal fluid(47), the mean global signal, and task-evoked coactivations via AFNI TENT(38). The study also implemented bandpass filtering between 0.008 and 0.1 Hz and censored motion artefacts with thresholds of 0.35-mm framewise displacement and 1.55 standardised DVARS. For each of the four tasks, we used the same parcellation, Pearson’s correlation, r-to-z transformation and PCA as we did for HCP Young Adults and HCP Aging. Accordingly, we obtained four sets of 75 task FC features for Dunedin Study.

**Resting-state fMRI functional connectivity (Rest FC)**

**HCP Young Adults**: We applied two denoising strategies on the fMRI time series during rest. The first strategy was ICA-FIX(9) which was done by the study. Specifically, we high-pass-filtered the provided ICA-FIX-denoised time series at 0.008Hz using nilearn(51) and concatenated them across four runs. The second strategy was aCompCor(47). We applied the same pre-processing steps on rest FC as we did on task FC for HCP Young Adults (see above), except that here we did not use HRF-convolved task events as regressors of no interest since there were no events. We chose to apply this aCompCor strategy in addition to ICA-FIX, so that rest FC and task FC for HCP Young Adults were consistent with each other.

**HCP Aging**: The study applied ICA-FIX(9) to denoise fMRI time series during both rest and tasks, meaning that the denoising strategies done by the study for rest FC and task FC were already consistent with each other. Accordingly, we only used ICA-FIX(9) here. Like HCP Young Adults, we further applied high-pass filtered at 0.008Hz on the provided ICA-FIX-denoised time series and concatenated them across four runs.

**Dunedin Study**: The Dunedin Study investigators applied the same denoising strategies to Rest FC as they did to Task FC(35) (see above), leaving bandpass-filtered time series.

For each of the three datasets, we took the denoised, filtered time series and applied the same parcellation, Pearson’s correlation, r-to-z transformation and PCA as we did for Task FC. Accordingly, we obtained one set of 75 rest FC features for each of the datasets.

**Structural MRI**

For cortical thickness and cortical surface area, we parcellated the cortical regions into 148 ROIs using the Destrieux atlas(52), leaving two sets of 148 features. For subcortical volume, we parcellated the subcortical regions into 19 ROIs using the ASEG atlas(10), leaving one set of 19 features. For total brain volume, we included five summary statistics from Freesurfer(10): total cortical grey matter volume (FS_TotCort_GM_Vol), total cortical white matter volume (FS_Tot_WM_Vol), total subcortical grey matter volume (FS_SubCort_GM_Vol), estimated intra-cranial volume (FS_IntraCranial_Vol) and ratio of brain segmentation volume to estimated total intracranial volume (FS_BrainSegVol_eTIV_Ratio). This left another set of five features.

**Predictive Algorithms**

Elastic Net(53) is a general form of penalized regression. Elastic Net has two hyperparameters: 1) 'α', determining the degree of penalty to the sum of the feature's slopes and 2) 'l1 ratio', determining the degree to which the sum of either the squared (known as ‘Ridge’; l1 ratio=0) or absolute (known as ‘Lasso’; l1 ratio=1) slopes is penalised. We performed a logarithmic scale grid search on the 70 possible α values, ranging from 10^−1^ to 10^2^, and a linear scale grid search on the 25 possible l1 ratio values, ranging from 0 to 1.

Support Vector Regression (SVR) (54, 55) is a kernel-based algorithm. Unlike Elastic Net, SVR with the Radial Basis Function (RBF) allows for non-linearity between each feature and the target and interaction among features. We tuned the following hyperparameters: 1) ‘ε’, determining the margin of tolerance where no penalty is given to errors, 2) ‘γ’, determining the kernel coefficient, 3) ‘C’ or ‘complexity,’ determining penalty for high complexity. We performed a linear scale grid search on the 10 possible ε values, ranging from 0.02 to 0.22. We also performed a grid search on 20 possible γ values, using the following set: 10^-8^, 10^-7^, 10^-6^, 10^-5^, 10^-4^, 10^-3^, 3*10^-8^, 3*10^-7^, 3*10^-6^, 3*10^-5^, 3*10^-4^, 3*10^-3^, 6*10^-8^, 6*10^-7^, 6*10^-6^, 6*10^-5^, 6*10^-4^, 6*10^-3^ as well as one over the multiplication of the number of features and variance of features (known as ‘scale’ in sklearn) and one over the number of features (known as ‘auto’ in sklearn). Finally, we also performed a grid search on eight possible C values using the following set: 1, 6, 9, 10, 12, 15, 20 and 50.

Random Forest(56) is a tree-based algorithm. The algorithm bootstraps observations and incorporates a random subset of features at each split of tree building. The algorithm made predictions based on an aggregation of predicted values across bootstrapped trees. Like SVR with RBF, Random Forest allows for a non-linear relationship between each feature and the target and for interactions amongst features. Here we used 5000 trees (‘n_estimators’) and tuned two hyperparameters: 1) ‘max_depth’, determining the maximum depth of each tree and 2) ‘max_feature’, determining the number of features that are randomly sampled at each split. We performed a grid search for 10 possible max_depth values, using the integers ranging from 1 to 10. We also performed a grid search for three possible max_feature values: the number of features itself, the square root of the number of features and log-based 2 of the number of features.

XGBoost(57) is another tree-based algorithm, which also allows for non-linearity and interaction. It generates sequential trees where a current tree adapts from the gradients of previous trees. We used ‘gbtree’ as a booster and tuned four hyperparameters: 1) ‘max_depth’, determining the maximum depth of each tree, 2) ‘γ’, determining a minimum loss reduction required to make a further partition on a leaf node of the tree, 3) ‘subsample’, determining the ratio of the training instance, 4) ‘learning_rate’, determining the speed of tree adaptation. We performed a linear scale grid search on nine possible max_depth values using integers, ranging from 1 to 9. We also performed a logarithmic scale grid search on the six possible γ values, ranging from 10^-5^ to 0.7. Next, we performed a grid search on three possible subsample values: 0.6, 0.8 and 1. Lastly, performed a logarithmic scale grid search on the five learning_rate values, ranging from 10^-5^ to 10^-0.1^. For other hyperparameters, we used their default values (see <https://xgboost.readthedocs.io/>).

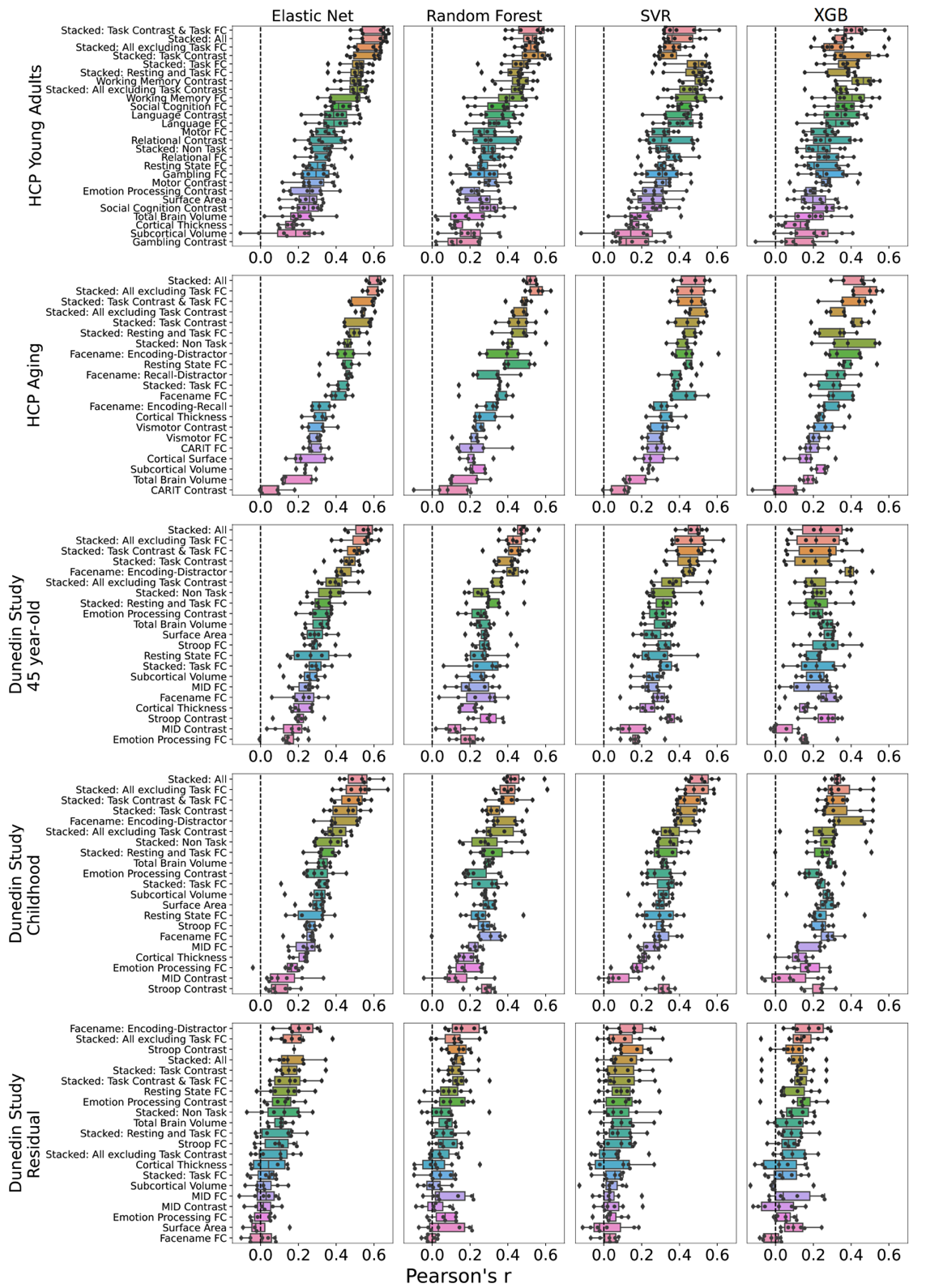

***Figure S1****.* ***Predictability (Pearson’s correlation, r) of stacked and non-stacked models for each predictive-modelling algorithm and dataset.*** *Higher is better.* *Each dot represents predictive performance at each outer-fold test set. Note that, for the stacked models, we only showed those with the same predictive algorithm across both layers here. For Dunedin Study, childhood scores reflect cognitive abilities, averaged across 7, 9 and 11 years old, and negative residual scores reflect a stronger decline in cognitive abilities, as expected from childhood cognitive abilities, compared to participants’ peers. SVR = Support Vector Regression; XGB = XGBoost; FC = Functional Connectivity.*

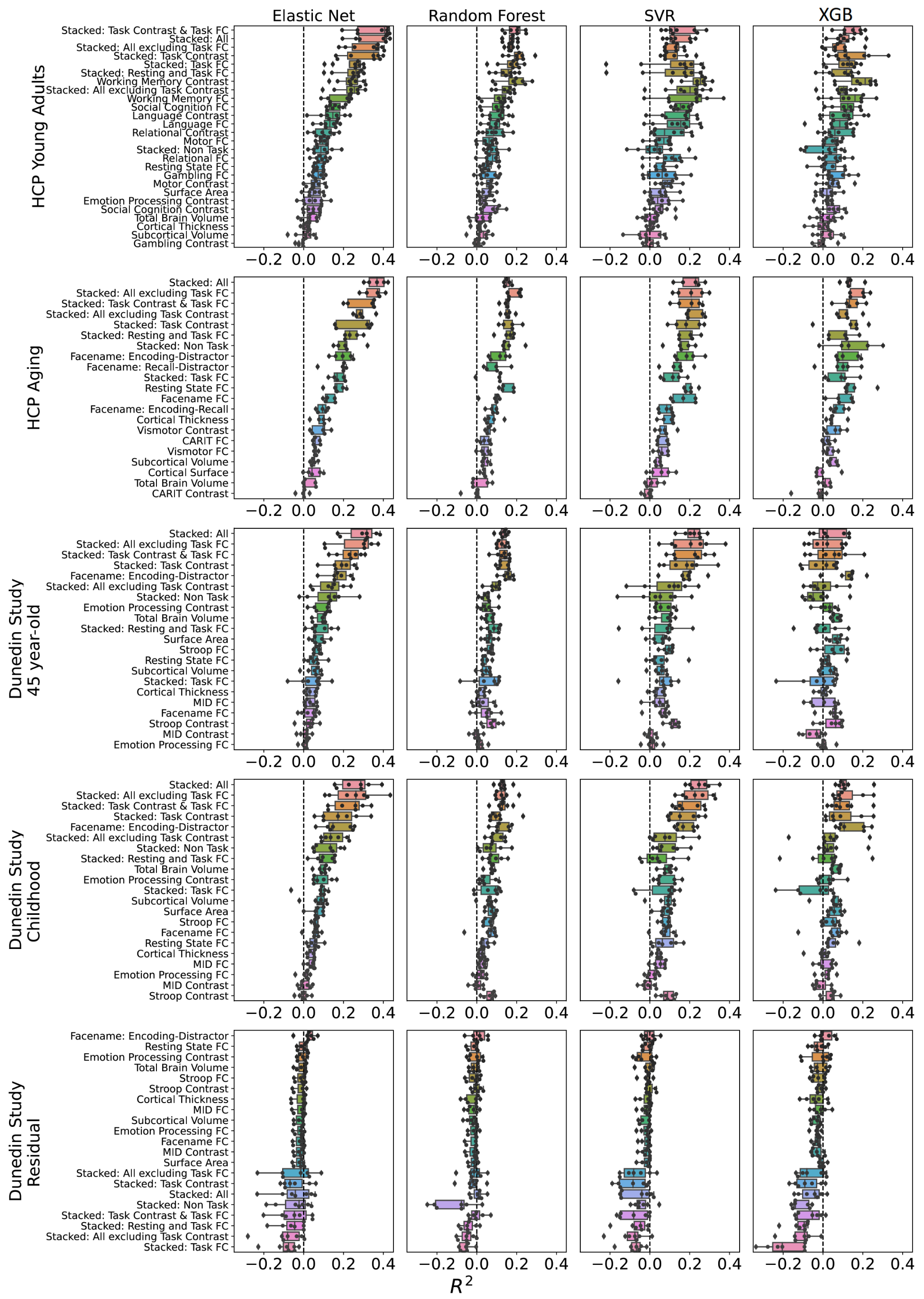

**Figure S2**. **Predictability (coefficient of determination, R^2^) of stacked and non-stacked models for each predictive-modelling algorithm and dataset.** Higher is better. Each dot represents predictive performance at each outer-fold test set. Note that, for the stacked models here, we only showed those with the same predictive algorithm across both layers. For stacked models with different predictive algorithms between layers, please see Figures S5-S6. For Dunedin Study, childhood scores reflect cognitive abilities, averaged across 7, 9 and 11 years old, and negative residual scores reflect a stronger decline in cognitive abilities, as expected from childhood cognitive abilities, compared to participants’ peers. SVR = Support Vector Regression; XGB = XGBoost; FC = Functional Connectivity.

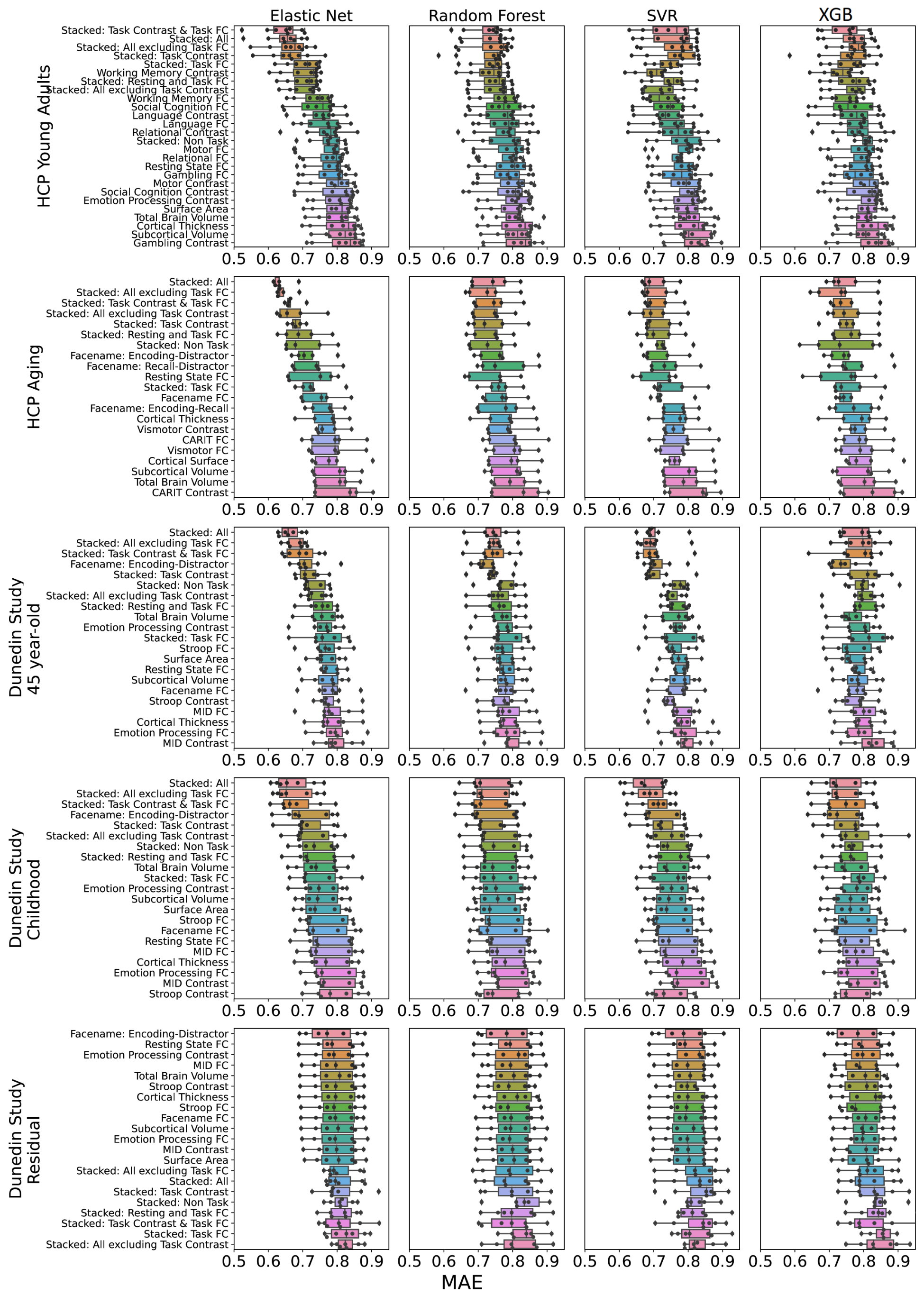

**Figure S3**. **Predictability (mean absolute error, MAE) of stacked and non-stacked models for each predictive-modelling algorithm and dataset.** Lower is better. Each dot represents predictive performance at each outer-fold test set. Note that, for the stacked models here, we only showed those with the same predictive algorithm across both layers. For stacked models with different predictive algorithms between layers, please see Figures S7-S8. For Dunedin Study, childhood scores reflect cognitive abilities, averaged across 7, 9 and 11 years old, and negative residual scores reflect a stronger decline in cognitive abilities, as expected from childhood cognitive abilities, compared to participants’ peers. SVR = Support Vector Regression; XGB = XGBoost; FC = Functional Connectivity.

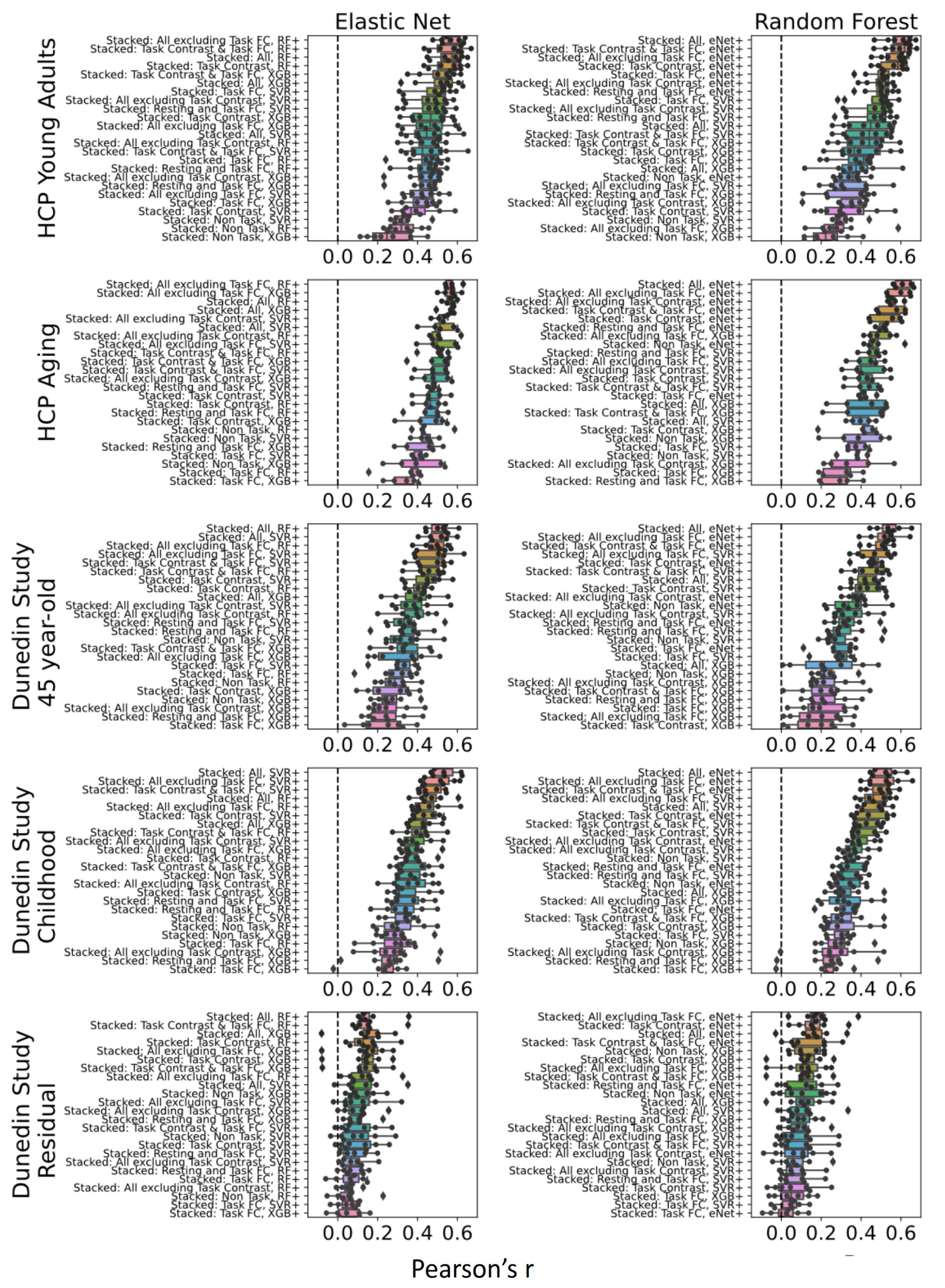

**Figure S4**. **Predictability (Pearson’s correlation, *r*) of stacked models having different predictive algorithms between layers with Elastic Net or Random Forest at the second layer (indicated by top labels).** Higher is better. The predictive-modelling algorithms at the first layer are indicated by left-side labels. Each dot represents predictive performance at each outer-fold test set. For Dunedin Study, childhood scores reflect cognitive abilities, averaged across 7, 9 and 11 years old, and negative residual scores reflect a stronger decline in cognitive abilities, as expected from childhood cognitive abilities, compared to participants’ peers. eNet = Elastic Net; RF = Random Forest; SVR = Support Vector Regression; XGB = XGBoost; FC = Functional Connectivity.

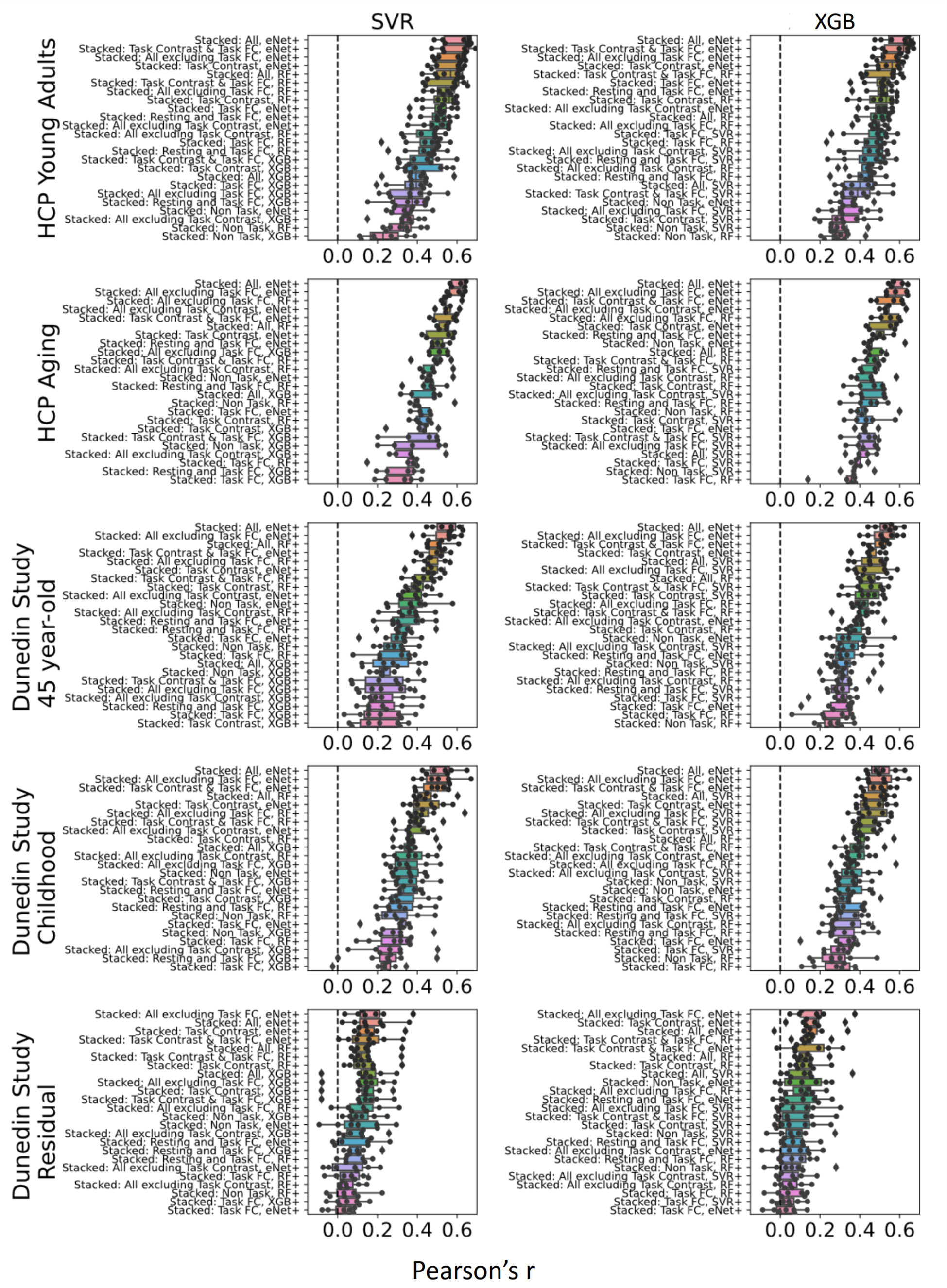

**Figure S5**. **Predictability (Pearson’s correlation, *r*) of stacked models having different predictive algorithms between layers with Support Vector Regression or XGBoost at the second layer (indicated by top labels).** Higher is better. The predictive-modelling algorithms at the first layer are indicated by left-side labels. Each dot represents predictive performance at each outer-fold test set. For Dunedin Study, childhood scores reflect cognitive abilities, averaged across 7, 9 and 11 years old, and negative residual scores reflect a stronger decline in cognitive abilities, as expected from childhood cognitive abilities, compared to participants’ peers. eNet = Elastic Net; RF = Random Forest; SVR = Support Vector Regression; XGB = XGBoost; FC = Functional Connectivity.

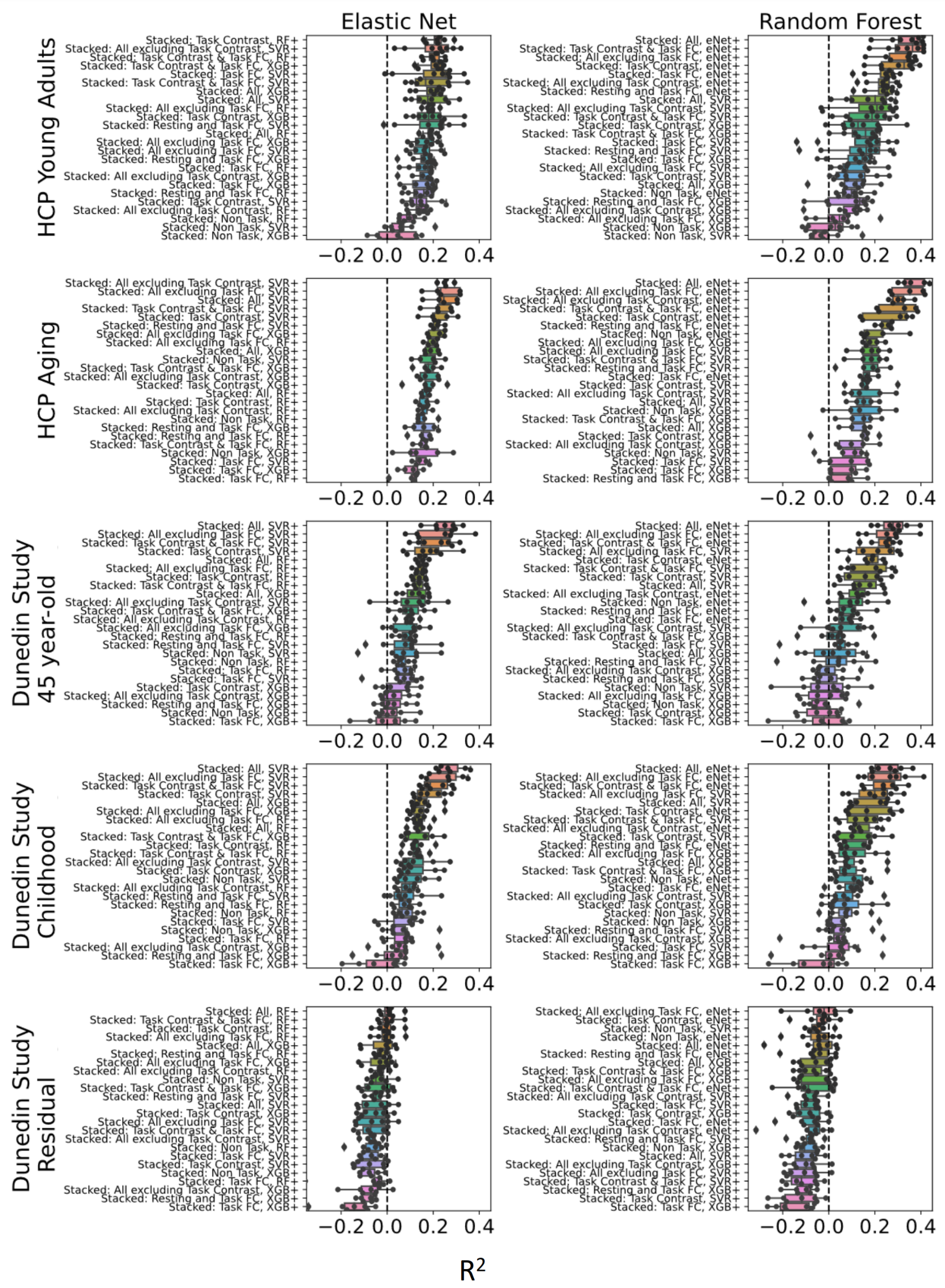

**Figure S6**. **Predictability (coefficient of determination, *R*^2^) of stacked models having different predictive algorithms between layers with Elastic Net or Random Forest at the second layer (indicated by top labels).** Higher is better. The predictive-modelling algorithms at the first layer are indicated by left-side labels. Each dot represents predictive performance at each outer-fold test set. For Dunedin Study, childhood scores reflect cognitive abilities, averaged across 7, 9 and 11 years old, and negative residual scores reflect a stronger decline in cognitive abilities, as expected from childhood cognitive abilities, compared to participants’ peers. eNet = Elastic Net; RF = Random Forest; SVR = Support Vector Regression; XGB = XGBoost; FC = Functional Connectivity.

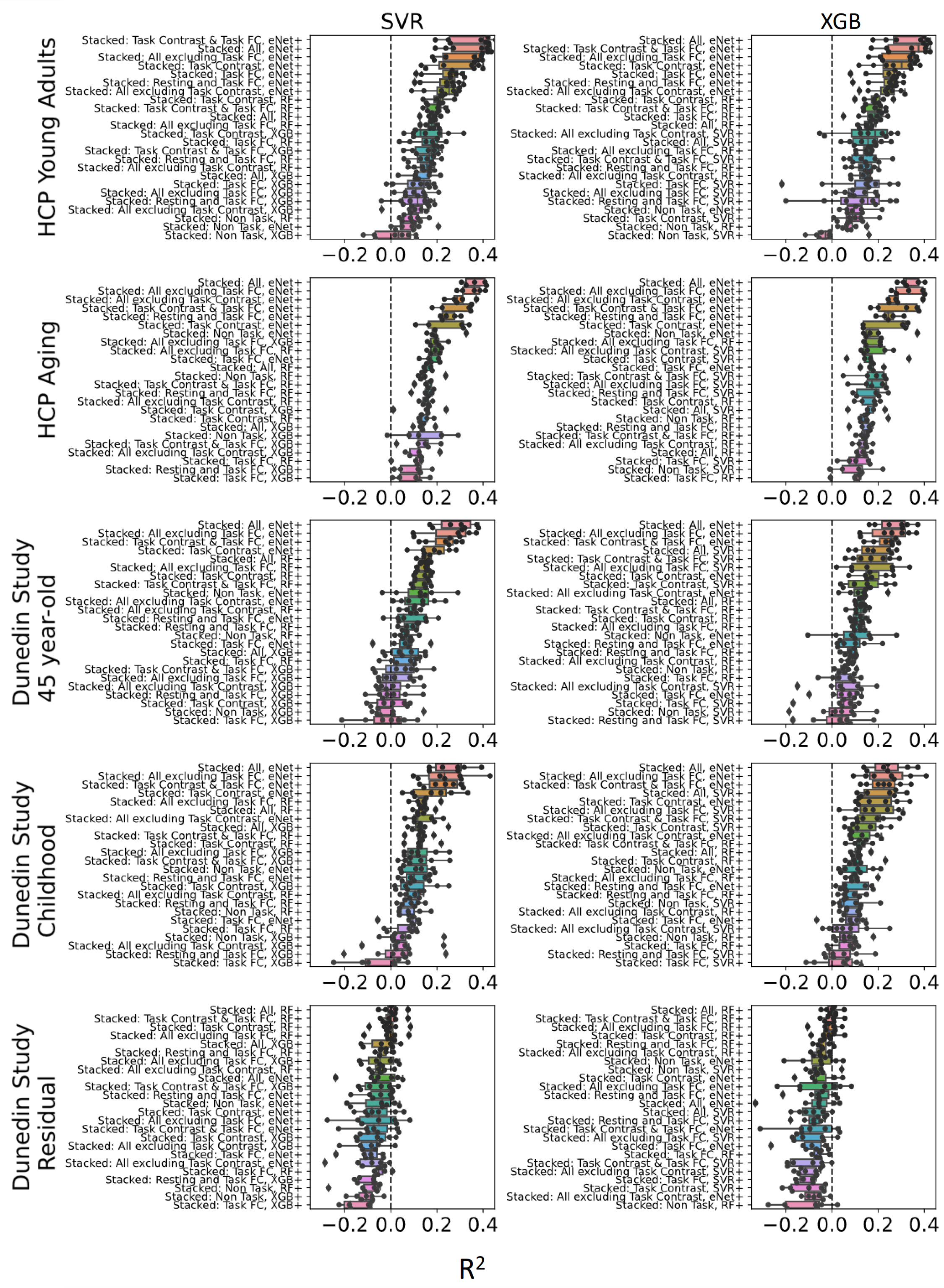

**Figure S7**. **Predictability (coefficient of determination, *R*^2^)** **of stacked models having different predictive algorithms between layers with Support Vector Regression or XGBoost** **at the second layer (indicated by top labels).** Higher is better. The predictive-modelling algorithms at the first layer are indicated by left-side labels. Each dot represents predictive performance at each outer-fold test set. For Dunedin Study, childhood scores reflect cognitive abilities, averaged across 7, 9 and 11 years old, and negative residual scores reflect a stronger decline in cognitive abilities, as expected from childhood cognitive abilities, compared to participants’ peers. eNet = Elastic Net; RF = Random Forest; SVR = Support Vector Regression; XGB = XGBoost; FC = Functional Connectivity.

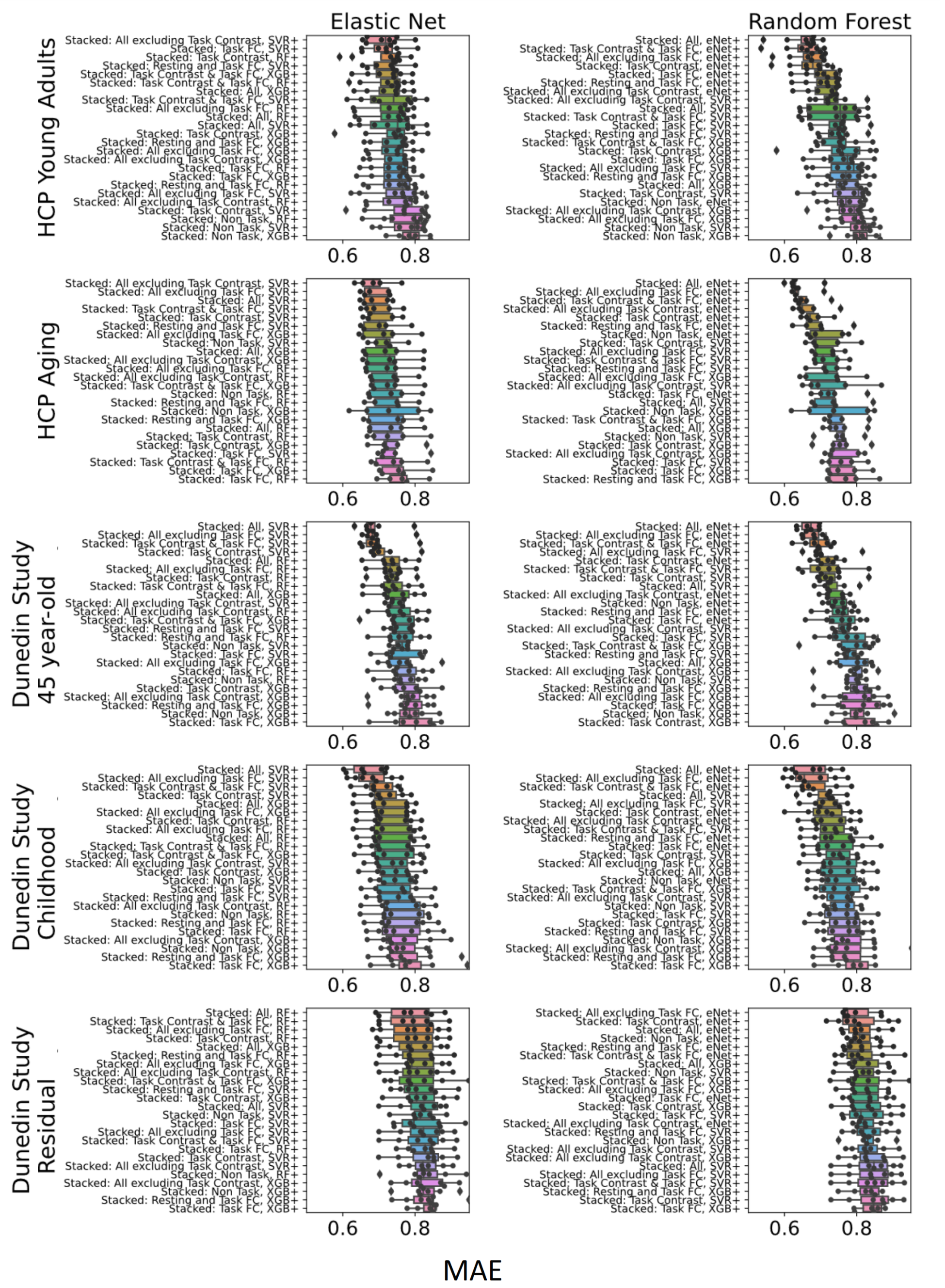

**Figure S8**. **Predictability (mean absolute error, MAE) of stacked models having different predictive algorithms between layers with Elastic Net or Random Forest at the second layer (indicated by top labels).** Lower is better. The predictive-modelling algorithms at the first layer are indicated by left-side labels. Each dot represents predictive performance at each outer-fold test set. For Dunedin Study, childhood scores reflect cognitive abilities, averaged across 7, 9 and 11 years old, and negative residual scores reflect a stronger decline in cognitive abilities, as expected from childhood cognitive abilities, compared to participants’ peers. eNet = Elastic Net; RF = Random Forest; SVR = Support Vector Regression; XGB = XGBoost; FC = Functional Connectivity.

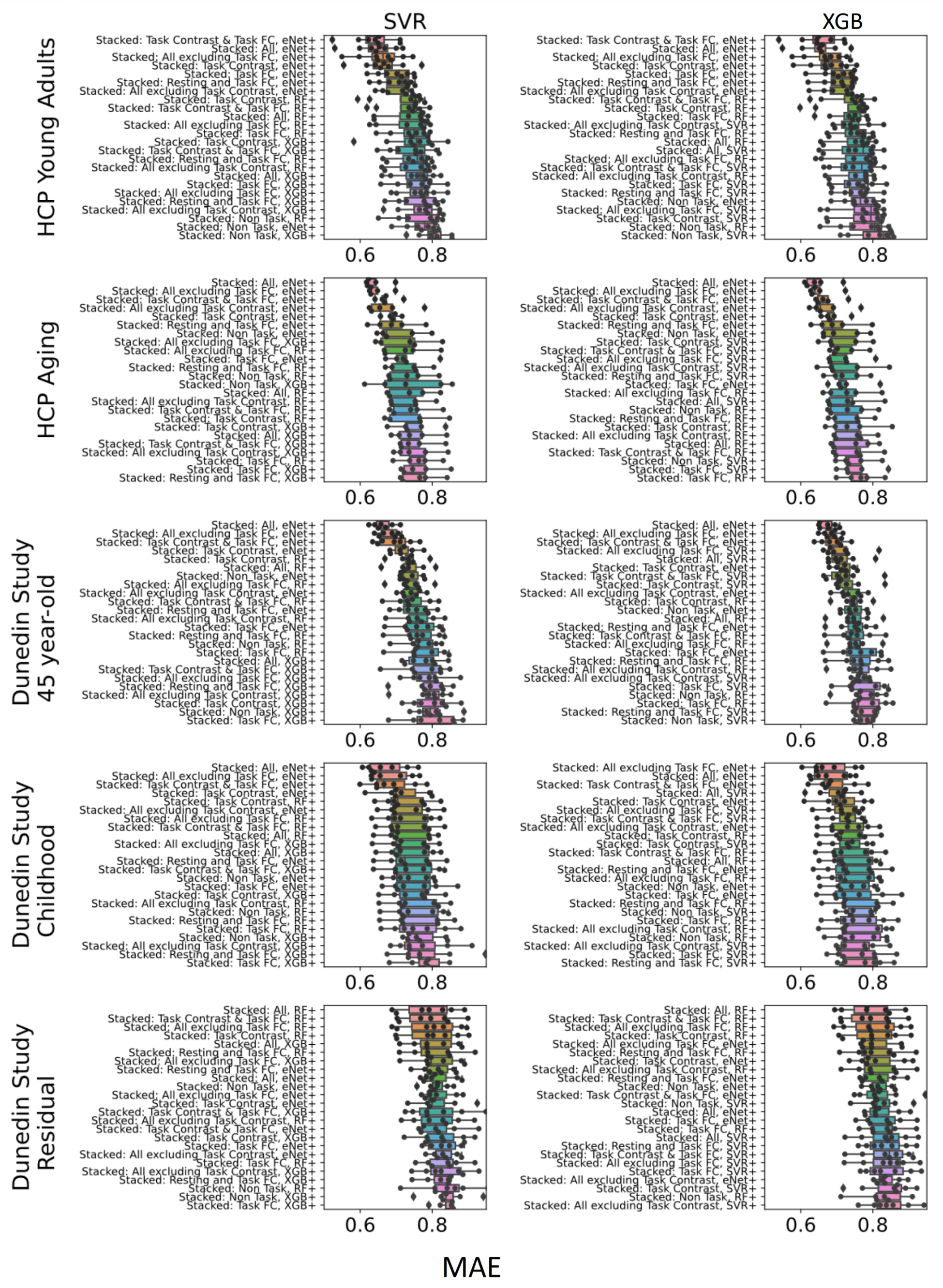

**Figure S9**. **Predictability (mean absolute error, MAE) of stacked models having different predictive algorithms between layers with Support Vector Regression or XGBoost** **at the second layer (indicated by top labels).** Lower is better. The predictive-modelling algorithms at the first layer are indicated by left-side labels. Each dot represents predictive performance at each outer-fold test set. For Dunedin Study, childhood scores reflect cognitive abilities, averaged across 7, 9 and 11 years old, and negative residual scores reflect a stronger decline in cognitive abilities, as expected from childhood cognitive abilities, compared to participants’ peers. eNet = Elastic Net; RF = Random Forest; SVR = Support Vector Regression; XGB = XGBoost; FC = Functional Connectivity.

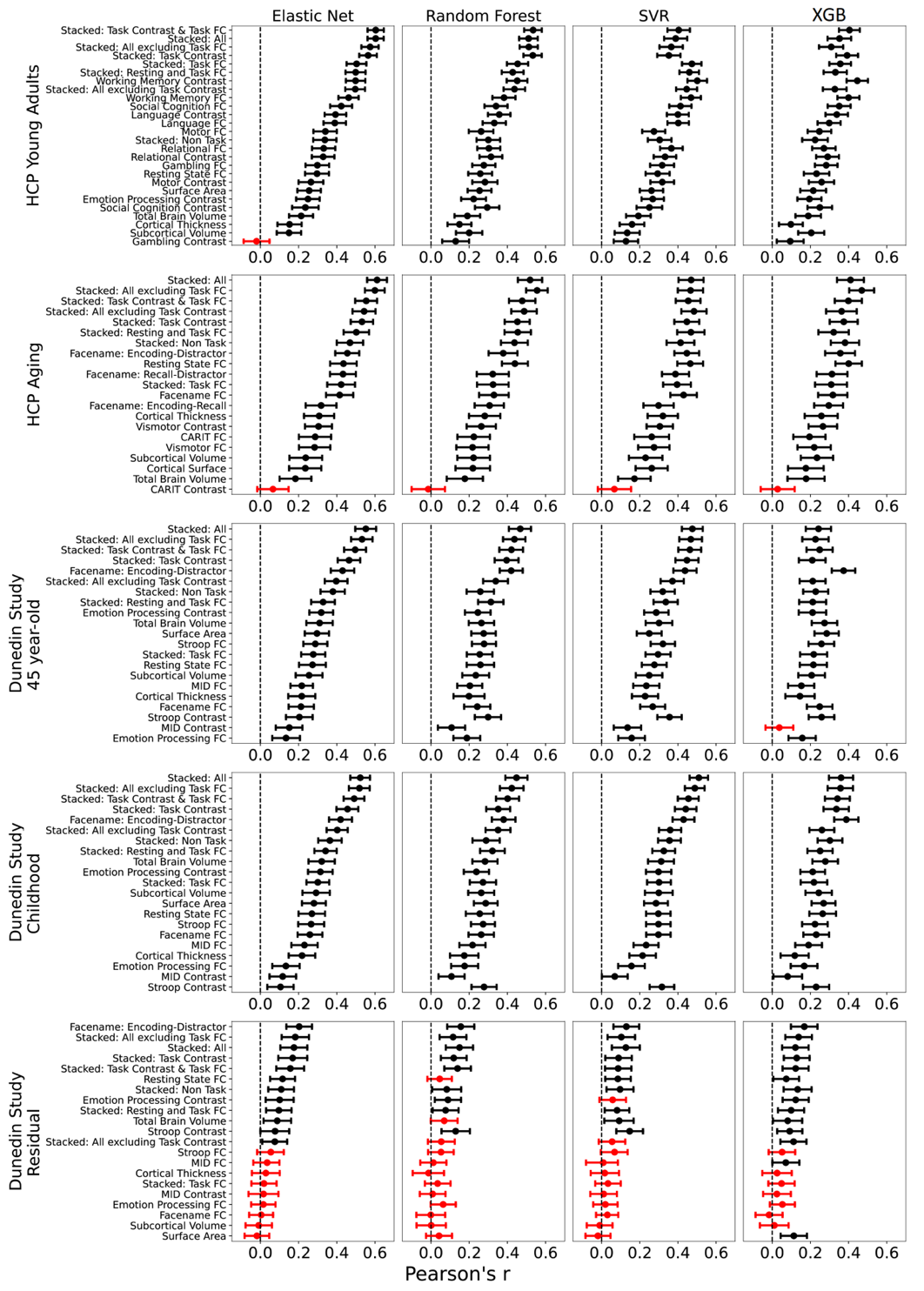

***Figure S10. Bootstrapped predictability (Pearson’s correlation, r) of stacked and non-stacked models for each predictive-modelling algorithm and dataset.*** *Higher is better. Each dot and bar represent the median and 95% confidence intervals (CI) of bootstrapped distributions, respectively. If 95% CI was higher than zero (indicated by the black colour), then predictability from a particular prediction model was better than chance. Note that, for the stacked models here, we only showed those with the same predictive algorithm across both layers. For Dunedin Study, childhood scores reflect cognitive abilities, averaged across 7, 9 and 11 years old, and negative residual scores reflect a stronger decline in cognitive abilities, as expected from childhood cognitive abilities, compared to participants’ peers. SVR = Support Vector Regression; XGB = XGBoost; FC = Functional Connectivity.*

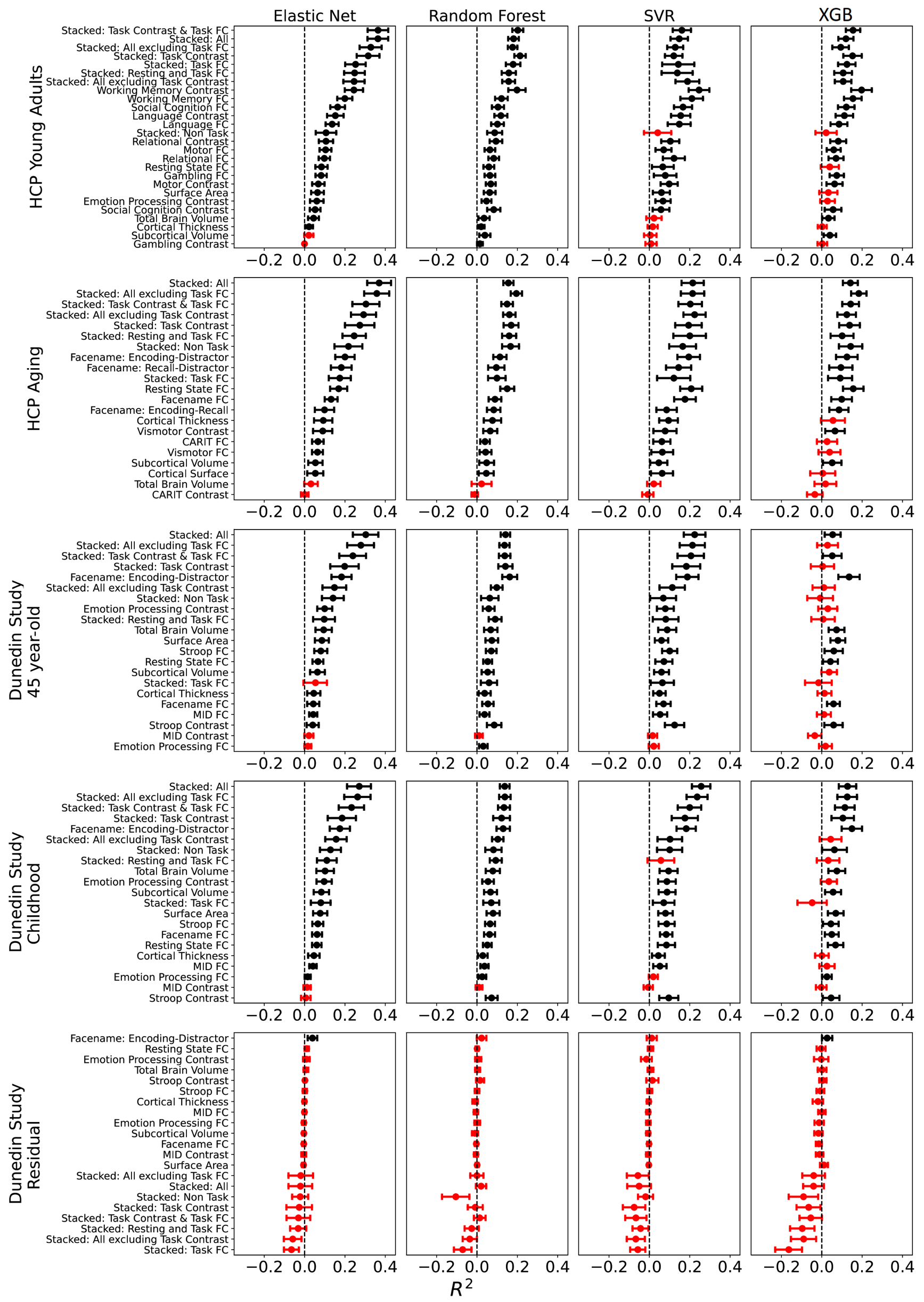

**Figure S11. Bootstrapped predictability (coefficient of determination, *R*^2^) of stacked and non-stacked models for each predictive-modelling algorithm and dataset.** Higher is better. Each dot and bar represent the median and 95% confidence intervals (CI) of bootstrapped distributions, respectively. If 95% CI was higher than zero (indicated by the black colour), then predictability from a particular prediction model was better than chance. Note that, for the stacked models here, we only showed those with the same predictive algorithm across both layers. For stacked models with different predictive algorithms between layers, please see Figures S11-S16. For Dunedin Study, childhood scores reflect cognitive abilities, averaged across 7, 9 and 11 years old, and negative residual scores reflect a stronger decline in cognitive abilities, as expected from childhood cognitive abilities, compared to participants’ peers. SVR = Support Vector Regression; XGB = XGBoost; FC = Functional Connectivity.

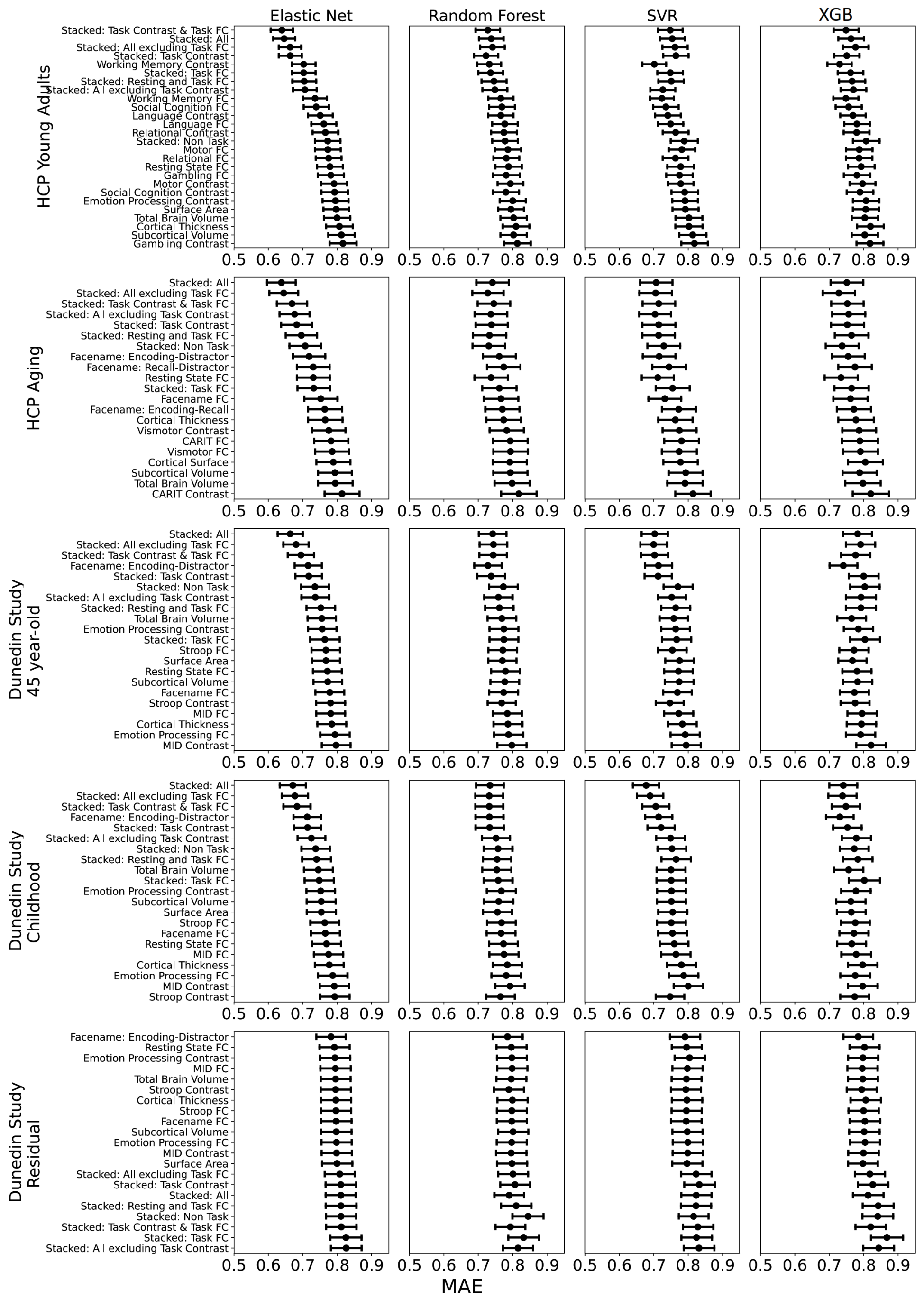

**Figure S12. Bootstrapped predictability (mean absolute error, MAE) of stacked and non-stacked models for each predictive-modelling algorithm and dataset.** Lower is better. Each dot and bar represent the median and 95% confidence intervals (CI) of bootstrapped distributions, respectively. Note that, for the stacked models here, we only showed those with the same predictive algorithm across both layers. For stacked models with different predictive algorithms between layers, please see Figures S11-S16. For Dunedin Study, childhood scores reflect cognitive abilities, averaged across 7, 9 and 11 years old, and negative residual scores reflect a stronger decline in cognitive abilities, as expected from childhood cognitive abilities, compared to participants’ peers. SVR = Support Vector Regression; XGB = XGBoost; FC = Functional Connectivity.

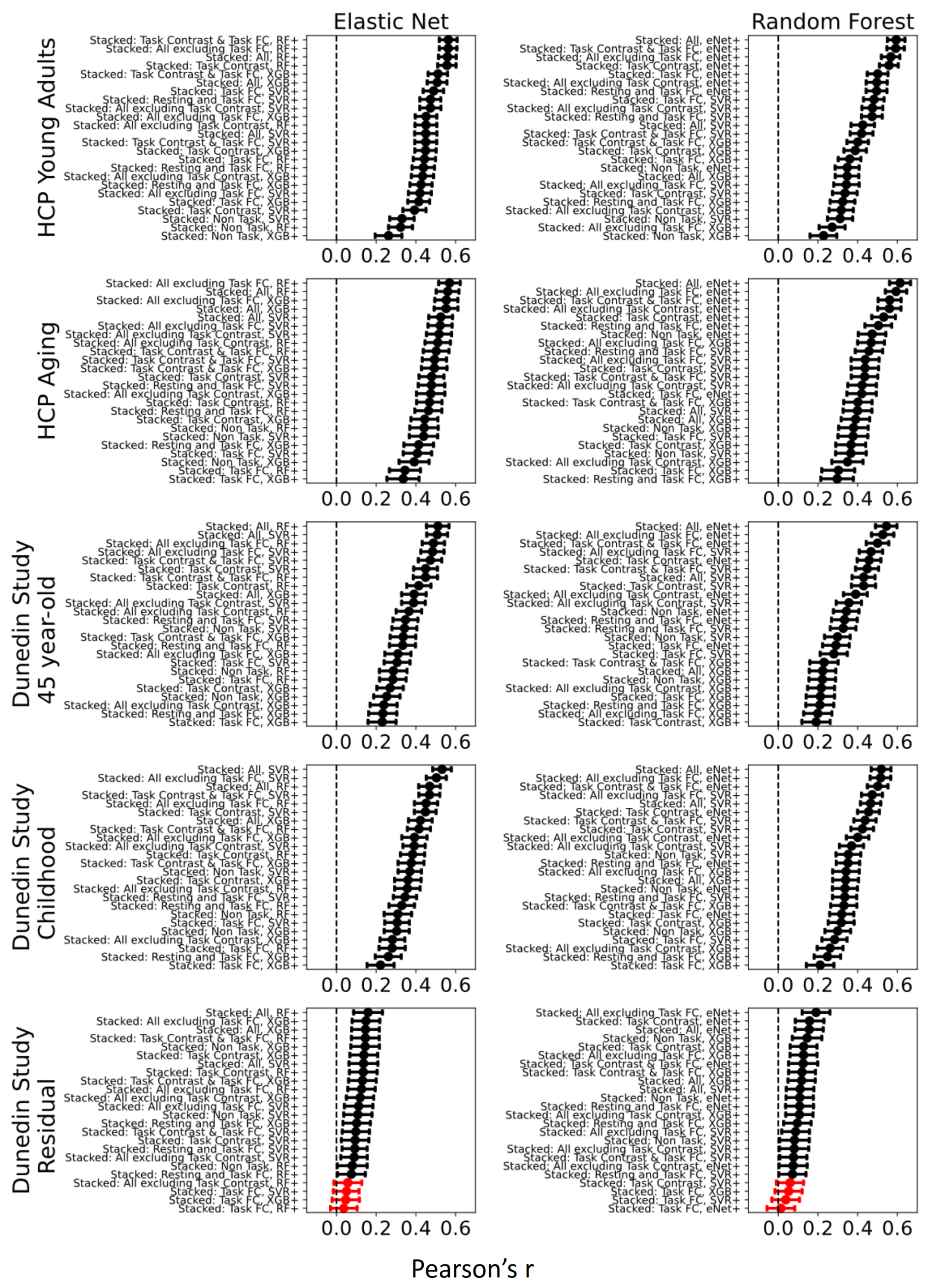

**Figure S13**. **Bootstrapped predictability (Pearson’s correlation, *r*) of stacked models having different predictive algorithms between layers with Elastic Net or Random Forest at the second layer (indicated by top labels).** Higher is better. The predictive-modelling algorithms at the first layer are indicated by left-side labels. Each dot and bar represent the median and 95% confidence intervals (CI) of bootstrapped distributions, respectively. If 95% CI was higher than zero (indicated by the black colour), then predictability from a particular prediction model was better than chance. For Dunedin Study, childhood scores reflect cognitive abilities, averaged across 7, 9 and 11 years old, and negative residual scores reflect a stronger decline in cognitive abilities, as expected from childhood cognitive abilities, compared to participants’ peers. eNet = Elastic Net; RF = Random Forest; SVR = Support Vector Regression; XGB = XGBoost; FC = Functional Connectivity.

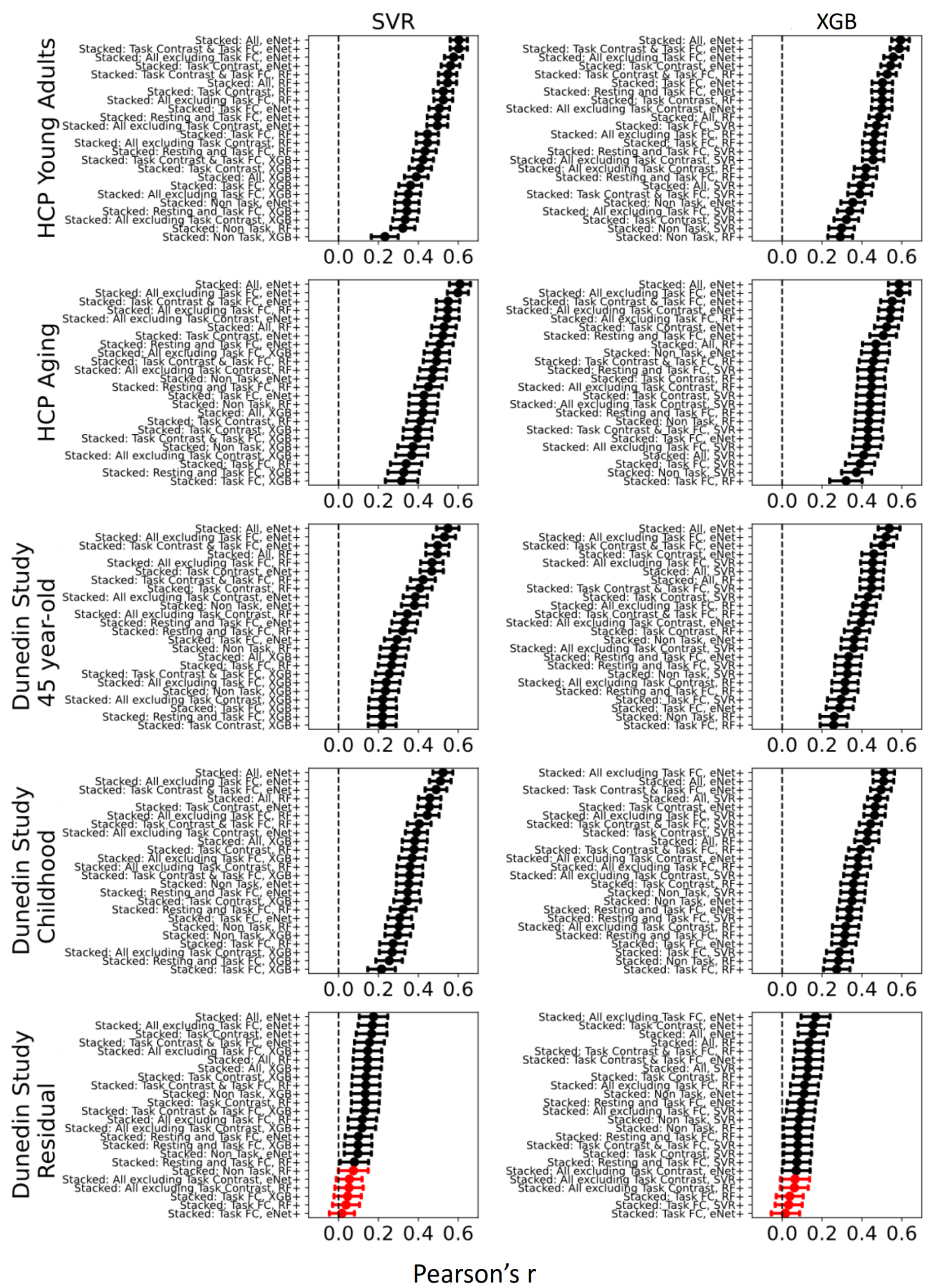

**Figure S14**. **Bootstrapped predictability (Pearson’s correlation, *r*) of stacked models having different predictive algorithms between layers with Support Vector Regression or XGBoost at the second layer (indicated by top labels).** Higher is better. The predictive-modelling algorithms at the first layer are indicated by left-side labels. Each dot and bar represent the median and 95% confidence intervals (CI) of bootstrapped distributions, respectively. If 95% CI was higher than zero (indicated by the black colour), then predictability from a particular prediction model was better than chance. For Dunedin Study, childhood scores reflect cognitive abilities, averaged across 7, 9 and 11 years old, and negative residual scores reflect a stronger decline in cognitive abilities, as expected from childhood cognitive abilities, compared to participants’ peers. eNet = Elastic Net; RF = Random Forest; SVR = Support Vector Regression; XGB = XGBoost; FC = Functional Connectivity.

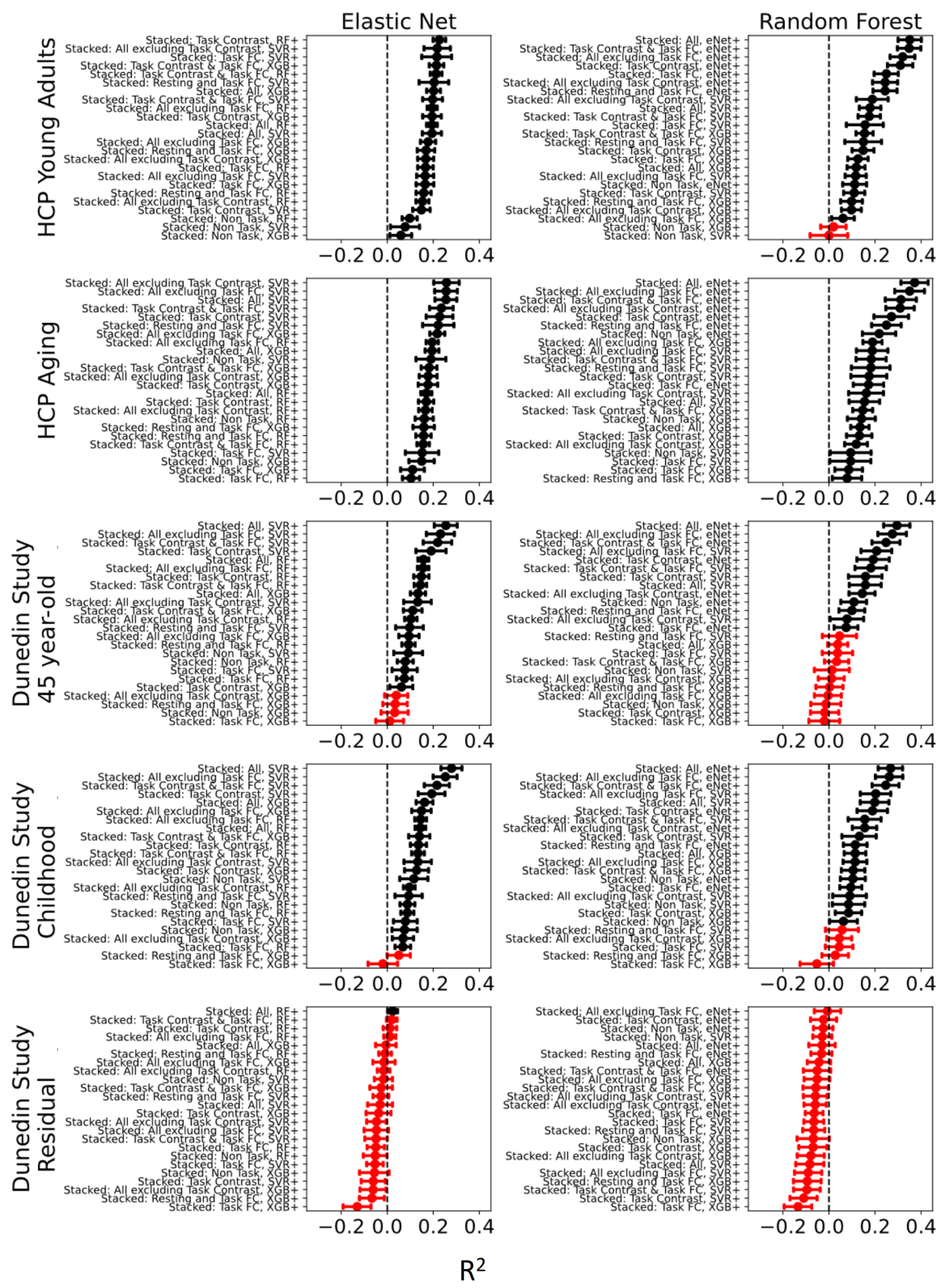

**Figure S15**. **Bootstrapped predictability (coefficient of determination, R^2^) of stacked models having different predictive algorithms between layers with Elastic Net or Random Forest at the second layer (indicated by top labels).** Higher is better. The predictive-modelling algorithms at the first layer are indicated by left-side labels. Each dot and bar represent the median and 95% confidence intervals (CI) of bootstrapped distributions, respectively. If 95% CI was higher than zero (indicated by the black colour), then predictability from a particular prediction model was better than chance. For Dunedin Study, childhood scores reflect cognitive abilities, averaged across 7, 9 and 11 years old, and negative residual scores reflect a stronger decline in cognitive abilities, as expected from childhood cognitive abilities, compared to participants’ peers. eNet = Elastic Net; RF = Random Forest; SVR = Support Vector Regression; XGB = XGBoost; FC = Functional Connectivity.

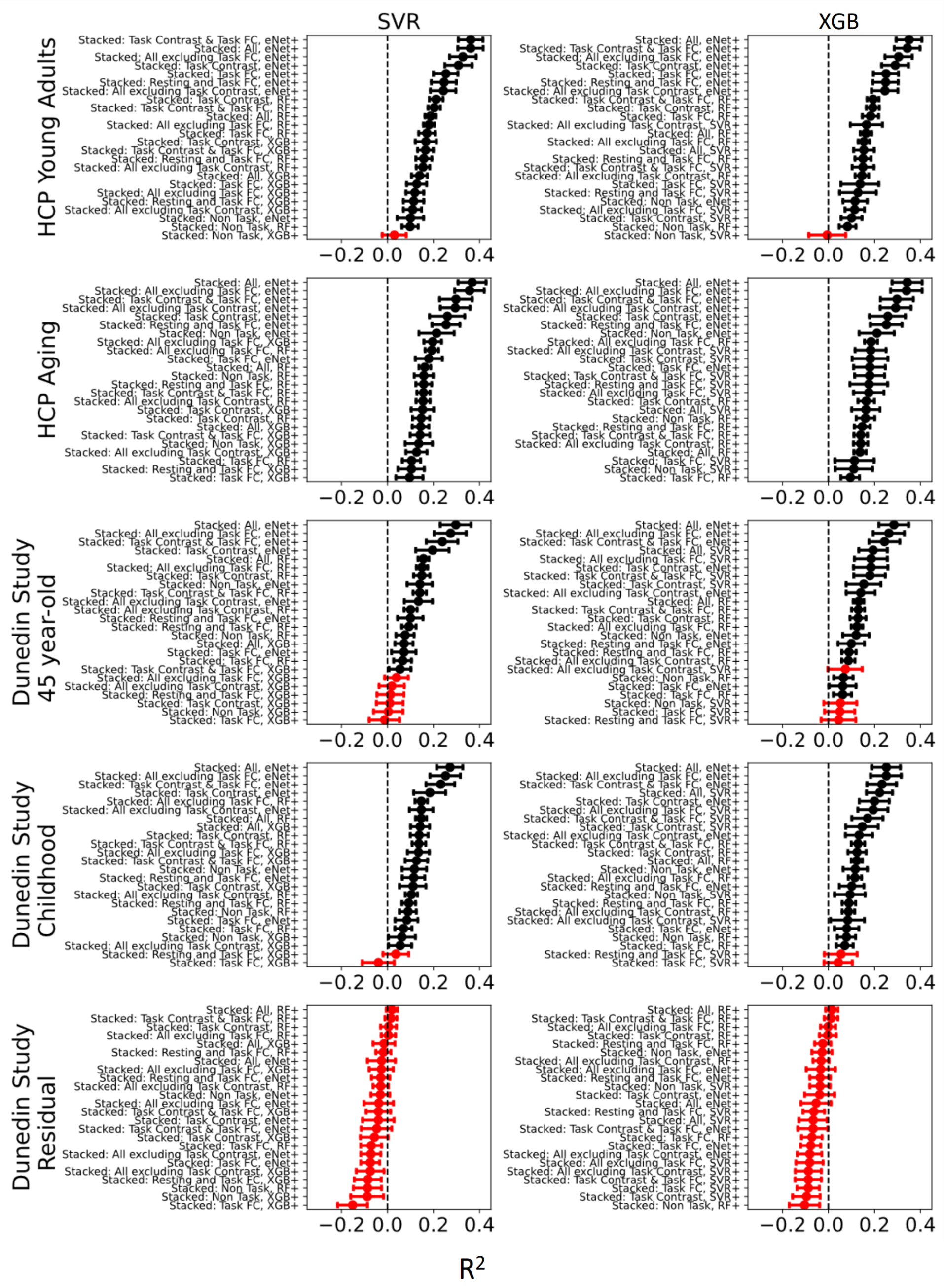

**Figure S16**. **Bootstrapped predictability (coefficient of determination, R^2^) of stacked models having different predictive algorithms between layers with Support Vector Regression or XGBoost at the second layer (indicated by top labels).** Higher is better. The predictive-modelling algorithms at the first layer are indicated by left-side labels. Each dot and bar represent the median and 95% confidence intervals (CI) of bootstrapped distributions, respectively. If 95% CI was higher than zero (indicated by the black colour), then predictability from a particular prediction model was better than chance. For Dunedin Study, childhood scores reflect cognitive abilities, averaged across 7, 9 and 11 years old, and negative residual scores reflect a stronger decline in cognitive abilities, as expected from childhood cognitive abilities, compared to participants’ peers. eNet = Elastic Net; RF = Random Forest; SVR = Support Vector Regression; XGB = XGBoost; FC = Functional Connectivity.

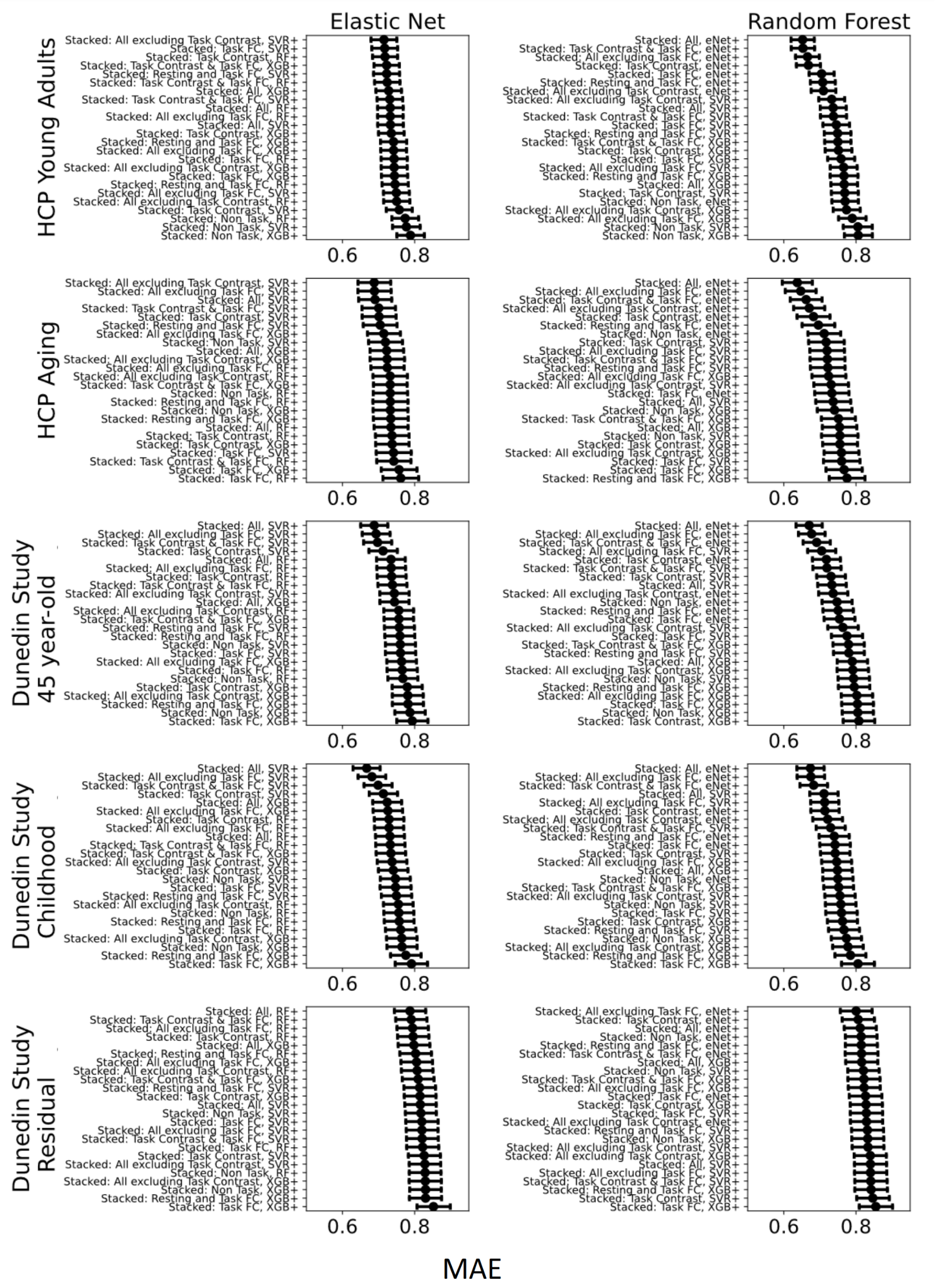

**Figure S17**. **Bootstrapped predictability (mean absolute error, MAE) of stacked models having different predictive algorithms between layers with Elastic Net or Random Forest at the second layer (indicated by top labels).** Lower is better. The predictive-modelling algorithms at the first layer are indicated by left-side labels. Each dot and bar represent the median and 95% confidence intervals (CI) of bootstrapped distributions, respectively. For Dunedin Study, childhood scores reflect cognitive abilities, averaged across 7, 9 and 11 years old, and negative residual scores reflect a stronger decline in cognitive abilities, as expected from childhood cognitive abilities, compared to participants’ peers. eNet = Elastic Net; RF = Random Forest; SVR = Support Vector Regression; XGB = XGBoost; FC = Functional Connectivity.

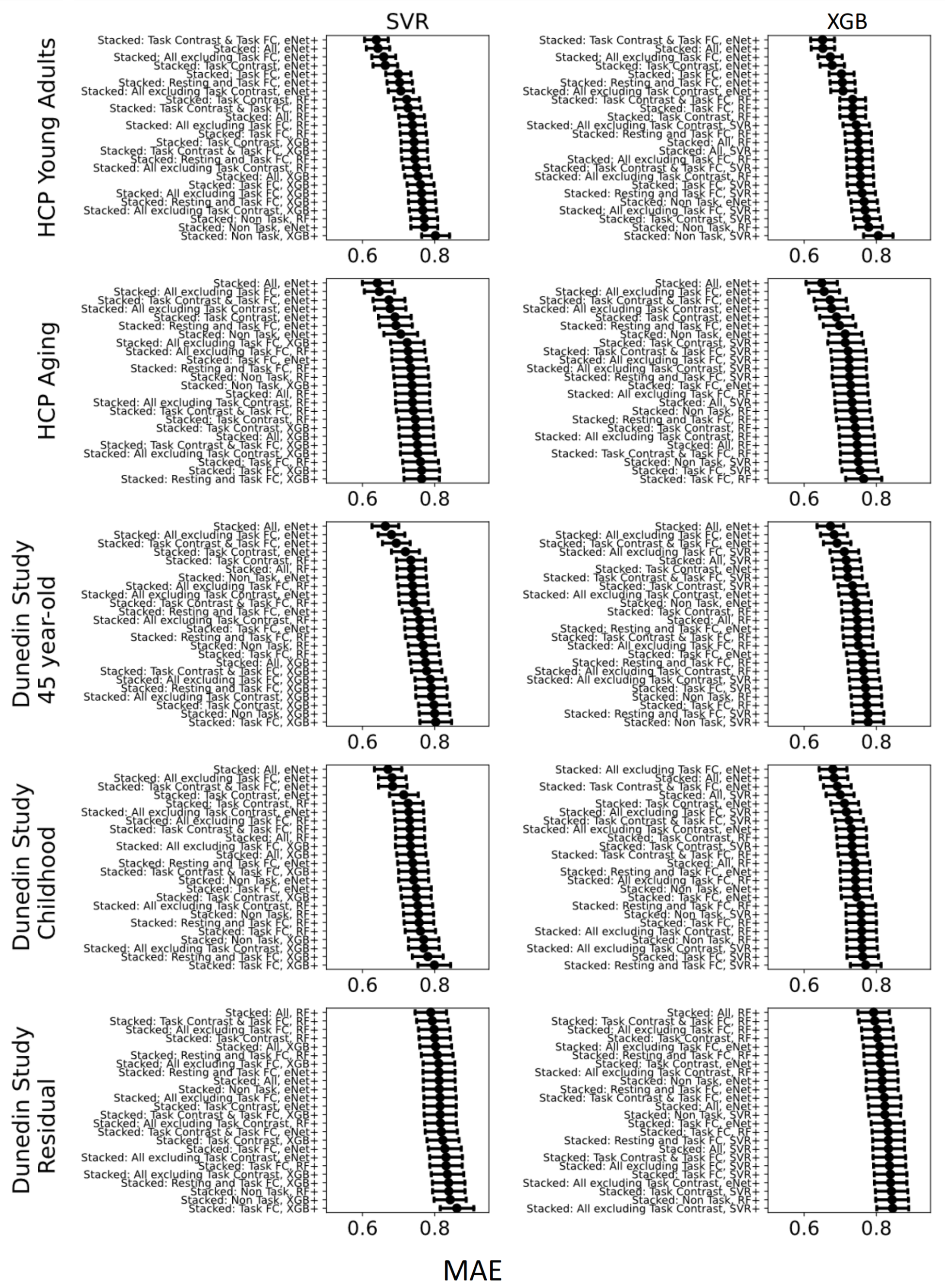

**Figure S18**. **Bootstrapped predictability (mean absolute error, MAE) of stacked models having different predictive algorithms between layers with Support Vector Regression or XGBoost at the second layer (indicated by top labels).** Lower is better. The predictive-modelling algorithms at the first layer are indicated by left-side labels. Each dot and bar represent the median and 95% confidence intervals (CI) of bootstrapped distributions, respectively. For Dunedin Study, childhood scores reflect cognitive abilities, averaged across 7, 9 and 11 years old, and negative residual scores reflect a stronger decline in cognitive abilities, as expected from childhood cognitive abilities, compared to participants’ peers. eNet = Elastic Net; RF = Random Forest; SVR = Support Vector Regression; XGB = XGBoost; FC = Functional Connectivity.

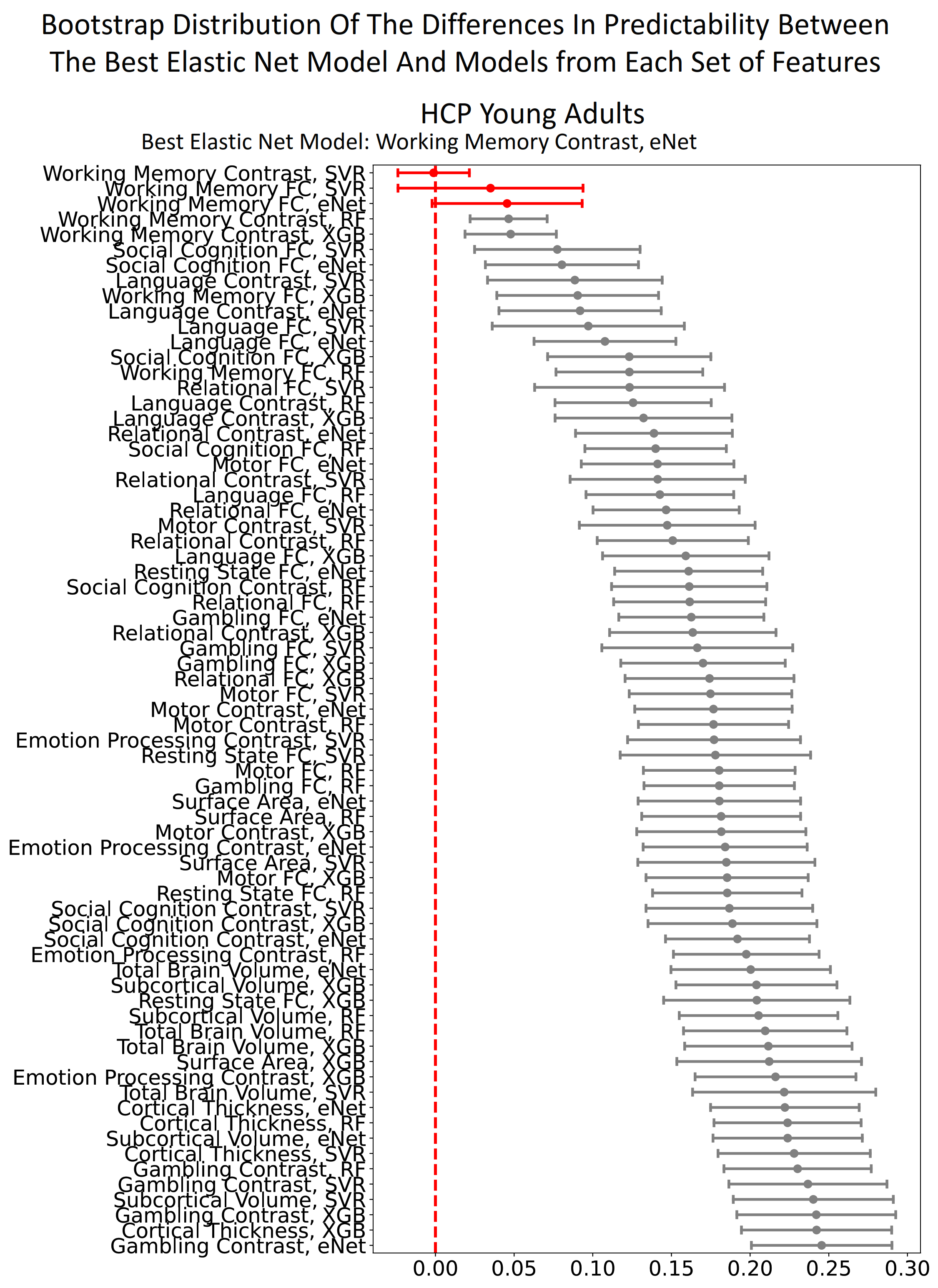

**Figure S19. Bootstrapped distribution of the differences in predictability (Pearson’s correlation, *r*) between the best non-stacked model with Elastic Net, “Working Memory task contrast”, and other non-stacked models in HCP Young Adults.** Each dot and bar represent the median and 95% confidence intervals (CI) of bootstrapped distributions, respectively. If 95% CI was higher than zero (indicated by the black colour), then the predictability of the best non-stacked model with Elastic Net was significantly better than the comparing model. eNet = Elastic Net; RF = Random Forest; SVR = Support Vector Regression; XGB = XGBoost; FC = Functional Connectivity.

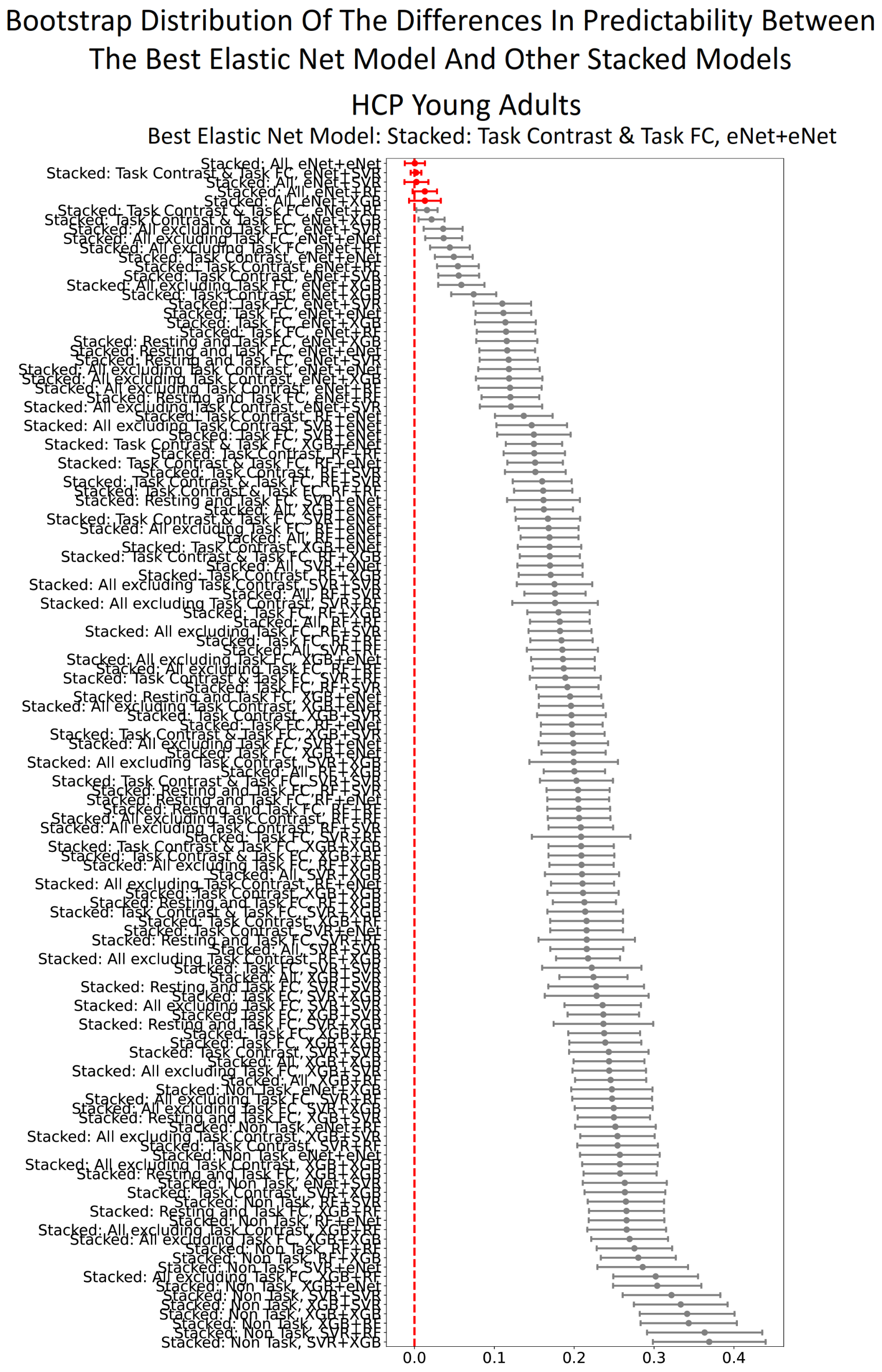

**Figure S20. Bootstrapped distribution of the differences in predictability (Pearson’s correlation, *r*) between the best stacked model with Elastic Net, “Stacked: Task Contrast & Task FC”, and other non-stacked models in HCP Young Adults.** Each dot and bar represent the median and 95% confidence intervals (CI) of bootstrapped distributions, respectively. If 95% CI was higher than zero (indicated by the black colour), then the predictability of the best stacked model with Elastic Net was significantly better than the comparing model. eNet = Elastic Net; RF = Random Forest; SVR = Support Vector Regression; XGB = XGBoost; FC = Functional Connectivity.

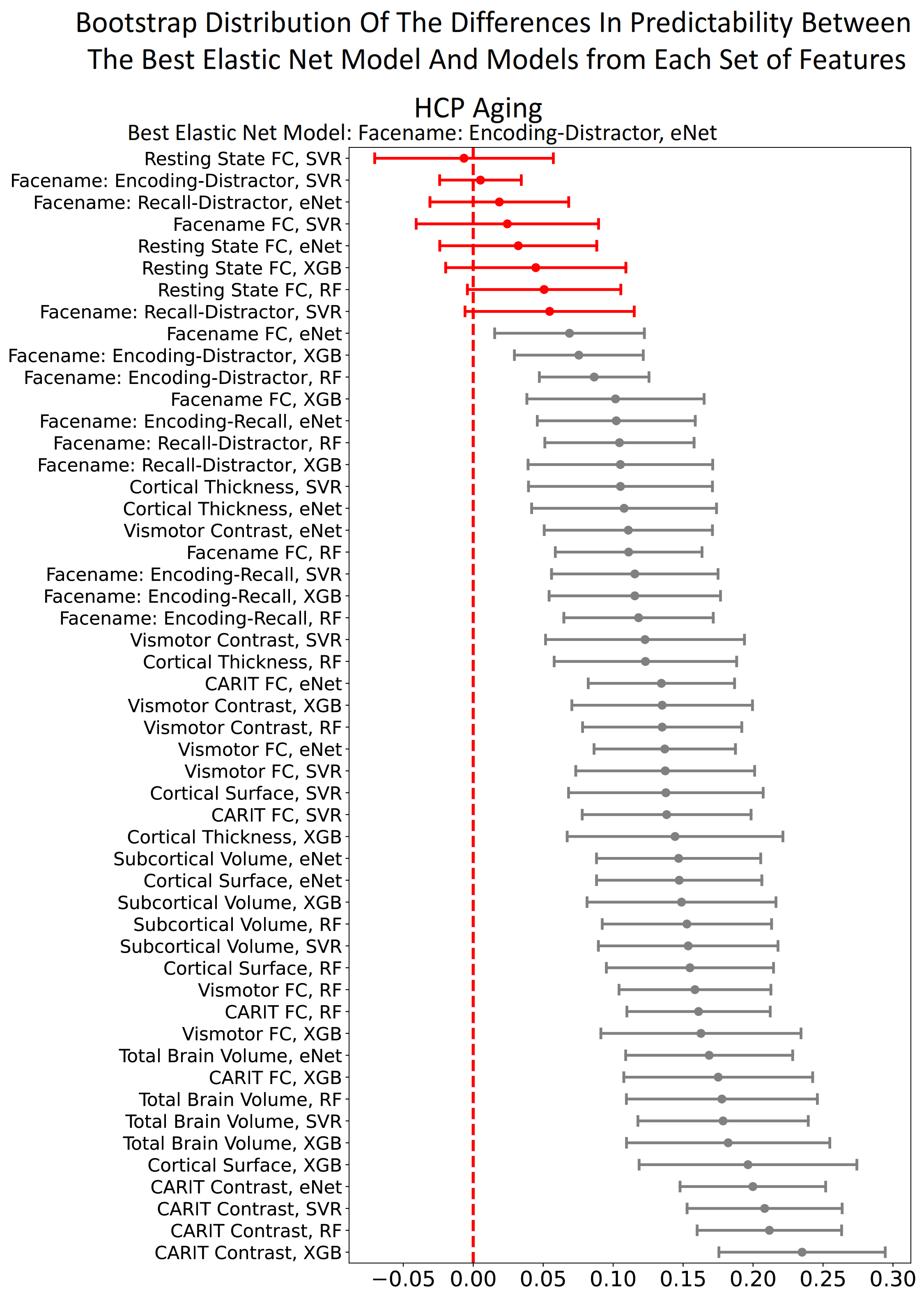

**Figure S21. Bootstrapped distribution of the differences in predictability (Pearson’s correlation, *r*) between the best non-stacked model with Elastic Net, “Face-name: Encoding vs Distractor task contrast”, and other non-stacked models in HCP Aging.** Each dot and bar represent the median and 95% confidence intervals (CI) of bootstrapped distributions, respectively. If 95% CI was higher than zero (indicated by the black colour), then the predictability of the best non-stacked model with Elastic Net was significantly better than the comparing model. eNet = Elastic Net; RF = Random Forest; SVR = Support Vector Regression; XGB = XGBoost; FC = Functional Connectivity.

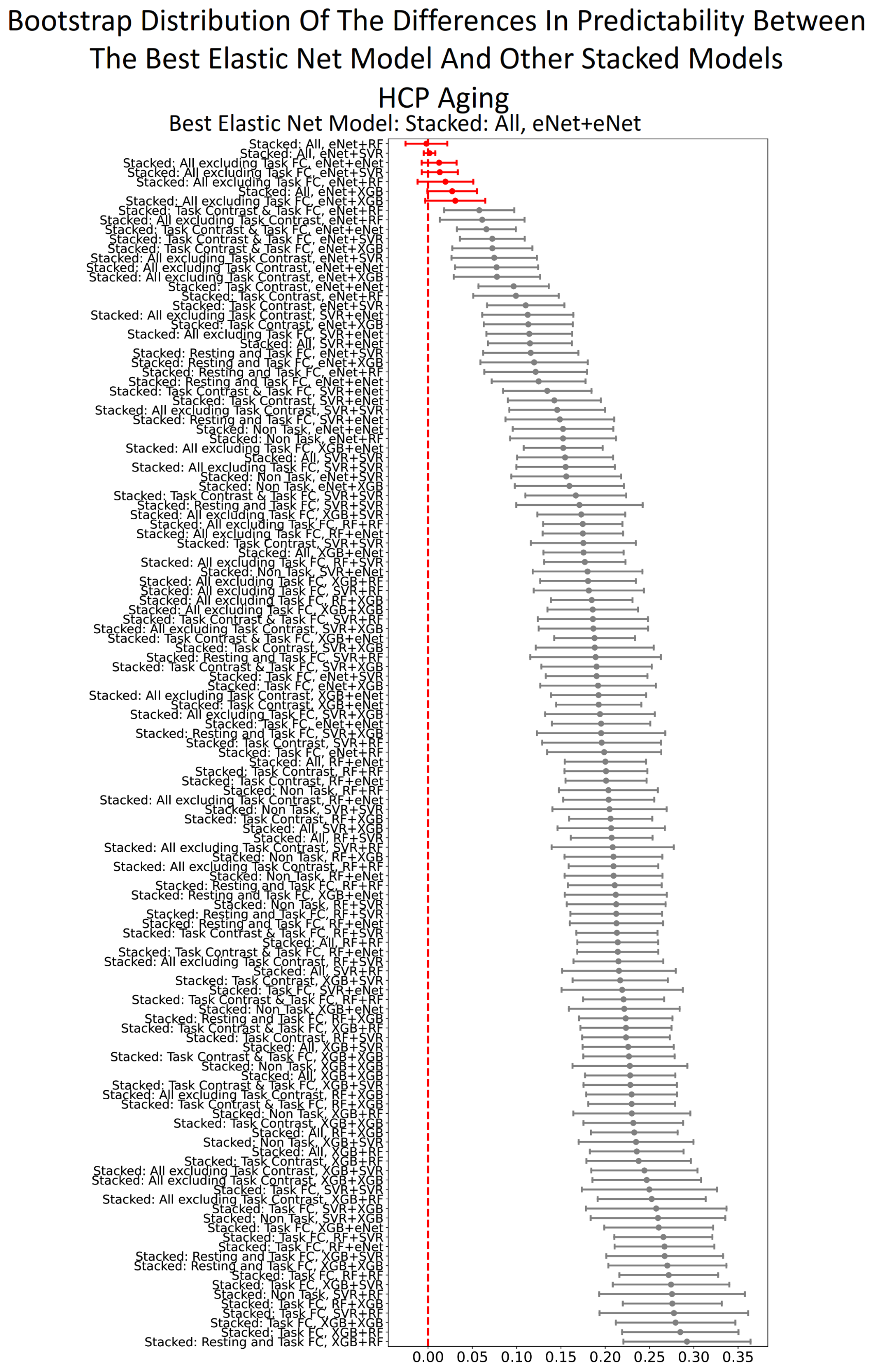

**Figure S22. Bootstrapped distribution of the differences in predictability (Pearson’s correlation, *r*) between the best stacked model with Elastic Net, “Stacked: All”, and other stacked models in HCP Aging.** Each dot and bar represent the median and 95% confidence intervals (CI) of bootstrapped distributions, respectively. If 95% CI was higher than zero (indicated by the black colour), then the predictability of the best stacked model with Elastic Net was significantly better than the comparing model. eNet = Elastic Net; RF = Random Forest; SVR = Support Vector Regression; XGB = XGBoost; FC = Functional Connectivity.

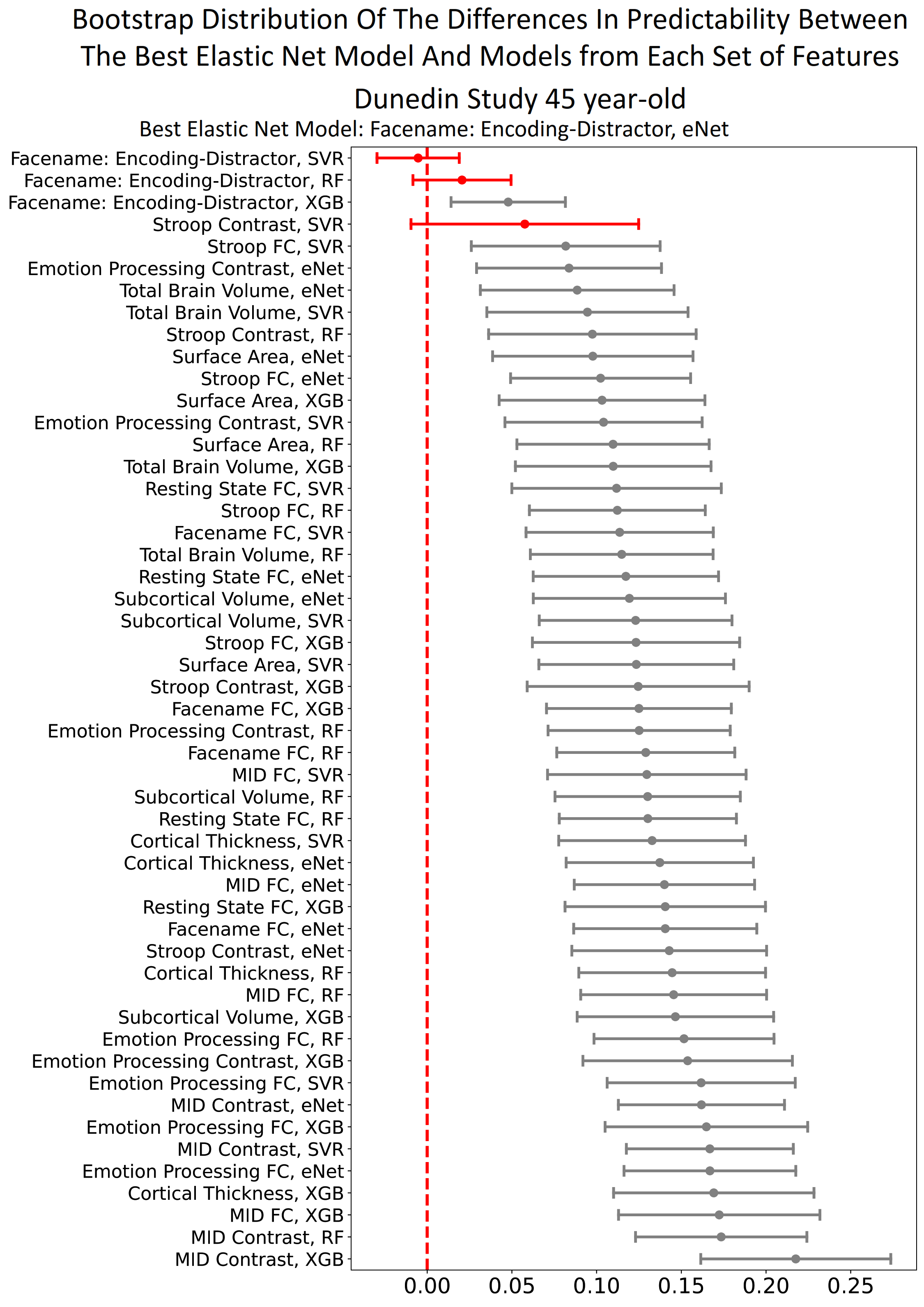

**Figure S23. Bootstrapped distribution of the differences in predictability (Pearson’s correlation, *r*) between the best non-stacked model with Elastic Net, “Face-name: Encoding vs Distractor task contrast”, and other non-stacked models in Dunedin Study when predicting cognitive abilities at the same time of MRI scanning.** Each dot and bar represent the median and 95% confidence intervals (CI) of bootstrapped distributions, respectively. If 95% CI was higher than zero (indicated by the black colour), then the predictability of the best non-stacked model with Elastic Net was significantly better than the comparing model. eNet = Elastic Net; RF = Random Forest; SVR = Support Vector Regression; XGB = XGBoost; FC = Functional Connectivity.

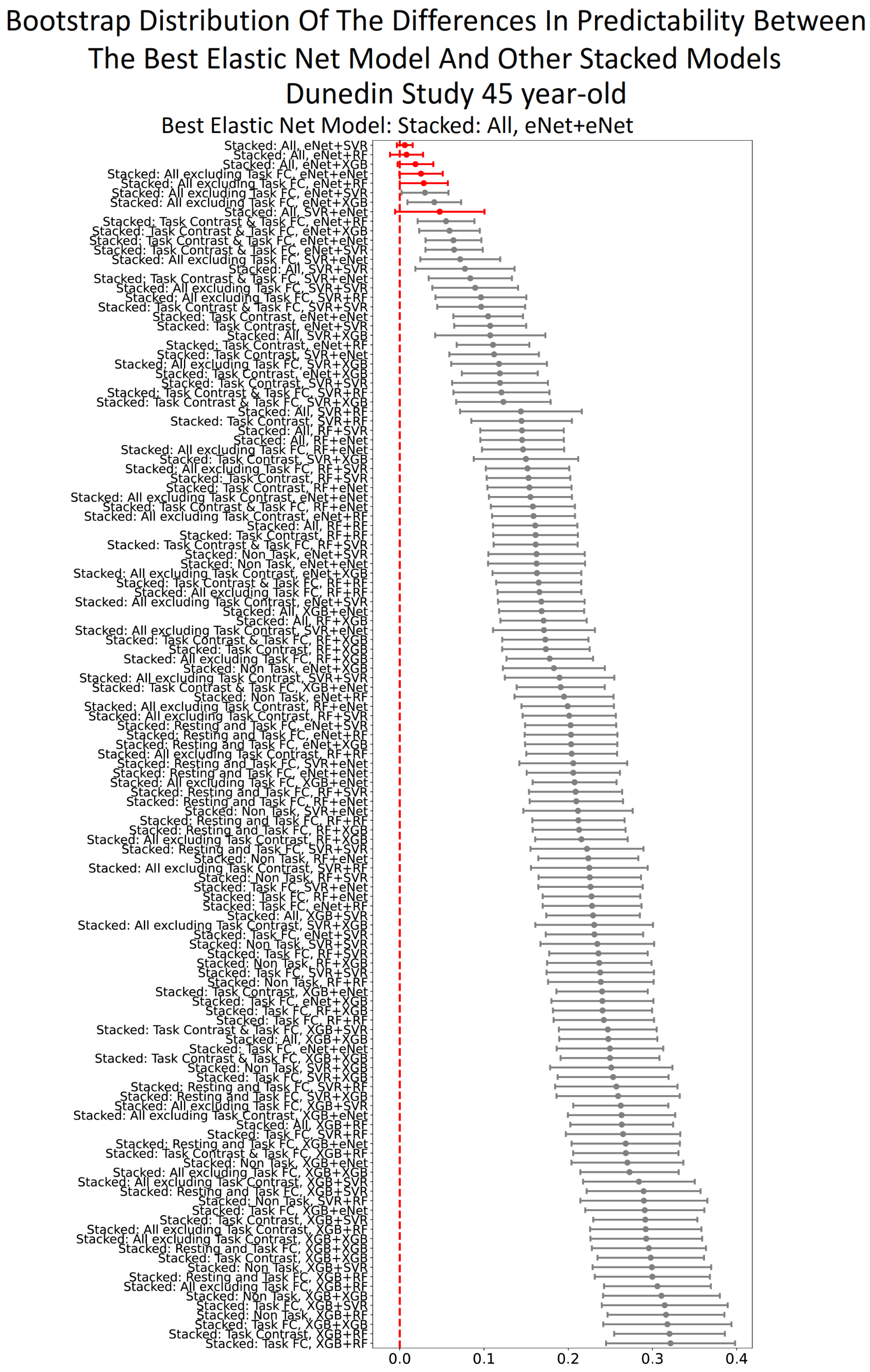

**Figure S24. Bootstrapped distribution of the differences in predictability (Pearson’s correlation, *r*) between the best stacked model with Elastic Net, “Stacked: All”, and other stacked models in Dunedin Study when predicting cognitive abilities at the same time of MRI scanning.** Each dot and bar represent the median and 95% confidence intervals (CI) of bootstrapped distributions, respectively. If 95% CI was higher than zero (indicated by the black colour), then the predictability of the best non-stacked model with Elastic Net was significantly better than the comparing model. eNet = Elastic Net; RF = Random Forest; SVR = Support Vector Regression; XGB = XGBoost; FC = Functional Connectivity.

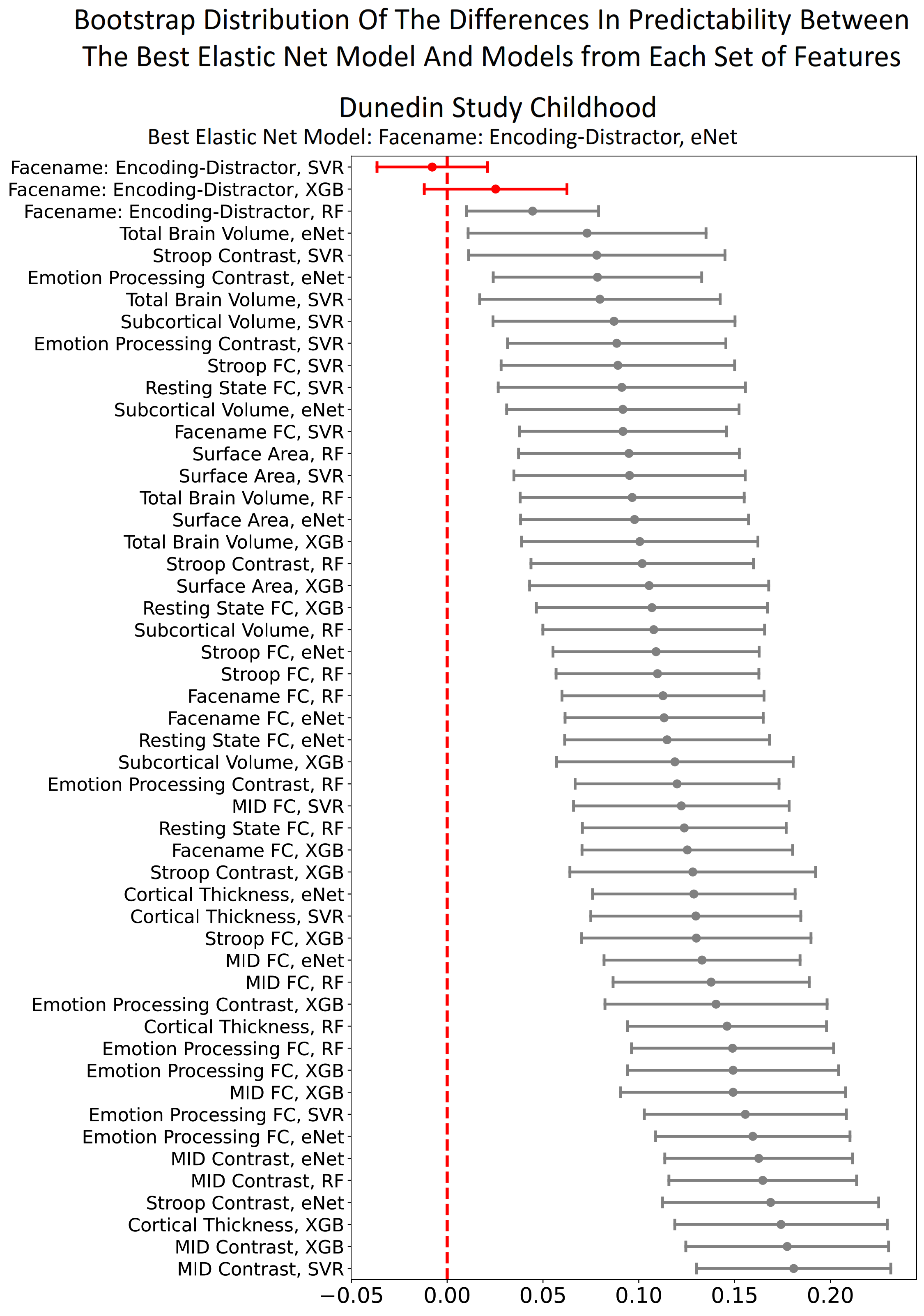

**Figure S25. Bootstrapped distribution of the differences in predictability (Pearson’s correlation, *r*) between the best non-stacked model with Elastic Net, “Face-name: Encoding vs Distractor task contrast”, and other non-stacked models in Dunedin Study when predicting cognitive abilities earlier in life (7, 9 and 11 years).** Each dot and bar represent the median and 95% confidence intervals (CI) of bootstrapped distributions, respectively. If 95% CI was higher than zero (indicated by the black colour), then the predictability of the best non-stacked model with Elastic Net was significantly better than the comparing model. eNet = Elastic Net; RF = Random Forest; SVR = Support Vector Regression; XGB = XGBoost; FC = Functional Connectivity.

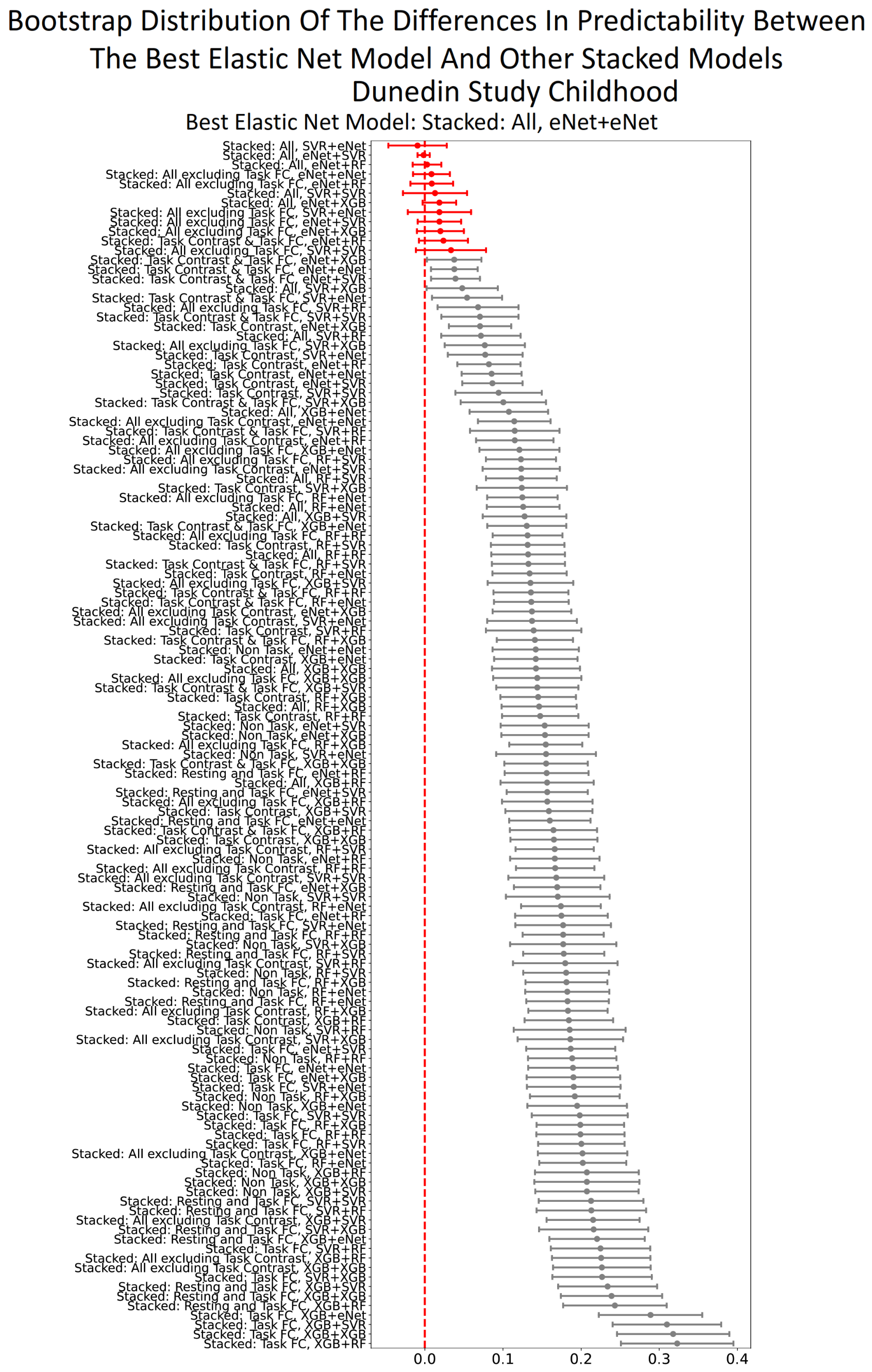

**Figure S26. Bootstrapped distribution of the differences in predictability (Pearson’s correlation, *r*) between the best stacked model with Elastic Net, “Stacked: All”, and other stacked models in Dunedin Study when predicting cognitive abilities earlier in life (7, 9 and 11 years).** Each dot and bar represent the median and 95% confidence intervals (CI) of bootstrapped distributions, respectively. If 95% CI was higher than zero (indicated by the black colour), then the predictability of the best stacked model with Elastic Net was significantly better than the comparing model. eNet = Elastic Net; RF = Random Forest; SVR = Support Vector Regression; XGB = XGBoost; FC = Functional Connectivity.

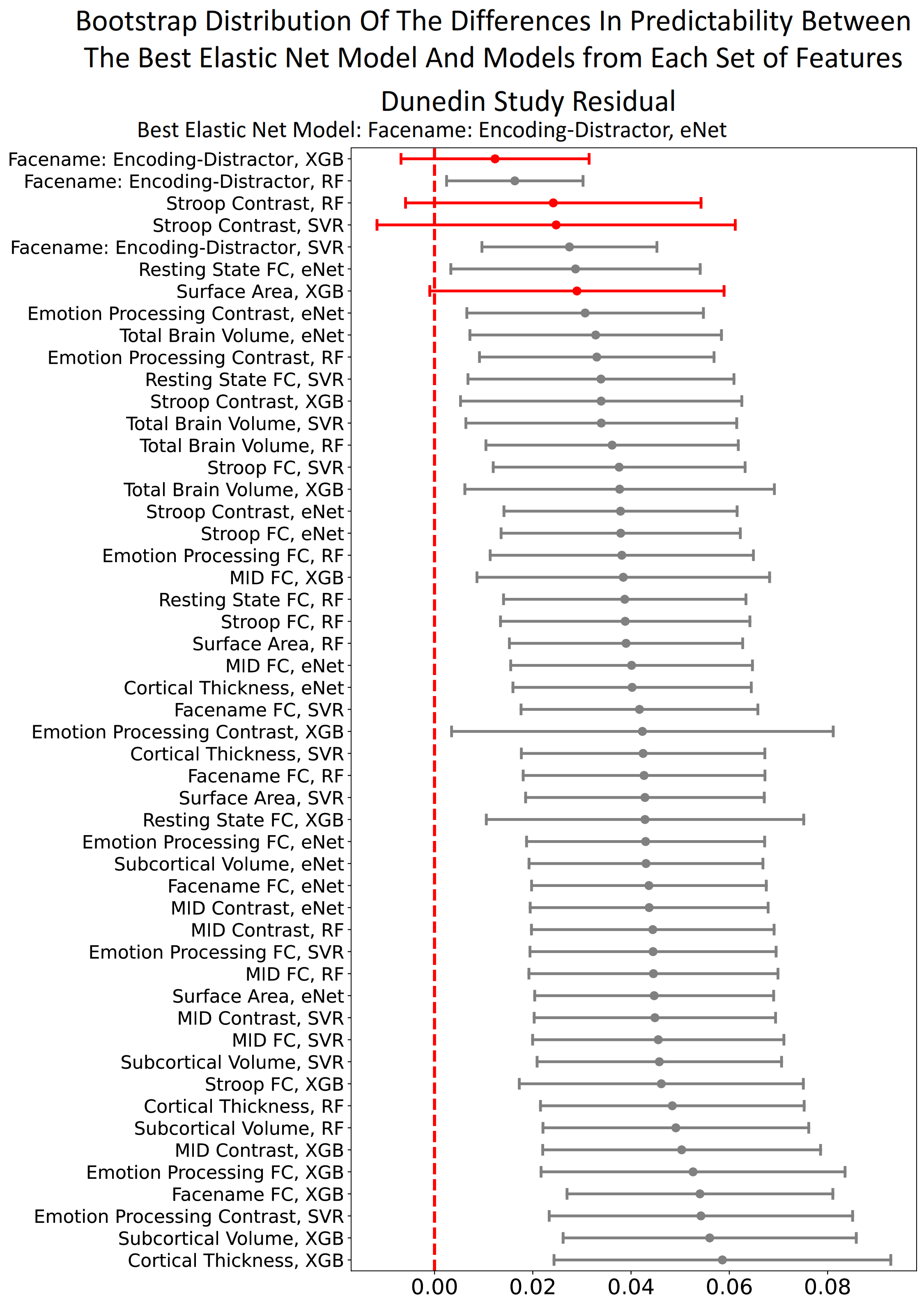

**Figure S27. Bootstrapped distribution of the differences in predictability (Pearson’s correlation, *r*) between the best non-stacked model with Elastic Net, “Face-name: Encoding vs Distractor task contrast”, and other non-stacked models in Dunedin Study when predicting the residual scores for cognitive abilities.** Each dot and bar represent the median and 95% confidence intervals (CI) of bootstrapped distributions, respectively. If 95% CI was higher than zero (indicated by the black colour), then the predictability of the best non-stacked model with Elastic Net was significantly better than the comparing model. eNet = Elastic Net; RF = Random Forest; SVR = Support Vector Regression; XGB = XGBoost; FC = Functional Connectivity.

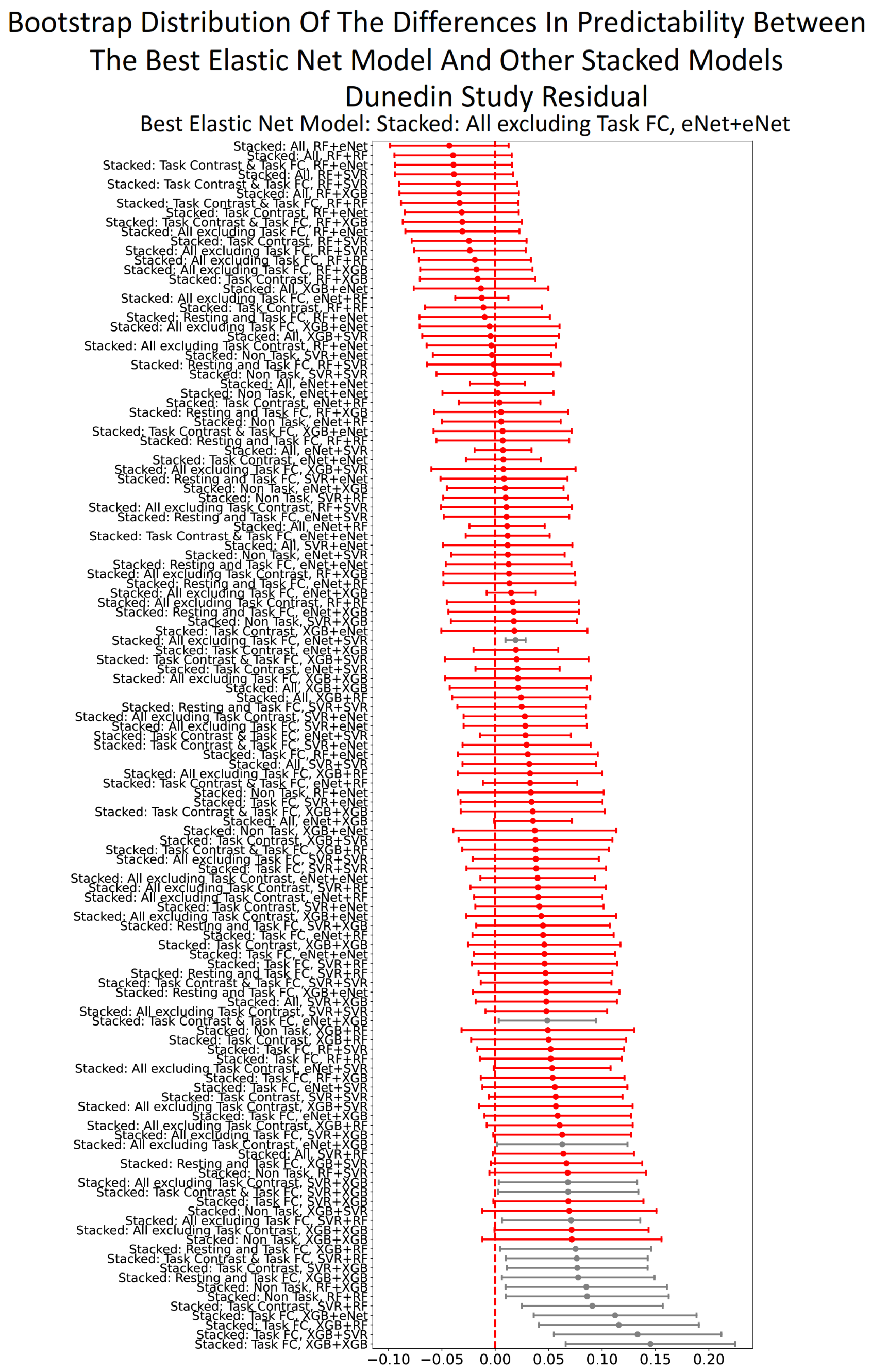

**Figure S28. Bootstrapped distribution of the differences in predictability (Pearson’s correlation, *r*) between the best stacked model with Elastic Net, “Stacked: All excluding Task FC”, and other stacked models in Dunedin Study when predicting the residual scores for cognitive abilities.** Each dot and bar represent the median and 95% confidence intervals (CI) of bootstrapped distributions, respectively. If 95% CI was higher than zero (indicated by the black colour), then the predictability of the best stacked model with Elastic Net was significantly better than the comparing model. eNet = Elastic Net; RF = Random Forest; SVR = Support Vector Regression; XGB = XGBoost; FC = Functional Connectivity.

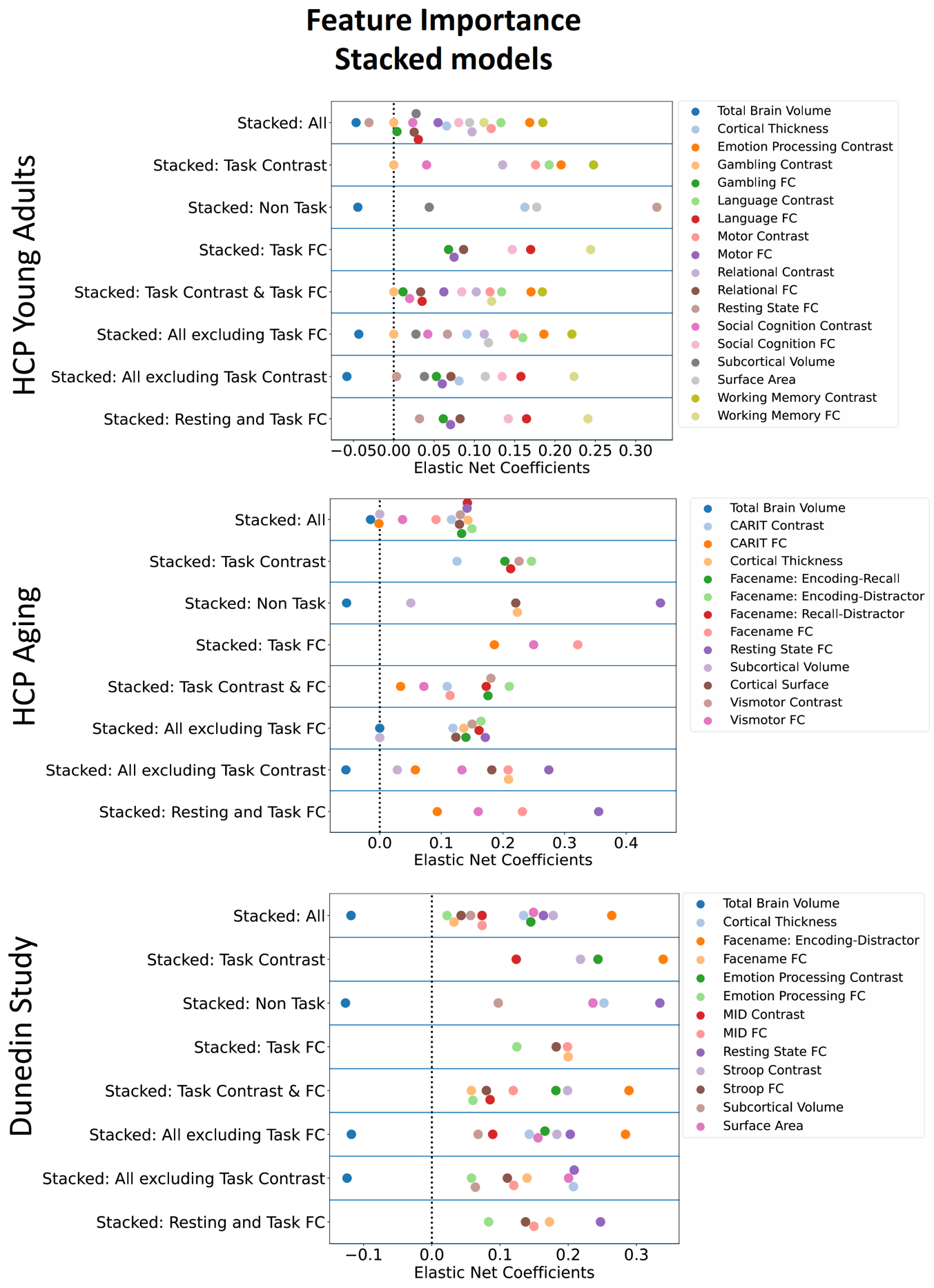

***Figure S29. Feature importance of stacked models with Elastic Net, indicated by Elastic Net Coefficients, for each dataset, when predicting cognitive abilities at the time of scanning.*** *A higher magnitude of a coefficient indicates a stronger contribution to the prediction. FC = Functional Connectivity*

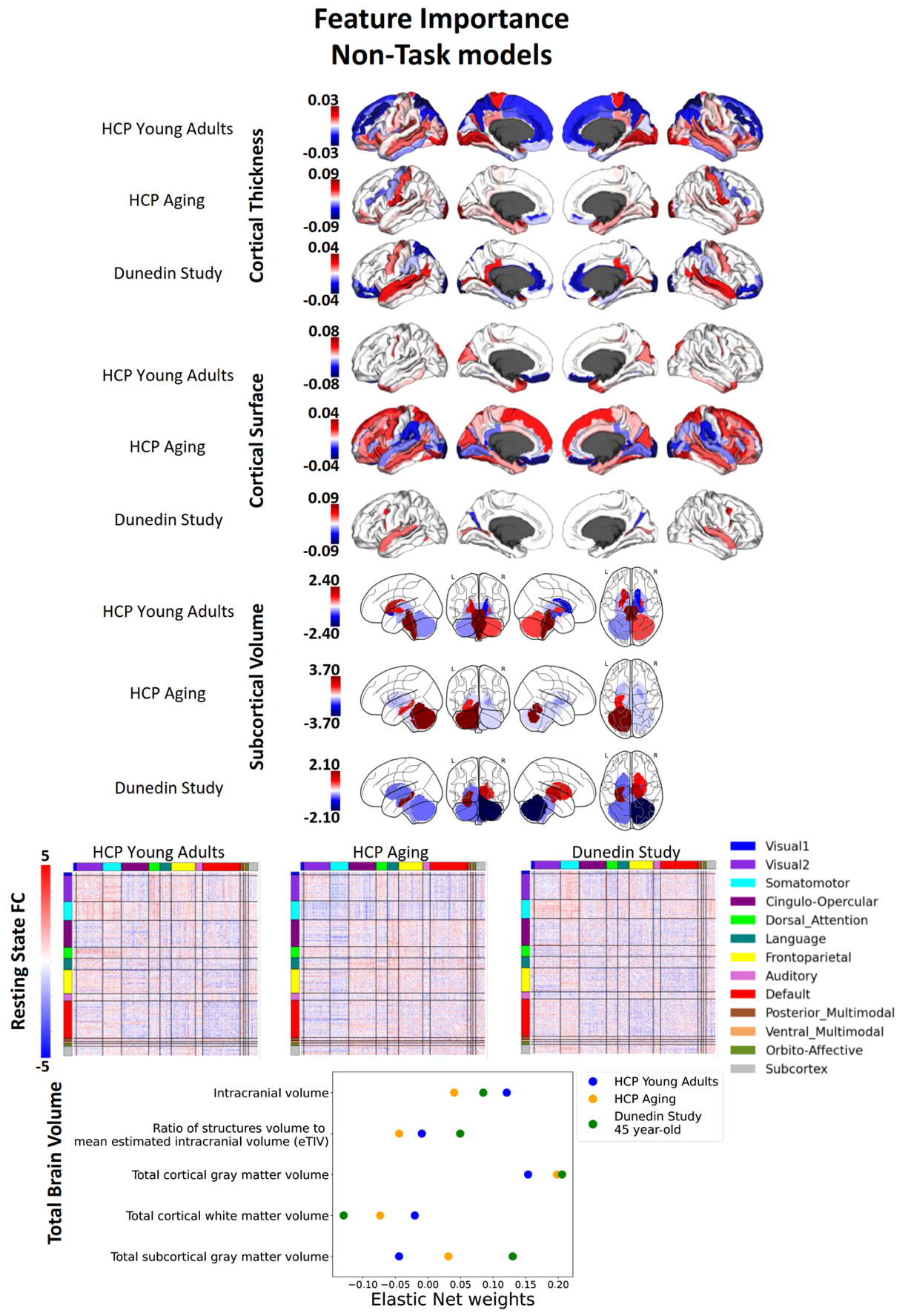

**Figure S30. Feature importance of non-stacked, non-task models with Elastic Net, indicated by Elastic Net Coefficients, for each dataset, when predicting cognitive abilities at the time of scanning.** A higher magnitude of a coefficient indicates a stronger contribution to the prediction. FC = Functional Connectivity.

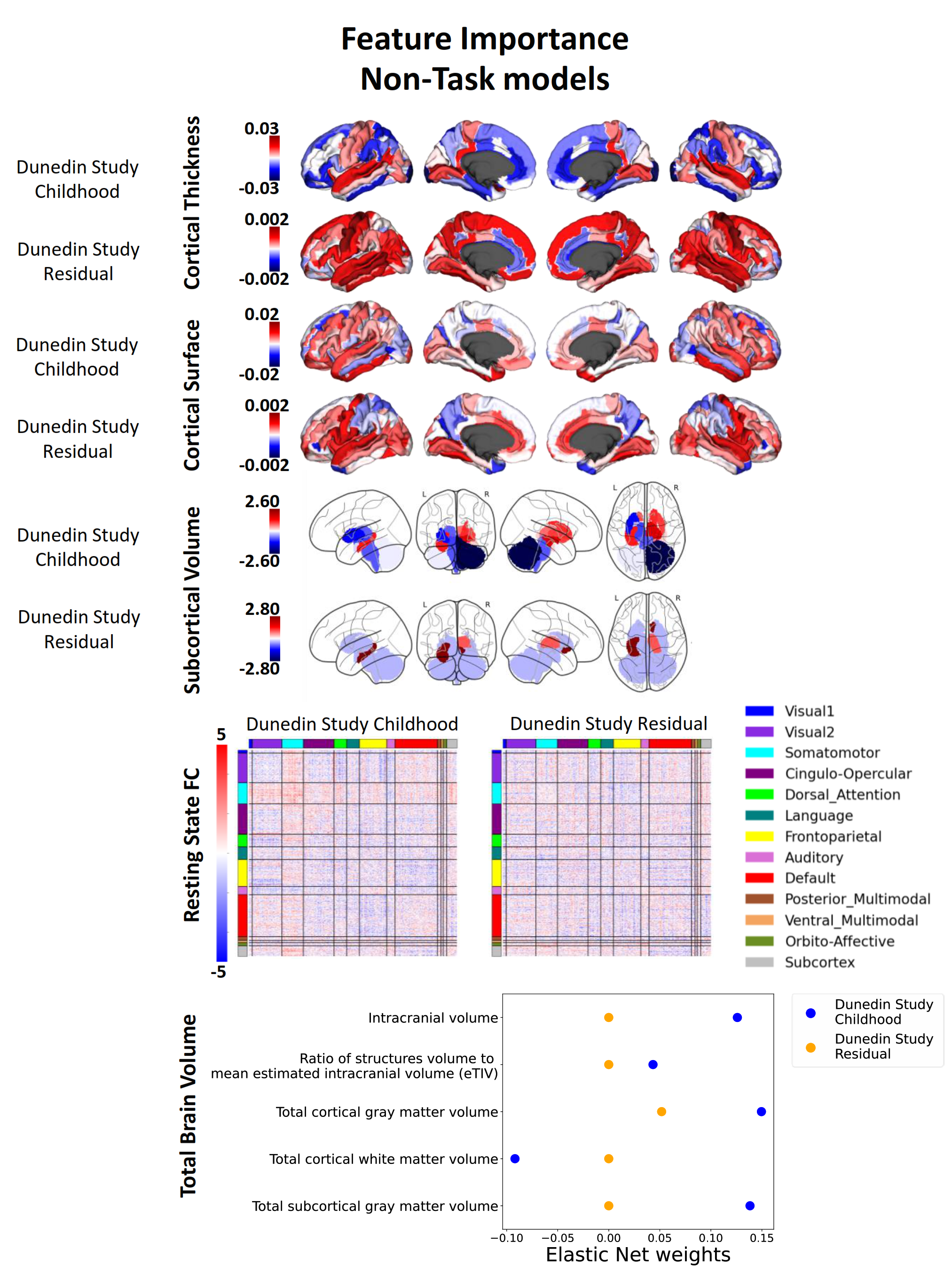

**Figure S30 (Cont.). Feature importance of non-stacked, non-task models with Elastic Net, indicated by Elastic Net Coefficients, for each dataset, when predicting cognitive abilities at the time of scanning.** A higher magnitude of a coefficient indicates a stronger contribution to the prediction. FC = Functional Connectivity.

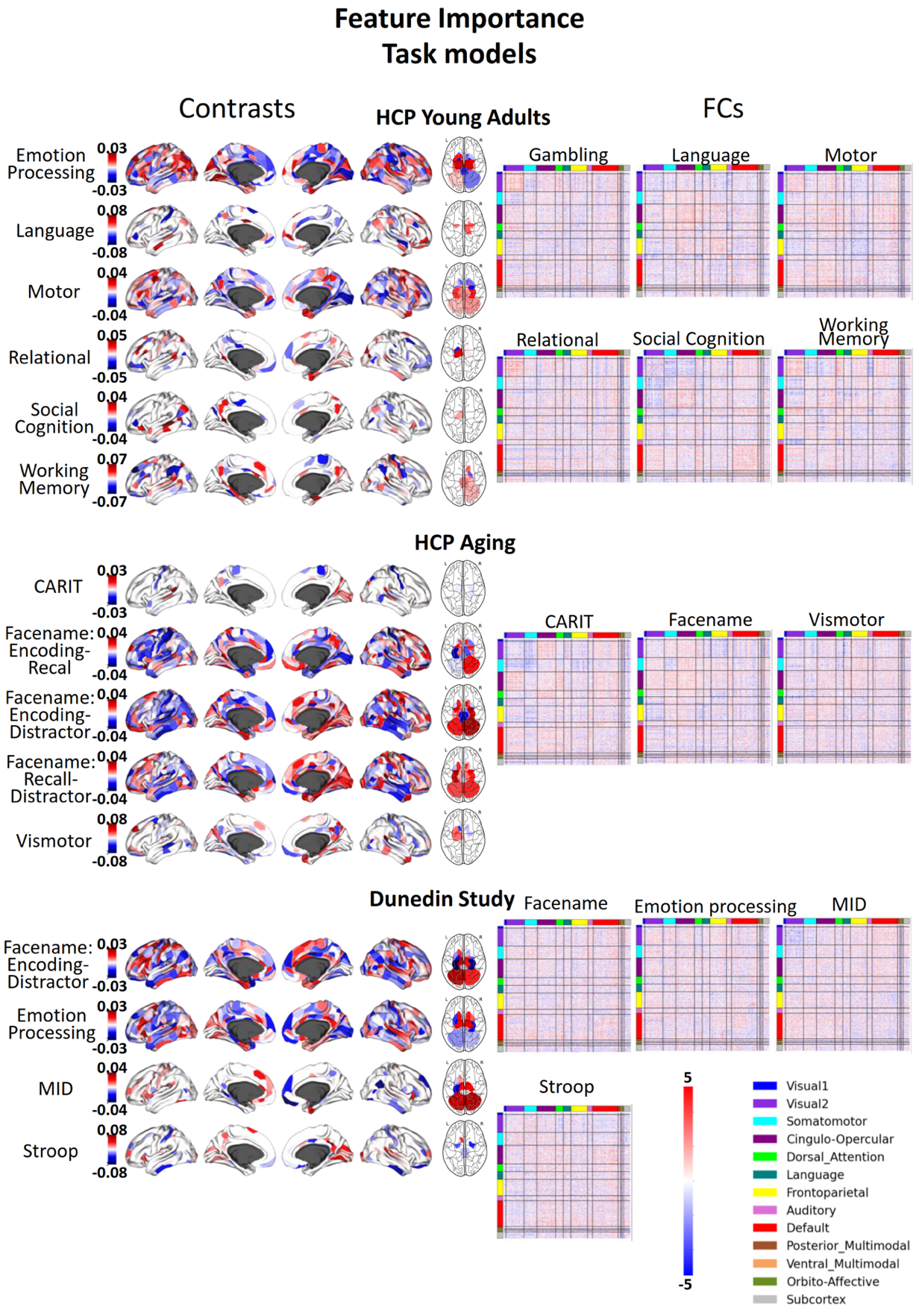

**Figure S31. Feature importance of non-stacked, task models (including Task Contrasts and Task FC) with Elastic Net, indicated by Elastic Net Coefficients, for each dataset, when predicting cognitive abilities at the time of scanning.** A higher magnitude of a coefficient indicates a stronger contribution to the prediction. FC = Functional Connectivity.

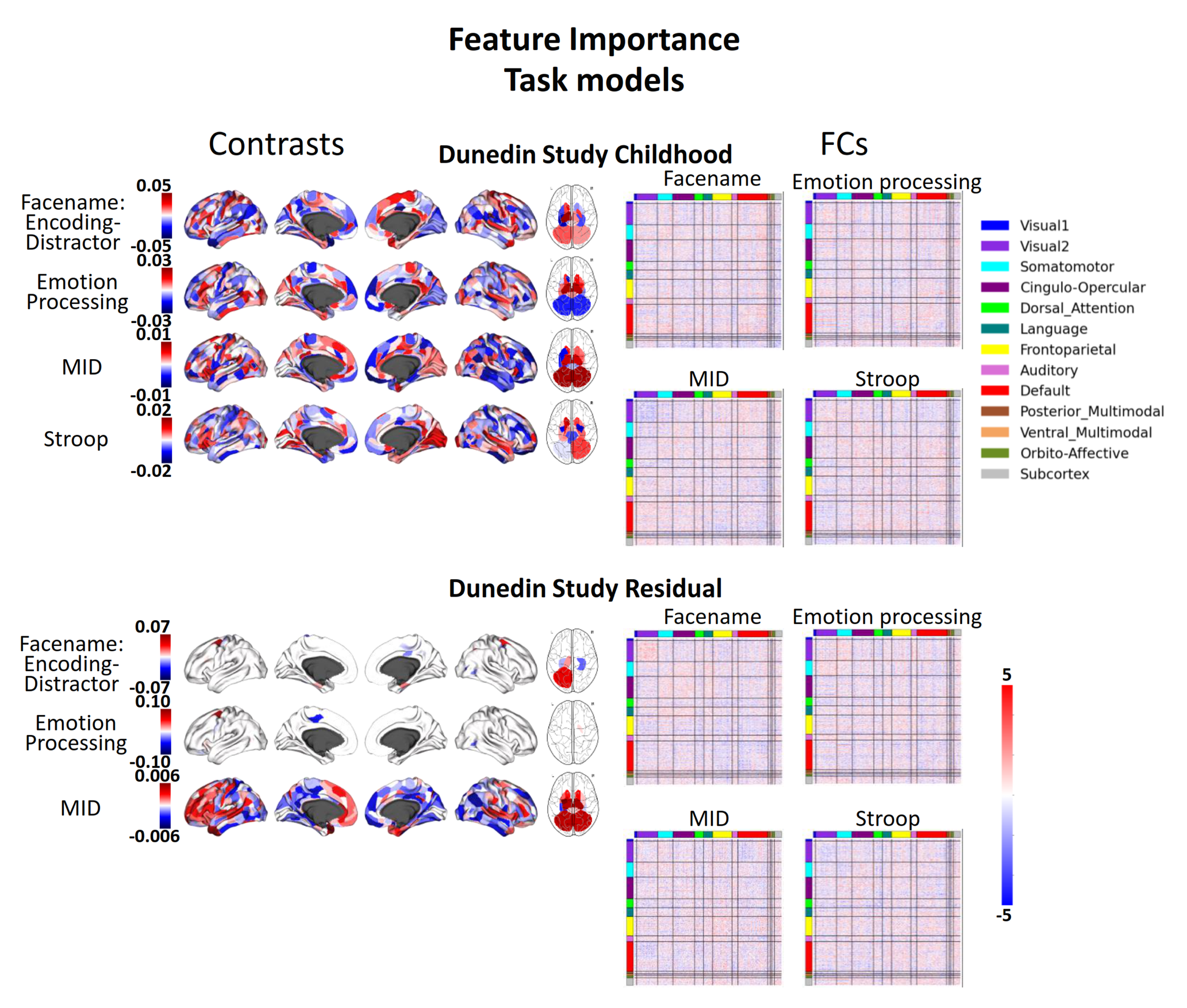

**Figure S31 (Cont.). Feature importance of non-stacked, task models (including Task Contrasts and Task FC) with Elastic Net, indicated by Elastic Net Coefficients, for each dataset, when predicting cognitive abilities at the time of scanning.** A higher magnitude of a coefficient indicates a stronger contribution to the prediction. FC = Functional Connectivity.

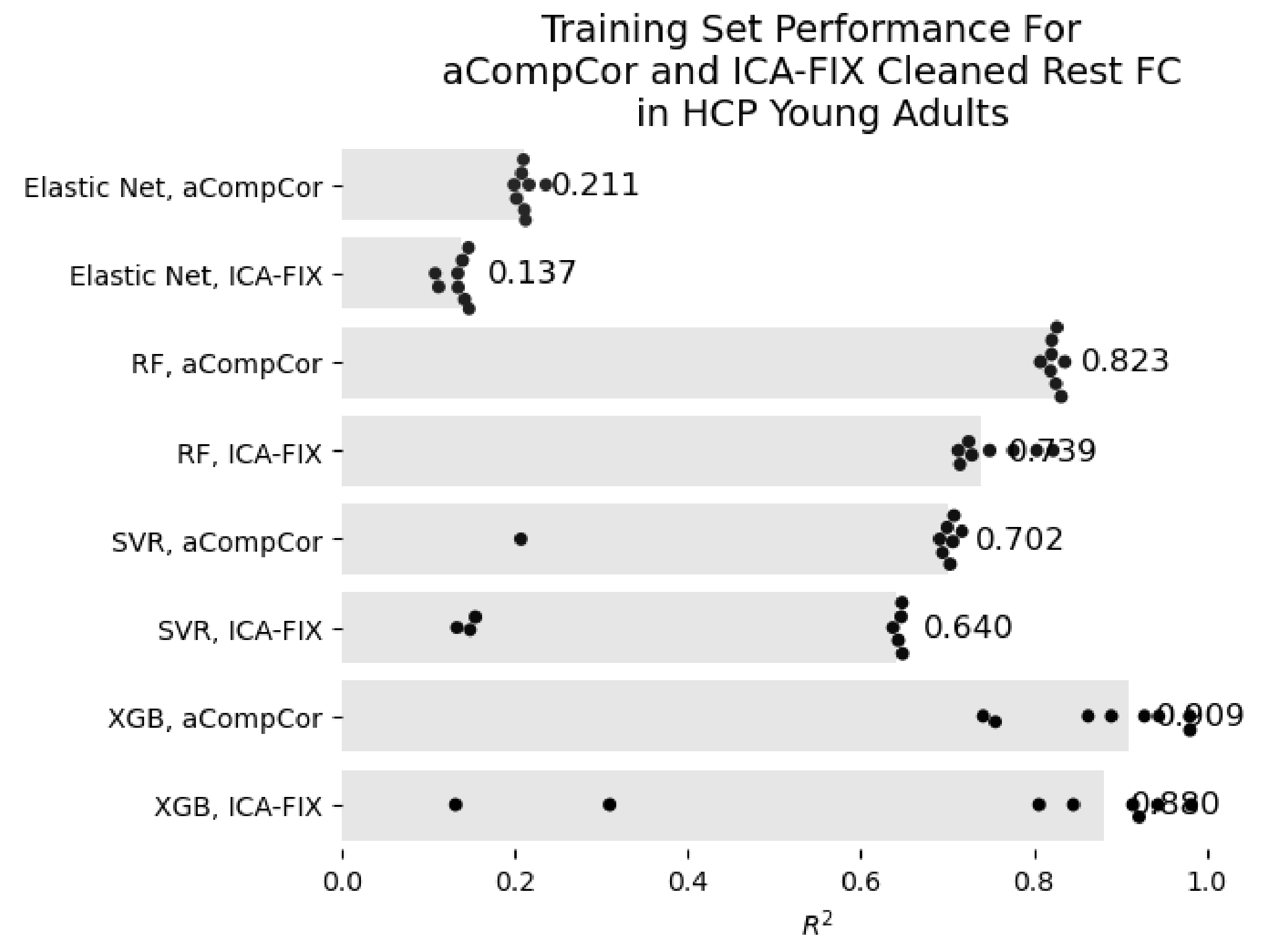

**Figure S32. Comparison in the predictive performance (*R^2^*) within training sets for rest FC in HCP Young Adults between two denoising strategies: aCompCor and ICA-FIX.** Each dot represents predictive performance at each training set, and the number to the right is the median across sets. eNet = Elastic Net; RF = Random Forest; SVR = Support Vector Regression; XGB = XGBoost; FC = Functional Connectivity.

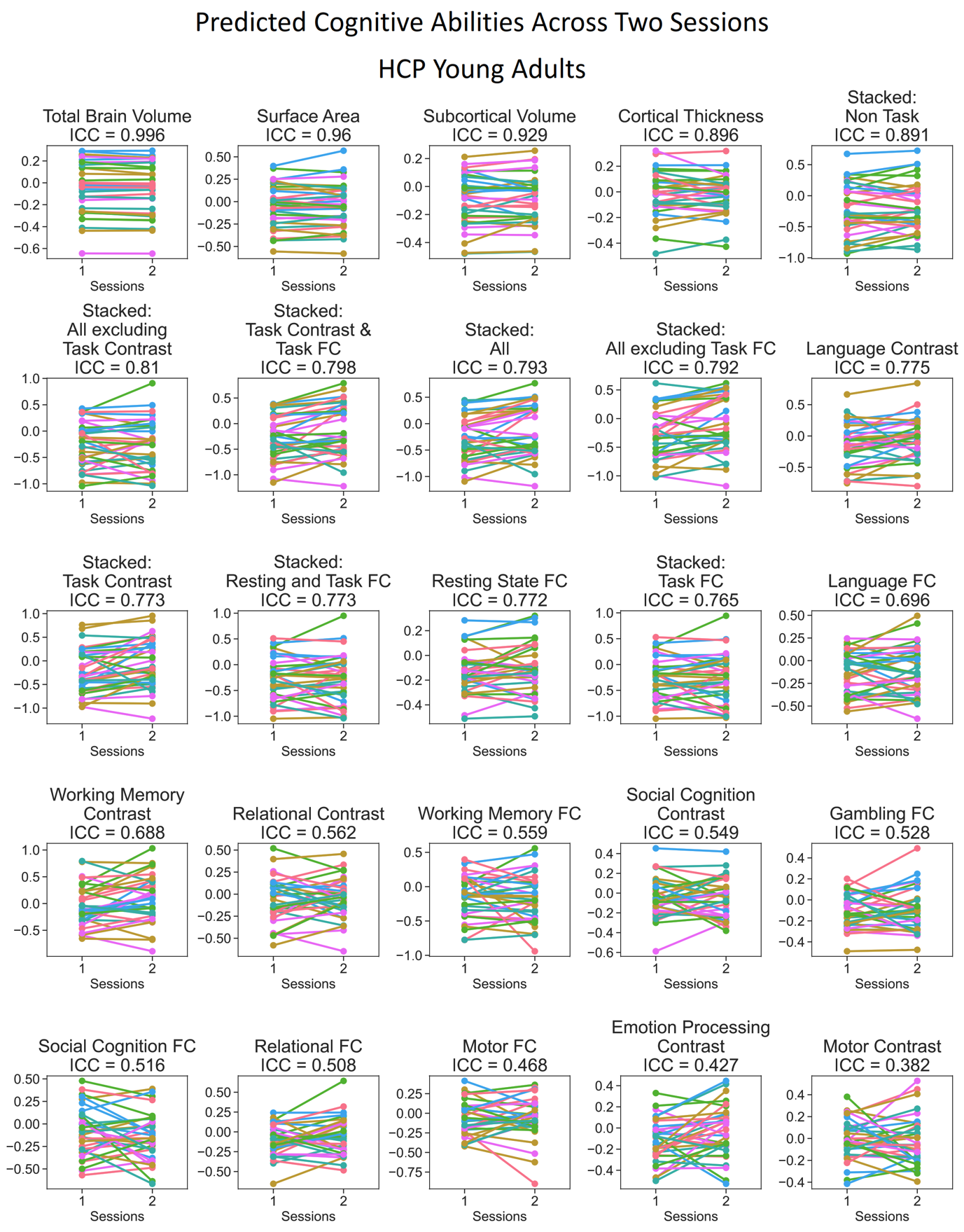

***Figure S33. Predicted values of stacked and non-stacked models across two scanning sessions, ranked by interclass correlation (ICC) for HCP Young Adults.*** *Each line represents each participant. Lines would be completely parallel with each other in the case of perfect test-retest reliability.*

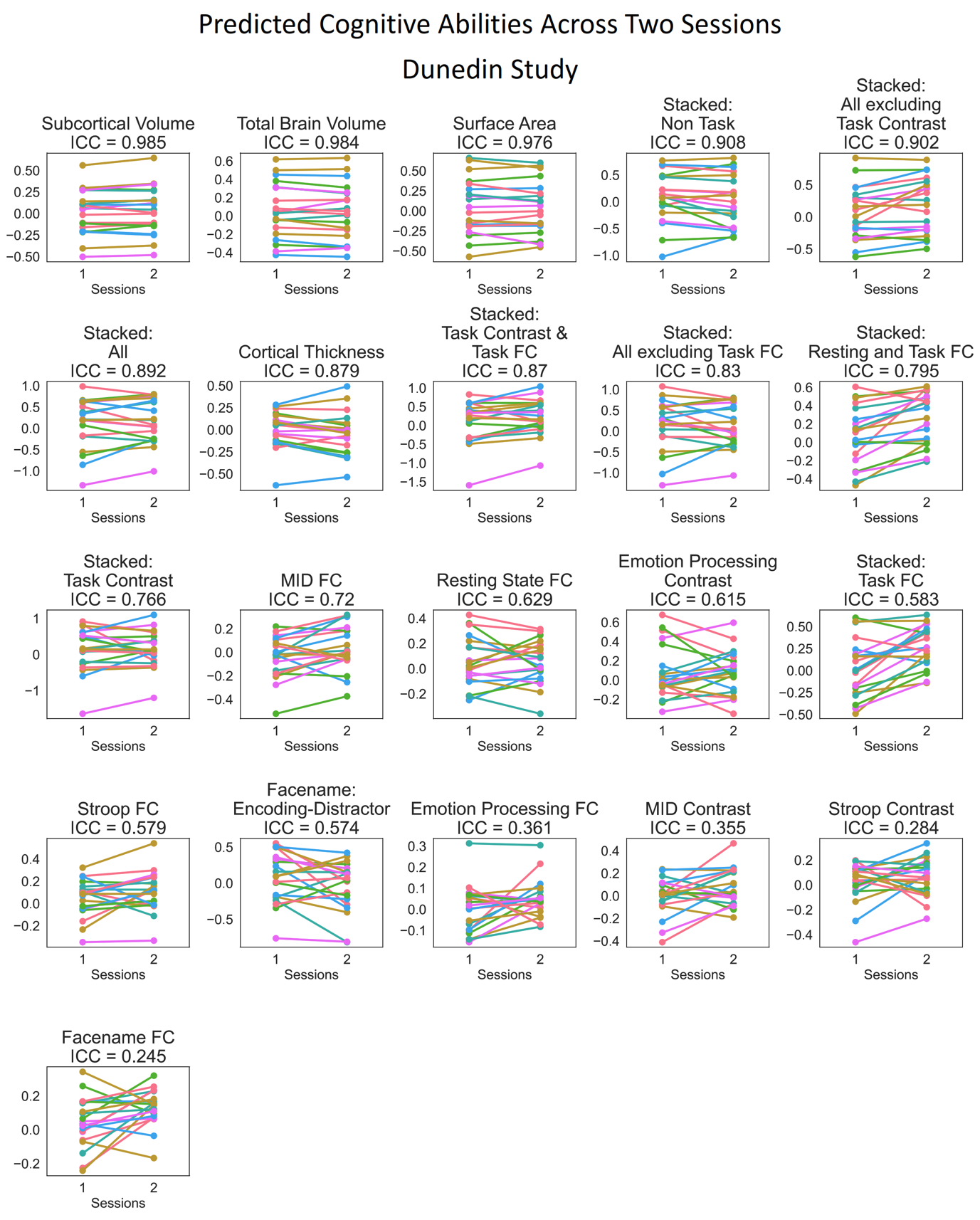

***Figure S34. Predicted values of stacked and non-stacked models across two scanning sessions, ranked by interclass correlation (ICC) for Dunedin Study.*** *Each line represents each participant. Lines would be completely parallel with each other in the case of perfect test-retest reliability.*

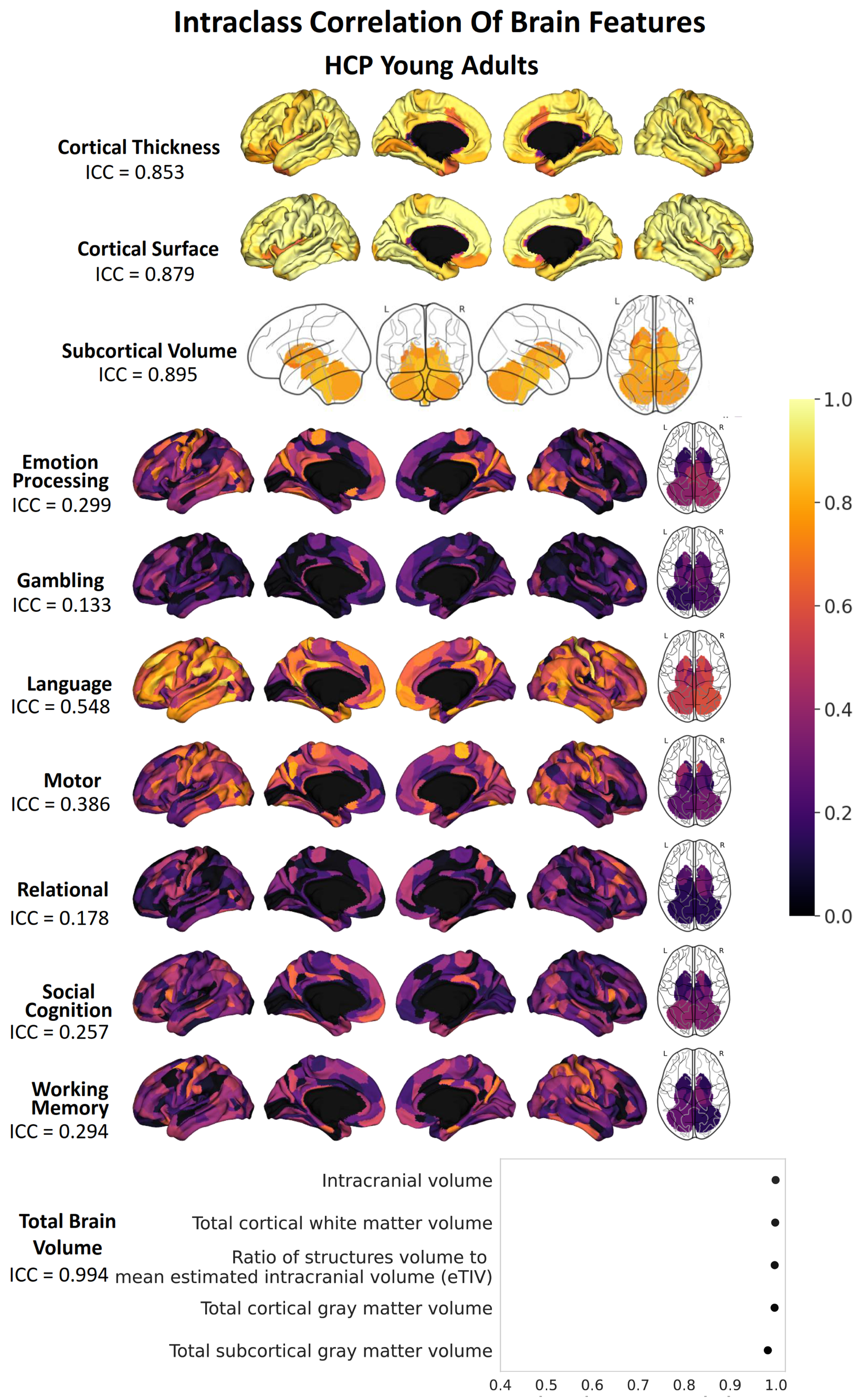

**Figure S35. Interclass correlation (ICC) of each brain feature, excluding those from Rest and Task FC, before prediction modelling for HCP Young Adults.** The ICC number indicates the mean of ICC for that particular set of features.

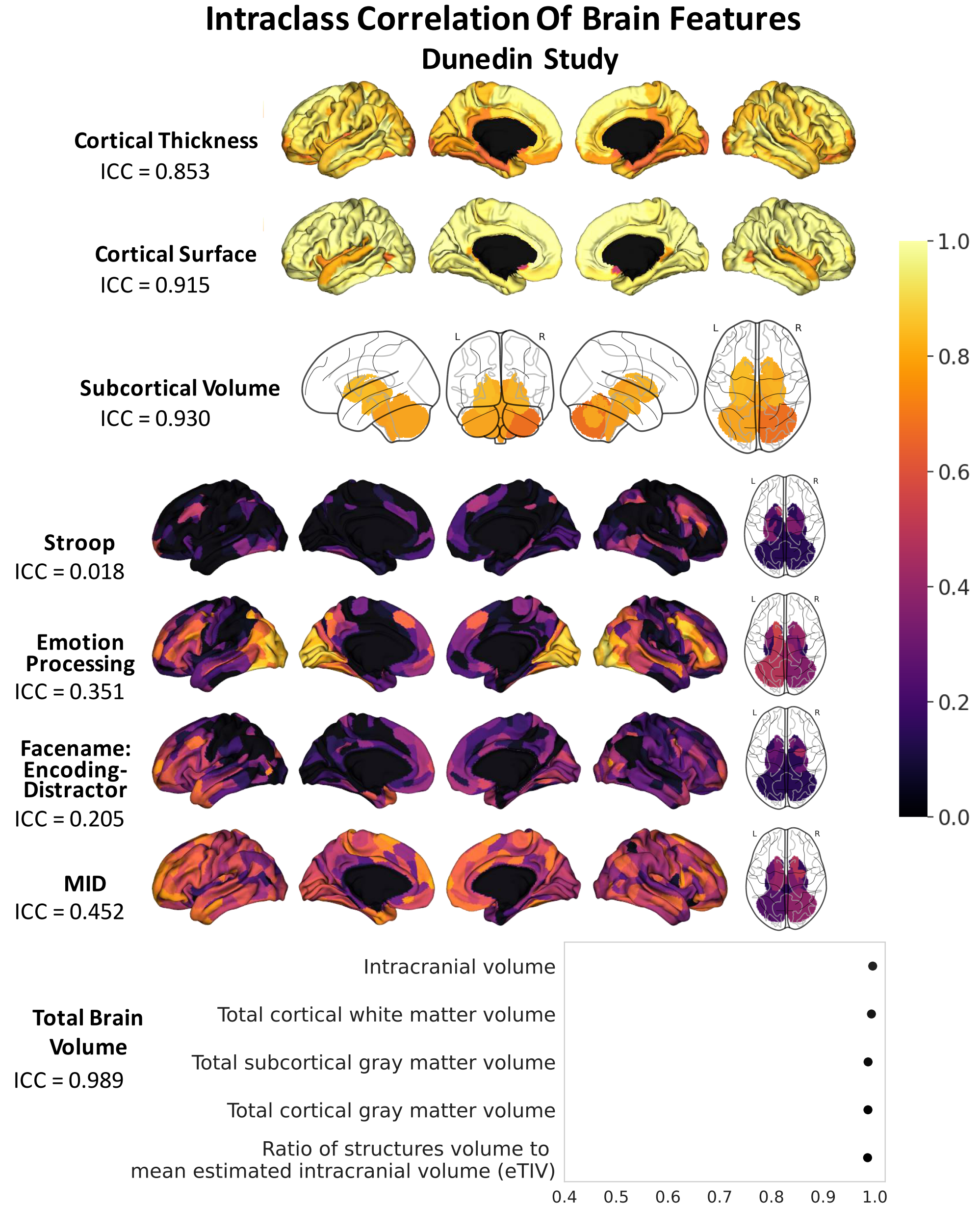

**Figure S36 Interclass correlation (ICC) of each brain feature, excluding those from Rest and Task FC, before prediction modelling for Dunedin Study.** The ICC number indicates the mean of ICC for that particular set of features.

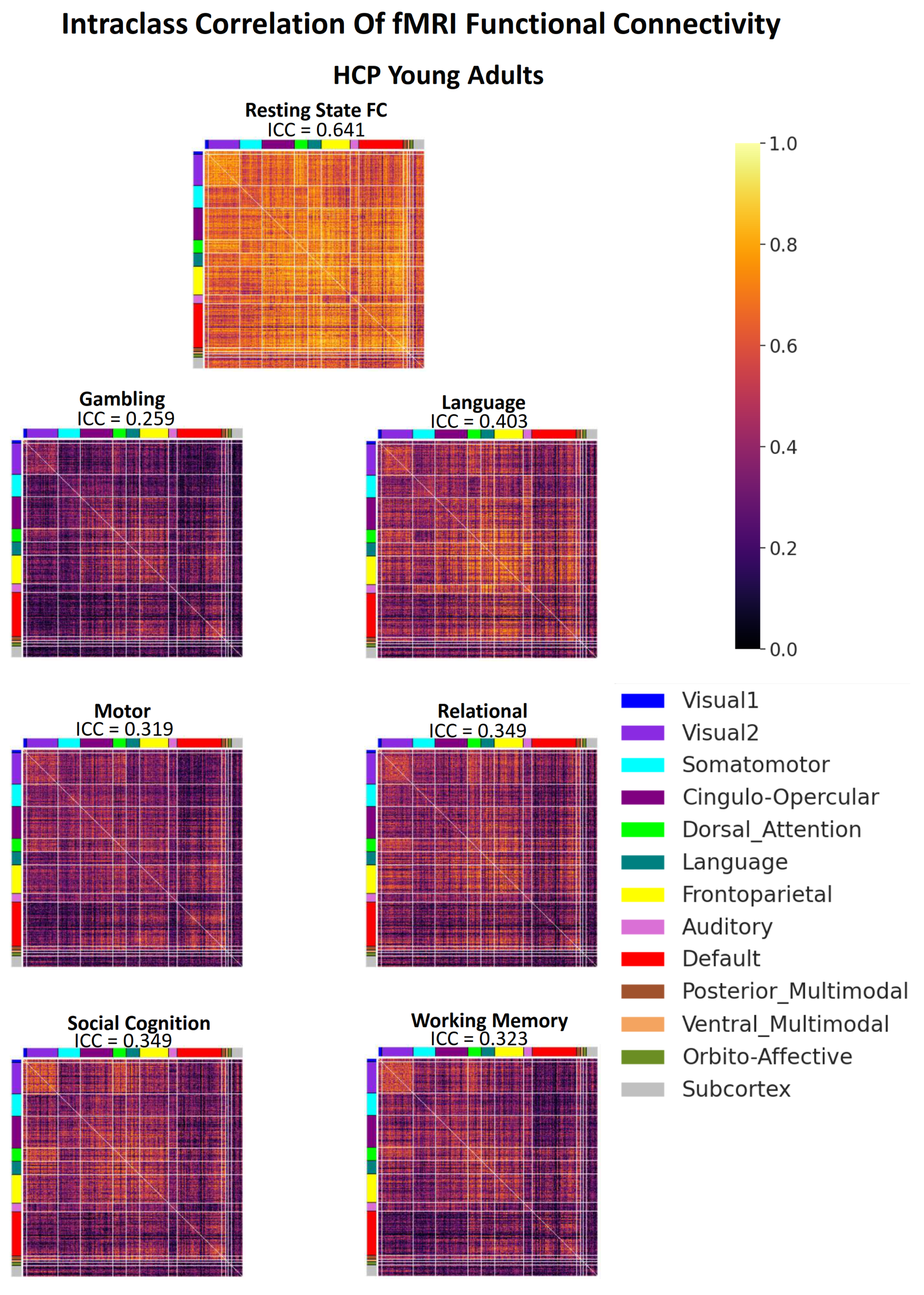

**Figure S37 Interclass correlation (ICC) of each brain feature of Rest and Task FC before prediction modelling for HCP Aging.** The ICC number indicates the mean of ICC for that particular set of features.

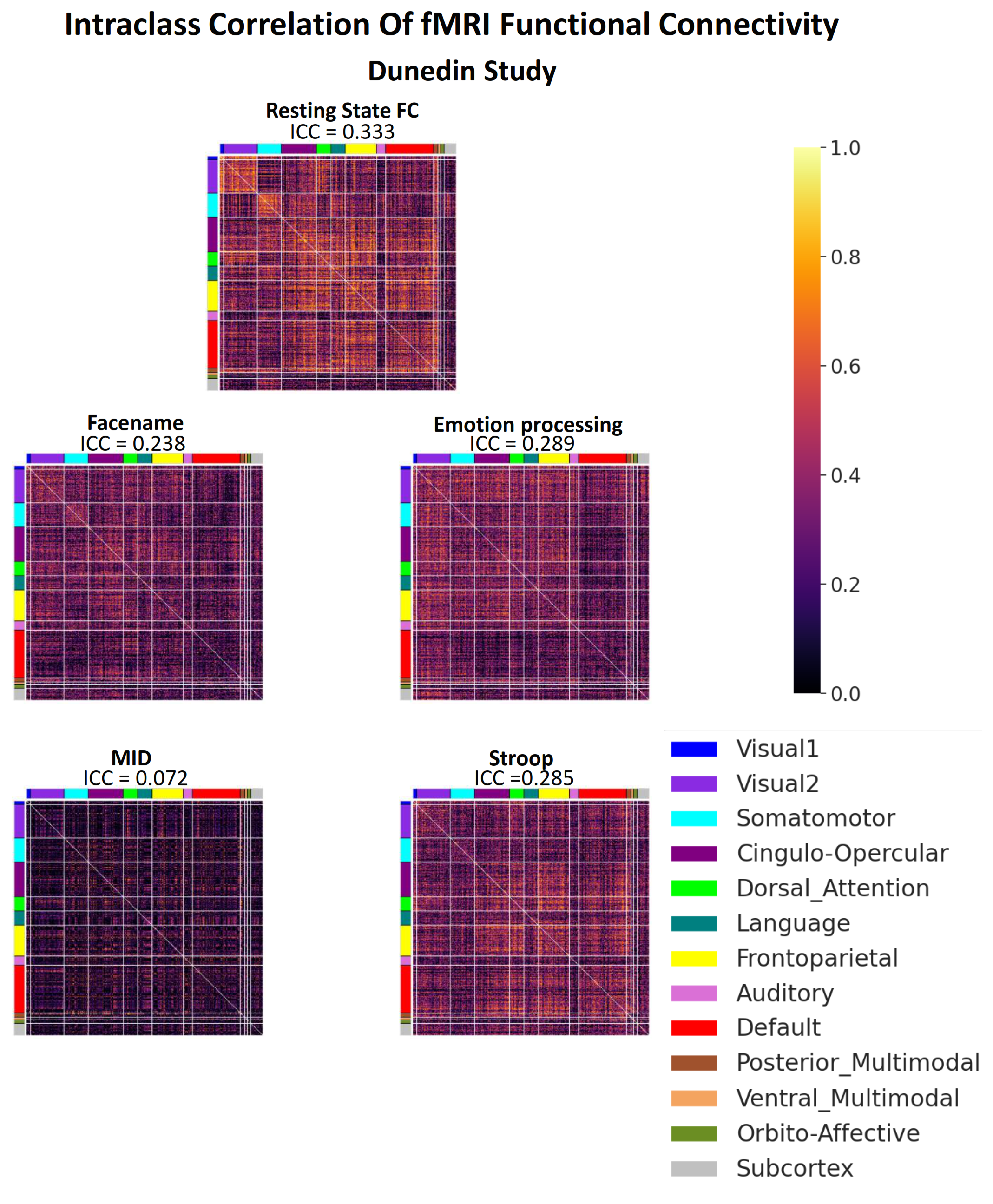

**Figure S38 Interclass correlation (ICC) of each brain feature of Rest and Task FC before prediction modelling for Dunedin Study.** The ICC number indicates the mean of ICC for that particular set of features.

***Table S1. Feature importance of the top-performing non-stacked models with with Elastic Net, as indicated by Elastic Net coefficients for the HCP Young Adult dataset: Working Memory Task Contrast***

| **Glasser's label** | **Brain Region** | **Hemisphere** | **Network** | **Amplitude** |
| --- | --- | --- | --- | --- |
| R_8BL | Dorsolateral Prefrontal, 8BL | Right | Default | -0.0699 |
| L_AIP | Superior Parietal, AIP | Left | Dorsal_Attention | 0.0567 |
| R_PFop | Inferior Parietal, PFop | Right | Cingulo-Opercular | -0.0537 |
| L_PCV | Posterior Cingulate, PCV | Left | Posterior_Multimodal | 0.0508 |
| L_PF | Inferior Parietal, PF | Left | Cingulo-Opercular | -0.0463 |
| L_PGs | Inferior Parietal, PGs | Left | Default | -0.0432 |
| R_PF | Inferior Parietal, PF | Right | Cingulo-Opercular | -0.0425 |
| R_PCV | Posterior Cingulate, PCV | Right | Posterior_Multimodal | 0.0421 |
| R_VIP | Superior Parietal, VIP | Right | Visual2 | 0.0409 |
| R_PHA1 | Medial Temporal, PHA1 | Right | Default | -0.0406 |
| R_33pr | Anterior Cingulate and Medial Prefrontal, 33pr | Right | Frontoparietal | 0.0405 |
| R_EC | Medial Temporal, EC | Right | Default | 0.0397 |
| R_d32 | Anterior Cingulate and Medial Prefrontal, d32 | Right | Frontoparietal | 0.0396 |
| R_7Pm | Superior Parietal, 7Pm | Right | Frontoparietal | 0.0392 |
| L_31pv | Posterior Cingulate, 31pv | Left | Default | -0.0373 |
| R_AAIC | Insular and Frontal Opercular, AAIC | Right | Orbito-Affective | 0.036 |
| L_47s | Orbital and Polar Frontal, 47s | Left | Default | -0.0355 |
| L_V8 | Ventral Stream Visual, V8 | Left | Visual2 | -0.0342 |
| L_FST | MT+ Complex and Neighboring Visual Areas, FST | Left | Visual2 | 0.0335 |
| L_7Pm | Superior Parietal, 7Pm | Left | Frontoparietal | 0.0335 |
| R_IFSa | Inferior Frontal, IFSa | Right | Cingulo-Opercular | -0.0321 |
| R_6v | Premotor, 6v | Right | Somatomotor | -0.0317 |
| L_6mp | Paracentral Lobular and Mid Cingulate, 6mp | Left | Somatomotor | -0.0311 |
| R_PGp | Inferior Parietal, PGp | Right | Dorsal_Attention | -0.0287 |
| R_5mv | Paracentral Lobular and Mid Cingulate, 5mv | Right | Cingulo-Opercular | -0.0287 |
| R_i6-8 | Dorsolateral Prefrontal, i6-8 | Right | Frontoparietal | 0.0286 |
| R_TPOJ1 | Temporo-Parieto-Occipital Junction, TPOJ1 | Right | Language | 0.0286 |
| R_IFJp | Inferior Frontal, IFJp | Right | Frontoparietal | 0.028 |
| R_6r | Premotor, 6r | Right | Cingulo-Opercular | -0.0277 |
| R_13l | Orbital and Polar Frontal, 13l | Right | Frontoparietal | -0.0276 |
| L_IP1 | Inferior Parietal, IP1 | Left | Frontoparietal | 0.0274 |
| R_p32 | Anterior Cingulate and Medial Prefrontal, p32 | Right | Default | 0.0267 |
| L_MIP | Superior Parietal, MIP | Left | Dorsal_Attention | 0.0264 |
| R_AVI | Insular and Frontal Opercular, AVI | Right | Frontoparietal | 0.0263 |
| R_DVT | Posterior Cingulate, DVT | Right | Visual1 | -0.025 |
| L_LIPd | Superior Parietal, LIPd | Left | Dorsal_Attention | 0.0222 |
| PALLIDUM_RIGHT | Pallidum Right | Right | Subcortex | -0.0221 |
| R_6a | Premotor, 6a | Right | Dorsal_Attention | 0.0216 |
| R_p10p | Orbital and Polar Frontal, p10p | Right | Frontoparietal | 0.0215 |
| R_8Av | Dorsolateral Prefrontal, 8Av | Right | Default | -0.0214 |
| R_LIPd | Superior Parietal, LIPd | Right | Dorsal_Attention | 0.0213 |
| L_TE2p | Lateral Temporal, TE2p | Left | Dorsal_Attention | 0.0212 |
| CAUDATE_RIGHT | Caudate Right | Right | Subcortex | 0.0212 |
| R_AIP | Superior Parietal, AIP | Right | Dorsal_Attention | 0.0196 |
| L_33pr | Anterior Cingulate and Medial Prefrontal, 33pr | Left | Cingulo-Opercular | 0.0195 |
| L_FOP3 | Insular and Frontal Opercular, FOP3 | Left | Cingulo-Opercular | -0.0194 |
| L_p47r | Inferior Frontal, p47r | Left | Frontoparietal | -0.0192 |
| R_A1 | Early Auditory, A1 | Right | Auditory | 0.0191 |
| L_LO2 | MT+ Complex and Neighboring Visual Areas, LO2 | Left | Visual2 | -0.019 |
| R_LBelt | Early Auditory, LBelt | Right | Auditory | 0.019 |
| L_PreS | Medial Temporal, PreS | Left | Default | 0.0185 |
| R_IFSp | Inferior Frontal, IFSp | Right | Frontoparietal | -0.0181 |
| R_a24 | Anterior Cingulate and Medial Prefrontal, a24 | Right | Default | 0.018 |
| L_7AL | Superior Parietal, 7AL | Left | Somatomotor | -0.018 |
| BRAIN_STEM | Brain Stem | None | Subcortex | 0.0179 |
| R_7m | Posterior Cingulate, 7m | Right | Default | 0.0179 |
| L_TPOJ3 | Temporo-Parieto-Occipital Junction, TPOJ3 | Left | Posterior_Multimodal | 0.0177 |
| R_STSdp | Auditory Association, STSdp | Right | Language | 0.0175 |
| R_FOP3 | Insular and Frontal Opercular, FOP3 | Right | Cingulo-Opercular | -0.0174 |
| R_A5 | Auditory Association, A5 | Right | Language | 0.0167 |
| R_9p | Dorsolateral Prefrontal, 9p | Right | Default | -0.0165 |
| R_47m | Orbital and Polar Frontal, 47m | Right | Default | -0.0164 |
| L_PIT | Ventral Stream Visual, PIT | Left | Visual2 | -0.0164 |
| R_PeEc | Medial Temporal, PeEc | Right | Ventral_Multimodal | 0.0157 |
| L_V4t | MT+ Complex and Neighboring Visual Areas, V4t | Left | Visual2 | 0.0155 |
| L_10d | Orbital and Polar Frontal, 10d | Left | Default | -0.0154 |
| R_IP2 | Inferior Parietal, IP2 | Right | Frontoparietal | 0.0153 |
| R_VMV2 | Ventral Stream Visual, VMV2 | Right | Visual2 | -0.0152 |
| R_PHT | Lateral Temporal, PHT | Right | Dorsal_Attention | -0.0144 |
| L_7PL | Superior Parietal, 7Pl | Left | Dorsal_Attention | 0.0131 |
| L_v23ab | Posterior Cingulate, v23ab | Left | Default | -0.0119 |
| L_V3B | Dorsal Stream Visual, V3B | Left | Visual2 | -0.0119 |
| L_PFt | Inferior Parietal, PFt | Left | Dorsal_Attention | 0.0113 |
| R_FOP2 | Insular and Frontal Opercular, FOP2 | Right | Somatomotor | 0.0111 |
| R_V1 | Primary Visual, V1 | Right | Visual1 | 0.011 |
| CEREBELLUM_RIGHT | Cerebellum Right | Right | Subcortex | 0.0106 |
| R_OFC | Orbital and Polar Frontal, OFC | Right | Frontoparietal | -0.0105 |
| R_V3CD | MT+ Complex and Neighboring Visual Areas, V3CD | Right | Visual2 | -0.01 |
| R_TE1a | Lateral Temporal, TE1a | Right | Default | -0.0098 |
| R_10pp | Orbital and Polar Frontal, 10pp | Right | Default | -0.0095 |
| R_MBelt | Early Auditory, MBelt | Right | Auditory | 0.0094 |
| R_STV | Temporo-Parieto-Occipital Junction, STV | Right | Posterior_Multimodal | 0.0093 |
| L_FOP1 | Posterior Opercular, FOP1 | Left | Cingulo-Opercular | -0.0093 |
| L_23c | Paracentral Lobular and Mid Cingulate, 23c | Left | Cingulo-Opercular | -0.0092 |
| L_V2 | Early Visual, V2 | Left | Visual2 | 0.0091 |
| L_44 | Inferior Frontal, 44 | Left | Language | -0.0091 |
| THALAMUS_RIGHT | Thalamus Right | Right | Subcortex | 0.0089 |
| HIPPOCAMPUS_RIGHT | Hippocampus Right | Right | Subcortex | 0.0089 |
| R_LO3 | MT+ Complex and Neighboring Visual Areas, LO3 | Right | Visual2 | 0.0086 |
| L_V1 | Primary Visual, V1 | Left | Visual1 | 0.0085 |
| L_TE1a | Lateral Temporal, TE1a | Left | Default | -0.0082 |
| R_TE2p | Lateral Temporal, TE2p | Right | Dorsal_Attention | 0.0081 |
| L_PFcm | Early Auditory, PFcm | Left | Cingulo-Opercular | -0.0081 |
| L_OP2-3 | Posterior Opercular, OP2-3 | Left | Somatomotor | -0.0076 |
| L_IFJp | Inferior Frontal, IFJp | Left | Frontoparietal | 0.0076 |
| L_3b | Somatosensory and Motor, 3b | Left | Somatomotor | -0.0075 |
| R_a47r | Inferior Frontal, a47r | Right | Frontoparietal | 0.0072 |
| L_5m | Paracentral Lobular and Mid Cingulate, 5m | Left | Somatomotor | -0.0071 |
| R_d23ab | Posterior Cingulate, d23ab | Right | Default | -0.0069 |
| R_FEF | Premotor, FEF | Right | Cingulo-Opercular | 0.0067 |
| L_FEF | Premotor, FEF | Left | Cingulo-Opercular | 0.0066 |
| R_SFL | Dorsolateral Prefrontal, SFL | Right | Language | -0.0066 |
| L_PHA3 | Medial Temporal, PHA3 | Left | Dorsal_Attention | 0.0065 |
| R_FOP1 | Posterior Opercular, FOP1 | Right | Cingulo-Opercular | -0.006 |
| L_ProS | Posterior Cingulate, ProS | Left | Visual1 | 0.0059 |
| L_p24pr | Anterior Cingulate and Medial Prefrontal, p24pr | Left | Cingulo-Opercular | -0.0058 |
| R_MIP | Superior Parietal, MIP | Right | Dorsal_Attention | 0.0057 |
| L_STSva | Auditory Association, STSva | Left | Default | -0.0057 |
| R_43 | Posterior Opercular, 43 | Right | Cingulo-Opercular | -0.0056 |
| L_MBelt | Early Auditory, MBelt | Left | Auditory | -0.0056 |
| ACCUMBENS_RIGHT | Accumbens Right | Right | Subcortex | 0.0055 |
| R_v23ab | Posterior Cingulate, v23ab | Right | Default | -0.0054 |
| L_SCEF | Paracentral Lobular and Mid Cingulate, SCEF | Left | Cingulo-Opercular | 0.0054 |
| R_PSL | Temporo-Parieto Occipital Junction, PSL | Right | Cingulo-Opercular | -0.0047 |
| L_8BL | Dorsolateral Prefrontal, 8BL | Left | Default | -0.0046 |
| L_9-46d | Dorsolateral Prefrontal, 9-46d | Left | Cingulo-Opercular | 0.0045 |
| R_V8 | Ventral Stream Visual, V8 | Right | Visual2 | -0.0041 |
| R_10v | Anterior Cingulate and Medial Prefrontal, 10v | Right | Default | -0.0036 |
| R_STSda | Auditory Association, STSda | Right | Default | 0.0036 |
| R_11l | Orbital and Polar Frontal, 11l | Right | Frontoparietal | 0.0034 |
| R_23c | Paracentral Lobular and Mid Cingulate, 23c | Right | Cingulo-Opercular | -0.0034 |
| L_LBelt | Early Auditory, LBelt | Left | Auditory | 0.0034 |
| R_POS2 | Posterior Cingulate, POS2 | Right | Frontoparietal | -0.0031 |
| R_TE1m | Lateral Temporal, TE1m | Right | Frontoparietal | 0.003 |
| L_OFC | Orbital and Polar Frontal, OFC | Left | Default | -0.0029 |
| L_9a | Dorsolateral Prefrontal, 9a | Left | Default | -0.0026 |
| L_55b | Premotor, 55b | Left | Language | 0.0026 |
| L_MI | Insular and Frontal Opercular, MI | Left | Cingulo-Opercular | -0.0024 |
| L_STGa | Auditory Association, STGa | Left | Language | -0.0021 |
| L_13l | Orbital and Polar Frontal, 13l | Left | Frontoparietal | -0.0021 |
| L_POS1 | Posterior Cingulate, POS1 | Left | Default | -0.002 |
| L_7PC | Superior Parietal, 7PC | Left | Somatomotor | -0.0019 |
| R_44 | Inferior Frontal, 44 | Right | Frontoparietal | -0.0019 |
| L_9m | Anterior Cingulate and Medial Prefrontal, 9m | Left | Default | -0.0016 |
| R_FST | MT+ Complex and Neighboring Visual Areas, FST | Right | Visual2 | 0.0016 |
| R_V3B | Dorsal Stream Visual, V3B | Right | Visual2 | -0.0016 |
| L_pOFC | Anterior Cingulate and Medial Prefrontal, pOFC | Left | Orbito-Affective | -0.0015 |
| R_PHA3 | Medial Temporal, PHA3 | Right | Dorsal_Attention | -0.0013 |
| R_TGv | Lateral Temporal, TGv | Right | Language | -0.0013 |
| L_d23ab | Posterior Cingulate, d23ab | Left | Default | -0.0013 |
| L_31a | Posterior Cingulate, 31a | Left | Default | -0.0012 |
| L_LO3 | MT+ Complex and Neighboring Visual Areas, LO3 | Left | Visual2 | 0.0012 |
| PUTAMEN_RIGHT | Putamen Right | Right | Subcortex | 0.001 |
| R_8Ad | Dorsolateral Prefrontal, 8Ad | Right | Default | -0.001 |
| L_p32pr | Anterior Cingulate and Medial Prefrontal, p32pr | Left | Cingulo-Opercular | -0.0009 |
| L_STSda | Auditory Association, STSda | Left | Language | -0.0009 |
| DIENCEPHALON_VENTRAL_LEFT | Diencephalon Ventral Left | Left | Subcortex | -0.0009 |
| R_V4t | MT+ Complex and Neighboring Visual Areas, V4t | Right | Visual2 | 0.0008 |
| L_PFm | Inferior Parietal, PFm | Left | Frontoparietal | -0.0007 |
| L_31pd | Posterior Cingulate, 31pd | Left | Default | -0.0005 |
| L_TE1p | Lateral Temporal, TE1p | Left | Frontoparietal | -0.0004 |
| L_PoI1 | Insular and Frontal Opercular, PoI1 | Left | Cingulo-Opercular | -0.0004 |
| L_V3 | Early Visual, V3 | Left | Visual2 | 0.0004 |
| L_DVT | Posterior Cingulate, DVT | Left | Visual1 | -0.0004 |
| R_FFC | Ventral Stream Visual, FFC | Right | Visual2 | 0 |
| L_FOP2 | Insular and Frontal Opercular, FOP2 | Left | Somatomotor | 0 |
| L_H | Medial Temporal, H | Left | Default | 0 |
| R_IPS1 | Dorsal Stream Visual, IPS1 | Right | Visual2 | 0 |
| R_V7 | Dorsal Stream Visual, V7 | Right | Visual2 | 0 |
| L_EC | Medial Temporal, EC | Left | Default | 0 |
| L_AVI | Insular and Frontal Opercular, AVI | Left | Frontoparietal | 0 |
| R_s6-8 | Dorsolateral Prefrontal, s6-8 | Right | Frontoparietal | 0 |
| L_PeEc | Medial Temporal, PeEc | Left | Ventral_Multimodal | 0 |
| L_PBelt | Early Auditory, PBelt | Left | Auditory | 0 |
| L_A5 | Auditory Association, A5 | Left | Language | 0 |
| L_PHA1 | Medial Temporal, PHA1 | Left | Default | 0 |
| R_IFJa | Inferior Frontal, IFJa | Right | Language | 0 |
| L_AAIC | Insular and Frontal Opercular, AAIC | Left | Orbito-Affective | 0 |
| R_OP4 | Posterior Opercular, OP4 | Right | Somatomotor | 0 |
| L_Pir | Insular and Frontal Opercular, Pir | Left | Orbito-Affective | 0 |
| R_LO2 | MT+ Complex and Neighboring Visual Areas, LO2 | Right | Visual2 | 0 |
| L_IFJa | Inferior Frontal, IFJa | Left | Language | 0 |
| R_52 | Early Auditory, 52 | Right | Auditory | 0 |
| L_IFSp | Inferior Frontal, IFSp | Left | Language | 0 |
| L_IFSa | Inferior Frontal, IFSa | Left | Frontoparietal | 0 |
| L_p9-46v | Dorsolateral Prefrontal, p9-46v | Left | Frontoparietal | 0 |
| L_46 | Dorsolateral Prefrontal, 46 | Left | Cingulo-Opercular | 0 |
| L_a9-46v | Dorsolateral Prefrontal, a9-46v | Left | Frontoparietal | 0 |
| R_OP2-3 | Posterior Opercular, OP2-3 | Right | Somatomotor | 0 |
| L_10v | Anterior Cingulate and Medial Prefrontal, 10v | Left | Default | 0 |
| L_a10p | Orbital and Polar Frontal, a10p | Left | Frontoparietal | 0 |
| L_10pp | Orbital and Polar Frontal, 10pp | Left | Default | 0 |
| L_11l | Orbital and Polar Frontal, 11l | Left | Frontoparietal | 0 |
| R_LO1 | MT+ Complex and Neighboring Visual Areas, LO1 | Right | Visual2 | 0 |
| L_FOP4 | Insular and Frontal Opercular, FOP4 | Left | Cingulo-Opercular | 0 |
| L_6a | Premotor, 6a | Left | Dorsal_Attention | 0 |
| L_i6-8 | Dorsolateral Prefrontal, i6-8 | Left | Frontoparietal | 0 |
| L_s6-8 | Dorsolateral Prefrontal, s6-8 | Left | Frontoparietal | 0 |
| L_43 | Posterior Opercular, 43 | Left | Cingulo-Opercular | 0 |
| L_OP4 | Posterior Opercular, OP4 | Left | Somatomotor | 0 |
| L_OP1 | Posterior Opercular, OP1 | Left | Somatomotor | 0 |
| R_OP1 | Posterior Opercular, OP1 | Right | Somatomotor | 0 |
| L_52 | Early Auditory, 52 | Left | Auditory | 0 |
| L_RI | Early Auditory, RI | Left | Auditory | 0 |
| L_STSvp | Auditory Association, STSvp | Left | Default | 0 |
| L_PoI2 | Insular and Frontal Opercular, PoI2 | Left | Cingulo-Opercular | 0 |
| L_TA2 | Auditory Association, TA2 | Left | Auditory | 0 |
| L_STSdp | Auditory Association, STSdp | Left | Language | 0 |
| L_TPOJ2 | Temporo-Parieto-Occipital Junction, TPOJ2 | Left | Posterior_Multimodal | 0 |
| L_TGd | Lateral Temporal, TGd | Left | Default | 0 |
| L_Ig | Insular and Frontal Opercular, Ig | Left | Somatomotor | 0 |
| L_p10p | Orbital and Polar Frontal, p10p | Left | Frontoparietal | 0 |
| R_47s | Orbital and Polar Frontal, 47s | Right | Default | 0 |
| L_TGv | Lateral Temporal, TGv | Left | Language | 0 |
| R_a9-46v | Dorsolateral Prefrontal, a9-46v | Right | Frontoparietal | 0 |
| L_A4 | Auditory Association, A4 | Left | Auditory | 0 |
| R_9-46d | Dorsolateral Prefrontal, 9-46d | Right | Cingulo-Opercular | 0 |
| L_TE1m | Lateral Temporal, TE1m | Left | Default | 0 |
| L_PI | Insular and Frontal Opercular, PI | Left | Cingulo-Opercular | 0 |
| L_a32pr | Anterior Cingulate and Medial Prefrontal, a32pr | Left | Cingulo-Opercular | 0 |
| L_p24 | Anterior Cingulate and Medial Prefrontal, p24 | Left | Cingulo-Opercular | 0 |
| ACCUMBENS_LEFT | Accumbens Left | Left | Subcortex | 0 |
| AMYGDALA_LEFT | Amygdala Left | Left | Subcortex | 0 |
| AMYGDALA_RIGHT | Amygdala Right | Right | Subcortex | 0 |
| R_V4 | Early Visual, V4 | Right | Visual2 | 0 |
| CAUDATE_LEFT | Caudate Left | Left | Subcortex | 0 |
| R_V3 | Early Visual, V3 | Right | Visual2 | 0 |
| CEREBELLUM_LEFT | Cerebellum Left | Left | Subcortex | 0 |
| R_V2 | Early Visual, V2 | Right | Visual2 | 0 |
| DIENCEPHALON_VENTRAL_RIGHT | Diencephalon Ventral Right | Right | Subcortex | 0 |
| HIPPOCAMPUS_LEFT | Hippocampus Left | Left | Subcortex | 0 |
| R_9a | Dorsolateral Prefrontal, 9a | Right | Default | 0 |
| PALLIDUM_LEFT | Pallidum Left | Left | Subcortex | 0 |
| R_V6 | Dorsal Stream Visual, V6 | Right | Visual2 | 0 |
| PUTAMEN_LEFT | Putamen Left | Left | Subcortex | 0 |
| THALAMUS_LEFT | Thalamus Left | Left | Subcortex | 0 |
| L_FOP5 | Insular and Frontal Opercular, FOP5 | Left | Cingulo-Opercular | 0 |
| L_s32 | Anterior Cingulate and Medial Prefrontal, s32 | Left | Default | 0 |
| R_p9-46v | Dorsolateral Prefrontal, p9-46v | Right | Frontoparietal | 0 |
| L_25 | Anterior Cingulate and Medial Prefrontal, 25 | Left | Default | 0 |
| L_TE2a | Lateral Temporal, TE2a | Left | Default | 0 |
| L_TF | Medial Temporal, TF | Left | Ventral_Multimodal | 0 |
| R_RSC | Posterior Cingulate, RSC | Right | Frontoparietal | 0 |
| L_PHT | Lateral Temporal, PHT | Left | Dorsal_Attention | 0 |
| L_PH | MT+ Complex and Neighboring Visual Areas, PH | Left | Visual2 | 0 |
| L_TPOJ1 | Temporo-Parieto-Occipital Junction, TPOJ1 | Left | Language | 0 |
| L_a47r | Inferior Frontal, a47r | Left | Frontoparietal | 0 |
| R_V3A | Dorsal Stream Visual, V3A | Right | Visual2 | 0 |
| L_PGp | Inferior Parietal, PGp | Left | Dorsal_Attention | 0 |
| L_IP2 | Inferior Parietal, IP2 | Left | Frontoparietal | 0 |
| R_55b | Premotor, 55b | Right | Language | 0 |
| L_IP0 | Inferior Parietal, IP0 | Left | Dorsal_Attention | 0 |
| L_PFop | Inferior Parietal, PFop | Left | Cingulo-Opercular | 0 |
| R_PEF | Premotor, PEF | Right | Cingulo-Opercular | 0 |
| L_PGi | Inferior Parietal, PGi | Left | Default | 0 |
| R_46 | Dorsolateral Prefrontal, 46 | Right | Cingulo-Opercular | 0 |
| L_V6A | Dorsal Stream Visual, V6A | Left | Visual2 | 0 |
| L_VMV1 | Ventral Stream Visual, VMV1 | Left | Visual2 | 0 |
| L_VMV3 | Ventral Stream Visual, VMV3 | Left | Visual2 | 0 |
| L_PHA2 | Medial Temporal, PHA2 | Left | Default | 0 |
| R_3b | Somatosensory and Motor, 3b | Right | Somatomotor | 0 |
| R_4 | Somatosensory and Motor, 4 | Right | Somatomotor | 0 |
| L_V3CD | MT+ Complex and Neighboring Visual Areas, V3CD | Left | Visual2 | 0 |
| L_VMV2 | Ventral Stream Visual, VMV2 | Left | Visual2 | 0 |
| L_VVC | Ventral Stream Visual, VVC | Left | Visual2 | 0 |
| L_6r | Premotor, 6r | Left | Cingulo-Opercular | 0 |
| R_PreS | Medial Temporal, PreS | Right | Default | 0 |
| L_47l | Inferior Frontal, 47l | Left | Default | 0 |
| R_VMV1 | Ventral Stream Visual, VMV1 | Right | Visual2 | 0 |
| R_PHA2 | Medial Temporal, PHA2 | Right | Default | 0 |
| R_8C | Dorsolateral Prefrontal, 8C | Right | Frontoparietal | 0 |
| R_PFt | Inferior Parietal, PFt | Right | Dorsal_Attention | 0 |
| R_LIPv | Superior Parietal, LIPv | Right | Visual2 | 0 |
| R_31pd | Posterior Cingulate, 31pd | Right | Default | 0 |
| R_31a | Posterior Cingulate, 31a | Right | Frontoparietal | 0 |
| R_VVC | Ventral Stream Visual, VVC | Right | Visual2 | 0 |
| R_25 | Anterior Cingulate and Medial Prefrontal, 25 | Right | Default | 0 |
| R_s32 | Anterior Cingulate and Medial Prefrontal, s32 | Right | Default | 0 |
| R_pOFC | Anterior Cingulate and Medial Prefrontal, pOFC | Right | Orbito-Affective | 0 |
| R_PoI1 | Insular and Frontal Opercular, PoI1 | Right | Cingulo-Opercular | 0 |
| R_Ig | Insular and Frontal Opercular, Ig | Right | Somatomotor | 0 |
| R_FOP5 | Insular and Frontal Opercular, FOP5 | Right | Cingulo-Opercular | 0 |
| R_7PC | Superior Parietal, 7PC | Right | Somatomotor | 0 |
| R_p47r | Inferior Frontal, p47r | Right | Frontoparietal | 0 |
| R_7PL | Superior Parietal, 7Pl | Right | Dorsal_Attention | 0 |
| R_7Am | Superior Parietal, 7Am | Right | Cingulo-Opercular | 0 |
| R_A4 | Auditory Association, A4 | Right | Auditory | 0 |
| R_STSva | Auditory Association, STSva | Right | Default | 0 |
| R_PI | Insular and Frontal Opercular, PI | Right | Cingulo-Opercular | 0 |
| R_a32pr | Anterior Cingulate and Medial Prefrontal, a32pr | Right | Cingulo-Opercular | 0 |
| R_p24 | Anterior Cingulate and Medial Prefrontal, p24 | Right | Cingulo-Opercular | 0 |
| L_MST | MT+ Complex and Neighboring Visual Areas, MST | Left | Visual2 | 0 |
| L_V6 | Dorsal Stream Visual, V6 | Left | Visual2 | 0 |
| R_6ma | Paracentral Lobular and Mid Cingulate, 6ma | Right | Cingulo-Opercular | 0 |
| L_V4 | Early Visual, V4 | Left | Visual2 | 0 |
| R_SCEF | Paracentral Lobular and Mid Cingulate, SCEF | Right | Cingulo-Opercular | 0 |
| R_VMV3 | Ventral Stream Visual, VMV3 | Right | Visual2 | 0 |
| R_V6A | Dorsal Stream Visual, V6A | Right | Visual2 | 0 |
| L_45 | Inferior Frontal, 45 | Left | Language | 0 |
| R_PGs | Inferior Parietal, PGs | Right | Default | 0 |
| R_ProS | Posterior Cingulate, ProS | Right | Visual1 | 0 |
| R_9m | Anterior Cingulate and Medial Prefrontal, 9m | Right | Default | 0 |
| R_STGa | Auditory Association, STGa | Right | Language | 0 |
| R_PBelt | Early Auditory, PBelt | Right | Auditory | 0 |
| R_10r | Anterior Cingulate and Medial Prefrontal, 10r | Right | Default | 0 |
| R_8BM | Anterior Cingulate and Medial Prefrontal, 8BM | Right | Frontoparietal | 0 |
| R_STSvp | Auditory Association, STSvp | Right | Default | 0 |
| R_TGd | Lateral Temporal, TGd | Right | Default | 0 |
| R_p32pr | Anterior Cingulate and Medial Prefrontal, p32pr | Right | Cingulo-Opercular | 0 |
| R_TE1p | Lateral Temporal, TE1p | Right | Frontoparietal | 0 |
| R_TE2a | Lateral Temporal, TE2a | Right | Default | 0 |
| R_TF | Medial Temporal, TF | Right | Ventral_Multimodal | 0 |
| R_10d | Orbital and Polar Frontal, 10d | Right | Default | 0 |
| R_a24pr | Anterior Cingulate and Medial Prefrontal, a24pr | Right | Cingulo-Opercular | 0 |
| R_PH | MT+ Complex and Neighboring Visual Areas, PH | Right | Visual2 | 0 |
| R_p24pr | Anterior Cingulate and Medial Prefrontal, p24pr | Right | Cingulo-Opercular | 0 |
| R_TPOJ2 | Temporo-Parieto-Occipital Junction, TPOJ2 | Right | Posterior_Multimodal | 0 |
| R_TPOJ3 | Temporo-Parieto-Occipital Junction, TPOJ3 | Right | Posterior_Multimodal | 0 |
| R_6mp | Paracentral Lobular and Mid Cingulate, 6mp | Right | Somatomotor | 0 |
| R_6d | Premotor, 6d | Right | Somatomotor | 0 |
| R_3a | Somatosensory and Motor, 3a | Right | Somatomotor | 0 |
| R_IP1 | Inferior Parietal, IP1 | Right | Frontoparietal | 0 |
| R_IP0 | Inferior Parietal, IP0 | Right | Dorsal_Attention | 0 |
| R_2 | Somatosensory and Motor, 2 | Right | Somatomotor | 0 |
| R_1 | Somatosensory and Motor, 1 | Right | Somatomotor | 0 |
| R_PFm | Inferior Parietal, PFm | Right | Frontoparietal | 0 |
| R_PGi | Inferior Parietal, PGi | Right | Default | 0 |
| L_4 | Somatosensory and Motor, 4 | Left | Somatomotor | 0 |
| R_45 | Inferior Frontal, 45 | Right | Language | 0 |
| R_MST | MT+ Complex and Neighboring Visual Areas, MST | Right | Visual2 | 0 |
| L_PEF | Premotor, PEF | Left | Dorsal_Attention | 0 |
| R_23d | Posterior Cingulate, 23d | Right | Default | 0 |
| L_LIPv | Superior Parietal, LIPv | Left | Visual2 | 0 |
| L_VIP | Superior Parietal, VIP | Left | Visual2 | 0 |
| R_POS1 | Posterior Cingulate, POS1 | Right | Default | 0 |
| L_1 | Somatosensory and Motor, 1 | Left | Somatomotor | 0 |
| L_2 | Somatosensory and Motor, 2 | Left | Somatomotor | 0 |
| L_3a | Somatosensory and Motor, 3a | Left | Somatomotor | 0 |
| R_a10p | Orbital and Polar Frontal, a10p | Right | Frontoparietal | 0 |
| R_TA2 | Auditory Association, TA2 | Right | Auditory | 0 |
| L_6v | Premotor, 6v | Left | Somatomotor | 0 |
| R_PoI2 | Insular and Frontal Opercular, PoI2 | Right | Cingulo-Opercular | 0 |
| R_PFcm | Early Auditory, PFcm | Right | Cingulo-Opercular | 0 |
| L_a24pr | Anterior Cingulate and Medial Prefrontal, a24pr | Left | Cingulo-Opercular | 0 |
| L_a24 | Anterior Cingulate and Medial Prefrontal, a24 | Left | Default | 0 |
| L_d32 | Anterior Cingulate and Medial Prefrontal, d32 | Left | Default | 0 |
| L_8BM | Anterior Cingulate and Medial Prefrontal, 8BM | Left | Frontoparietal | 0 |
| L_p32 | Anterior Cingulate and Medial Prefrontal, p32 | Left | Default | 0 |
| L_10r | Anterior Cingulate and Medial Prefrontal, 10r | Left | Default | 0 |
| L_47m | Orbital and Polar Frontal, 47m | Left | Default | 0 |
| L_8Av | Dorsolateral Prefrontal, 8Av | Left | Default | 0 |
| L_8Ad | Dorsolateral Prefrontal, 8Ad | Left | Default | 0 |
| R_H | Medial Temporal, H | Right | Default | 0 |
| R_RI | Early Auditory, RI | Right | Somatomotor | 0 |
| L_9p | Dorsolateral Prefrontal, 9p | Left | Default | 0 |
| R_MT | MT+ Complex and Neighboring Visual Areas, MT | Right | Visual2 | 0 |
| L_8C | Dorsolateral Prefrontal, 8C | Left | Frontoparietal | 0 |
| R_PIT | Ventral Stream Visual, PIT | Right | Visual2 | 0 |
| L_7Am | Superior Parietal, 7Am | Left | Cingulo-Opercular | 0 |
| L_6ma | Paracentral Lobular and Mid Cingulate, 6ma | Left | Cingulo-Opercular | 0 |
| R_FOP4 | Insular and Frontal Opercular, FOP4 | Right | Cingulo-Opercular | 0 |
| L_PSL | Temporo-Parieto Occipital Junction, PSL | Left | Language | 0 |
| L_V3A | Dorsal Stream Visual, V3A | Left | Visual2 | 0 |
| L_RSC | Posterior Cingulate, RSC | Left | Frontoparietal | 0 |
| L_POS2 | Posterior Cingulate, POS2 | Left | Frontoparietal | 0 |
| L_V7 | Dorsal Stream Visual, V7 | Left | Visual2 | 0 |
| L_IPS1 | Dorsal Stream Visual, IPS1 | Left | Visual2 | 0 |
| L_FFC | Ventral Stream Visual, FFC | Left | Visual2 | 0 |
| R_7AL | Superior Parietal, 7AL | Right | Somatomotor | 0 |
| L_LO1 | MT+ Complex and Neighboring Visual Areas, LO1 | Left | Visual2 | 0 |
| R_24dv | Paracentral Lobular and Mid Cingulate, 24dv | Right | Somatomotor | 0 |
| R_24dd | Paracentral Lobular and Mid Cingulate, 24dd | Right | Somatomotor | 0 |
| L_MT | MT+ Complex and Neighboring Visual Areas, MT | Left | Visual2 | 0 |
| L_A1 | Early Auditory, A1 | Left | Auditory | 0 |
| L_SFL | Dorsolateral Prefrontal, SFL | Left | Language | 0 |
| R_MI | Insular and Frontal Opercular, MI | Right | Cingulo-Opercular | 0 |
| R_5L | Paracentral Lobular and Mid Cingulate, 5L | Right | Somatomotor | 0 |
| L_STV | Temporo-Parieto-Occipital Junction, STV | Left | Language | 0 |
| L_7m | Posterior Cingulate, 7m | Left | Default | 0 |
| L_23d | Posterior Cingulate, 23d | Left | Default | 0 |
| R_5m | Paracentral Lobular and Mid Cingulate, 5m | Right | Somatomotor | 0 |
| R_31pv | Posterior Cingulate, 31pv | Right | Default | 0 |
| R_47l | Inferior Frontal, 47l | Right | Default | 0 |
| L_5mv | Paracentral Lobular and Mid Cingulate, 5mv | Left | Cingulo-Opercular | 0 |
| R_Pir | Insular and Frontal Opercular, Pir | Right | Orbito-Affective | 0 |
| L_5L | Paracentral Lobular and Mid Cingulate, 5L | Left | Somatomotor | 0 |
| L_24dd | Paracentral Lobular and Mid Cingulate, 24dd | Left | Somatomotor | 0 |
| L_24dv | Paracentral Lobular and Mid Cingulate, 24dv | Left | Somatomotor | 0 |
| L_6d | Premotor, 6d | Left | Somatomotor | 0 |

***Table S2. Feature importance of the top-performing non-stacked models with with Elastic Net, as indicated by Elastic Net coefficients for the HCP Aging dataset dataset: Facename Encoding vs Distractor Task Contrast***

| **Glasser's label** | **Brain Region** | **Hemisphere** | **Network** | **Amplitude** |
| --- | --- | --- | --- | --- |
| L_VMV1 | Ventral Stream Visual, VMV1 | Left | Visual2 | 0.0377 |
| L_V3CD | MT+ Complex and Neighboring Visual Areas, V3CD | Left | Visual2 | -0.0359 |
| L_MT | MT+ Complex and Neighboring Visual Areas, MT | Left | Visual2 | 0.0328 |
| R_LO3 | MT+ Complex and Neighboring Visual Areas, LO3 | Right | Visual2 | -0.0325 |
| R_LBelt | Early Auditory, LBelt | Right | Auditory | -0.0324 |
| AMYGDALA_RIGHT | Amygdala Right | Right | Subcortex | 0.0321 |
| R_MBelt | Early Auditory, MBelt | Right | Auditory | -0.0319 |
| CEREBELLUM_RIGHT | Cerebellum Right | Right | Subcortex | 0.0314 |
| R_TE1p | Lateral Temporal, TE1p | Right | Frontoparietal | -0.0312 |
| L_VMV2 | Ventral Stream Visual, VMV2 | Left | Visual2 | 0.031 |
| L_6d | Premotor, 6d | Left | Somatomotor | 0.0309 |
| R_FOP1 | Posterior Opercular, FOP1 | Right | Cingulo-Opercular | -0.0306 |
| L_8BM | Anterior Cingulate and Medial Prefrontal, 8BM | Left | Frontoparietal | 0.0299 |
| L_2 | Somatosensory and Motor, 2 | Left | Somatomotor | 0.0295 |
| L_OP1 | Posterior Opercular, OP1 | Left | Somatomotor | 0.0294 |
| L_PCV | Posterior Cingulate, PCV | Left | Posterior_Multimodal | 0.0288 |
| R_LO1 | MT+ Complex and Neighboring Visual Areas, LO1 | Right | Visual2 | -0.0284 |
| R_A1 | Early Auditory, A1 | Right | Auditory | -0.0282 |
| L_6v | Premotor, 6v | Left | Somatomotor | -0.0281 |
| L_H | Medial Temporal, H | Left | Default | 0.0277 |
| L_OP4 | Posterior Opercular, OP4 | Left | Somatomotor | -0.0276 |
| L_d32 | Anterior Cingulate and Medial Prefrontal, d32 | Left | Default | 0.0274 |
| L_TE2a | Lateral Temporal, TE2a | Left | Default | 0.0273 |
| L_6r | Premotor, 6r | Left | Cingulo-Opercular | -0.0272 |
| HIPPOCAMPUS_RIGHT | Hippocampus Right | Right | Subcortex | 0.0272 |
| L_43 | Posterior Opercular, 43 | Left | Cingulo-Opercular | -0.0271 |
| R_VMV3 | Ventral Stream Visual, VMV3 | Right | Visual2 | -0.027 |
| L_47s | Orbital and Polar Frontal, 47s | Left | Default | 0.027 |
| L_FOP2 | Insular and Frontal Opercular, FOP2 | Left | Somatomotor | 0.0265 |
| R_V6A | Dorsal Stream Visual, V6A | Right | Visual2 | -0.0261 |
| R_43 | Posterior Opercular, 43 | Right | Cingulo-Opercular | -0.0261 |
| L_LIPd | Superior Parietal, LIPd | Left | Dorsal_Attention | 0.0257 |
| L_24dd | Paracentral Lobular and Mid Cingulate, 24dd | Left | Somatomotor | 0.0252 |
| R_AIP | Superior Parietal, AIP | Right | Dorsal_Attention | 0.0243 |
| ACCUMBENS_RIGHT | Accumbens Right | Right | Subcortex | 0.0241 |
| R_V3A | Dorsal Stream Visual, V3A | Right | Visual2 | -0.0232 |
| R_1 | Somatosensory and Motor, 1 | Right | Somatomotor | -0.0232 |
| L_PIT | Ventral Stream Visual, PIT | Left | Visual2 | -0.023 |
| L_EC | Medial Temporal, EC | Left | Default | 0.0229 |
| R_ProS | Posterior Cingulate, ProS | Right | Visual1 | 0.0229 |
| R_MST | MT+ Complex and Neighboring Visual Areas, MST | Right | Visual2 | -0.0225 |
| PALLIDUM_LEFT | Pallidum Left | Left | Subcortex | 0.0223 |
| L_STSva | Auditory Association, STSva | Left | Default | -0.0222 |
| L_V2 | Early Visual, V2 | Left | Visual2 | 0.0222 |
| L_24dv | Paracentral Lobular and Mid Cingulate, 24dv | Left | Somatomotor | 0.022 |
| AMYGDALA_LEFT | Amygdala Left | Left | Subcortex | 0.0217 |
| R_TPOJ2 | Temporo-Parieto-Occipital Junction, TPOJ2 | Right | Posterior_Multimodal | 0.0216 |
| R_6ma | Paracentral Lobular and Mid Cingulate, 6ma | Right | Cingulo-Opercular | 0.0213 |
| L_PEF | Premotor, PEF | Left | Dorsal_Attention | 0.0212 |
| L_MI | Insular and Frontal Opercular, MI | Left | Cingulo-Opercular | -0.0211 |
| R_5L | Paracentral Lobular and Mid Cingulate, 5L | Right | Somatomotor | -0.0209 |
| PALLIDUM_RIGHT | Pallidum Right | Right | Subcortex | 0.0205 |
| L_LBelt | Early Auditory, LBelt | Left | Auditory | -0.0204 |
| R_VMV1 | Ventral Stream Visual, VMV1 | Right | Visual2 | 0.0204 |
| L_FOP5 | Insular and Frontal Opercular, FOP5 | Left | Cingulo-Opercular | 0.0203 |
| R_A4 | Auditory Association, A4 | Right | Auditory | -0.02 |
| L_LO1 | MT+ Complex and Neighboring Visual Areas, LO1 | Left | Visual2 | -0.02 |
| L_FEF | Premotor, FEF | Left | Cingulo-Opercular | -0.0197 |
| L_TE2p | Lateral Temporal, TE2p | Left | Dorsal_Attention | 0.0197 |
| BRAIN_STEM | Brain Stem | None | Subcortex | -0.0197 |
| L_PHT | Lateral Temporal, PHT | Left | Dorsal_Attention | 0.0195 |
| R_STV | Temporo-Parieto-Occipital Junction, STV | Right | Posterior_Multimodal | -0.0194 |
| L_1 | Somatosensory and Motor, 1 | Left | Somatomotor | 0.0192 |
| L_TGv | Lateral Temporal, TGv | Left | Language | -0.0191 |
| R_VIP | Superior Parietal, VIP | Right | Visual2 | -0.019 |
| L_TE1m | Lateral Temporal, TE1m | Left | Default | -0.0186 |
| L_3b | Somatosensory and Motor, 3b | Left | Somatomotor | 0.0185 |
| R_PreS | Medial Temporal, PreS | Right | Default | 0.0185 |
| R_LIPv | Superior Parietal, LIPv | Right | Visual2 | -0.0185 |
| R_4 | Somatosensory and Motor, 4 | Right | Somatomotor | -0.0185 |
| L_TE1a | Lateral Temporal, TE1a | Left | Default | -0.0183 |
| R_10v | Anterior Cingulate and Medial Prefrontal, 10v | Right | Default | -0.0181 |
| R_9p | Dorsolateral Prefrontal, 9p | Right | Default | -0.018 |
| R_a10p | Orbital and Polar Frontal, a10p | Right | Frontoparietal | 0.018 |
| L_STSda | Auditory Association, STSda | Left | Language | -0.018 |
| R_i6-8 | Dorsolateral Prefrontal, i6-8 | Right | Frontoparietal | 0.0179 |
| CEREBELLUM_LEFT | Cerebellum Left | Left | Subcortex | 0.0178 |
| R_STSva | Auditory Association, STSva | Right | Default | -0.0177 |
| R_p32 | Anterior Cingulate and Medial Prefrontal, p32 | Right | Default | 0.0176 |
| R_SCEF | Paracentral Lobular and Mid Cingulate, SCEF | Right | Cingulo-Opercular | 0.0176 |
| L_FOP1 | Posterior Opercular, FOP1 | Left | Cingulo-Opercular | 0.0175 |
| CAUDATE_LEFT | Caudate Left | Left | Subcortex | 0.0174 |
| R_8Ad | Dorsolateral Prefrontal, 8Ad | Right | Default | -0.0173 |
| L_PHA1 | Medial Temporal, PHA1 | Left | Default | 0.0173 |
| R_TF | Medial Temporal, TF | Right | Ventral_Multimodal | -0.0173 |
| R_10r | Anterior Cingulate and Medial Prefrontal, 10r | Right | Default | -0.017 |
| R_13l | Orbital and Polar Frontal, 13l | Right | Frontoparietal | 0.017 |
| R_OP4 | Posterior Opercular, OP4 | Right | Somatomotor | -0.0169 |
| R_AVI | Insular and Frontal Opercular, AVI | Right | Frontoparietal | -0.0169 |
| R_6v | Premotor, 6v | Right | Somatomotor | -0.0166 |
| R_p24pr | Anterior Cingulate and Medial Prefrontal, p24pr | Right | Cingulo-Opercular | 0.0165 |
| L_TPOJ1 | Temporo-Parieto-Occipital Junction, TPOJ1 | Left | Language | 0.0163 |
| R_RI | Early Auditory, RI | Right | Somatomotor | -0.0163 |
| L_VIP | Superior Parietal, VIP | Left | Visual2 | -0.0162 |
| R_POS1 | Posterior Cingulate, POS1 | Right | Default | 0.0162 |
| L_7m | Posterior Cingulate, 7m | Left | Default | -0.0161 |
| L_IPS1 | Dorsal Stream Visual, IPS1 | Left | Visual2 | -0.0161 |
| L_PBelt | Early Auditory, PBelt | Left | Auditory | -0.016 |
| R_11l | Orbital and Polar Frontal, 11l | Right | Frontoparietal | 0.0158 |
| L_AVI | Insular and Frontal Opercular, AVI | Left | Frontoparietal | -0.0157 |
| R_FOP5 | Insular and Frontal Opercular, FOP5 | Right | Cingulo-Opercular | -0.0155 |
| L_11l | Orbital and Polar Frontal, 11l | Left | Frontoparietal | 0.0155 |
| L_PHA3 | Medial Temporal, PHA3 | Left | Dorsal_Attention | -0.0155 |
| R_10pp | Orbital and Polar Frontal, 10pp | Right | Default | 0.0154 |
| L_TE1p | Lateral Temporal, TE1p | Left | Frontoparietal | -0.0154 |
| L_TA2 | Auditory Association, TA2 | Left | Auditory | 0.0154 |
| R_45 | Inferior Frontal, 45 | Right | Language | 0.0154 |
| R_DVT | Posterior Cingulate, DVT | Right | Visual1 | -0.0152 |
| L_p24pr | Anterior Cingulate and Medial Prefrontal, p24pr | Left | Cingulo-Opercular | 0.015 |
| R_V2 | Early Visual, V2 | Right | Visual2 | 0.0149 |
| R_PeEc | Medial Temporal, PeEc | Right | Ventral_Multimodal | 0.0149 |
| R_PGi | Inferior Parietal, PGi | Right | Default | -0.0148 |
| R_PBelt | Early Auditory, PBelt | Right | Auditory | -0.0147 |
| R_23c | Paracentral Lobular and Mid Cingulate, 23c | Right | Cingulo-Opercular | 0.0147 |
| R_LO2 | MT+ Complex and Neighboring Visual Areas, LO2 | Right | Visual2 | 0.0146 |
| R_VVC | Ventral Stream Visual, VVC | Right | Visual2 | 0.0145 |
| L_31pv | Posterior Cingulate, 31pv | Left | Default | -0.0145 |
| R_H | Medial Temporal, H | Right | Default | 0.0145 |
| R_V3CD | MT+ Complex and Neighboring Visual Areas, V3CD | Right | Visual2 | -0.0145 |
| R_V7 | Dorsal Stream Visual, V7 | Right | Visual2 | -0.0144 |
| R_52 | Early Auditory, 52 | Right | Auditory | -0.0144 |
| R_V3 | Early Visual, V3 | Right | Visual2 | -0.0144 |
| R_PGs | Inferior Parietal, PGs | Right | Default | 0.0142 |
| L_7PC | Superior Parietal, 7PC | Left | Somatomotor | 0.0142 |
| L_V4 | Early Visual, V4 | Left | Visual2 | -0.0142 |
| R_TE1a | Lateral Temporal, TE1a | Right | Default | -0.0141 |
| R_PSL | Temporo-Parieto Occipital Junction, PSL | Right | Cingulo-Opercular | -0.0141 |
| L_p32pr | Anterior Cingulate and Medial Prefrontal, p32pr | Left | Cingulo-Opercular | -0.0137 |
| L_6mp | Paracentral Lobular and Mid Cingulate, 6mp | Left | Somatomotor | 0.0136 |
| L_MIP | Superior Parietal, MIP | Left | Dorsal_Attention | -0.0136 |
| R_31pd | Posterior Cingulate, 31pd | Right | Default | -0.0136 |
| R_V3B | Dorsal Stream Visual, V3B | Right | Visual2 | -0.0135 |
| R_TPOJ1 | Temporo-Parieto-Occipital Junction, TPOJ1 | Right | Language | -0.0135 |
| L_A5 | Auditory Association, A5 | Left | Language | 0.0135 |
| L_V6 | Dorsal Stream Visual, V6 | Left | Visual2 | 0.0134 |
| R_TGv | Lateral Temporal, TGv | Right | Language | -0.0134 |
| L_STGa | Auditory Association, STGa | Left | Language | 0.0133 |
| L_V3 | Early Visual, V3 | Left | Visual2 | -0.0133 |
| L_FOP4 | Insular and Frontal Opercular, FOP4 | Left | Cingulo-Opercular | -0.0132 |
| L_IFSp | Inferior Frontal, IFSp | Left | Language | 0.0132 |
| R_PH | MT+ Complex and Neighboring Visual Areas, PH | Right | Visual2 | 0.0131 |
| R_FST | MT+ Complex and Neighboring Visual Areas, FST | Right | Visual2 | -0.0131 |
| R_p47r | Inferior Frontal, p47r | Right | Frontoparietal | -0.013 |
| R_STSda | Auditory Association, STSda | Right | Default | -0.013 |
| L_FFC | Ventral Stream Visual, FFC | Left | Visual2 | -0.0129 |
| R_TE2p | Lateral Temporal, TE2p | Right | Dorsal_Attention | 0.0128 |
| R_5mv | Paracentral Lobular and Mid Cingulate, 5mv | Right | Cingulo-Opercular | -0.0127 |
| R_PFcm | Early Auditory, PFcm | Right | Cingulo-Opercular | -0.0127 |
| R_SFL | Dorsolateral Prefrontal, SFL | Right | Language | 0.0127 |
| L_PFm | Inferior Parietal, PFm | Left | Frontoparietal | -0.0126 |
| L_MBelt | Early Auditory, MBelt | Left | Auditory | -0.0125 |
| CAUDATE_RIGHT | Caudate Right | Right | Subcortex | 0.0124 |
| L_p10p | Orbital and Polar Frontal, p10p | Left | Frontoparietal | 0.0124 |
| R_46 | Dorsolateral Prefrontal, 46 | Right | Cingulo-Opercular | 0.0124 |
| R_3b | Somatosensory and Motor, 3b | Right | Somatomotor | -0.0124 |
| L_i6-8 | Dorsolateral Prefrontal, i6-8 | Left | Frontoparietal | -0.0123 |
| R_23d | Posterior Cingulate, 23d | Right | Default | -0.0123 |
| R_IP0 | Inferior Parietal, IP0 | Right | Dorsal_Attention | -0.0122 |
| L_STSdp | Auditory Association, STSdp | Left | Language | 0.0122 |
| L_V3B | Dorsal Stream Visual, V3B | Left | Visual2 | 0.0121 |
| R_A5 | Auditory Association, A5 | Right | Language | -0.0118 |
| R_MT | MT+ Complex and Neighboring Visual Areas, MT | Right | Visual2 | 0.0116 |
| R_MI | Insular and Frontal Opercular, MI | Right | Cingulo-Opercular | -0.0116 |
| THALAMUS_LEFT | Thalamus Left | Left | Subcortex | 0.0115 |
| R_PFm | Inferior Parietal, PFm | Right | Frontoparietal | -0.0115 |
| L_TPOJ3 | Temporo-Parieto-Occipital Junction, TPOJ3 | Left | Posterior_Multimodal | 0.0115 |
| L_a32pr | Anterior Cingulate and Medial Prefrontal, a32pr | Left | Cingulo-Opercular | 0.0114 |
| L_FOP3 | Insular and Frontal Opercular, FOP3 | Left | Cingulo-Opercular | 0.0114 |
| R_33pr | Anterior Cingulate and Medial Prefrontal, 33pr | Right | Frontoparietal | 0.0113 |
| R_5m | Paracentral Lobular and Mid Cingulate, 5m | Right | Somatomotor | -0.0113 |
| R_VMV2 | Ventral Stream Visual, VMV2 | Right | Visual2 | -0.0113 |
| R_TPOJ3 | Temporo-Parieto-Occipital Junction, TPOJ3 | Right | Posterior_Multimodal | 0.0112 |
| L_9m | Anterior Cingulate and Medial Prefrontal, 9m | Left | Default | 0.011 |
| R_p10p | Orbital and Polar Frontal, p10p | Right | Frontoparietal | 0.011 |
| L_PHA2 | Medial Temporal, PHA2 | Left | Default | -0.011 |
| L_PGi | Inferior Parietal, PGi | Left | Default | -0.011 |
| R_47l | Inferior Frontal, 47l | Right | Default | -0.0109 |
| R_a9-46v | Dorsolateral Prefrontal, a9-46v | Right | Frontoparietal | 0.0108 |
| L_V3A | Dorsal Stream Visual, V3A | Left | Visual2 | 0.0107 |
| R_IFSp | Inferior Frontal, IFSp | Right | Frontoparietal | -0.0106 |
| R_PHA3 | Medial Temporal, PHA3 | Right | Dorsal_Attention | 0.0105 |
| L_d23ab | Posterior Cingulate, d23ab | Left | Default | 0.0104 |
| R_24dd | Paracentral Lobular and Mid Cingulate, 24dd | Right | Somatomotor | -0.0104 |
| R_3a | Somatosensory and Motor, 3a | Right | Somatomotor | -0.0102 |
| R_8BM | Anterior Cingulate and Medial Prefrontal, 8BM | Right | Frontoparietal | 0.0102 |
| L_5L | Paracentral Lobular and Mid Cingulate, 5L | Left | Somatomotor | -0.0101 |
| L_IP2 | Inferior Parietal, IP2 | Left | Frontoparietal | 0.01 |
| L_PFop | Inferior Parietal, PFop | Left | Cingulo-Opercular | -0.01 |
| R_7PC | Superior Parietal, 7PC | Right | Somatomotor | -0.01 |
| R_AAIC | Insular and Frontal Opercular, AAIC | Right | Orbito-Affective | -0.0099 |
| R_25 | Anterior Cingulate and Medial Prefrontal, 25 | Right | Default | -0.0098 |
| L_5mv | Paracentral Lobular and Mid Cingulate, 5mv | Left | Cingulo-Opercular | -0.0098 |
| L_3a | Somatosensory and Motor, 3a | Left | Somatomotor | 0.0097 |
| L_a10p | Orbital and Polar Frontal, a10p | Left | Frontoparietal | -0.0097 |
| R_47m | Orbital and Polar Frontal, 47m | Right | Default | 0.0095 |
| R_9-46d | Dorsolateral Prefrontal, 9-46d | Right | Cingulo-Opercular | -0.0093 |
| L_Pir | Insular and Frontal Opercular, Pir | Left | Orbito-Affective | 0.0093 |
| R_IFJa | Inferior Frontal, IFJa | Right | Language | -0.0093 |
| L_a24 | Anterior Cingulate and Medial Prefrontal, a24 | Left | Default | 0.0092 |
| R_MIP | Superior Parietal, MIP | Right | Dorsal_Attention | -0.0091 |
| R_FOP3 | Insular and Frontal Opercular, FOP3 | Right | Cingulo-Opercular | 0.0091 |
| L_13l | Orbital and Polar Frontal, 13l | Left | Frontoparietal | 0.009 |
| R_V6 | Dorsal Stream Visual, V6 | Right | Visual2 | 0.0088 |
| R_FOP2 | Insular and Frontal Opercular, FOP2 | Right | Somatomotor | 0.0087 |
| L_ProS | Posterior Cingulate, ProS | Left | Visual1 | 0.0087 |
| L_RSC | Posterior Cingulate, RSC | Left | Frontoparietal | -0.0087 |
| R_PCV | Posterior Cingulate, PCV | Right | Posterior_Multimodal | -0.0087 |
| R_FEF | Premotor, FEF | Right | Cingulo-Opercular | 0.0087 |
| R_8Av | Dorsolateral Prefrontal, 8Av | Right | Default | -0.0086 |
| R_LIPd | Superior Parietal, LIPd | Right | Dorsal_Attention | 0.0086 |
| R_RSC | Posterior Cingulate, RSC | Right | Frontoparietal | 0.0086 |
| L_p9-46v | Dorsolateral Prefrontal, p9-46v | Left | Frontoparietal | -0.0086 |
| L_IFSa | Inferior Frontal, IFSa | Left | Frontoparietal | -0.0084 |
| L_45 | Inferior Frontal, 45 | Left | Language | 0.0084 |
| R_6mp | Paracentral Lobular and Mid Cingulate, 6mp | Right | Somatomotor | -0.0084 |
| R_7m | Posterior Cingulate, 7m | Right | Default | -0.0082 |
| L_PGp | Inferior Parietal, PGp | Left | Dorsal_Attention | 0.0082 |
| R_PHT | Lateral Temporal, PHT | Right | Dorsal_Attention | 0.0082 |
| THALAMUS_RIGHT | Thalamus Right | Right | Subcortex | -0.0081 |
| L_pOFC | Anterior Cingulate and Medial Prefrontal, pOFC | Left | Orbito-Affective | 0.008 |
| L_s6-8 | Dorsolateral Prefrontal, s6-8 | Left | Frontoparietal | -0.0079 |
| R_PoI2 | Insular and Frontal Opercular, PoI2 | Right | Cingulo-Opercular | 0.0078 |
| L_IFJp | Inferior Frontal, IFJp | Left | Frontoparietal | 0.0078 |
| L_PSL | Temporo-Parieto Occipital Junction, PSL | Left | Language | 0.0078 |
| L_PFcm | Early Auditory, PFcm | Left | Cingulo-Opercular | -0.0077 |
| L_8Av | Dorsolateral Prefrontal, 8Av | Left | Default | -0.0077 |
| L_a9-46v | Dorsolateral Prefrontal, a9-46v | Left | Frontoparietal | 0.0077 |
| L_v23ab | Posterior Cingulate, v23ab | Left | Default | -0.0076 |
| L_52 | Early Auditory, 52 | Left | Auditory | 0.0076 |
| L_V6A | Dorsal Stream Visual, V6A | Left | Visual2 | 0.0072 |
| L_TPOJ2 | Temporo-Parieto-Occipital Junction, TPOJ2 | Left | Posterior_Multimodal | 0.0072 |
| L_POS1 | Posterior Cingulate, POS1 | Left | Default | 0.0071 |
| R_IFSa | Inferior Frontal, IFSa | Right | Cingulo-Opercular | -0.007 |
| L_PGs | Inferior Parietal, PGs | Left | Default | -0.007 |
| L_STSvp | Auditory Association, STSvp | Left | Default | 0.0069 |
| L_V1 | Primary Visual, V1 | Left | Visual1 | 0.0068 |
| L_5m | Paracentral Lobular and Mid Cingulate, 5m | Left | Somatomotor | -0.0068 |
| L_23c | Paracentral Lobular and Mid Cingulate, 23c | Left | Cingulo-Opercular | 0.0068 |
| L_AIP | Superior Parietal, AIP | Left | Dorsal_Attention | 0.0068 |
| R_TGd | Lateral Temporal, TGd | Right | Default | -0.0067 |
| R_EC | Medial Temporal, EC | Right | Default | -0.0067 |
| L_STV | Temporo-Parieto-Occipital Junction, STV | Left | Language | 0.0067 |
| L_25 | Anterior Cingulate and Medial Prefrontal, 25 | Left | Default | 0.0067 |
| DIENCEPHALON_VENTRAL_RIGHT | Diencephalon Ventral Right | Right | Subcortex | 0.0066 |
| R_d32 | Anterior Cingulate and Medial Prefrontal, d32 | Right | Frontoparietal | -0.0066 |
| HIPPOCAMPUS_LEFT | Hippocampus Left | Left | Subcortex | 0.0065 |
| L_VVC | Ventral Stream Visual, VVC | Left | Visual2 | 0.0064 |
| R_V1 | Primary Visual, V1 | Right | Visual1 | 0.0063 |
| R_IFJp | Inferior Frontal, IFJp | Right | Frontoparietal | 0.0063 |
| L_IP1 | Inferior Parietal, IP1 | Left | Frontoparietal | -0.0063 |
| L_9p | Dorsolateral Prefrontal, 9p | Left | Default | -0.0062 |
| R_a24 | Anterior Cingulate and Medial Prefrontal, a24 | Right | Default | 0.0061 |
| R_PF | Inferior Parietal, PF | Right | Cingulo-Opercular | -0.006 |
| R_p24 | Anterior Cingulate and Medial Prefrontal, p24 | Right | Cingulo-Opercular | 0.006 |
| R_TE1m | Lateral Temporal, TE1m | Right | Frontoparietal | -0.0059 |
| ACCUMBENS_LEFT | Accumbens Left | Left | Subcortex | 0.0059 |
| R_8C | Dorsolateral Prefrontal, 8C | Right | Frontoparietal | -0.0058 |
| R_9a | Dorsolateral Prefrontal, 9a | Right | Default | -0.0058 |
| L_7PL | Superior Parietal, 7Pl | Left | Dorsal_Attention | -0.0057 |
| R_IP1 | Inferior Parietal, IP1 | Right | Frontoparietal | 0.0057 |
| L_PeEc | Medial Temporal, PeEc | Left | Ventral_Multimodal | -0.0056 |
| R_PEF | Premotor, PEF | Right | Cingulo-Opercular | -0.0056 |
| R_PoI1 | Insular and Frontal Opercular, PoI1 | Right | Cingulo-Opercular | 0.0055 |
| R_7AL | Superior Parietal, 7AL | Right | Somatomotor | -0.0054 |
| L_p32 | Anterior Cingulate and Medial Prefrontal, p32 | Left | Default | 0.0053 |
| R_OFC | Orbital and Polar Frontal, OFC | Right | Frontoparietal | 0.0053 |
| L_MST | MT+ Complex and Neighboring Visual Areas, MST | Left | Visual2 | -0.005 |
| L_OFC | Orbital and Polar Frontal, OFC | Left | Default | -0.0049 |
| L_33pr | Anterior Cingulate and Medial Prefrontal, 33pr | Left | Cingulo-Opercular | 0.0049 |
| R_STGa | Auditory Association, STGa | Right | Language | -0.0049 |
| L_10d | Orbital and Polar Frontal, 10d | Left | Default | -0.0048 |
| R_a32pr | Anterior Cingulate and Medial Prefrontal, a32pr | Right | Cingulo-Opercular | 0.0048 |
| R_d23ab | Posterior Cingulate, d23ab | Right | Default | -0.0048 |
| L_a24pr | Anterior Cingulate and Medial Prefrontal, a24pr | Left | Cingulo-Opercular | -0.0048 |
| L_SCEF | Paracentral Lobular and Mid Cingulate, SCEF | Left | Cingulo-Opercular | 0.0047 |
| L_PI | Insular and Frontal Opercular, PI | Left | Cingulo-Opercular | 0.0046 |
| R_31pv | Posterior Cingulate, 31pv | Right | Default | 0.0046 |
| L_7Pm | Superior Parietal, 7Pm | Left | Frontoparietal | 0.0046 |
| L_PoI2 | Insular and Frontal Opercular, PoI2 | Left | Cingulo-Opercular | 0.0046 |
| L_RI | Early Auditory, RI | Left | Auditory | -0.0044 |
| R_s32 | Anterior Cingulate and Medial Prefrontal, s32 | Right | Default | -0.0044 |
| R_PFop | Inferior Parietal, PFop | Right | Cingulo-Opercular | -0.0044 |
| PUTAMEN_RIGHT | Putamen Right | Right | Subcortex | -0.0043 |
| L_47m | Orbital and Polar Frontal, 47m | Left | Default | 0.0042 |
| R_55b | Premotor, 55b | Right | Language | -0.0042 |
| L_A4 | Auditory Association, A4 | Left | Auditory | -0.0042 |
| R_STSvp | Auditory Association, STSvp | Right | Default | -0.0041 |
| L_POS2 | Posterior Cingulate, POS2 | Left | Frontoparietal | -0.0041 |
| R_FFC | Ventral Stream Visual, FFC | Right | Visual2 | -0.0041 |
| R_PHA1 | Medial Temporal, PHA1 | Right | Default | -0.004 |
| R_p9-46v | Dorsolateral Prefrontal, p9-46v | Right | Frontoparietal | 0.004 |
| R_47s | Orbital and Polar Frontal, 47s | Right | Default | 0.0039 |
| R_IP2 | Inferior Parietal, IP2 | Right | Frontoparietal | 0.0038 |
| L_TF | Medial Temporal, TF | Left | Ventral_Multimodal | 0.0038 |
| L_p47r | Inferior Frontal, p47r | Left | Frontoparietal | -0.0038 |
| L_DVT | Posterior Cingulate, DVT | Left | Visual1 | -0.0037 |
| L_4 | Somatosensory and Motor, 4 | Left | Somatomotor | -0.0037 |
| R_PFt | Inferior Parietal, PFt | Right | Dorsal_Attention | 0.0037 |
| L_SFL | Dorsolateral Prefrontal, SFL | Left | Language | -0.0036 |
| PUTAMEN_LEFT | Putamen Left | Left | Subcortex | -0.0036 |
| R_V4 | Early Visual, V4 | Right | Visual2 | 0.0036 |
| L_23d | Posterior Cingulate, 23d | Left | Default | -0.0035 |
| R_FOP4 | Insular and Frontal Opercular, FOP4 | Right | Cingulo-Opercular | -0.0035 |
| L_OP2-3 | Posterior Opercular, OP2-3 | Left | Somatomotor | 0.0034 |
| L_AAIC | Insular and Frontal Opercular, AAIC | Left | Orbito-Affective | -0.0034 |
| L_PoI1 | Insular and Frontal Opercular, PoI1 | Left | Cingulo-Opercular | 0.0034 |
| R_7Am | Superior Parietal, 7Am | Right | Cingulo-Opercular | 0.0033 |
| R_IPS1 | Dorsal Stream Visual, IPS1 | Right | Visual2 | 0.0033 |
| R_V8 | Ventral Stream Visual, V8 | Right | Visual2 | -0.0032 |
| L_LIPv | Superior Parietal, LIPv | Left | Visual2 | -0.0032 |
| R_POS2 | Posterior Cingulate, POS2 | Right | Frontoparietal | 0.0032 |
| R_PHA2 | Medial Temporal, PHA2 | Right | Default | -0.003 |
| DIENCEPHALON_VENTRAL_LEFT | Diencephalon Ventral Left | Left | Subcortex | 0.003 |
| L_p24 | Anterior Cingulate and Medial Prefrontal, p24 | Left | Cingulo-Opercular | 0.003 |
| L_IP0 | Inferior Parietal, IP0 | Left | Dorsal_Attention | 0.0029 |
| R_v23ab | Posterior Cingulate, v23ab | Right | Default | 0.0029 |
| L_31a | Posterior Cingulate, 31a | Left | Default | -0.0029 |
| L_LO2 | MT+ Complex and Neighboring Visual Areas, LO2 | Left | Visual2 | -0.0029 |
| R_2 | Somatosensory and Motor, 2 | Right | Somatomotor | -0.0028 |
| L_46 | Dorsolateral Prefrontal, 46 | Left | Cingulo-Opercular | 0.0028 |
| L_A1 | Early Auditory, A1 | Left | Auditory | -0.0027 |
| R_OP1 | Posterior Opercular, OP1 | Right | Somatomotor | -0.0027 |
| L_55b | Premotor, 55b | Left | Language | 0.0026 |
| L_PF | Inferior Parietal, PF | Left | Cingulo-Opercular | 0.0026 |
| R_TA2 | Auditory Association, TA2 | Right | Auditory | 0.0026 |
| L_7Am | Superior Parietal, 7Am | Left | Cingulo-Opercular | -0.0025 |
| L_s32 | Anterior Cingulate and Medial Prefrontal, s32 | Left | Default | -0.0025 |
| R_STSdp | Auditory Association, STSdp | Right | Language | 0.0025 |
| L_a47r | Inferior Frontal, a47r | Left | Frontoparietal | -0.0025 |
| L_V4t | MT+ Complex and Neighboring Visual Areas, V4t | Left | Visual2 | -0.0024 |
| L_31pd | Posterior Cingulate, 31pd | Left | Default | 0.0024 |
| R_Pir | Insular and Frontal Opercular, Pir | Right | Orbito-Affective | 0.0023 |
| R_44 | Inferior Frontal, 44 | Right | Frontoparietal | -0.0023 |
| R_PI | Insular and Frontal Opercular, PI | Right | Cingulo-Opercular | -0.0022 |
| L_6ma | Paracentral Lobular and Mid Cingulate, 6ma | Left | Cingulo-Opercular | -0.0022 |
| R_8BL | Dorsolateral Prefrontal, 8BL | Right | Default | -0.0021 |
| R_TE2a | Lateral Temporal, TE2a | Right | Default | -0.0021 |
| L_8C | Dorsolateral Prefrontal, 8C | Left | Frontoparietal | 0.0021 |
| R_V4t | MT+ Complex and Neighboring Visual Areas, V4t | Right | Visual2 | -0.002 |
| L_10pp | Orbital and Polar Frontal, 10pp | Left | Default | 0.002 |
| L_V8 | Ventral Stream Visual, V8 | Left | Visual2 | 0.002 |
| R_31a | Posterior Cingulate, 31a | Right | Frontoparietal | 0.0019 |
| R_24dv | Paracentral Lobular and Mid Cingulate, 24dv | Right | Somatomotor | -0.0019 |
| L_IFJa | Inferior Frontal, IFJa | Left | Language | -0.0018 |
| R_10d | Orbital and Polar Frontal, 10d | Right | Default | -0.0018 |
| R_9m | Anterior Cingulate and Medial Prefrontal, 9m | Right | Default | -0.0018 |
| L_47l | Inferior Frontal, 47l | Left | Default | -0.0017 |
| L_PFt | Inferior Parietal, PFt | Left | Dorsal_Attention | -0.0017 |
| L_6a | Premotor, 6a | Left | Dorsal_Attention | -0.0017 |
| R_7Pm | Superior Parietal, 7Pm | Right | Frontoparietal | -0.0016 |
| R_a47r | Inferior Frontal, a47r | Right | Frontoparietal | -0.0015 |
| R_s6-8 | Dorsolateral Prefrontal, s6-8 | Right | Frontoparietal | 0.0015 |
| R_6r | Premotor, 6r | Right | Cingulo-Opercular | 0.0013 |
| R_6a | Premotor, 6a | Right | Dorsal_Attention | 0.0013 |
| L_8Ad | Dorsolateral Prefrontal, 8Ad | Left | Default | 0.0012 |
| L_V7 | Dorsal Stream Visual, V7 | Left | Visual2 | -0.001 |
| L_10v | Anterior Cingulate and Medial Prefrontal, 10v | Left | Default | 0.0009 |
| L_10r | Anterior Cingulate and Medial Prefrontal, 10r | Left | Default | 0.0008 |
| L_PH | MT+ Complex and Neighboring Visual Areas, PH | Left | Visual2 | 0.0008 |
| R_Ig | Insular and Frontal Opercular, Ig | Right | Somatomotor | 0.0006 |
| L_9-46d | Dorsolateral Prefrontal, 9-46d | Left | Cingulo-Opercular | -0.0006 |
| R_PGp | Inferior Parietal, PGp | Right | Dorsal_Attention | 0.0006 |
| L_9a | Dorsolateral Prefrontal, 9a | Left | Default | -0.0005 |
| L_VMV3 | Ventral Stream Visual, VMV3 | Left | Visual2 | 0.0005 |
| L_8BL | Dorsolateral Prefrontal, 8BL | Left | Default | 0.0004 |
| L_44 | Inferior Frontal, 44 | Left | Language | 0.0004 |
| R_p32pr | Anterior Cingulate and Medial Prefrontal, p32pr | Right | Cingulo-Opercular | -0.0003 |
| R_a24pr | Anterior Cingulate and Medial Prefrontal, a24pr | Right | Cingulo-Opercular | -0.0002 |
| L_LO3 | MT+ Complex and Neighboring Visual Areas, LO3 | Left | Visual2 | -0.0002 |
| R_OP2-3 | Posterior Opercular, OP2-3 | Right | Somatomotor | -0.0002 |
| L_TGd | Lateral Temporal, TGd | Left | Default | 0.0002 |
| L_Ig | Insular and Frontal Opercular, Ig | Left | Somatomotor | 0.0002 |
| R_6d | Premotor, 6d | Right | Somatomotor | 0.0001 |
| L_FST | MT+ Complex and Neighboring Visual Areas, FST | Left | Visual2 | 0.0001 |
| L_PreS | Medial Temporal, PreS | Left | Default | -0.0001 |
| R_pOFC | Anterior Cingulate and Medial Prefrontal, pOFC | Right | Orbito-Affective | -0.0001 |
| R_PIT | Ventral Stream Visual, PIT | Right | Visual2 | -0.0001 |
| L_7AL | Superior Parietal, 7AL | Left | Somatomotor | 0 |
| R_7PL | Superior Parietal, 7Pl | Right | Dorsal_Attention | 0 |

***Table S3. Feature importance of the top-performing non-stacked models with with Elastic Net, as indicated by Elastic Net coefficients for the Dunedin Study dataset, predicting cognitive abilities at 45 years old.: Facename Encoding vs Distractor Task Contrast***

| **Glasser's label** | **Brain Region** | **Hemisphere** | **Network** | **Amplitude** |
| --- | --- | --- | --- | --- |
| HIPPOCAMPUS_LEFT | Hippocampus Left | Left | Subcortex | -0.0319 |
| R_LIPd | Superior Parietal, LIPd | Right | Dorsal_Attention | 0.0304 |
| R_FST | MT+ Complex and Neighboring Visual Areas, FST | Right | Visual2 | 0.0302 |
| L_V3B | Dorsal Stream Visual, V3B | Left | Visual2 | -0.0292 |
| L_6v | Premotor, 6v | Left | Somatomotor | -0.0281 |
| HIPPOCAMPUS_RIGHT | Hippocampus Right | Right | Subcortex | -0.0271 |
| L_AVI | Insular and Frontal Opercular, AVI | Left | Frontoparietal | 0.025 |
| CEREBELLUM_LEFT | Cerebellum Left | Left | Subcortex | 0.025 |
| DIENCEPHALON_VENTRAL_LEFT | Diencephalon Ventral Left | Left | Subcortex | 0.0243 |
| L_IP2 | Inferior Parietal, IP2 | Left | Frontoparietal | 0.0239 |
| R_IFJp | Inferior Frontal, IFJp | Right | Frontoparietal | 0.0239 |
| L_43 | Posterior Opercular, 43 | Left | Cingulo-Opercular | -0.0232 |
| L_a32pr | Anterior Cingulate and Medial Prefrontal, a32pr | Left | Cingulo-Opercular | 0.0232 |
| R_LO1 | MT+ Complex and Neighboring Visual Areas, LO1 | Right | Visual2 | 0.0228 |
| R_IFSa | Inferior Frontal, IFSa | Right | Cingulo-Opercular | 0.0221 |
| R_MIP | Superior Parietal, MIP | Right | Dorsal_Attention | 0.0218 |
| R_PH | MT+ Complex and Neighboring Visual Areas, PH | Right | Visual2 | 0.0214 |
| L_LBelt | Early Auditory, LBelt | Left | Auditory | 0.0211 |
| R_IPS1 | Dorsal Stream Visual, IPS1 | Right | Visual2 | 0.0211 |
| L_RI | Early Auditory, RI | Left | Auditory | 0.021 |
| L_LO3 | MT+ Complex and Neighboring Visual Areas, LO3 | Left | Visual2 | 0.0209 |
| L_LIPd | Superior Parietal, LIPd | Left | Dorsal_Attention | 0.0209 |
| R_VMV2 | Ventral Stream Visual, VMV2 | Right | Visual2 | -0.0207 |
| L_PSL | Temporo-Parieto Occipital Junction, PSL | Left | Language | 0.0206 |
| L_LO1 | MT+ Complex and Neighboring Visual Areas, LO1 | Left | Visual2 | 0.0204 |
| R_7PL | Superior Parietal, 7Pl | Right | Dorsal_Attention | 0.0204 |
| L_8Ad | Dorsolateral Prefrontal, 8Ad | Left | Default | -0.0203 |
| L_IFJp | Inferior Frontal, IFJp | Left | Frontoparietal | 0.0202 |
| R_8C | Dorsolateral Prefrontal, 8C | Right | Frontoparietal | 0.0196 |
| R_3a | Somatosensory and Motor, 3a | Right | Somatomotor | -0.0191 |
| R_STSvp | Auditory Association, STSvp | Right | Default | -0.019 |
| R_AIP | Superior Parietal, AIP | Right | Dorsal_Attention | 0.0188 |
| L_MST | MT+ Complex and Neighboring Visual Areas, MST | Left | Visual2 | -0.0183 |
| L_PGs | Inferior Parietal, PGs | Left | Default | -0.0182 |
| L_TPOJ1 | Temporo-Parieto-Occipital Junction, TPOJ1 | Left | Language | 0.018 |
| L_AIP | Superior Parietal, AIP | Left | Dorsal_Attention | 0.0176 |
| L_IFJa | Inferior Frontal, IFJa | Left | Language | 0.0175 |
| R_LIPv | Superior Parietal, LIPv | Right | Visual2 | 0.0175 |
| R_AVI | Insular and Frontal Opercular, AVI | Right | Frontoparietal | 0.0174 |
| R_PGi | Inferior Parietal, PGi | Right | Default | -0.0173 |
| L_8BM | Anterior Cingulate and Medial Prefrontal, 8BM | Left | Frontoparietal | 0.0169 |
| L_V2 | Early Visual, V2 | Left | Visual2 | -0.0169 |
| L_V7 | Dorsal Stream Visual, V7 | Left | Visual2 | -0.0168 |
| R_8BM | Anterior Cingulate and Medial Prefrontal, 8BM | Right | Frontoparietal | 0.0167 |
| L_V1 | Primary Visual, V1 | Left | Visual1 | -0.0165 |
| R_13l | Orbital and Polar Frontal, 13l | Right | Frontoparietal | -0.0165 |
| L_PFm | Inferior Parietal, PFm | Left | Frontoparietal | -0.0164 |
| R_6r | Premotor, 6r | Right | Cingulo-Opercular | 0.0164 |
| R_TE1m | Lateral Temporal, TE1m | Right | Frontoparietal | -0.0164 |
| R_8BL | Dorsolateral Prefrontal, 8BL | Right | Default | -0.0163 |
| L_p24pr | Anterior Cingulate and Medial Prefrontal, p24pr | Left | Cingulo-Opercular | 0.0161 |
| L_52 | Early Auditory, 52 | Left | Auditory | -0.016 |
| R_IFJa | Inferior Frontal, IFJa | Right | Language | 0.0159 |
| L_33pr | Anterior Cingulate and Medial Prefrontal, 33pr | Left | Cingulo-Opercular | 0.0157 |
| R_i6-8 | Dorsolateral Prefrontal, i6-8 | Right | Frontoparietal | 0.0156 |
| L_POS1 | Posterior Cingulate, POS1 | Left | Default | -0.0156 |
| R_a32pr | Anterior Cingulate and Medial Prefrontal, a32pr | Right | Cingulo-Opercular | 0.0154 |
| R_6v | Premotor, 6v | Right | Somatomotor | -0.0154 |
| L_8C | Dorsolateral Prefrontal, 8C | Left | Frontoparietal | 0.0152 |
| R_6a | Premotor, 6a | Right | Dorsal_Attention | 0.0152 |
| THALAMUS_RIGHT | Thalamus Right | Right | Subcortex | 0.0151 |
| PALLIDUM_LEFT | Pallidum Left | Left | Subcortex | -0.0151 |
| R_31pd | Posterior Cingulate, 31pd | Right | Default | -0.0149 |
| R_A4 | Auditory Association, A4 | Right | Auditory | -0.0149 |
| R_V6A | Dorsal Stream Visual, V6A | Right | Visual2 | -0.0149 |
| L_FOP1 | Posterior Opercular, FOP1 | Left | Cingulo-Opercular | -0.0147 |
| R_OP4 | Posterior Opercular, OP4 | Right | Somatomotor | -0.0146 |
| R_43 | Posterior Opercular, 43 | Right | Cingulo-Opercular | -0.0146 |
| L_PIT | Ventral Stream Visual, PIT | Left | Visual2 | -0.0145 |
| CEREBELLUM_RIGHT | Cerebellum Right | Right | Subcortex | 0.0145 |
| L_TPOJ3 | Temporo-Parieto-Occipital Junction, TPOJ3 | Left | Posterior_Multimodal | 0.0144 |
| R_IP2 | Inferior Parietal, IP2 | Right | Frontoparietal | 0.0143 |
| R_9p | Dorsolateral Prefrontal, 9p | Right | Default | -0.0143 |
| R_MT | MT+ Complex and Neighboring Visual Areas, MT | Right | Visual2 | -0.014 |
| R_VVC | Ventral Stream Visual, VVC | Right | Visual2 | 0.014 |
| R_ProS | Posterior Cingulate, ProS | Right | Visual1 | 0.0139 |
| R_TPOJ3 | Temporo-Parieto-Occipital Junction, TPOJ3 | Right | Posterior_Multimodal | 0.0138 |
| ACCUMBENS_LEFT | Accumbens Left | Left | Subcortex | 0.0137 |
| L_VMV3 | Ventral Stream Visual, VMV3 | Left | Visual2 | -0.0136 |
| L_OP2-3 | Posterior Opercular, OP2-3 | Left | Somatomotor | -0.0135 |
| L_A5 | Auditory Association, A5 | Left | Language | 0.0135 |
| L_VMV2 | Ventral Stream Visual, VMV2 | Left | Visual2 | -0.0133 |
| R_Pir | Insular and Frontal Opercular, Pir | Right | Orbito-Affective | 0.0133 |
| R_TE1a | Lateral Temporal, TE1a | Right | Default | -0.0132 |
| R_TGv | Lateral Temporal, TGv | Right | Language | -0.0132 |
| L_TGd | Lateral Temporal, TGd | Left | Default | -0.0132 |
| L_d32 | Anterior Cingulate and Medial Prefrontal, d32 | Left | Default | 0.013 |
| R_V7 | Dorsal Stream Visual, V7 | Right | Visual2 | -0.013 |
| R_SFL | Dorsolateral Prefrontal, SFL | Right | Language | -0.013 |
| R_4 | Somatosensory and Motor, 4 | Right | Somatomotor | -0.0129 |
| L_PoI1 | Insular and Frontal Opercular, PoI1 | Left | Cingulo-Opercular | -0.0129 |
| L_6ma | Paracentral Lobular and Mid Cingulate, 6ma | Left | Cingulo-Opercular | -0.0128 |
| R_PHT | Lateral Temporal, PHT | Right | Dorsal_Attention | 0.0128 |
| R_s6-8 | Dorsolateral Prefrontal, s6-8 | Right | Frontoparietal | -0.0128 |
| DIENCEPHALON_VENTRAL_RIGHT | Diencephalon Ventral Right | Right | Subcortex | 0.0128 |
| L_6d | Premotor, 6d | Left | Somatomotor | 0.0128 |
| R_STGa | Auditory Association, STGa | Right | Language | -0.0127 |
| L_SCEF | Paracentral Lobular and Mid Cingulate, SCEF | Left | Cingulo-Opercular | 0.0127 |
| R_46 | Dorsolateral Prefrontal, 46 | Right | Cingulo-Opercular | 0.0127 |
| R_31a | Posterior Cingulate, 31a | Right | Frontoparietal | -0.0126 |
| L_OP1 | Posterior Opercular, OP1 | Left | Somatomotor | -0.0125 |
| R_TE1p | Lateral Temporal, TE1p | Right | Frontoparietal | 0.0125 |
| L_TE1a | Lateral Temporal, TE1a | Left | Default | -0.0125 |
| L_TF | Medial Temporal, TF | Left | Ventral_Multimodal | -0.0124 |
| L_FFC | Ventral Stream Visual, FFC | Left | Visual2 | 0.0123 |
| CAUDATE_LEFT | Caudate Left | Left | Subcortex | 0.0121 |
| R_VMV3 | Ventral Stream Visual, VMV3 | Right | Visual2 | -0.012 |
| L_FEF | Premotor, FEF | Left | Cingulo-Opercular | 0.012 |
| L_STSdp | Auditory Association, STSdp | Left | Language | 0.0119 |
| L_TE1m | Lateral Temporal, TE1m | Left | Default | -0.0117 |
| L_v23ab | Posterior Cingulate, v23ab | Left | Default | 0.0117 |
| R_PoI1 | Insular and Frontal Opercular, PoI1 | Right | Cingulo-Opercular | -0.0116 |
| R_p32 | Anterior Cingulate and Medial Prefrontal, p32 | Right | Default | -0.0116 |
| R_PFm | Inferior Parietal, PFm | Right | Frontoparietal | -0.0116 |
| R_IFSp | Inferior Frontal, IFSp | Right | Frontoparietal | 0.0116 |
| L_s6-8 | Dorsolateral Prefrontal, s6-8 | Left | Frontoparietal | -0.0115 |
| R_V3B | Dorsal Stream Visual, V3B | Right | Visual2 | -0.0115 |
| R_FFC | Ventral Stream Visual, FFC | Right | Visual2 | 0.0115 |
| R_PFop | Inferior Parietal, PFop | Right | Cingulo-Opercular | -0.0114 |
| R_OP1 | Posterior Opercular, OP1 | Right | Somatomotor | -0.0114 |
| R_7PC | Superior Parietal, 7PC | Right | Somatomotor | 0.0113 |
| L_PBelt | Early Auditory, PBelt | Left | Auditory | 0.0112 |
| L_TGv | Lateral Temporal, TGv | Left | Language | -0.0112 |
| L_9p | Dorsolateral Prefrontal, 9p | Left | Default | -0.0111 |
| L_31a | Posterior Cingulate, 31a | Left | Default | -0.011 |
| R_v23ab | Posterior Cingulate, v23ab | Right | Default | -0.011 |
| R_9m | Anterior Cingulate and Medial Prefrontal, 9m | Right | Default | -0.0109 |
| L_PGi | Inferior Parietal, PGi | Left | Default | -0.0109 |
| L_a24 | Anterior Cingulate and Medial Prefrontal, a24 | Left | Default | 0.0108 |
| R_MST | MT+ Complex and Neighboring Visual Areas, MST | Right | Visual2 | -0.0108 |
| AMYGDALA_RIGHT | Amygdala Right | Right | Subcortex | -0.0107 |
| L_5L | Paracentral Lobular and Mid Cingulate, 5L | Left | Somatomotor | -0.0107 |
| L_5m | Paracentral Lobular and Mid Cingulate, 5m | Left | Somatomotor | -0.0106 |
| L_A1 | Early Auditory, A1 | Left | Auditory | -0.0106 |
| R_STSva | Auditory Association, STSva | Right | Default | -0.0105 |
| R_TGd | Lateral Temporal, TGd | Right | Default | -0.0105 |
| L_7Am | Superior Parietal, 7Am | Left | Cingulo-Opercular | 0.0105 |
| L_V8 | Ventral Stream Visual, V8 | Left | Visual2 | 0.0104 |
| L_s32 | Anterior Cingulate and Medial Prefrontal, s32 | Left | Default | -0.0104 |
| R_TE2a | Lateral Temporal, TE2a | Right | Default | -0.0103 |
| THALAMUS_LEFT | Thalamus Left | Left | Subcortex | 0.0101 |
| R_LO3 | MT+ Complex and Neighboring Visual Areas, LO3 | Right | Visual2 | 0.0101 |
| L_6a | Premotor, 6a | Left | Dorsal_Attention | -0.0101 |
| L_V6A | Dorsal Stream Visual, V6A | Left | Visual2 | 0.01 |
| L_3b | Somatosensory and Motor, 3b | Left | Somatomotor | -0.0099 |
| L_10d | Orbital and Polar Frontal, 10d | Left | Default | -0.0099 |
| L_31pv | Posterior Cingulate, 31pv | Left | Default | -0.0098 |
| R_p9-46v | Dorsolateral Prefrontal, p9-46v | Right | Frontoparietal | 0.0097 |
| L_V4 | Early Visual, V4 | Left | Visual2 | -0.0096 |
| L_PFop | Inferior Parietal, PFop | Left | Cingulo-Opercular | 0.0095 |
| R_POS1 | Posterior Cingulate, POS1 | Right | Default | 0.0095 |
| R_V2 | Early Visual, V2 | Right | Visual2 | -0.0094 |
| R_PIT | Ventral Stream Visual, PIT | Right | Visual2 | -0.0094 |
| R_LO2 | MT+ Complex and Neighboring Visual Areas, LO2 | Right | Visual2 | 0.0093 |
| R_10v | Anterior Cingulate and Medial Prefrontal, 10v | Right | Default | -0.0093 |
| L_STSvp | Auditory Association, STSvp | Left | Default | 0.0093 |
| R_POS2 | Posterior Cingulate, POS2 | Right | Frontoparietal | 0.0092 |
| R_6d | Premotor, 6d | Right | Somatomotor | 0.0091 |
| L_p24 | Anterior Cingulate and Medial Prefrontal, p24 | Left | Cingulo-Opercular | 0.0091 |
| R_PFt | Inferior Parietal, PFt | Right | Dorsal_Attention | 0.0091 |
| R_45 | Inferior Frontal, 45 | Right | Language | 0.0091 |
| R_FEF | Premotor, FEF | Right | Cingulo-Opercular | 0.009 |
| L_STSva | Auditory Association, STSva | Left | Default | -0.0089 |
| L_IP0 | Inferior Parietal, IP0 | Left | Dorsal_Attention | -0.0087 |
| R_MI | Insular and Frontal Opercular, MI | Right | Cingulo-Opercular | 0.0087 |
| PALLIDUM_RIGHT | Pallidum Right | Right | Subcortex | 0.0086 |
| R_STSdp | Auditory Association, STSdp | Right | Language | 0.0086 |
| L_STGa | Auditory Association, STGa | Left | Language | -0.0085 |
| L_a10p | Orbital and Polar Frontal, a10p | Left | Frontoparietal | 0.0083 |
| R_PSL | Temporo-Parieto Occipital Junction, PSL | Right | Cingulo-Opercular | -0.0083 |
| R_MBelt | Early Auditory, MBelt | Right | Auditory | 0.0083 |
| L_DVT | Posterior Cingulate, DVT | Left | Visual1 | 0.0083 |
| L_7AL | Superior Parietal, 7AL | Left | Somatomotor | 0.0083 |
| L_i6-8 | Dorsolateral Prefrontal, i6-8 | Left | Frontoparietal | 0.0082 |
| L_FOP5 | Insular and Frontal Opercular, FOP5 | Left | Cingulo-Opercular | 0.0081 |
| L_FOP4 | Insular and Frontal Opercular, FOP4 | Left | Cingulo-Opercular | 0.0081 |
| L_ProS | Posterior Cingulate, ProS | Left | Visual1 | 0.0081 |
| R_p10p | Orbital and Polar Frontal, p10p | Right | Frontoparietal | -0.008 |
| L_24dd | Paracentral Lobular and Mid Cingulate, 24dd | Left | Somatomotor | -0.0079 |
| L_V6 | Dorsal Stream Visual, V6 | Left | Visual2 | -0.0078 |
| L_VVC | Ventral Stream Visual, VVC | Left | Visual2 | 0.0078 |
| R_pOFC | Anterior Cingulate and Medial Prefrontal, pOFC | Right | Orbito-Affective | 0.0078 |
| R_3b | Somatosensory and Motor, 3b | Right | Somatomotor | -0.0078 |
| L_p32 | Anterior Cingulate and Medial Prefrontal, p32 | Left | Default | -0.0077 |
| L_OFC | Orbital and Polar Frontal, OFC | Left | Default | -0.0076 |
| R_10pp | Orbital and Polar Frontal, 10pp | Right | Default | -0.0076 |
| L_LO2 | MT+ Complex and Neighboring Visual Areas, LO2 | Left | Visual2 | 0.0075 |
| L_9m | Anterior Cingulate and Medial Prefrontal, 9m | Left | Default | 0.0075 |
| L_pOFC | Anterior Cingulate and Medial Prefrontal, pOFC | Left | Orbito-Affective | -0.0074 |
| L_a9-46v | Dorsolateral Prefrontal, a9-46v | Left | Frontoparietal | -0.0073 |
| L_13l | Orbital and Polar Frontal, 13l | Left | Frontoparietal | 0.0073 |
| L_10v | Anterior Cingulate and Medial Prefrontal, 10v | Left | Default | -0.0073 |
| L_IPS1 | Dorsal Stream Visual, IPS1 | Left | Visual2 | 0.0073 |
| L_8BL | Dorsolateral Prefrontal, 8BL | Left | Default | -0.0072 |
| L_9-46d | Dorsolateral Prefrontal, 9-46d | Left | Cingulo-Opercular | -0.0072 |
| R_EC | Medial Temporal, EC | Right | Default | 0.0071 |
| R_5m | Paracentral Lobular and Mid Cingulate, 5m | Right | Somatomotor | 0.007 |
| L_7PC | Superior Parietal, 7PC | Left | Somatomotor | -0.007 |
| R_TA2 | Auditory Association, TA2 | Right | Auditory | -0.0069 |
| R_PreS | Medial Temporal, PreS | Right | Default | 0.0069 |
| L_PFcm | Early Auditory, PFcm | Left | Cingulo-Opercular | -0.0069 |
| R_FOP3 | Insular and Frontal Opercular, FOP3 | Right | Cingulo-Opercular | -0.0068 |
| R_11l | Orbital and Polar Frontal, 11l | Right | Frontoparietal | 0.0067 |
| R_a24 | Anterior Cingulate and Medial Prefrontal, a24 | Right | Default | -0.0067 |
| R_RSC | Posterior Cingulate, RSC | Right | Frontoparietal | 0.0067 |
| L_45 | Inferior Frontal, 45 | Left | Language | -0.0066 |
| L_TE1p | Lateral Temporal, TE1p | Left | Frontoparietal | -0.0066 |
| L_VMV1 | Ventral Stream Visual, VMV1 | Left | Visual2 | -0.0066 |
| R_5L | Paracentral Lobular and Mid Cingulate, 5L | Right | Somatomotor | 0.0066 |
| L_d23ab | Posterior Cingulate, d23ab | Left | Default | 0.0065 |
| PUTAMEN_LEFT | Putamen Left | Left | Subcortex | -0.0065 |
| R_d32 | Anterior Cingulate and Medial Prefrontal, d32 | Right | Frontoparietal | 0.0064 |
| R_A5 | Auditory Association, A5 | Right | Language | -0.0064 |
| R_OP2-3 | Posterior Opercular, OP2-3 | Right | Somatomotor | 0.0064 |
| R_V3A | Dorsal Stream Visual, V3A | Right | Visual2 | -0.0063 |
| L_PCV | Posterior Cingulate, PCV | Left | Posterior_Multimodal | -0.0063 |
| L_47s | Orbital and Polar Frontal, 47s | Left | Default | -0.0063 |
| L_POS2 | Posterior Cingulate, POS2 | Left | Frontoparietal | 0.0062 |
| L_PHA2 | Medial Temporal, PHA2 | Left | Default | -0.0062 |
| L_Pir | Insular and Frontal Opercular, Pir | Left | Orbito-Affective | 0.0061 |
| L_p9-46v | Dorsolateral Prefrontal, p9-46v | Left | Frontoparietal | -0.0061 |
| L_8Av | Dorsolateral Prefrontal, 8Av | Left | Default | -0.006 |
| L_23d | Posterior Cingulate, 23d | Left | Default | 0.006 |
| R_V6 | Dorsal Stream Visual, V6 | Right | Visual2 | -0.006 |
| L_31pd | Posterior Cingulate, 31pd | Left | Default | -0.006 |
| R_8Ad | Dorsolateral Prefrontal, 8Ad | Right | Default | -0.006 |
| L_PEF | Premotor, PEF | Left | Dorsal_Attention | -0.006 |
| L_3a | Somatosensory and Motor, 3a | Left | Somatomotor | -0.0059 |
| R_PEF | Premotor, PEF | Right | Cingulo-Opercular | 0.0059 |
| R_PHA1 | Medial Temporal, PHA1 | Right | Default | 0.0058 |
| R_STV | Temporo-Parieto-Occipital Junction, STV | Right | Posterior_Multimodal | -0.0057 |
| R_7Pm | Superior Parietal, 7Pm | Right | Frontoparietal | 0.0057 |
| R_55b | Premotor, 55b | Right | Language | -0.0057 |
| R_PCV | Posterior Cingulate, PCV | Right | Posterior_Multimodal | -0.0056 |
| R_47m | Orbital and Polar Frontal, 47m | Right | Default | -0.0056 |
| R_TE2p | Lateral Temporal, TE2p | Right | Dorsal_Attention | 0.0055 |
| R_PGp | Inferior Parietal, PGp | Right | Dorsal_Attention | -0.0055 |
| R_a47r | Inferior Frontal, a47r | Right | Frontoparietal | 0.0055 |
| R_DVT | Posterior Cingulate, DVT | Right | Visual1 | 0.0055 |
| L_44 | Inferior Frontal, 44 | Left | Language | -0.0054 |
| R_V8 | Ventral Stream Visual, V8 | Right | Visual2 | -0.0052 |
| R_10d | Orbital and Polar Frontal, 10d | Right | Default | -0.0052 |
| R_5mv | Paracentral Lobular and Mid Cingulate, 5mv | Right | Cingulo-Opercular | -0.0051 |
| L_STV | Temporo-Parieto-Occipital Junction, STV | Left | Language | -0.0051 |
| L_24dv | Paracentral Lobular and Mid Cingulate, 24dv | Left | Somatomotor | -0.0051 |
| L_PHT | Lateral Temporal, PHT | Left | Dorsal_Attention | 0.005 |
| L_TE2a | Lateral Temporal, TE2a | Left | Default | -0.005 |
| L_FST | MT+ Complex and Neighboring Visual Areas, FST | Left | Visual2 | 0.0049 |
| R_Ig | Insular and Frontal Opercular, Ig | Right | Somatomotor | -0.0049 |
| L_55b | Premotor, 55b | Left | Language | 0.0049 |
| L_2 | Somatosensory and Motor, 2 | Left | Somatomotor | -0.0048 |
| R_23c | Paracentral Lobular and Mid Cingulate, 23c | Right | Cingulo-Opercular | -0.0048 |
| L_STSda | Auditory Association, STSda | Left | Language | -0.0048 |
| R_V1 | Primary Visual, V1 | Right | Visual1 | 0.0048 |
| R_8Av | Dorsolateral Prefrontal, 8Av | Right | Default | 0.0048 |
| R_9a | Dorsolateral Prefrontal, 9a | Right | Default | 0.0048 |
| R_7AL | Superior Parietal, 7AL | Right | Somatomotor | 0.0047 |
| R_FOP5 | Insular and Frontal Opercular, FOP5 | Right | Cingulo-Opercular | 0.0047 |
| R_23d | Posterior Cingulate, 23d | Right | Default | -0.0047 |
| R_52 | Early Auditory, 52 | Right | Auditory | -0.0046 |
| L_OP4 | Posterior Opercular, OP4 | Left | Somatomotor | -0.0046 |
| L_p32pr | Anterior Cingulate and Medial Prefrontal, p32pr | Left | Cingulo-Opercular | 0.0046 |
| CAUDATE_RIGHT | Caudate Right | Right | Subcortex | -0.0044 |
| L_PoI2 | Insular and Frontal Opercular, PoI2 | Left | Cingulo-Opercular | -0.0044 |
| R_V4 | Early Visual, V4 | Right | Visual2 | 0.0044 |
| L_6r | Premotor, 6r | Left | Cingulo-Opercular | 0.0043 |
| R_TF | Medial Temporal, TF | Right | Ventral_Multimodal | -0.0043 |
| R_6mp | Paracentral Lobular and Mid Cingulate, 6mp | Right | Somatomotor | -0.0043 |
| L_10r | Anterior Cingulate and Medial Prefrontal, 10r | Left | Default | -0.0042 |
| L_6mp | Paracentral Lobular and Mid Cingulate, 6mp | Left | Somatomotor | -0.0042 |
| R_a10p | Orbital and Polar Frontal, a10p | Right | Frontoparietal | 0.0041 |
| L_TE2p | Lateral Temporal, TE2p | Left | Dorsal_Attention | 0.004 |
| L_FOP3 | Insular and Frontal Opercular, FOP3 | Left | Cingulo-Opercular | -0.004 |
| R_PHA3 | Medial Temporal, PHA3 | Right | Dorsal_Attention | -0.0039 |
| R_7m | Posterior Cingulate, 7m | Right | Default | 0.0039 |
| L_PGp | Inferior Parietal, PGp | Left | Dorsal_Attention | -0.0037 |
| L_IP1 | Inferior Parietal, IP1 | Left | Frontoparietal | -0.0037 |
| R_d23ab | Posterior Cingulate, d23ab | Right | Default | 0.0037 |
| L_IFSp | Inferior Frontal, IFSp | Left | Language | 0.0037 |
| L_a24pr | Anterior Cingulate and Medial Prefrontal, a24pr | Left | Cingulo-Opercular | 0.0037 |
| L_TPOJ2 | Temporo-Parieto-Occipital Junction, TPOJ2 | Left | Posterior_Multimodal | 0.0036 |
| R_H | Medial Temporal, H | Right | Default | -0.0036 |
| L_46 | Dorsolateral Prefrontal, 46 | Left | Cingulo-Opercular | 0.0036 |
| L_V3A | Dorsal Stream Visual, V3A | Left | Visual2 | -0.0035 |
| R_p24pr | Anterior Cingulate and Medial Prefrontal, p24pr | Right | Cingulo-Opercular | -0.0035 |
| L_V3CD | MT+ Complex and Neighboring Visual Areas, V3CD | Left | Visual2 | 0.0035 |
| R_FOP4 | Insular and Frontal Opercular, FOP4 | Right | Cingulo-Opercular | -0.0035 |
| L_MT | MT+ Complex and Neighboring Visual Areas, MT | Left | Visual2 | 0.0035 |
| L_PreS | Medial Temporal, PreS | Left | Default | 0.0034 |
| R_2 | Somatosensory and Motor, 2 | Right | Somatomotor | -0.0033 |
| R_PHA2 | Medial Temporal, PHA2 | Right | Default | -0.0033 |
| L_PeEc | Medial Temporal, PeEc | Left | Ventral_Multimodal | 0.0033 |
| L_47m | Orbital and Polar Frontal, 47m | Left | Default | 0.0032 |
| R_A1 | Early Auditory, A1 | Right | Auditory | 0.0032 |
| R_OFC | Orbital and Polar Frontal, OFC | Right | Frontoparietal | -0.0032 |
| R_33pr | Anterior Cingulate and Medial Prefrontal, 33pr | Right | Frontoparietal | 0.0031 |
| L_PF | Inferior Parietal, PF | Left | Cingulo-Opercular | 0.0031 |
| L_MIP | Superior Parietal, MIP | Left | Dorsal_Attention | -0.003 |
| R_31pv | Posterior Cingulate, 31pv | Right | Default | 0.003 |
| R_SCEF | Paracentral Lobular and Mid Cingulate, SCEF | Right | Cingulo-Opercular | 0.0029 |
| R_PGs | Inferior Parietal, PGs | Right | Default | -0.0028 |
| R_PF | Inferior Parietal, PF | Right | Cingulo-Opercular | -0.0028 |
| R_PI | Insular and Frontal Opercular, PI | Right | Cingulo-Opercular | 0.0028 |
| R_24dv | Paracentral Lobular and Mid Cingulate, 24dv | Right | Somatomotor | -0.0028 |
| L_AAIC | Insular and Frontal Opercular, AAIC | Left | Orbito-Affective | 0.0027 |
| L_RSC | Posterior Cingulate, RSC | Left | Frontoparietal | 0.0027 |
| L_PH | MT+ Complex and Neighboring Visual Areas, PH | Left | Visual2 | 0.0027 |
| R_V3 | Early Visual, V3 | Right | Visual2 | -0.0026 |
| L_5mv | Paracentral Lobular and Mid Cingulate, 5mv | Left | Cingulo-Opercular | -0.0026 |
| R_RI | Early Auditory, RI | Right | Somatomotor | -0.0026 |
| BRAIN_STEM | Brain Stem | None | Subcortex | -0.0025 |
| L_V4t | MT+ Complex and Neighboring Visual Areas, V4t | Left | Visual2 | -0.0025 |
| R_47s | Orbital and Polar Frontal, 47s | Right | Default | -0.0025 |
| R_VIP | Superior Parietal, VIP | Right | Visual2 | 0.0025 |
| R_STSda | Auditory Association, STSda | Right | Default | 0.0024 |
| R_VMV1 | Ventral Stream Visual, VMV1 | Right | Visual2 | -0.0024 |
| L_11l | Orbital and Polar Frontal, 11l | Left | Frontoparietal | 0.0024 |
| L_MBelt | Early Auditory, MBelt | Left | Auditory | 0.0023 |
| L_10pp | Orbital and Polar Frontal, 10pp | Left | Default | -0.0021 |
| R_44 | Inferior Frontal, 44 | Right | Frontoparietal | 0.002 |
| L_PFt | Inferior Parietal, PFt | Left | Dorsal_Attention | 0.002 |
| L_PHA3 | Medial Temporal, PHA3 | Left | Dorsal_Attention | 0.002 |
| R_TPOJ2 | Temporo-Parieto-Occipital Junction, TPOJ2 | Right | Posterior_Multimodal | 0.0019 |
| R_PFcm | Early Auditory, PFcm | Right | Cingulo-Opercular | 0.0018 |
| L_47l | Inferior Frontal, 47l | Left | Default | 0.0018 |
| L_FOP2 | Insular and Frontal Opercular, FOP2 | Left | Somatomotor | -0.0018 |
| R_LBelt | Early Auditory, LBelt | Right | Auditory | -0.0018 |
| R_V4t | MT+ Complex and Neighboring Visual Areas, V4t | Right | Visual2 | -0.0018 |
| R_9-46d | Dorsolateral Prefrontal, 9-46d | Right | Cingulo-Opercular | 0.0018 |
| L_PI | Insular and Frontal Opercular, PI | Left | Cingulo-Opercular | 0.0016 |
| R_24dd | Paracentral Lobular and Mid Cingulate, 24dd | Right | Somatomotor | -0.0016 |
| L_EC | Medial Temporal, EC | Left | Default | -0.0014 |
| R_10r | Anterior Cingulate and Medial Prefrontal, 10r | Right | Default | 0.0014 |
| R_a9-46v | Dorsolateral Prefrontal, a9-46v | Right | Frontoparietal | 0.0014 |
| L_9a | Dorsolateral Prefrontal, 9a | Left | Default | -0.0014 |
| R_1 | Somatosensory and Motor, 1 | Right | Somatomotor | -0.0014 |
| R_p32pr | Anterior Cingulate and Medial Prefrontal, p32pr | Right | Cingulo-Opercular | -0.0014 |
| R_FOP1 | Posterior Opercular, FOP1 | Right | Cingulo-Opercular | -0.0014 |
| R_25 | Anterior Cingulate and Medial Prefrontal, 25 | Right | Default | -0.0013 |
| L_25 | Anterior Cingulate and Medial Prefrontal, 25 | Left | Default | -0.0013 |
| L_4 | Somatosensory and Motor, 4 | Left | Somatomotor | -0.0012 |
| L_VIP | Superior Parietal, VIP | Left | Visual2 | -0.0012 |
| L_TA2 | Auditory Association, TA2 | Left | Auditory | 0.0012 |
| L_a47r | Inferior Frontal, a47r | Left | Frontoparietal | -0.0012 |
| L_A4 | Auditory Association, A4 | Left | Auditory | -0.0012 |
| L_MI | Insular and Frontal Opercular, MI | Left | Cingulo-Opercular | 0.0012 |
| L_p47r | Inferior Frontal, p47r | Left | Frontoparietal | -0.0012 |
| L_LIPv | Superior Parietal, LIPv | Left | Visual2 | -0.0012 |
| L_IFSa | Inferior Frontal, IFSa | Left | Frontoparietal | 0.0011 |
| R_IP0 | Inferior Parietal, IP0 | Right | Dorsal_Attention | -0.0011 |
| R_V3CD | MT+ Complex and Neighboring Visual Areas, V3CD | Right | Visual2 | -0.0011 |
| L_V3 | Early Visual, V3 | Left | Visual2 | -0.001 |
| R_TPOJ1 | Temporo-Parieto-Occipital Junction, TPOJ1 | Right | Language | -0.0009 |
| AMYGDALA_LEFT | Amygdala Left | Left | Subcortex | 0.0009 |
| R_47l | Inferior Frontal, 47l | Right | Default | 0.0008 |
| PUTAMEN_RIGHT | Putamen Right | Right | Subcortex | 0.0008 |
| L_1 | Somatosensory and Motor, 1 | Left | Somatomotor | 0.0008 |
| R_IP1 | Inferior Parietal, IP1 | Right | Frontoparietal | -0.0007 |
| L_SFL | Dorsolateral Prefrontal, SFL | Left | Language | -0.0007 |
| R_PoI2 | Insular and Frontal Opercular, PoI2 | Right | Cingulo-Opercular | 0.0005 |
| L_23c | Paracentral Lobular and Mid Cingulate, 23c | Left | Cingulo-Opercular | 0.0005 |
| R_AAIC | Insular and Frontal Opercular, AAIC | Right | Orbito-Affective | 0.0005 |
| R_PBelt | Early Auditory, PBelt | Right | Auditory | 0.0005 |
| L_Ig | Insular and Frontal Opercular, Ig | Left | Somatomotor | -0.0004 |
| R_PeEc | Medial Temporal, PeEc | Right | Ventral_Multimodal | 0.0004 |
| R_6ma | Paracentral Lobular and Mid Cingulate, 6ma | Right | Cingulo-Opercular | -0.0004 |
| R_p47r | Inferior Frontal, p47r | Right | Frontoparietal | -0.0003 |
| R_a24pr | Anterior Cingulate and Medial Prefrontal, a24pr | Right | Cingulo-Opercular | 0.0003 |
| L_7Pm | Superior Parietal, 7Pm | Left | Frontoparietal | 0.0003 |
| ACCUMBENS_RIGHT | Accumbens Right | Right | Subcortex | -0.0003 |
| L_H | Medial Temporal, H | Left | Default | -0.0003 |
| R_7Am | Superior Parietal, 7Am | Right | Cingulo-Opercular | 0.0003 |
| R_p24 | Anterior Cingulate and Medial Prefrontal, p24 | Right | Cingulo-Opercular | -0.0002 |
| R_s32 | Anterior Cingulate and Medial Prefrontal, s32 | Right | Default | 0.0001 |
| L_p10p | Orbital and Polar Frontal, p10p | Left | Frontoparietal | 0.0001 |
| L_7PL | Superior Parietal, 7Pl | Left | Dorsal_Attention | 0.0001 |
| L_PHA1 | Medial Temporal, PHA1 | Left | Default | 0.0001 |
| R_FOP2 | Insular and Frontal Opercular, FOP2 | Right | Somatomotor | 0.0001 |
| L_7m | Posterior Cingulate, 7m | Left | Default | 0 |

***Table S4. Feature importance of the top-performing non-stacked models with with Elastic Net, as indicated by Elastic Net coefficients for the Dunedin Study dataset, predicting cognitive abilities at 7, 9, 11 years old.: Facename Encoding vs Distractor Task Contrast***

| **Glasser's label** | **Brain Region** | **Hemisphere** | **Network** | **Amplitude** |
| --- | --- | --- | --- | --- |
| L_6v | Premotor, 6v | Left | Somatomotor | -0.0492 |
| R_PH | MT+ Complex and Neighboring Visual Areas, PH | Right | Visual2 | 0.0437 |
| R_FST | MT+ Complex and Neighboring Visual Areas, FST | Right | Visual2 | 0.043 |
| DIENCEPHALON_VENTRAL_LEFT | Diencephalon Ventral Left | Left | Subcortex | 0.0428 |
| R_STSvp | Auditory Association, STSvp | Right | Default | -0.0411 |
| L_AVI | Insular and Frontal Opercular, AVI | Left | Frontoparietal | 0.0382 |
| R_MIP | Superior Parietal, MIP | Right | Dorsal_Attention | 0.0379 |
| R_7PL | Superior Parietal, 7Pl | Right | Dorsal_Attention | 0.0373 |
| R_LO1 | MT+ Complex and Neighboring Visual Areas, LO1 | Right | Visual2 | 0.036 |
| R_8BM | Anterior Cingulate and Medial Prefrontal, 8BM | Right | Frontoparietal | 0.0357 |
| HIPPOCAMPUS_LEFT | Hippocampus Left | Left | Subcortex | -0.0352 |
| L_IFJa | Inferior Frontal, IFJa | Left | Language | 0.0346 |
| L_RI | Early Auditory, RI | Left | Auditory | 0.0334 |
| R_AVI | Insular and Frontal Opercular, AVI | Right | Frontoparietal | 0.0332 |
| L_a32pr | Anterior Cingulate and Medial Prefrontal, a32pr | Left | Cingulo-Opercular | 0.0332 |
| L_V8 | Ventral Stream Visual, V8 | Left | Visual2 | 0.0322 |
| L_43 | Posterior Opercular, 43 | Left | Cingulo-Opercular | -0.031 |
| L_V2 | Early Visual, V2 | Left | Visual2 | -0.0309 |
| L_OP1 | Posterior Opercular, OP1 | Left | Somatomotor | -0.0307 |
| R_FFC | Ventral Stream Visual, FFC | Right | Visual2 | 0.0307 |
| R_VMV2 | Ventral Stream Visual, VMV2 | Right | Visual2 | -0.0305 |
| L_IP0 | Inferior Parietal, IP0 | Left | Dorsal_Attention | -0.0304 |
| L_PSL | Temporo-Parieto Occipital Junction, PSL | Left | Language | 0.0303 |
| R_PFm | Inferior Parietal, PFm | Right | Frontoparietal | -0.0301 |
| L_PCV | Posterior Cingulate, PCV | Left | Posterior_Multimodal | -0.0299 |
| R_IPS1 | Dorsal Stream Visual, IPS1 | Right | Visual2 | 0.0291 |
| L_31a | Posterior Cingulate, 31a | Left | Default | -0.029 |
| L_A1 | Early Auditory, A1 | Left | Auditory | -0.0289 |
| R_STSdp | Auditory Association, STSdp | Right | Language | 0.0288 |
| R_STGa | Auditory Association, STGa | Right | Language | -0.0283 |
| L_V3B | Dorsal Stream Visual, V3B | Left | Visual2 | -0.0283 |
| ACCUMBENS_LEFT | Accumbens Left | Left | Subcortex | 0.0281 |
| L_7AL | Superior Parietal, 7AL | Left | Somatomotor | 0.0277 |
| L_PFm | Inferior Parietal, PFm | Left | Frontoparietal | -0.0276 |
| R_PCV | Posterior Cingulate, PCV | Right | Posterior_Multimodal | -0.0273 |
| R_8C | Dorsolateral Prefrontal, 8C | Right | Frontoparietal | 0.0271 |
| R_A4 | Auditory Association, A4 | Right | Auditory | -0.0268 |
| R_V7 | Dorsal Stream Visual, V7 | Right | Visual2 | -0.0266 |
| L_6d | Premotor, 6d | Left | Somatomotor | 0.0266 |
| L_8C | Dorsolateral Prefrontal, 8C | Left | Frontoparietal | 0.0266 |
| L_v23ab | Posterior Cingulate, v23ab | Left | Default | 0.0263 |
| L_A5 | Auditory Association, A5 | Left | Language | 0.0248 |
| L_PoI1 | Insular and Frontal Opercular, PoI1 | Left | Cingulo-Opercular | -0.0247 |
| R_IFJa | Inferior Frontal, IFJa | Right | Language | 0.0246 |
| L_LO3 | MT+ Complex and Neighboring Visual Areas, LO3 | Left | Visual2 | 0.0245 |
| R_PGi | Inferior Parietal, PGi | Right | Default | -0.0243 |
| R_9p | Dorsolateral Prefrontal, 9p | Right | Default | -0.0243 |
| R_6a | Premotor, 6a | Right | Dorsal_Attention | 0.0241 |
| L_STSvp | Auditory Association, STSvp | Left | Default | 0.0239 |
| R_RSC | Posterior Cingulate, RSC | Right | Frontoparietal | 0.0238 |
| R_OP4 | Posterior Opercular, OP4 | Right | Somatomotor | -0.0237 |
| R_SFL | Dorsolateral Prefrontal, SFL | Right | Language | -0.0236 |
| R_11l | Orbital and Polar Frontal, 11l | Right | Frontoparietal | 0.0235 |
| L_PoI2 | Insular and Frontal Opercular, PoI2 | Left | Cingulo-Opercular | -0.0234 |
| L_7Am | Superior Parietal, 7Am | Left | Cingulo-Opercular | 0.0233 |
| L_VMV3 | Ventral Stream Visual, VMV3 | Left | Visual2 | -0.0233 |
| L_p10p | Orbital and Polar Frontal, p10p | Left | Frontoparietal | -0.0232 |
| L_TPOJ1 | Temporo-Parieto-Occipital Junction, TPOJ1 | Left | Language | 0.0232 |
| L_10d | Orbital and Polar Frontal, 10d | Left | Default | -0.0228 |
| L_Pir | Insular and Frontal Opercular, Pir | Left | Orbito-Affective | 0.0225 |
| R_TF | Medial Temporal, TF | Right | Ventral_Multimodal | 0.0224 |
| L_LBelt | Early Auditory, LBelt | Left | Auditory | 0.0224 |
| L_MST | MT+ Complex and Neighboring Visual Areas, MST | Left | Visual2 | -0.0223 |
| L_V4t | MT+ Complex and Neighboring Visual Areas, V4t | Left | Visual2 | 0.0219 |
| R_TPOJ3 | Temporo-Parieto-Occipital Junction, TPOJ3 | Right | Posterior_Multimodal | 0.0219 |
| L_STSdp | Auditory Association, STSdp | Left | Language | 0.0219 |
| R_6r | Premotor, 6r | Right | Cingulo-Opercular | 0.0218 |
| R_VMV3 | Ventral Stream Visual, VMV3 | Right | Visual2 | -0.0218 |
| R_p32pr | Anterior Cingulate and Medial Prefrontal, p32pr | Right | Cingulo-Opercular | -0.0217 |
| L_IFJp | Inferior Frontal, IFJp | Left | Frontoparietal | 0.0217 |
| R_IFSa | Inferior Frontal, IFSa | Right | Cingulo-Opercular | 0.0214 |
| R_v23ab | Posterior Cingulate, v23ab | Right | Default | -0.0214 |
| R_a9-46v | Dorsolateral Prefrontal, a9-46v | Right | Frontoparietal | 0.0213 |
| R_45 | Inferior Frontal, 45 | Right | Language | 0.0212 |
| R_6v | Premotor, 6v | Right | Somatomotor | -0.0212 |
| R_i6-8 | Dorsolateral Prefrontal, i6-8 | Right | Frontoparietal | 0.021 |
| L_a24 | Anterior Cingulate and Medial Prefrontal, a24 | Left | Default | 0.0208 |
| L_23d | Posterior Cingulate, 23d | Left | Default | 0.0207 |
| L_SCEF | Paracentral Lobular and Mid Cingulate, SCEF | Left | Cingulo-Opercular | 0.0207 |
| L_FOP1 | Posterior Opercular, FOP1 | Left | Cingulo-Opercular | -0.0206 |
| R_VVC | Ventral Stream Visual, VVC | Right | Visual2 | 0.0206 |
| L_VVC | Ventral Stream Visual, VVC | Left | Visual2 | 0.0206 |
| L_FEF | Premotor, FEF | Left | Cingulo-Opercular | 0.0205 |
| R_VIP | Superior Parietal, VIP | Right | Visual2 | 0.0204 |
| R_PIT | Ventral Stream Visual, PIT | Right | Visual2 | -0.0202 |
| R_8BL | Dorsolateral Prefrontal, 8BL | Right | Default | -0.0202 |
| L_PBelt | Early Auditory, PBelt | Left | Auditory | 0.0201 |
| R_MST | MT+ Complex and Neighboring Visual Areas, MST | Right | Visual2 | -0.0198 |
| R_TE1m | Lateral Temporal, TE1m | Right | Frontoparietal | -0.0197 |
| PALLIDUM_RIGHT | Pallidum Right | Right | Subcortex | 0.0194 |
| R_6d | Premotor, 6d | Right | Somatomotor | 0.0193 |
| R_PHA1 | Medial Temporal, PHA1 | Right | Default | 0.0193 |
| R_PGp | Inferior Parietal, PGp | Right | Dorsal_Attention | -0.0193 |
| R_s6-8 | Dorsolateral Prefrontal, s6-8 | Right | Frontoparietal | -0.0192 |
| R_p10p | Orbital and Polar Frontal, p10p | Right | Frontoparietal | -0.0192 |
| R_V6 | Dorsal Stream Visual, V6 | Right | Visual2 | -0.0191 |
| R_46 | Dorsolateral Prefrontal, 46 | Right | Cingulo-Opercular | 0.0189 |
| L_IP2 | Inferior Parietal, IP2 | Left | Frontoparietal | 0.0189 |
| R_MT | MT+ Complex and Neighboring Visual Areas, MT | Right | Visual2 | -0.0188 |
| CAUDATE_LEFT | Caudate Left | Left | Subcortex | 0.0187 |
| L_p24pr | Anterior Cingulate and Medial Prefrontal, p24pr | Left | Cingulo-Opercular | 0.0186 |
| L_52 | Early Auditory, 52 | Left | Auditory | -0.0185 |
| L_PGi | Inferior Parietal, PGi | Left | Default | -0.0184 |
| R_13l | Orbital and Polar Frontal, 13l | Right | Frontoparietal | -0.0184 |
| R_p32 | Anterior Cingulate and Medial Prefrontal, p32 | Right | Default | -0.0183 |
| R_pOFC | Anterior Cingulate and Medial Prefrontal, pOFC | Right | Orbito-Affective | 0.0182 |
| HIPPOCAMPUS_RIGHT | Hippocampus Right | Right | Subcortex | -0.0182 |
| R_AAIC | Insular and Frontal Opercular, AAIC | Right | Orbito-Affective | -0.0181 |
| PUTAMEN_LEFT | Putamen Left | Left | Subcortex | -0.018 |
| R_PSL | Temporo-Parieto Occipital Junction, PSL | Right | Cingulo-Opercular | -0.0178 |
| L_24dd | Paracentral Lobular and Mid Cingulate, 24dd | Left | Somatomotor | -0.0177 |
| R_PoI1 | Insular and Frontal Opercular, PoI1 | Right | Cingulo-Opercular | -0.0177 |
| PALLIDUM_LEFT | Pallidum Left | Left | Subcortex | -0.0177 |
| L_6a | Premotor, 6a | Left | Dorsal_Attention | -0.0175 |
| L_FFC | Ventral Stream Visual, FFC | Left | Visual2 | 0.0174 |
| R_IFJp | Inferior Frontal, IFJp | Right | Frontoparietal | 0.017 |
| L_p9-46v | Dorsolateral Prefrontal, p9-46v | Left | Frontoparietal | -0.017 |
| R_TA2 | Auditory Association, TA2 | Right | Auditory | -0.0168 |
| L_Ig | Insular and Frontal Opercular, Ig | Left | Somatomotor | 0.0167 |
| R_IFSp | Inferior Frontal, IFSp | Right | Frontoparietal | 0.0167 |
| CEREBELLUM_LEFT | Cerebellum Left | Left | Subcortex | 0.0166 |
| R_PFop | Inferior Parietal, PFop | Right | Cingulo-Opercular | -0.0165 |
| R_TE1p | Lateral Temporal, TE1p | Right | Frontoparietal | 0.0164 |
| L_9m | Anterior Cingulate and Medial Prefrontal, 9m | Left | Default | 0.0164 |
| R_PF | Inferior Parietal, PF | Right | Cingulo-Opercular | -0.0164 |
| R_8Ad | Dorsolateral Prefrontal, 8Ad | Right | Default | -0.0164 |
| L_V6 | Dorsal Stream Visual, V6 | Left | Visual2 | -0.0162 |
| R_31a | Posterior Cingulate, 31a | Right | Frontoparietal | -0.016 |
| L_OFC | Orbital and Polar Frontal, OFC | Left | Default | -0.016 |
| L_s6-8 | Dorsolateral Prefrontal, s6-8 | Left | Frontoparietal | -0.0159 |
| L_FOP2 | Insular and Frontal Opercular, FOP2 | Left | Somatomotor | -0.0158 |
| R_52 | Early Auditory, 52 | Right | Auditory | -0.0158 |
| R_EC | Medial Temporal, EC | Right | Default | 0.0156 |
| L_8BM | Anterior Cingulate and Medial Prefrontal, 8BM | Left | Frontoparietal | 0.0156 |
| R_STSva | Auditory Association, STSva | Right | Default | -0.0155 |
| R_TE2p | Lateral Temporal, TE2p | Right | Dorsal_Attention | 0.0153 |
| R_TE2a | Lateral Temporal, TE2a | Right | Default | -0.0153 |
| R_5m | Paracentral Lobular and Mid Cingulate, 5m | Right | Somatomotor | 0.0152 |
| L_47m | Orbital and Polar Frontal, 47m | Left | Default | 0.0151 |
| L_d32 | Anterior Cingulate and Medial Prefrontal, d32 | Left | Default | 0.0151 |
| R_IP2 | Inferior Parietal, IP2 | Right | Frontoparietal | 0.0149 |
| R_47m | Orbital and Polar Frontal, 47m | Right | Default | -0.0149 |
| R_V8 | Ventral Stream Visual, V8 | Right | Visual2 | -0.0148 |
| R_a10p | Orbital and Polar Frontal, a10p | Right | Frontoparietal | -0.0148 |
| R_STV | Temporo-Parieto-Occipital Junction, STV | Right | Posterior_Multimodal | -0.0147 |
| L_TGd | Lateral Temporal, TGd | Left | Default | -0.0146 |
| R_A5 | Auditory Association, A5 | Right | Language | -0.0145 |
| R_5L | Paracentral Lobular and Mid Cingulate, 5L | Right | Somatomotor | 0.0145 |
| R_TE1a | Lateral Temporal, TE1a | Right | Default | -0.0144 |
| L_8Ad | Dorsolateral Prefrontal, 8Ad | Left | Default | -0.0144 |
| L_7Pm | Superior Parietal, 7Pm | Left | Frontoparietal | -0.0143 |
| R_IP1 | Inferior Parietal, IP1 | Right | Frontoparietal | -0.0143 |
| R_10v | Anterior Cingulate and Medial Prefrontal, 10v | Right | Default | -0.0142 |
| R_p24pr | Anterior Cingulate and Medial Prefrontal, p24pr | Right | Cingulo-Opercular | -0.0142 |
| R_FEF | Premotor, FEF | Right | Cingulo-Opercular | 0.0141 |
| R_TGd | Lateral Temporal, TGd | Right | Default | -0.0141 |
| L_LO1 | MT+ Complex and Neighboring Visual Areas, LO1 | Left | Visual2 | 0.0141 |
| R_PFt | Inferior Parietal, PFt | Right | Dorsal_Attention | 0.0139 |
| L_PGs | Inferior Parietal, PGs | Left | Default | -0.0135 |
| L_MIP | Superior Parietal, MIP | Left | Dorsal_Attention | 0.0135 |
| L_V7 | Dorsal Stream Visual, V7 | Left | Visual2 | -0.0134 |
| R_55b | Premotor, 55b | Right | Language | -0.0133 |
| R_PHA3 | Medial Temporal, PHA3 | Right | Dorsal_Attention | -0.0133 |
| L_FOP4 | Insular and Frontal Opercular, FOP4 | Left | Cingulo-Opercular | 0.013 |
| R_LO2 | MT+ Complex and Neighboring Visual Areas, LO2 | Right | Visual2 | 0.0128 |
| L_V1 | Primary Visual, V1 | Left | Visual1 | -0.0128 |
| R_ProS | Posterior Cingulate, ProS | Right | Visual1 | 0.0127 |
| R_a24 | Anterior Cingulate and Medial Prefrontal, a24 | Right | Default | -0.0127 |
| L_LO2 | MT+ Complex and Neighboring Visual Areas, LO2 | Left | Visual2 | 0.0127 |
| L_IP1 | Inferior Parietal, IP1 | Left | Frontoparietal | -0.0125 |
| R_25 | Anterior Cingulate and Medial Prefrontal, 25 | Right | Default | -0.0124 |
| R_V3B | Dorsal Stream Visual, V3B | Right | Visual2 | -0.0123 |
| L_a9-46v | Dorsolateral Prefrontal, a9-46v | Left | Frontoparietal | -0.0123 |
| L_4 | Somatosensory and Motor, 4 | Left | Somatomotor | 0.0123 |
| L_9p | Dorsolateral Prefrontal, 9p | Left | Default | -0.0122 |
| L_IPS1 | Dorsal Stream Visual, IPS1 | Left | Visual2 | -0.0122 |
| R_3a | Somatosensory and Motor, 3a | Right | Somatomotor | -0.0121 |
| AMYGDALA_LEFT | Amygdala Left | Left | Subcortex | -0.012 |
| L_44 | Inferior Frontal, 44 | Left | Language | -0.0119 |
| R_7Am | Superior Parietal, 7Am | Right | Cingulo-Opercular | -0.0118 |
| L_PHA1 | Medial Temporal, PHA1 | Left | Default | 0.0117 |
| R_TGv | Lateral Temporal, TGv | Right | Language | -0.0117 |
| R_V4 | Early Visual, V4 | Right | Visual2 | 0.0116 |
| R_V3A | Dorsal Stream Visual, V3A | Right | Visual2 | -0.0116 |
| R_33pr | Anterior Cingulate and Medial Prefrontal, 33pr | Right | Frontoparietal | 0.0116 |
| R_OP1 | Posterior Opercular, OP1 | Right | Somatomotor | -0.0115 |
| R_POS1 | Posterior Cingulate, POS1 | Right | Default | 0.0113 |
| CAUDATE_RIGHT | Caudate Right | Right | Subcortex | -0.0111 |
| L_MT | MT+ Complex and Neighboring Visual Areas, MT | Left | Visual2 | 0.0111 |
| DIENCEPHALON_VENTRAL_RIGHT | Diencephalon Ventral Right | Right | Subcortex | 0.0109 |
| L_FOP3 | Insular and Frontal Opercular, FOP3 | Left | Cingulo-Opercular | 0.0109 |
| L_FST | MT+ Complex and Neighboring Visual Areas, FST | Left | Visual2 | 0.0109 |
| L_PH | MT+ Complex and Neighboring Visual Areas, PH | Left | Visual2 | 0.0109 |
| L_V3A | Dorsal Stream Visual, V3A | Left | Visual2 | 0.0108 |
| R_p24 | Anterior Cingulate and Medial Prefrontal, p24 | Right | Cingulo-Opercular | -0.0108 |
| L_8BL | Dorsolateral Prefrontal, 8BL | Left | Default | -0.0107 |
| CEREBELLUM_RIGHT | Cerebellum Right | Right | Subcortex | 0.0106 |
| L_9-46d | Dorsolateral Prefrontal, 9-46d | Left | Cingulo-Opercular | -0.0105 |
| R_7m | Posterior Cingulate, 7m | Right | Default | 0.0104 |
| L_VMV1 | Ventral Stream Visual, VMV1 | Left | Visual2 | -0.0104 |
| R_POS2 | Posterior Cingulate, POS2 | Right | Frontoparietal | 0.0103 |
| L_s32 | Anterior Cingulate and Medial Prefrontal, s32 | Left | Default | -0.0103 |
| L_55b | Premotor, 55b | Left | Language | 0.0102 |
| R_MBelt | Early Auditory, MBelt | Right | Auditory | 0.0102 |
| L_VMV2 | Ventral Stream Visual, VMV2 | Left | Visual2 | -0.0102 |
| L_TPOJ2 | Temporo-Parieto-Occipital Junction, TPOJ2 | Left | Posterior_Multimodal | -0.0102 |
| L_IFSa | Inferior Frontal, IFSa | Left | Frontoparietal | -0.0101 |
| L_TE1a | Lateral Temporal, TE1a | Left | Default | -0.0101 |
| R_Ig | Insular and Frontal Opercular, Ig | Right | Somatomotor | -0.0101 |
| L_RSC | Posterior Cingulate, RSC | Left | Frontoparietal | 0.0099 |
| L_PIT | Ventral Stream Visual, PIT | Left | Visual2 | -0.0098 |
| R_PEF | Premotor, PEF | Right | Cingulo-Opercular | 0.0098 |
| R_a32pr | Anterior Cingulate and Medial Prefrontal, a32pr | Right | Cingulo-Opercular | 0.0098 |
| L_V4 | Early Visual, V4 | Left | Visual2 | -0.0098 |
| R_8Av | Dorsolateral Prefrontal, 8Av | Right | Default | 0.0097 |
| R_31pd | Posterior Cingulate, 31pd | Right | Default | -0.0097 |
| L_33pr | Anterior Cingulate and Medial Prefrontal, 33pr | Left | Cingulo-Opercular | 0.0097 |
| R_AIP | Superior Parietal, AIP | Right | Dorsal_Attention | 0.0096 |
| R_MI | Insular and Frontal Opercular, MI | Right | Cingulo-Opercular | 0.0095 |
| R_PGs | Inferior Parietal, PGs | Right | Default | -0.0095 |
| R_1 | Somatosensory and Motor, 1 | Right | Somatomotor | 0.0095 |
| L_25 | Anterior Cingulate and Medial Prefrontal, 25 | Left | Default | -0.0094 |
| THALAMUS_LEFT | Thalamus Left | Left | Subcortex | -0.0093 |
| L_23c | Paracentral Lobular and Mid Cingulate, 23c | Left | Cingulo-Opercular | -0.0093 |
| L_PFt | Inferior Parietal, PFt | Left | Dorsal_Attention | -0.0091 |
| L_STGa | Auditory Association, STGa | Left | Language | -0.0091 |
| L_TE2a | Lateral Temporal, TE2a | Left | Default | 0.0089 |
| R_LIPv | Superior Parietal, LIPv | Right | Visual2 | 0.0088 |
| L_6mp | Paracentral Lobular and Mid Cingulate, 6mp | Left | Somatomotor | 0.0087 |
| R_OP2-3 | Posterior Opercular, OP2-3 | Right | Somatomotor | 0.0085 |
| L_pOFC | Anterior Cingulate and Medial Prefrontal, pOFC | Left | Orbito-Affective | -0.0084 |
| R_V2 | Early Visual, V2 | Right | Visual2 | -0.0083 |
| R_LIPd | Superior Parietal, LIPd | Right | Dorsal_Attention | 0.0082 |
| L_OP4 | Posterior Opercular, OP4 | Left | Somatomotor | -0.0082 |
| R_PeEc | Medial Temporal, PeEc | Right | Ventral_Multimodal | -0.008 |
| L_LIPv | Superior Parietal, LIPv | Left | Visual2 | -0.008 |
| R_24dd | Paracentral Lobular and Mid Cingulate, 24dd | Right | Somatomotor | 0.008 |
| R_LBelt | Early Auditory, LBelt | Right | Auditory | -0.008 |
| L_5mv | Paracentral Lobular and Mid Cingulate, 5mv | Left | Cingulo-Opercular | -0.008 |
| L_PreS | Medial Temporal, PreS | Left | Default | 0.0079 |
| L_PGp | Inferior Parietal, PGp | Left | Dorsal_Attention | -0.0077 |
| L_IFSp | Inferior Frontal, IFSp | Left | Language | -0.0077 |
| R_9m | Anterior Cingulate and Medial Prefrontal, 9m | Right | Default | -0.0077 |
| L_TPOJ3 | Temporo-Parieto-Occipital Junction, TPOJ3 | Left | Posterior_Multimodal | 0.0077 |
| R_PI | Insular and Frontal Opercular, PI | Right | Cingulo-Opercular | -0.0076 |
| R_10pp | Orbital and Polar Frontal, 10pp | Right | Default | -0.0076 |
| L_1 | Somatosensory and Motor, 1 | Left | Somatomotor | 0.0075 |
| L_SFL | Dorsolateral Prefrontal, SFL | Left | Language | -0.0075 |
| L_p24 | Anterior Cingulate and Medial Prefrontal, p24 | Left | Cingulo-Opercular | 0.0075 |
| L_STV | Temporo-Parieto-Occipital Junction, STV | Left | Language | -0.0074 |
| R_RI | Early Auditory, RI | Right | Somatomotor | -0.0073 |
| L_PHA2 | Medial Temporal, PHA2 | Left | Default | -0.0072 |
| R_V1 | Primary Visual, V1 | Right | Visual1 | 0.0071 |
| L_AAIC | Insular and Frontal Opercular, AAIC | Left | Orbito-Affective | 0.007 |
| L_TA2 | Auditory Association, TA2 | Left | Auditory | -0.0069 |
| R_5mv | Paracentral Lobular and Mid Cingulate, 5mv | Right | Cingulo-Opercular | -0.0069 |
| L_MI | Insular and Frontal Opercular, MI | Left | Cingulo-Opercular | -0.0069 |
| R_7AL | Superior Parietal, 7AL | Right | Somatomotor | 0.0068 |
| L_EC | Medial Temporal, EC | Left | Default | 0.0067 |
| L_DVT | Posterior Cingulate, DVT | Left | Visual1 | 0.0066 |
| R_6ma | Paracentral Lobular and Mid Cingulate, 6ma | Right | Cingulo-Opercular | -0.0066 |
| L_a10p | Orbital and Polar Frontal, a10p | Left | Frontoparietal | 0.0066 |
| L_MBelt | Early Auditory, MBelt | Left | Auditory | 0.0066 |
| L_OP2-3 | Posterior Opercular, OP2-3 | Left | Somatomotor | -0.0064 |
| ACCUMBENS_RIGHT | Accumbens Right | Right | Subcortex | -0.0064 |
| L_47l | Inferior Frontal, 47l | Left | Default | 0.0063 |
| L_ProS | Posterior Cingulate, ProS | Left | Visual1 | 0.0063 |
| THALAMUS_RIGHT | Thalamus Right | Right | Subcortex | 0.0063 |
| L_V6A | Dorsal Stream Visual, V6A | Left | Visual2 | 0.0062 |
| R_DVT | Posterior Cingulate, DVT | Right | Visual1 | 0.0061 |
| R_FOP5 | Insular and Frontal Opercular, FOP5 | Right | Cingulo-Opercular | 0.0061 |
| L_10pp | Orbital and Polar Frontal, 10pp | Left | Default | 0.006 |
| L_6r | Premotor, 6r | Left | Cingulo-Opercular | 0.006 |
| R_47l | Inferior Frontal, 47l | Right | Default | 0.0059 |
| L_13l | Orbital and Polar Frontal, 13l | Left | Frontoparietal | 0.0059 |
| R_47s | Orbital and Polar Frontal, 47s | Right | Default | 0.0059 |
| R_FOP4 | Insular and Frontal Opercular, FOP4 | Right | Cingulo-Opercular | 0.0058 |
| R_PreS | Medial Temporal, PreS | Right | Default | 0.0056 |
| PUTAMEN_RIGHT | Putamen Right | Right | Subcortex | -0.0055 |
| L_TGv | Lateral Temporal, TGv | Left | Language | -0.0054 |
| L_31pv | Posterior Cingulate, 31pv | Left | Default | -0.0054 |
| R_FOP2 | Insular and Frontal Opercular, FOP2 | Right | Somatomotor | 0.0053 |
| R_d23ab | Posterior Cingulate, d23ab | Right | Default | -0.0052 |
| R_V3CD | MT+ Complex and Neighboring Visual Areas, V3CD | Right | Visual2 | 0.0052 |
| R_VMV1 | Ventral Stream Visual, VMV1 | Right | Visual2 | 0.0051 |
| L_47s | Orbital and Polar Frontal, 47s | Left | Default | 0.005 |
| L_p47r | Inferior Frontal, p47r | Left | Frontoparietal | -0.005 |
| R_V6A | Dorsal Stream Visual, V6A | Right | Visual2 | -0.0049 |
| R_10r | Anterior Cingulate and Medial Prefrontal, 10r | Right | Default | 0.0049 |
| R_44 | Inferior Frontal, 44 | Right | Frontoparietal | -0.0048 |
| L_p32 | Anterior Cingulate and Medial Prefrontal, p32 | Left | Default | 0.0048 |
| R_a24pr | Anterior Cingulate and Medial Prefrontal, a24pr | Right | Cingulo-Opercular | 0.0048 |
| R_43 | Posterior Opercular, 43 | Right | Cingulo-Opercular | -0.0046 |
| L_VIP | Superior Parietal, VIP | Left | Visual2 | 0.0046 |
| L_24dv | Paracentral Lobular and Mid Cingulate, 24dv | Left | Somatomotor | -0.0045 |
| L_11l | Orbital and Polar Frontal, 11l | Left | Frontoparietal | 0.0043 |
| L_TE1p | Lateral Temporal, TE1p | Left | Frontoparietal | -0.0042 |
| L_2 | Somatosensory and Motor, 2 | Left | Somatomotor | -0.0042 |
| R_31pv | Posterior Cingulate, 31pv | Right | Default | 0.0041 |
| L_PFcm | Early Auditory, PFcm | Left | Cingulo-Opercular | -0.0041 |
| L_AIP | Superior Parietal, AIP | Left | Dorsal_Attention | 0.004 |
| R_FOP3 | Insular and Frontal Opercular, FOP3 | Right | Cingulo-Opercular | -0.004 |
| R_p9-46v | Dorsolateral Prefrontal, p9-46v | Right | Frontoparietal | -0.0039 |
| L_PF | Inferior Parietal, PF | Left | Cingulo-Opercular | 0.0039 |
| L_7PC | Superior Parietal, 7PC | Left | Somatomotor | -0.0039 |
| R_10d | Orbital and Polar Frontal, 10d | Right | Default | -0.0038 |
| R_3b | Somatosensory and Motor, 3b | Right | Somatomotor | -0.0037 |
| L_3a | Somatosensory and Motor, 3a | Left | Somatomotor | -0.0037 |
| R_7Pm | Superior Parietal, 7Pm | Right | Frontoparietal | -0.0036 |
| R_SCEF | Paracentral Lobular and Mid Cingulate, SCEF | Right | Cingulo-Opercular | -0.0036 |
| R_LO3 | MT+ Complex and Neighboring Visual Areas, LO3 | Right | Visual2 | 0.0036 |
| L_i6-8 | Dorsolateral Prefrontal, i6-8 | Left | Frontoparietal | 0.0036 |
| R_PHA2 | Medial Temporal, PHA2 | Right | Default | 0.0036 |
| R_9-46d | Dorsolateral Prefrontal, 9-46d | Right | Cingulo-Opercular | -0.0034 |
| L_PEF | Premotor, PEF | Left | Dorsal_Attention | -0.0034 |
| L_V3 | Early Visual, V3 | Left | Visual2 | -0.0033 |
| AMYGDALA_RIGHT | Amygdala Right | Right | Subcortex | 0.0032 |
| R_FOP1 | Posterior Opercular, FOP1 | Right | Cingulo-Opercular | -0.0031 |
| L_TE2p | Lateral Temporal, TE2p | Left | Dorsal_Attention | 0.0031 |
| L_3b | Somatosensory and Motor, 3b | Left | Somatomotor | 0.003 |
| L_LIPd | Superior Parietal, LIPd | Left | Dorsal_Attention | -0.003 |
| R_TPOJ2 | Temporo-Parieto-Occipital Junction, TPOJ2 | Right | Posterior_Multimodal | -0.003 |
| BRAIN_STEM | Brain Stem | None | Subcortex | 0.003 |
| R_24dv | Paracentral Lobular and Mid Cingulate, 24dv | Right | Somatomotor | 0.0029 |
| L_7PL | Superior Parietal, 7Pl | Left | Dorsal_Attention | 0.0028 |
| L_TF | Medial Temporal, TF | Left | Ventral_Multimodal | -0.0028 |
| L_FOP5 | Insular and Frontal Opercular, FOP5 | Left | Cingulo-Opercular | 0.0028 |
| L_H | Medial Temporal, H | Left | Default | -0.0027 |
| L_a24pr | Anterior Cingulate and Medial Prefrontal, a24pr | Left | Cingulo-Opercular | 0.0027 |
| R_H | Medial Temporal, H | Right | Default | 0.0026 |
| L_45 | Inferior Frontal, 45 | Left | Language | -0.0026 |
| R_Pir | Insular and Frontal Opercular, Pir | Right | Orbito-Affective | 0.0026 |
| R_23c | Paracentral Lobular and Mid Cingulate, 23c | Right | Cingulo-Opercular | -0.0025 |
| R_6mp | Paracentral Lobular and Mid Cingulate, 6mp | Right | Somatomotor | -0.0024 |
| L_STSva | Auditory Association, STSva | Left | Default | -0.0023 |
| L_8Av | Dorsolateral Prefrontal, 8Av | Left | Default | -0.0023 |
| L_PHT | Lateral Temporal, PHT | Left | Dorsal_Attention | 0.0021 |
| R_PoI2 | Insular and Frontal Opercular, PoI2 | Right | Cingulo-Opercular | 0.002 |
| L_POS2 | Posterior Cingulate, POS2 | Left | Frontoparietal | 0.002 |
| R_IP0 | Inferior Parietal, IP0 | Right | Dorsal_Attention | 0.002 |
| L_p32pr | Anterior Cingulate and Medial Prefrontal, p32pr | Left | Cingulo-Opercular | -0.0018 |
| R_STSda | Auditory Association, STSda | Right | Default | 0.0018 |
| L_V3CD | MT+ Complex and Neighboring Visual Areas, V3CD | Left | Visual2 | 0.0018 |
| L_A4 | Auditory Association, A4 | Left | Auditory | 0.0018 |
| R_2 | Somatosensory and Motor, 2 | Right | Somatomotor | -0.0018 |
| L_46 | Dorsolateral Prefrontal, 46 | Left | Cingulo-Opercular | 0.0017 |
| L_7m | Posterior Cingulate, 7m | Left | Default | -0.0016 |
| L_PeEc | Medial Temporal, PeEc | Left | Ventral_Multimodal | -0.0016 |
| L_10v | Anterior Cingulate and Medial Prefrontal, 10v | Left | Default | -0.0014 |
| R_9a | Dorsolateral Prefrontal, 9a | Right | Default | -0.0014 |
| R_OFC | Orbital and Polar Frontal, OFC | Right | Frontoparietal | 0.0013 |
| L_PHA3 | Medial Temporal, PHA3 | Left | Dorsal_Attention | 0.0013 |
| L_d23ab | Posterior Cingulate, d23ab | Left | Default | -0.0012 |
| R_s32 | Anterior Cingulate and Medial Prefrontal, s32 | Right | Default | 0.0012 |
| L_5m | Paracentral Lobular and Mid Cingulate, 5m | Left | Somatomotor | -0.0012 |
| R_d32 | Anterior Cingulate and Medial Prefrontal, d32 | Right | Frontoparietal | -0.0012 |
| L_5L | Paracentral Lobular and Mid Cingulate, 5L | Left | Somatomotor | -0.0012 |
| L_a47r | Inferior Frontal, a47r | Left | Frontoparietal | -0.0011 |
| R_7PC | Superior Parietal, 7PC | Right | Somatomotor | 0.0008 |
| R_PFcm | Early Auditory, PFcm | Right | Cingulo-Opercular | -0.0007 |
| L_9a | Dorsolateral Prefrontal, 9a | Left | Default | 0.0006 |
| R_23d | Posterior Cingulate, 23d | Right | Default | 0.0006 |
| L_31pd | Posterior Cingulate, 31pd | Left | Default | -0.0005 |
| R_TPOJ1 | Temporo-Parieto-Occipital Junction, TPOJ1 | Right | Language | 0.0005 |
| R_PBelt | Early Auditory, PBelt | Right | Auditory | -0.0005 |
| R_p47r | Inferior Frontal, p47r | Right | Frontoparietal | 0.0005 |
| R_V3 | Early Visual, V3 | Right | Visual2 | -0.0005 |
| L_PI | Insular and Frontal Opercular, PI | Left | Cingulo-Opercular | -0.0005 |
| R_a47r | Inferior Frontal, a47r | Right | Frontoparietal | 0.0004 |
| L_10r | Anterior Cingulate and Medial Prefrontal, 10r | Left | Default | -0.0003 |
| R_PHT | Lateral Temporal, PHT | Right | Dorsal_Attention | -0.0003 |
| L_PFop | Inferior Parietal, PFop | Left | Cingulo-Opercular | -0.0003 |
| L_POS1 | Posterior Cingulate, POS1 | Left | Default | 0.0002 |
| L_STSda | Auditory Association, STSda | Left | Language | 0.0002 |
| R_A1 | Early Auditory, A1 | Right | Auditory | 0.0002 |
| R_V4t | MT+ Complex and Neighboring Visual Areas, V4t | Right | Visual2 | -0.0001 |
| L_6ma | Paracentral Lobular and Mid Cingulate, 6ma | Left | Cingulo-Opercular | 0.0001 |
| L_TE1m | Lateral Temporal, TE1m | Left | Default | 0 |
| R_4 | Somatosensory and Motor, 4 | Right | Somatomotor | 0 |

***Table S5. Feature importance of the top-performing non-stacked models with with Elastic Net, as indicated by Elastic Net coefficients for the Dunedin Study dataset, predicting residual scores for cognitive abilities: Facename Encoding vs Distractor Task Contrast***

| **Glasser's label** | **Brain Region** | **Hemisphere** | **Network** | **Amplitude** |
| --- | --- | --- | --- | --- |
| R_LIPd | Superior Parietal, LIPd | Right | Dorsal_Attention | 0.0734 |
| L_LIPd | Superior Parietal, LIPd | Left | Dorsal_Attention | 0.0535 |
| CEREBELLUM_LEFT | Cerebellum Left | Left | Subcortex | 0.0429 |
| L_IP2 | Inferior Parietal, IP2 | Left | Frontoparietal | 0.0362 |
| L_V3B | Dorsal Stream Visual, V3B | Left | Visual2 | -0.0259 |
| R_4 | Somatosensory and Motor, 4 | Right | Somatomotor | -0.0228 |
| HIPPOCAMPUS_RIGHT | Hippocampus Right | Right | Subcortex | -0.0226 |
| L_AIP | Superior Parietal, AIP | Left | Dorsal_Attention | 0.0213 |
| L_POS1 | Posterior Cingulate, POS1 | Left | Default | -0.0212 |
| L_V4t | MT+ Complex and Neighboring Visual Areas, V4t | Left | Visual2 | -0.0209 |
| R_TF | Medial Temporal, TF | Right | Ventral_Multimodal | -0.0208 |
| L_3b | Somatosensory and Motor, 3b | Left | Somatomotor | -0.0187 |
| L_TF | Medial Temporal, TF | Left | Ventral_Multimodal | -0.0184 |
| THALAMUS_LEFT | Thalamus Left | Left | Subcortex | 0.0148 |
| R_AIP | Superior Parietal, AIP | Right | Dorsal_Attention | 0.0136 |
| R_3a | Somatosensory and Motor, 3a | Right | Somatomotor | -0.0114 |
| HIPPOCAMPUS_LEFT | Hippocampus Left | Left | Subcortex | -0.011 |
| L_5L | Paracentral Lobular and Mid Cingulate, 5L | Left | Somatomotor | -0.008 |
| L_TE2a | Lateral Temporal, TE2a | Left | Default | -0.0078 |
| R_IFJp | Inferior Frontal, IFJp | Right | Frontoparietal | 0.0075 |
| L_TE1m | Lateral Temporal, TE1m | Left | Default | -0.0059 |
| L_5m | Paracentral Lobular and Mid Cingulate, 5m | Left | Somatomotor | -0.003 |
| L_VMV2 | Ventral Stream Visual, VMV2 | Left | Visual2 | -0.0026 |
| L_IPS1 | Dorsal Stream Visual, IPS1 | Left | Visual2 | 0.0023 |
| L_OP2-3 | Posterior Opercular, OP2-3 | Left | Somatomotor | -0.0021 |
| R_p9-46v | Dorsolateral Prefrontal, p9-46v | Right | Frontoparietal | 0.0004 |
| L_44 | Inferior Frontal, 44 | Left | Language | 0 |
| L_a47r | Inferior Frontal, a47r | Left | Frontoparietal | 0 |
| L_47l | Inferior Frontal, 47l | Left | Default | 0 |
| L_45 | Inferior Frontal, 45 | Left | Language | 0 |
| L_9m | Anterior Cingulate and Medial Prefrontal, 9m | Left | Default | 0 |
| L_8BL | Dorsolateral Prefrontal, 8BL | Left | Default | 0 |
| L_10d | Orbital and Polar Frontal, 10d | Left | Default | 0 |
| L_9p | Dorsolateral Prefrontal, 9p | Left | Default | 0 |
| L_47m | Orbital and Polar Frontal, 47m | Left | Default | 0 |
| L_8Av | Dorsolateral Prefrontal, 8Av | Left | Default | 0 |
| L_6r | Premotor, 6r | Left | Cingulo-Opercular | 0 |
| L_8Ad | Dorsolateral Prefrontal, 8Ad | Left | Default | 0 |
| L_8C | Dorsolateral Prefrontal, 8C | Left | Frontoparietal | 0 |
| L_a9-46v | Dorsolateral Prefrontal, a9-46v | Left | Frontoparietal | 0 |
| L_IFJa | Inferior Frontal, IFJa | Left | Language | 0 |
| L_IFJp | Inferior Frontal, IFJp | Left | Frontoparietal | 0 |
| L_s6-8 | Dorsolateral Prefrontal, s6-8 | Left | Frontoparietal | 0 |
| L_i6-8 | Dorsolateral Prefrontal, i6-8 | Left | Frontoparietal | 0 |
| L_6a | Premotor, 6a | Left | Dorsal_Attention | 0 |
| L_47s | Orbital and Polar Frontal, 47s | Left | Default | 0 |
| L_OFC | Orbital and Polar Frontal, OFC | Left | Default | 0 |
| L_13l | Orbital and Polar Frontal, 13l | Left | Frontoparietal | 0 |
| L_11l | Orbital and Polar Frontal, 11l | Left | Frontoparietal | 0 |
| L_10pp | Orbital and Polar Frontal, 10pp | Left | Default | 0 |
| L_a10p | Orbital and Polar Frontal, a10p | Left | Frontoparietal | 0 |
| L_10v | Anterior Cingulate and Medial Prefrontal, 10v | Left | Default | 0 |
| L_9a | Dorsolateral Prefrontal, 9a | Left | Default | 0 |
| L_9-46d | Dorsolateral Prefrontal, 9-46d | Left | Cingulo-Opercular | 0 |
| L_p32 | Anterior Cingulate and Medial Prefrontal, p32 | Left | Default | 0 |
| L_46 | Dorsolateral Prefrontal, 46 | Left | Cingulo-Opercular | 0 |
| L_p9-46v | Dorsolateral Prefrontal, p9-46v | Left | Frontoparietal | 0 |
| L_IFSa | Inferior Frontal, IFSa | Left | Frontoparietal | 0 |
| L_IFSp | Inferior Frontal, IFSp | Left | Language | 0 |
| L_10r | Anterior Cingulate and Medial Prefrontal, 10r | Left | Default | 0 |
| R_V1 | Primary Visual, V1 | Right | Visual1 | 0 |
| L_8BM | Anterior Cingulate and Medial Prefrontal, 8BM | Left | Frontoparietal | 0 |
| L_d32 | Anterior Cingulate and Medial Prefrontal, d32 | Left | Default | 0 |
| L_24dd | Paracentral Lobular and Mid Cingulate, 24dd | Left | Somatomotor | 0 |
| L_23c | Paracentral Lobular and Mid Cingulate, 23c | Left | Cingulo-Opercular | 0 |
| L_5mv | Paracentral Lobular and Mid Cingulate, 5mv | Left | Cingulo-Opercular | 0 |
| L_31pv | Posterior Cingulate, 31pv | Left | Default | 0 |
| L_d23ab | Posterior Cingulate, d23ab | Left | Default | 0 |
| L_v23ab | Posterior Cingulate, v23ab | Left | Default | 0 |
| L_23d | Posterior Cingulate, 23d | Left | Default | 0 |
| L_7m | Posterior Cingulate, 7m | Left | Default | 0 |
| L_7Pm | Superior Parietal, 7Pm | Left | Frontoparietal | 0 |
| L_STV | Temporo-Parieto-Occipital Junction, STV | Left | Language | 0 |
| L_PCV | Posterior Cingulate, PCV | Left | Posterior_Multimodal | 0 |
| L_SFL | Dorsolateral Prefrontal, SFL | Left | Language | 0 |
| L_PSL | Temporo-Parieto Occipital Junction, PSL | Left | Language | 0 |
| L_A1 | Early Auditory, A1 | Left | Auditory | 0 |
| L_MT | MT+ Complex and Neighboring Visual Areas, MT | Left | Visual2 | 0 |
| L_PIT | Ventral Stream Visual, PIT | Left | Visual2 | 0 |
| L_LO2 | MT+ Complex and Neighboring Visual Areas, LO2 | Left | Visual2 | 0 |
| L_LO1 | MT+ Complex and Neighboring Visual Areas, LO1 | Left | Visual2 | 0 |
| L_FFC | Ventral Stream Visual, FFC | Left | Visual2 | 0 |
| L_24dv | Paracentral Lobular and Mid Cingulate, 24dv | Left | Somatomotor | 0 |
| L_7AL | Superior Parietal, 7AL | Left | Somatomotor | 0 |
| L_SCEF | Paracentral Lobular and Mid Cingulate, SCEF | Left | Cingulo-Opercular | 0 |
| L_6d | Premotor, 6d | Left | Somatomotor | 0 |
| L_a24 | Anterior Cingulate and Medial Prefrontal, a24 | Left | Default | 0 |
| L_p32pr | Anterior Cingulate and Medial Prefrontal, p32pr | Left | Cingulo-Opercular | 0 |
| L_a24pr | Anterior Cingulate and Medial Prefrontal, a24pr | Left | Cingulo-Opercular | 0 |
| L_33pr | Anterior Cingulate and Medial Prefrontal, 33pr | Left | Cingulo-Opercular | 0 |
| L_OP4 | Posterior Opercular, OP4 | Left | Somatomotor | 0 |
| L_p24pr | Anterior Cingulate and Medial Prefrontal, p24pr | Left | Cingulo-Opercular | 0 |
| L_6v | Premotor, 6v | Left | Somatomotor | 0 |
| L_6mp | Paracentral Lobular and Mid Cingulate, 6mp | Left | Somatomotor | 0 |
| L_3a | Somatosensory and Motor, 3a | Left | Somatomotor | 0 |
| L_6ma | Paracentral Lobular and Mid Cingulate, 6ma | Left | Cingulo-Opercular | 0 |
| L_2 | Somatosensory and Motor, 2 | Left | Somatomotor | 0 |
| L_1 | Somatosensory and Motor, 1 | Left | Somatomotor | 0 |
| L_MIP | Superior Parietal, MIP | Left | Dorsal_Attention | 0 |
| L_VIP | Superior Parietal, VIP | Left | Visual2 | 0 |
| L_LIPv | Superior Parietal, LIPv | Left | Visual2 | 0 |
| L_7PC | Superior Parietal, 7PC | Left | Somatomotor | 0 |
| L_7PL | Superior Parietal, 7Pl | Left | Dorsal_Attention | 0 |
| L_7Am | Superior Parietal, 7Am | Left | Cingulo-Opercular | 0 |
| L_43 | Posterior Opercular, 43 | Left | Cingulo-Opercular | 0 |
| L_MI | Insular and Frontal Opercular, MI | Left | Cingulo-Opercular | 0 |
| L_OP1 | Posterior Opercular, OP1 | Left | Somatomotor | 0 |
| L_VVC | Ventral Stream Visual, VVC | Left | Visual2 | 0 |
| L_p47r | Inferior Frontal, p47r | Left | Frontoparietal | 0 |
| L_p10p | Orbital and Polar Frontal, p10p | Left | Frontoparietal | 0 |
| L_FOP5 | Insular and Frontal Opercular, FOP5 | Left | Cingulo-Opercular | 0 |
| L_Ig | Insular and Frontal Opercular, Ig | Left | Somatomotor | 0 |
| L_PoI1 | Insular and Frontal Opercular, PoI1 | Left | Cingulo-Opercular | 0 |
| L_pOFC | Anterior Cingulate and Medial Prefrontal, pOFC | Left | Orbito-Affective | 0 |
| L_s32 | Anterior Cingulate and Medial Prefrontal, s32 | Left | Default | 0 |
| L_25 | Anterior Cingulate and Medial Prefrontal, 25 | Left | Default | 0 |
| L_31a | Posterior Cingulate, 31a | Left | Default | 0 |
| L_MBelt | Early Auditory, MBelt | Left | Auditory | 0 |
| L_31pd | Posterior Cingulate, 31pd | Left | Default | 0 |
| L_LO3 | MT+ Complex and Neighboring Visual Areas, LO3 | Left | Visual2 | 0 |
| L_V3CD | MT+ Complex and Neighboring Visual Areas, V3CD | Left | Visual2 | 0 |
| L_FST | MT+ Complex and Neighboring Visual Areas, FST | Left | Visual2 | 0 |
| L_PHA2 | Medial Temporal, PHA2 | Left | Default | 0 |
| L_VMV3 | Ventral Stream Visual, VMV3 | Left | Visual2 | 0 |
| L_VMV1 | Ventral Stream Visual, VMV1 | Left | Visual2 | 0 |
| L_V6A | Dorsal Stream Visual, V6A | Left | Visual2 | 0 |
| L_TGv | Lateral Temporal, TGv | Left | Language | 0 |
| L_LBelt | Early Auditory, LBelt | Left | Auditory | 0 |
| L_PGi | Inferior Parietal, PGi | Left | Default | 0 |
| CAUDATE_LEFT | Caudate Left | Left | Subcortex | 0 |
| PUTAMEN_RIGHT | Putamen Right | Right | Subcortex | 0 |
| PUTAMEN_LEFT | Putamen Left | Left | Subcortex | 0 |
| PALLIDUM_RIGHT | Pallidum Right | Right | Subcortex | 0 |
| PALLIDUM_LEFT | Pallidum Left | Left | Subcortex | 0 |
| DIENCEPHALON_VENTRAL_RIGHT | Diencephalon Ventral Right | Right | Subcortex | 0 |
| DIENCEPHALON_VENTRAL_LEFT | Diencephalon Ventral Left | Left | Subcortex | 0 |
| CEREBELLUM_RIGHT | Cerebellum Right | Right | Subcortex | 0 |
| CAUDATE_RIGHT | Caudate Right | Right | Subcortex | 0 |
| BRAIN_STEM | Brain Stem | None | Subcortex | 0 |
| L_A4 | Auditory Association, A4 | Left | Auditory | 0 |
| AMYGDALA_RIGHT | Amygdala Right | Right | Subcortex | 0 |
| AMYGDALA_LEFT | Amygdala Left | Left | Subcortex | 0 |
| ACCUMBENS_RIGHT | Accumbens Right | Right | Subcortex | 0 |
| ACCUMBENS_LEFT | Accumbens Left | Left | Subcortex | 0 |
| L_p24 | Anterior Cingulate and Medial Prefrontal, p24 | Left | Cingulo-Opercular | 0 |
| L_a32pr | Anterior Cingulate and Medial Prefrontal, a32pr | Left | Cingulo-Opercular | 0 |
| L_PI | Insular and Frontal Opercular, PI | Left | Cingulo-Opercular | 0 |
| L_STSva | Auditory Association, STSva | Left | Default | 0 |
| L_PGs | Inferior Parietal, PGs | Left | Default | 0 |
| L_PFm | Inferior Parietal, PFm | Left | Frontoparietal | 0 |
| L_52 | Early Auditory, 52 | Left | Auditory | 0 |
| L_FOP1 | Posterior Opercular, FOP1 | Left | Cingulo-Opercular | 0 |
| L_PeEc | Medial Temporal, PeEc | Left | Ventral_Multimodal | 0 |
| L_ProS | Posterior Cingulate, ProS | Left | Visual1 | 0 |
| L_H | Medial Temporal, H | Left | Default | 0 |
| L_PreS | Medial Temporal, PreS | Left | Default | 0 |
| L_EC | Medial Temporal, EC | Left | Default | 0 |
| L_PFt | Inferior Parietal, PFt | Left | Dorsal_Attention | 0 |
| L_FOP2 | Insular and Frontal Opercular, FOP2 | Left | Somatomotor | 0 |
| L_FOP3 | Insular and Frontal Opercular, FOP3 | Left | Cingulo-Opercular | 0 |
| L_AAIC | Insular and Frontal Opercular, AAIC | Left | Orbito-Affective | 0 |
| L_PBelt | Early Auditory, PBelt | Left | Auditory | 0 |
| L_AVI | Insular and Frontal Opercular, AVI | Left | Frontoparietal | 0 |
| L_Pir | Insular and Frontal Opercular, Pir | Left | Orbito-Affective | 0 |
| L_POS2 | Posterior Cingulate, POS2 | Left | Frontoparietal | 0 |
| L_FOP4 | Insular and Frontal Opercular, FOP4 | Left | Cingulo-Opercular | 0 |
| L_TA2 | Auditory Association, TA2 | Left | Auditory | 0 |
| L_PoI2 | Insular and Frontal Opercular, PoI2 | Left | Cingulo-Opercular | 0 |
| L_PFcm | Early Auditory, PFcm | Left | Cingulo-Opercular | 0 |
| L_RI | Early Auditory, RI | Left | Auditory | 0 |
| L_STGa | Auditory Association, STGa | Left | Language | 0 |
| L_A5 | Auditory Association, A5 | Left | Language | 0 |
| L_PF | Inferior Parietal, PF | Left | Cingulo-Opercular | 0 |
| L_PH | MT+ Complex and Neighboring Visual Areas, PH | Left | Visual2 | 0 |
| L_PFop | Inferior Parietal, PFop | Left | Cingulo-Opercular | 0 |
| L_IP0 | Inferior Parietal, IP0 | Left | Dorsal_Attention | 0 |
| L_IP1 | Inferior Parietal, IP1 | Left | Frontoparietal | 0 |
| L_PGp | Inferior Parietal, PGp | Left | Dorsal_Attention | 0 |
| L_DVT | Posterior Cingulate, DVT | Left | Visual1 | 0 |
| L_TPOJ3 | Temporo-Parieto-Occipital Junction, TPOJ3 | Left | Posterior_Multimodal | 0 |
| L_TPOJ2 | Temporo-Parieto-Occipital Junction, TPOJ2 | Left | Posterior_Multimodal | 0 |
| L_TPOJ1 | Temporo-Parieto-Occipital Junction, TPOJ1 | Left | Language | 0 |
| L_PHT | Lateral Temporal, PHT | Left | Dorsal_Attention | 0 |
| L_PHA1 | Medial Temporal, PHA1 | Left | Default | 0 |
| L_TE2p | Lateral Temporal, TE2p | Left | Dorsal_Attention | 0 |
| L_TE1p | Lateral Temporal, TE1p | Left | Frontoparietal | 0 |
| L_TE1a | Lateral Temporal, TE1a | Left | Default | 0 |
| L_TGd | Lateral Temporal, TGd | Left | Default | 0 |
| L_STSvp | Auditory Association, STSvp | Left | Default | 0 |
| L_STSdp | Auditory Association, STSdp | Left | Language | 0 |
| L_STSda | Auditory Association, STSda | Left | Language | 0 |
| L_PHA3 | Medial Temporal, PHA3 | Left | Dorsal_Attention | 0 |
| L_V7 | Dorsal Stream Visual, V7 | Left | Visual2 | 0 |
| L_FEF | Premotor, FEF | Left | Cingulo-Opercular | 0 |
| L_RSC | Posterior Cingulate, RSC | Left | Frontoparietal | 0 |
| R_a24 | Anterior Cingulate and Medial Prefrontal, a24 | Right | Default | 0 |
| R_8BL | Dorsolateral Prefrontal, 8BL | Right | Default | 0 |
| R_9m | Anterior Cingulate and Medial Prefrontal, 9m | Right | Default | 0 |
| R_8Ad | Dorsolateral Prefrontal, 8Ad | Right | Default | 0 |
| R_8Av | Dorsolateral Prefrontal, 8Av | Right | Default | 0 |
| R_47m | Orbital and Polar Frontal, 47m | Right | Default | 0 |
| R_10r | Anterior Cingulate and Medial Prefrontal, 10r | Right | Default | 0 |
| R_p32 | Anterior Cingulate and Medial Prefrontal, p32 | Right | Default | 0 |
| R_8BM | Anterior Cingulate and Medial Prefrontal, 8BM | Right | Frontoparietal | 0 |
| R_d32 | Anterior Cingulate and Medial Prefrontal, d32 | Right | Frontoparietal | 0 |
| R_p32pr | Anterior Cingulate and Medial Prefrontal, p32pr | Right | Cingulo-Opercular | 0 |
| R_s6-8 | Dorsolateral Prefrontal, s6-8 | Right | Frontoparietal | 0 |
| R_a24pr | Anterior Cingulate and Medial Prefrontal, a24pr | Right | Cingulo-Opercular | 0 |
| R_33pr | Anterior Cingulate and Medial Prefrontal, 33pr | Right | Frontoparietal | 0 |
| R_p24pr | Anterior Cingulate and Medial Prefrontal, p24pr | Right | Cingulo-Opercular | 0 |
| R_6v | Premotor, 6v | Right | Somatomotor | 0 |
| R_6mp | Paracentral Lobular and Mid Cingulate, 6mp | Right | Somatomotor | 0 |
| R_6d | Premotor, 6d | Right | Somatomotor | 0 |
| R_2 | Somatosensory and Motor, 2 | Right | Somatomotor | 0 |
| R_1 | Somatosensory and Motor, 1 | Right | Somatomotor | 0 |
| R_MIP | Superior Parietal, MIP | Right | Dorsal_Attention | 0 |
| R_9p | Dorsolateral Prefrontal, 9p | Right | Default | 0 |
| R_10d | Orbital and Polar Frontal, 10d | Right | Default | 0 |
| R_8C | Dorsolateral Prefrontal, 8C | Right | Frontoparietal | 0 |
| R_44 | Inferior Frontal, 44 | Right | Frontoparietal | 0 |
| R_6a | Premotor, 6a | Right | Dorsal_Attention | 0 |
| R_47s | Orbital and Polar Frontal, 47s | Right | Default | 0 |
| R_OFC | Orbital and Polar Frontal, OFC | Right | Frontoparietal | 0 |
| R_13l | Orbital and Polar Frontal, 13l | Right | Frontoparietal | 0 |
| R_11l | Orbital and Polar Frontal, 11l | Right | Frontoparietal | 0 |
| R_10pp | Orbital and Polar Frontal, 10pp | Right | Default | 0 |
| R_a10p | Orbital and Polar Frontal, a10p | Right | Frontoparietal | 0 |
| R_10v | Anterior Cingulate and Medial Prefrontal, 10v | Right | Default | 0 |
| R_9a | Dorsolateral Prefrontal, 9a | Right | Default | 0 |
| R_9-46d | Dorsolateral Prefrontal, 9-46d | Right | Cingulo-Opercular | 0 |
| R_a9-46v | Dorsolateral Prefrontal, a9-46v | Right | Frontoparietal | 0 |
| R_46 | Dorsolateral Prefrontal, 46 | Right | Cingulo-Opercular | 0 |
| R_IFSa | Inferior Frontal, IFSa | Right | Cingulo-Opercular | 0 |
| R_IFSp | Inferior Frontal, IFSp | Right | Frontoparietal | 0 |
| R_IFJa | Inferior Frontal, IFJa | Right | Language | 0 |
| R_6r | Premotor, 6r | Right | Cingulo-Opercular | 0 |
| R_a47r | Inferior Frontal, a47r | Right | Frontoparietal | 0 |
| R_47l | Inferior Frontal, 47l | Right | Default | 0 |
| R_45 | Inferior Frontal, 45 | Right | Language | 0 |
| R_VIP | Superior Parietal, VIP | Right | Visual2 | 0 |
| R_LIPv | Superior Parietal, LIPv | Right | Visual2 | 0 |
| R_7PC | Superior Parietal, 7PC | Right | Somatomotor | 0 |
| R_A1 | Early Auditory, A1 | Right | Auditory | 0 |
| R_PIT | Ventral Stream Visual, PIT | Right | Visual2 | 0 |
| R_LO2 | MT+ Complex and Neighboring Visual Areas, LO2 | Right | Visual2 | 0 |
| R_LO1 | MT+ Complex and Neighboring Visual Areas, LO1 | Right | Visual2 | 0 |
| R_V3B | Dorsal Stream Visual, V3B | Right | Visual2 | 0 |
| R_FFC | Ventral Stream Visual, FFC | Right | Visual2 | 0 |
| R_IPS1 | Dorsal Stream Visual, IPS1 | Right | Visual2 | 0 |
| R_V7 | Dorsal Stream Visual, V7 | Right | Visual2 | 0 |
| R_POS2 | Posterior Cingulate, POS2 | Right | Frontoparietal | 0 |
| R_RSC | Posterior Cingulate, RSC | Right | Frontoparietal | 0 |
| R_V3A | Dorsal Stream Visual, V3A | Right | Visual2 | 0 |
| R_55b | Premotor, 55b | Right | Language | 0 |
| R_PEF | Premotor, PEF | Right | Cingulo-Opercular | 0 |
| R_FEF | Premotor, FEF | Right | Cingulo-Opercular | 0 |
| R_3b | Somatosensory and Motor, 3b | Right | Somatomotor | 0 |
| R_V8 | Ventral Stream Visual, V8 | Right | Visual2 | 0 |
| R_V4 | Early Visual, V4 | Right | Visual2 | 0 |
| R_V3 | Early Visual, V3 | Right | Visual2 | 0 |
| R_V2 | Early Visual, V2 | Right | Visual2 | 0 |
| R_V6 | Dorsal Stream Visual, V6 | Right | Visual2 | 0 |
| R_MT | MT+ Complex and Neighboring Visual Areas, MT | Right | Visual2 | 0 |
| R_PSL | Temporo-Parieto Occipital Junction, PSL | Right | Cingulo-Opercular | 0 |
| R_7PL | Superior Parietal, 7Pl | Right | Dorsal_Attention | 0 |
| R_SFL | Dorsolateral Prefrontal, SFL | Right | Language | 0 |
| R_7Am | Superior Parietal, 7Am | Right | Cingulo-Opercular | 0 |
| R_6ma | Paracentral Lobular and Mid Cingulate, 6ma | Right | Cingulo-Opercular | 0 |
| R_SCEF | Paracentral Lobular and Mid Cingulate, SCEF | Right | Cingulo-Opercular | 0 |
| R_7AL | Superior Parietal, 7AL | Right | Somatomotor | 0 |
| R_24dv | Paracentral Lobular and Mid Cingulate, 24dv | Right | Somatomotor | 0 |
| R_24dd | Paracentral Lobular and Mid Cingulate, 24dd | Right | Somatomotor | 0 |
| R_5L | Paracentral Lobular and Mid Cingulate, 5L | Right | Somatomotor | 0 |
| R_23c | Paracentral Lobular and Mid Cingulate, 23c | Right | Cingulo-Opercular | 0 |
| R_5mv | Paracentral Lobular and Mid Cingulate, 5mv | Right | Cingulo-Opercular | 0 |
| R_5m | Paracentral Lobular and Mid Cingulate, 5m | Right | Somatomotor | 0 |
| R_31pv | Posterior Cingulate, 31pv | Right | Default | 0 |
| R_d23ab | Posterior Cingulate, d23ab | Right | Default | 0 |
| R_v23ab | Posterior Cingulate, v23ab | Right | Default | 0 |
| R_23d | Posterior Cingulate, 23d | Right | Default | 0 |
| R_POS1 | Posterior Cingulate, POS1 | Right | Default | 0 |
| R_7m | Posterior Cingulate, 7m | Right | Default | 0 |
| R_7Pm | Superior Parietal, 7Pm | Right | Frontoparietal | 0 |
| R_STV | Temporo-Parieto-Occipital Junction, STV | Right | Posterior_Multimodal | 0 |
| R_PCV | Posterior Cingulate, PCV | Right | Posterior_Multimodal | 0 |
| R_i6-8 | Dorsolateral Prefrontal, i6-8 | Right | Frontoparietal | 0 |
| R_43 | Posterior Opercular, 43 | Right | Cingulo-Opercular | 0 |
| L_V3A | Dorsal Stream Visual, V3A | Left | Visual2 | 0 |
| R_LO3 | MT+ Complex and Neighboring Visual Areas, LO3 | Right | Visual2 | 0 |
| R_Ig | Insular and Frontal Opercular, Ig | Right | Somatomotor | 0 |
| R_PoI1 | Insular and Frontal Opercular, PoI1 | Right | Cingulo-Opercular | 0 |
| R_pOFC | Anterior Cingulate and Medial Prefrontal, pOFC | Right | Orbito-Affective | 0 |
| R_s32 | Anterior Cingulate and Medial Prefrontal, s32 | Right | Default | 0 |
| R_25 | Anterior Cingulate and Medial Prefrontal, 25 | Right | Default | 0 |
| R_VVC | Ventral Stream Visual, VVC | Right | Visual2 | 0 |
| R_31a | Posterior Cingulate, 31a | Right | Frontoparietal | 0 |
| R_31pd | Posterior Cingulate, 31pd | Right | Default | 0 |
| R_VMV2 | Ventral Stream Visual, VMV2 | Right | Visual2 | 0 |
| R_V3CD | MT+ Complex and Neighboring Visual Areas, V3CD | Right | Visual2 | 0 |
| R_OP4 | Posterior Opercular, OP4 | Right | Somatomotor | 0 |
| R_FST | MT+ Complex and Neighboring Visual Areas, FST | Right | Visual2 | 0 |
| R_V4t | MT+ Complex and Neighboring Visual Areas, V4t | Right | Visual2 | 0 |
| R_PHA2 | Medial Temporal, PHA2 | Right | Default | 0 |
| R_VMV3 | Ventral Stream Visual, VMV3 | Right | Visual2 | 0 |
| R_VMV1 | Ventral Stream Visual, VMV1 | Right | Visual2 | 0 |
| R_V6A | Dorsal Stream Visual, V6A | Right | Visual2 | 0 |
| R_PGs | Inferior Parietal, PGs | Right | Default | 0 |
| R_PGi | Inferior Parietal, PGi | Right | Default | 0 |
| R_PFm | Inferior Parietal, PFm | Right | Frontoparietal | 0 |
| R_FOP5 | Insular and Frontal Opercular, FOP5 | Right | Cingulo-Opercular | 0 |
| R_p10p | Orbital and Polar Frontal, p10p | Right | Frontoparietal | 0 |
| R_p47r | Inferior Frontal, p47r | Right | Frontoparietal | 0 |
| R_TGv | Lateral Temporal, TGv | Right | Language | 0 |
| L_55b | Premotor, 55b | Left | Language | 0 |
| L_PEF | Premotor, PEF | Left | Dorsal_Attention | 0 |
| R_MST | MT+ Complex and Neighboring Visual Areas, MST | Right | Visual2 | 0 |
| L_4 | Somatosensory and Motor, 4 | Left | Somatomotor | 0 |
| L_V8 | Ventral Stream Visual, V8 | Left | Visual2 | 0 |
| L_V4 | Early Visual, V4 | Left | Visual2 | 0 |
| L_V3 | Early Visual, V3 | Left | Visual2 | 0 |
| L_V2 | Early Visual, V2 | Left | Visual2 | 0 |
| L_V6 | Dorsal Stream Visual, V6 | Left | Visual2 | 0 |
| L_MST | MT+ Complex and Neighboring Visual Areas, MST | Left | Visual2 | 0 |
| L_V1 | Primary Visual, V1 | Left | Visual1 | 0 |
| R_p24 | Anterior Cingulate and Medial Prefrontal, p24 | Right | Cingulo-Opercular | 0 |
| R_a32pr | Anterior Cingulate and Medial Prefrontal, a32pr | Right | Cingulo-Opercular | 0 |
| R_PI | Insular and Frontal Opercular, PI | Right | Cingulo-Opercular | 0 |
| R_TE1m | Lateral Temporal, TE1m | Right | Frontoparietal | 0 |
| R_STSva | Auditory Association, STSva | Right | Default | 0 |
| R_A4 | Auditory Association, A4 | Right | Auditory | 0 |
| R_LBelt | Early Auditory, LBelt | Right | Auditory | 0 |
| R_MBelt | Early Auditory, MBelt | Right | Auditory | 0 |
| R_PF | Inferior Parietal, PF | Right | Cingulo-Opercular | 0 |
| R_PFop | Inferior Parietal, PFop | Right | Cingulo-Opercular | 0 |
| R_IP0 | Inferior Parietal, IP0 | Right | Dorsal_Attention | 0 |
| R_PeEc | Medial Temporal, PeEc | Right | Ventral_Multimodal | 0 |
| R_H | Medial Temporal, H | Right | Default | 0 |
| R_PreS | Medial Temporal, PreS | Right | Default | 0 |
| R_EC | Medial Temporal, EC | Right | Default | 0 |
| R_PFt | Inferior Parietal, PFt | Right | Dorsal_Attention | 0 |
| R_FOP2 | Insular and Frontal Opercular, FOP2 | Right | Somatomotor | 0 |
| R_FOP3 | Insular and Frontal Opercular, FOP3 | Right | Cingulo-Opercular | 0 |
| R_FOP1 | Posterior Opercular, FOP1 | Right | Cingulo-Opercular | 0 |
| R_AAIC | Insular and Frontal Opercular, AAIC | Right | Orbito-Affective | 0 |
| R_AVI | Insular and Frontal Opercular, AVI | Right | Frontoparietal | 0 |
| R_Pir | Insular and Frontal Opercular, Pir | Right | Orbito-Affective | 0 |
| R_MI | Insular and Frontal Opercular, MI | Right | Cingulo-Opercular | 0 |
| R_FOP4 | Insular and Frontal Opercular, FOP4 | Right | Cingulo-Opercular | 0 |
| R_TA2 | Auditory Association, TA2 | Right | Auditory | 0 |
| R_PoI2 | Insular and Frontal Opercular, PoI2 | Right | Cingulo-Opercular | 0 |
| R_PFcm | Early Auditory, PFcm | Right | Cingulo-Opercular | 0 |
| R_RI | Early Auditory, RI | Right | Somatomotor | 0 |
| R_52 | Early Auditory, 52 | Right | Auditory | 0 |
| R_OP2-3 | Posterior Opercular, OP2-3 | Right | Somatomotor | 0 |
| R_OP1 | Posterior Opercular, OP1 | Right | Somatomotor | 0 |
| R_ProS | Posterior Cingulate, ProS | Right | Visual1 | 0 |
| R_STGa | Auditory Association, STGa | Right | Language | 0 |
| R_IP1 | Inferior Parietal, IP1 | Right | Frontoparietal | 0 |
| R_PBelt | Early Auditory, PBelt | Right | Auditory | 0 |
| R_IP2 | Inferior Parietal, IP2 | Right | Frontoparietal | 0 |
| R_PGp | Inferior Parietal, PGp | Right | Dorsal_Attention | 0 |
| R_DVT | Posterior Cingulate, DVT | Right | Visual1 | 0 |
| R_TPOJ3 | Temporo-Parieto-Occipital Junction, TPOJ3 | Right | Posterior_Multimodal | 0 |
| R_TPOJ2 | Temporo-Parieto-Occipital Junction, TPOJ2 | Right | Posterior_Multimodal | 0 |
| R_TPOJ1 | Temporo-Parieto-Occipital Junction, TPOJ1 | Right | Language | 0 |
| R_PH | MT+ Complex and Neighboring Visual Areas, PH | Right | Visual2 | 0 |
| R_PHT | Lateral Temporal, PHT | Right | Dorsal_Attention | 0 |
| R_TE2p | Lateral Temporal, TE2p | Right | Dorsal_Attention | 0 |
| R_TE2a | Lateral Temporal, TE2a | Right | Default | 0 |
| R_TE1p | Lateral Temporal, TE1p | Right | Frontoparietal | 0 |
| R_TE1a | Lateral Temporal, TE1a | Right | Default | 0 |
| R_TGd | Lateral Temporal, TGd | Right | Default | 0 |
| R_STSvp | Auditory Association, STSvp | Right | Default | 0 |
| R_STSdp | Auditory Association, STSdp | Right | Language | 0 |
| R_STSda | Auditory Association, STSda | Right | Default | 0 |
| R_PHA3 | Medial Temporal, PHA3 | Right | Dorsal_Attention | 0 |
| R_PHA1 | Medial Temporal, PHA1 | Right | Default | 0 |
| R_A5 | Auditory Association, A5 | Right | Language | 0 |
| THALAMUS_RIGHT | Thalamus Right | Right | Subcortex | 0 |

***Table S6. Average feature importance of the top-performing non-stacked models with with Elastic Net, as indicated by Elastic Net coefficients for the HCP Young Adults dataset, predicting cognitive abilities: Working Memory Contrast.*** *The networks are ranked by the average amplitude.*

| **Network** | **Average Amplitude** | **SD Amplitude** | **Average Magnitude** | **SD Magnitude** | **Number of ROIs in Network** |
| --- | --- | --- | --- | --- | --- |
| **Posterior**  **Multimodal** | 0.017120 | 0.021203 | 0.017120 | 0.021203 | 7 |
| **Dorsal**  **Attention** | 0.008242 | 0.016623 | 0.012103 | 0.013936 | 23 |
| **Orbito-Affective** | 0.005750 | 0.014856 | 0.006266 | 0.014604 | 6 |
| **Frontoparietal** | 0.004750 | 0.014424 | 0.008096 | 0.012814 | 50 |
| **Ventral**  **Multimodal** | 0.003928 | 0.007857 | 0.003928 | 0.007857 | 4 |
| **Auditory** | 0.003019 | 0.007160 | 0.003760 | 0.006774 | 15 |
| **Subcortex** | 0.002688 | 0.008890 | 0.005102 | 0.007694 | 19 |
| **Language** | 0.001965 | 0.008123 | 0.003713 | 0.007457 | 23 |
| **Visual1** | 0.000013 | 0.013082 | 0.008475 | 0.009217 | 6 |
| **Visual2** | -0.000015 | 0.010176 | 0.004146 | 0.009276 | 54 |
| **Somatomotor** | -0.002406 | 0.007887 | 0.002975 | 0.007685 | 39 |
| **Default** | -0.003256 | 0.014204 | 0.006486 | 0.013034 | 77 |
| **Cingulo-Opercular** | -0.005016 | 0.013224 | 0.006542 | 0.012526 | 56 |

***Table S7. Average feature importance of the top-performing non-stacked models with with Elastic Net, as indicated by Elastic Net coefficients for the HCP Aging dataset, predicting cognitive abilities: Facename Encoding vs Distractor Task Contrast.*** *The networks are ranked by the average amplitude.*

| **Network** | **Average Amplitude** | **SD Amplitude** | **Average Magnitude** | **SD Magnitude** | **Number of ROIs in Network** |
| --- | --- | --- | --- | --- | --- |
| **Subcortex** | 0.011827 | 0.014116 | 0.015581 | 0.009522 | 19 |
| **Posterior**  **Multimodal** | 0.007449 | 0.016686 | 0.015482 | 0.007969 | 7 |
| **Dorsal**  **Attention** | 0.004975 | 0.011959 | 0.010149 | 0.007823 | 23 |
| **Visual1** | 0.004298 | 0.012815 | 0.010618 | 0.007147 | 6 |
| **Language** | 0.001100 | 0.011356 | 0.009755 | 0.005544 | 23 |
| **Orbito-Affective** | 0.001055 | 0.007184 | 0.005489 | 0.004097 | 6 |
| **Frontoparietal** | 0.000568 | 0.011527 | 0.009476 | 0.006448 | 50 |
| **Somatomotor** | -0.000321 | 0.016510 | 0.013234 | 0.009641 | 39 |
| **Default** | -0.000783 | 0.012169 | 0.009662 | 0.007356 | 77 |
| **Ventral**  **Multimodal** | -0.001050 | 0.013671 | 0.010389 | 0.006667 | 4 |
| **Cingulo-Opercular** | -0.001420 | 0.012573 | 0.010056 | 0.007561 | 56 |
| **Visual2** | -0.003188 | 0.016774 | 0.013629 | 0.010122 | 54 |
| **Auditory** | -0.011744 | 0.014127 | 0.015163 | 0.010052 | 15 |

***Table S8. Average feature importance of the top-performing non-stacked models with with Elastic Net, as indicated by Elastic Net coefficients for the Dunedin Study dataset, predicting cognitive abilities of 45years old: Facename Encoding vs Distractor Task Contrast.*** *The networks are ranked by the average amplitude.*

| **Network** | **Average Amplitude** | **SD Amplitude** | **Average Magnitude** | **SD Magnitude** | **Number of ROIs in Network** |
| --- | --- | --- | --- | --- | --- |
| **Dorsal**  **Attention** | 0.006244 | 0.011393 | 0.009894 | 0.008259 | 23 |
| **Visual1** | 0.004003 | 0.010554 | 0.009511 | 0.004696 | 6 |
| **Frontoparietal** | 0.003882 | 0.010994 | 0.009330 | 0.006887 | 50 |
| **Orbito-Affective** | 0.003838 | 0.007052 | 0.006305 | 0.004440 | 6 |
| **Posterior**  **Multimodal** | 0.002295 | 0.008981 | 0.007351 | 0.004875 | 7 |
| **Subcortex** | 0.002070 | 0.015654 | 0.012443 | 0.009281 | 19 |
| **Cingulo-Opercular** | 0.001577 | 0.009536 | 0.007451 | 0.006077 | 56 |
| **Language** | 0.001283 | 0.011044 | 0.009464 | 0.005482 | 23 |
| **Auditory** | 0.000851 | 0.011185 | 0.008318 | 0.007194 | 15 |
| **Visual2** | -0.000393 | 0.012931 | 0.010609 | 0.007259 | 54 |
| **Ventral**  **Multimodal** | -0.003259 | 0.006830 | 0.005075 | 0.005145 | 4 |
| **Somatomotor** | -0.004184 | 0.008661 | 0.007620 | 0.005778 | 39 |
| **Default** | -0.004205 | 0.007956 | 0.007522 | 0.004889 | 77 |

***Table S9. Average feature importance of the top-performing non-stacked models with with Elastic Net, as indicated by Elastic Net coefficients for the Dunedin Study dataset, predicting cognitive abilities of 7, 9, 11 years old: Facename Encoding vs Distractor Task Contrast.*** *The networks are ranked by the average amplitude.*

| **Network** | **Average Amplitude** | **SD Amplitude** | **Average Magnitude** | **SD Magnitude** | **Number of ROIs in Network** |
| --- | --- | --- | --- | --- | --- |
| **Visual1** | 0.004334 | 0.008775 | 0.008614 | 0.003241 | 6 |
| **Orbito-Affective** | 0.003967 | 0.015481 | 0.012804 | 0.007863 | 6 |
| **Language** | 0.003354 | 0.018817 | 0.015798 | 0.010242 | 23 |
| **Dorsal**  **Attention** | 0.003097 | 0.016435 | 0.012136 | 0.011232 | 23 |
| **Ventral**  **Multimodal** | 0.002503 | 0.013561 | 0.008698 | 0.009560 | 4 |
| **Frontoparietal** | 0.002390 | 0.017605 | 0.014725 | 0.009723 | 50 |
| **Subcortex** | 0.001371 | 0.019013 | 0.015418 | 0.010610 | 19 |
| **Visual2** | 0.001165 | 0.019374 | 0.016254 | 0.010371 | 54 |
| **Cingulo-Opercular** | -0.000224 | 0.013764 | 0.010888 | 0.008294 | 56 |
| **Somatomotor** | -0.001204 | 0.015127 | 0.011132 | 0.010154 | 39 |
| **Auditory** | -0.001840 | 0.018241 | 0.014450 | 0.010613 | 15 |
| **Default** | -0.003007 | 0.013203 | 0.010780 | 0.008108 | 77 |
| **Posterior**  **Multimodal** | -0.007922 | 0.018548 | 0.016367 | 0.010234 | 7 |

***Table S10. Average feature importance of the top-performing non-stacked models with with Elastic Net, as indicated by Elastic Net coefficients for the Dunedin Study dataset, predicting residual scores for cognitive abilities: Facename Encoding vs Distractor Task Contrast.*** *The networks are ranked by the average amplitude.*

| **Network** | **Average Amplitude** | **SD Amplitude** | **Average Magnitude** | **SD Magnitude** | **Number of ROIs in Network** |
| --- | --- | --- | --- | --- | --- |
| **Dorsal Attention** | 0.007034 | 0.018766 | 0.007034 | 0.018766 | 23 |
| **Subcortex** | 0.001268 | 0.012153 | 0.004805 | 0.011181 | 19 |
| **Frontoparietal** | 0.000881 | 0.005211 | 0.000881 | 0.005211 | 50 |
| **Auditory** | 0.000000 | 0.000000 | 0.000000 | 0.000000 | 15 |
| **Cingulo-Opercular** | 0.000000 | 0.000000 | 0.000000 | 0.000000 | 56 |
| **Language** | 0.000000 | 0.000000 | 0.000000 | 0.000000 | 23 |
| **Orbito-Affective** | 0.000000 | 0.000000 | 0.000000 | 0.000000 | 6 |
| **Posterior Multimodal** | 0.000000 | 0.000000 | 0.000000 | 0.000000 | 7 |
| **Visual1** | 0.000000 | 0.000000 | 0.000000 | 0.000000 | 6 |
| **Default** | -0.000453 | 0.002637 | 0.000453 | 0.002637 | 77 |
| **Visual2** | -0.000871 | 0.004512 | 0.000957 | 0.004494 | 54 |
| **Somatomotor** | -0.001692 | 0.005040 | 0.001692 | 0.005040 | 39 |
| **Ventral Multimodal** | -0.009811 | 0.011371 | 0.009811 | 0.011371 | 4 |
